# A multifaceted cellular damage repair and prevention pathway promotes high level tolerance to β-lactam antibiotics

**DOI:** 10.1101/777375

**Authors:** Jung-Ho Shin, Donghui Choe, Brett Ransegnola, Hye-Rim Hong, Ikenna Onyekwere, Trevor Cross, Qiaojuan Shi, Byung-Kwan Cho, Lars F. Westblade, Ilana L. Brito, Tobias Dörr

## Abstract

Bactericidal antibiotics are powerful agents due to their ability to convert essential bacterial functions into lethal processes. However, many important bacterial pathogens are remarkably tolerant against bactericidal antibiotics due to inducible damage repair responses. The cell wall damage response two-component system VxrAB of the gastrointestinal pathogen *Vibrio cholerae* promotes high-level β-lactam tolerance and controls a gene network encoding highly diverse functions, including negative control over multiple iron uptake systems. How this system contributes to tolerance is poorly understood. Here, we show that β-lactam antibiotics cause an increase in intracellular free iron levels and collateral oxidative damage, which is exacerbated in the Δ*vxrAB* mutant. Mutating major iron uptake systems drastically increased Δ*vxrAB* tolerance to β-lactams. We propose that VxrAB reduces antibiotic-induced toxic iron and concomitant metabolic perturbations by downregulating iron uptake transporters and show that iron sequestration enhances tolerance against β-lactam therapy in a mouse model of cholera infection. Our results suggest that a microorganism’s ability to counteract diverse antibiotic-induced stresses promotes high-level antibiotic tolerance, and highlights the complex secondary responses elicited by antibiotics.

## Introduction

The discovery and development of antibiotics is one of the most important biomedical breakthroughs in human history. The introduction of agents that selectively kill bacteria (bactericidal antibiotics) or inhibit their growth (bacteriostatic antibiotics) has directly saved countless lives by enabling treatment of once common causes of death (*e.g.*, pneumonia and sepsis) and indirectly facilitated other medical breakthroughs, including transplantation medicine and chemotherapy^1,2^. Unfortunately, these significant advances in medicine are threatened due to the rapid increase in the occurrence of antibiotic treatment failure. We urgently need to develop new therapeutics and improve the effectiveness of existing ones; this requires a precise understanding of antibiotic mechanisms of action, and especially of bacterial strategies to avoid their noxious effects. In principle, bacteria resist antibiotics by either developing the ability to grow in their presence (antibiotic resistance, ABR) or to simply stay alive in their presence for extended time periods (antibiotic tolerance/persistence)^3–8^. While the mechanisms and consequences of ABR are relatively well-established, antibiotic tolerance remains poorly understood, limiting our ability to effectively treat bacterial infections.

The β-lactam antibiotics (penicillins, cephalosporins, carbapenems, cephamycins and monobactams) are highly potent bactericidal agents. Typically, their lethal action results from their ability to simultaneously inhibit multiple targets (i.e., the transpeptidase domain of multiple penicillin-binding proteins [PBPs]), which ultimately causes bacterial cells to deplete essential cell wall precursors and self-destruct through the activity of endogenous, cell wall lytic enzymes (‘autolysins’; endopeptidases, amidases and lytic transglycosylases) ^9–12^. However, we and others have recently shown that many clinically significant Gram-negative pathogens are remarkably tolerant to β-lactams. The agent of cholera, *Vibrio cholerae*, the opportunistic pathogen *Pseudomonas aeruginosa* and members of the *Enterobacterales* all survive treatment with β-lactams (including the potent so-called “last resort” agent meropenem [a carbapenem]) by forming non-dividing, cell wall deficient spheroplasts ^11,13,14^. Upon removal of the β-lactam, spheroplasts rapidly revert to wild-type shape and growth, providing a possible explanation for treatment failure in the clinic. Spheroplasts are reminiscent of bacteria in the so-called “L-form” state ^15,16^ with the crucial distinction that spheroplasts, unlike L-forms, do not divide in the presence of antibiotics.

The extent of antibiotic tolerance is a reflection of the cell’s ability to mount a stress response to counteract primary damage resulting from antibiotic-target interactions ^17,18^. Numerous bacteria encode cell envelope damage response systems (alternative sigma factors and two-component response regulators) that sense damage to the cell wall or membranes and consequently upregulate complex regulons aimed at damage repair ^19–25^. In *V. cholerae*, high-level β-lactam tolerance depends on the cell wall damage sensing two-component system VxrAB (also known as WigKR)^17^, which is broadly conserved among *Vibrionaceae*. The vast VxrAB regulon is induced by cell wall-acting antibiotics and includes cell wall synthesis functions and those involved in biofilm formation and type VI secretion, *inter alia* ^17,26,27^. How VxrAB promotes high level tolerance against bactericidal β-lactams is unknown, but presumably relies on repair of cellular damage inflicted by β-lactam antibiotics, *i.e.*, primarily cell envelope damage. However, in addition to their primary mechanism of action, bactericidal antibiotics have also been proposed to generate toxic reactive oxygen species (ROS) as a byproduct, which are thought to enhance cell death ^28–31^. This idea has been highly controversial and the resulting debate is ongoing ^32–37^.

Here, we present evidence supporting a multifactorial role for VxrAB in promoting high-level β-lactam tolerance. We have found that the ability to upregulate cell wall synthesis only accounts for part of the high-level tolerance observed in wild-type *V. cholerae.* In addition, we propose that VxrAB’s role in promoting tolerance is to downregulate iron acquisition systems which are inappropriately induced due to the accumulation of ROS that are produced upon exposure to penicillin. Our data highlight the pleiotropic nature of antibiotic mechanism of action and provide a blueprint for the mechanistic underpinnings of the high-level β-lactam tolerance often observed in significant Gram-negative pathogens.

## Results

### Spheroplast-mediated β-lactam tolerance is widespread in human pathogens

Recently, we and others have reported that clinically significant bacterial pathogens exhibit high tolerance to β-lactam antibiotics via formation of cell wall deficient spheroplasts when exposed to antibiotics in various growth media, including human serum^11,13,14^. Here, we asked if spheroplast formation could also be observed in the standard bacterial growth medium LB (Lysogeny Broth). We tested a well-characterized^14^ panel of clinical isolates of *Enterobacterales* as well as known spheroplast-formers, *Pseudomonas aeruginosa* and *V. cholerae*, for their ability to survive exposure to the carbapenem antibiotic meropenem. Meropenem is not susceptible to hydrolysis by the AmpC β-lactamase which some members of the *Enterobacterales* encode^38^ and was thus appropriate for these experiments. Tolerance levels for almost all isolates (with the exception of *E. coli* lab strain MG1655), were very high (comparable to or higher than our previous observations in human serum and Brain-Heart Infusion broth^14^) (**Fig. 1A**). Crucially, tolerance in all cases coincided with the formation of membranous (membranes visualized by FM4-64 stain, **Fig. 1B**) spheroplasts after 3h of incubation. Thus, β-lactam tolerance phenotypes are recapitulated in LB, underscoring the usefulness of this growth medium to probe β-lactam tolerance.

**Figure 1.**
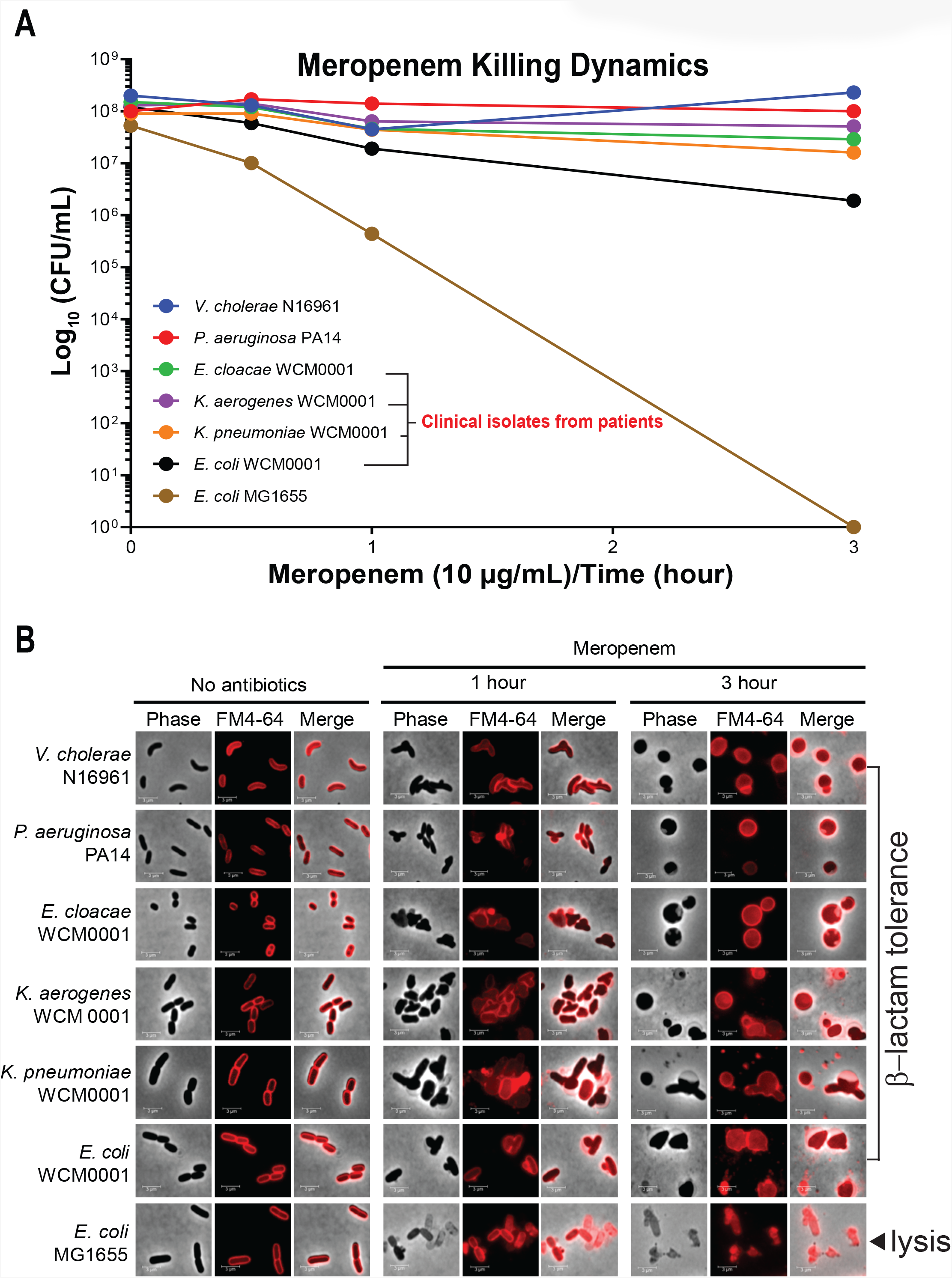
Spheroplast formation in diverse bacterial pathogens. **(A)** Time-dependent killing of a panel of bacterial isolates in the presence of meropenem (MEM10, 10 µg/ml). Cells were grown overnight and sub-cultured for 30 minutes at 37°C before being exposed to meropenem at 10 µg/mL and moved to a 37°C stationary incubator. Samples were taken at each indicated time point and spot-titered onto LB medium to quantify colony forming units (CFU/mL) throughout meropenem treatment. **(B)** images of cells corresponding to the time points in **(A)** were taken after labeling with membrane stain FM4-64. *E. cloacae, Enterobacter cloacae; K. aerogenes, Klebsiella aerogenes; K. pneumoniae, Klebsiella pneumoniae; E. coli, Escherichia coli*

### ChIP-Seq and RNA-Seq reveal a complex cell wall damage response

We previously elucidated the transcriptional network controlled by VxrAB upon overexpression of a phosphomimetic version of VxrB (VxrB^D78E^). Here, we directed our attention to genes directly controlled by the VxrAB response. To this end, we performed a ChIP-Seq experiment using a strain with functional, polyhistidine-tagged, chromosomally encoded VxrB assayed during penicillin (PenG) treatment. After 180 min of PenG exposure (100 µg/ml, 10 × MIC; same conditions used in our previous study^17^), 424 putative VxrB binding events were identified by comparison with a similarly treated control strain expressing wild-type, untagged VxrB (**Fig. 2A**, see **Table S3** for complete list). We then validated the ChIP-Seq results via electrophoretic mobility shift assays (EMSA) of the 10 DNAs from the strongest VxrB binding events (**Fig. S1**) and, for a subset, differential regulation via S1 nuclease assays of PenG-treated cells. We then integrated this ChIP-Seq dataset with our re-analyzed (with more stringent parameters, see Methods for details) previous RNA-Seq dataset of overexpressed VxrB^D78E^ ^17^ (**Fig. 2B**, see **Table S4** and **S5** for a complete list) to score VxrB binding events that coincided with transcriptional up- or downregulation of the associated gene(s) (**Table S5**). Our analysis resulted in the identification of 102 genes/operons that contained a putative VxrB binding site and exhibited significantly increased, VxrB-dependent expression levels, and 38 genes/operons that were scored as candidates for direct binding and downregulation (**Fig. 2C**). Preliminary functional assignments using the KEGG database ^39^ revealed that the VxrB regulon controlled a large number (76.4% 107/140) of genes of unknown function. Among the annotated upregulated genes, many coded for functions related to cell envelope biology (ABC transporters, peptidoglycan synthesis, and membrane synthesis) (**Fig. S2**). Among the downregulated genes, we observed genes encoding proteins involved in iron uptake systems, flagella biosynthesis, carbon metabolism, chemotaxis, cell wall lytic enzymes and lipopolysaccharide biosynthesis (**Fig. S2**). Importantly, genes encoding cell wall synthesis functions [*e.g.*, class A PBP1a (aPBP1a)], whose Vxr-dependent transcriptional regulation we have extensively validated^17^, exhibited strong VxrB binding sites (**Fig. 2D**) and differential, Vxr-dependent regulation in S1 nuclease assays, increasing our confidence in the robustness of the ChIP-Seq dataset. Both ChIP-Seq and RNA-Seq also corroborated autoregulation of the VxrAB operon (**Fig. 2B, Table S1**). We noted that while many of the putative Vxr-controlled genes exhibited VxrB binding in addition to differential regulation by phosphomimetic VxrB and VxrAB-dependent regulation upon exposure to PenG (**Fig. 2D**, *pbp1a* and *zapB*), some genes were apparently indirectly regulated as they clearly exhibited transcriptional downregulation (both upon overexpression of VxrB^D78E^ and exposure to PenG) without a clear signal in our VxrB ChIP-Seq experiment (**Fig. 2D, *flaA***). We speculate that this may reflect indirect regulation as a consequence of large scale metabolic alterations that might be expected from induction of the VxrAB regulon. Thus, the VxrAB cell wall stress response controls a gene network composed of diverse functions with an apparent enrichment for functions required for cell envelope homeostasis and iron acquisition.

**Figure 2.**
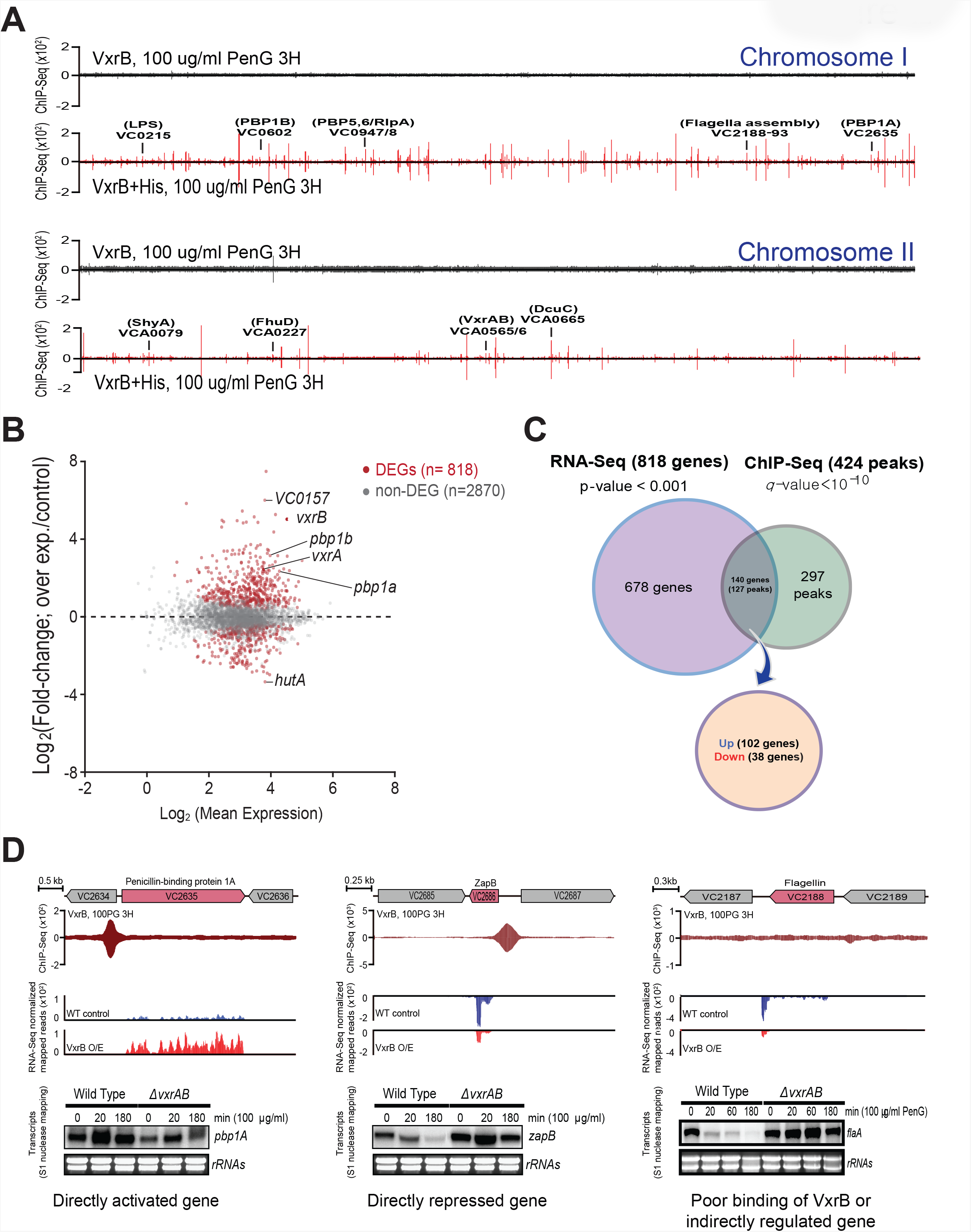
ChIP-Seq and RNA-Seq analysis reveals a complex VxrB regulon. **(A)** Overview of VxrB-binding profiles across the V. cholerae genome after β-lactam exposure (PenG 100μg/ml [10× MIC] for 3 hours). Red peaks indicate VxrB binding sites above the significance cutoff (q-value<10^−10^; MACS). **(B)** RNA-Seq MA plot is presented as fold change between VxrB^D78E^ overexpressed sample and control sample. Red dots indicate differentially expressed genes after VxrB^D78E^ overexpression (P-value < 0.001). Representative VxrB target genes are marked on both chromosomes and on the MA plot. **(C)** Venn diagram depicting overlap between RNA-seq and ChIP-Seq datasets **(D)** Regulatory modes of representative VxrB-controlled genes in response to PenG. Upper plot shows VxrB binding patterns (assessed via ChIP-Seq), lower panel shows transcriptional response upon overexpression of VxrB ^D78E^ (assessed via RNA-Seq). RNA expression was confirmed by S1 nuclease mapping assay in wild type and ΔvxrAB mutant strains after exposure to PenG (100 µg/ml for 3 hours).

### DNA footprinting analysis identifies putative VxrB binding sites

Based on an alignment of putative VxrB binding sites, we used the MEME suite ^40^ to identify the canonical VxrB consensus binding site (“Vxr box”). We based our analysis on the 102 upregulated and the 38 downregulated genes separately, reasoning that binding site variations may account for up-*versus* downregulation. This resulted in the putative VxrB binding motif TTGACAAAA-N2-TTGAC (N = any nucleotide) for the upregulated genes and CGCTCATTTTTTAACCAAGT for the downregulated genes. However, these putative binding motifs were poorly conserved across all Vxr-controlled genes. We hypothesize that this may be reflective of different binding characteristics of the phosphorylated vs. non-phosphorylated forms of VxrB; both forms might be expected to occur after PenG treatment and could bind to different motifs. To further characterize the VxrB binding site we conducted DNase I footprinting assays using purified VxrB and promoter regions of putative Vxr-controlled genes (**Fig. 3AB, Fig. S3**). We again chose genes with the strongest VxrB binding as indicated in our EMSA and ChIP-Seq experiments. We identified 10 promoters where VxrB protected a ~ 50 bp sequence from DNase I digestion at low µM concentrations (**Fig. 3AB, Fig. S3**), again corroborating direct binding to these promoters. However, apparent affinity varied between promoters, with the promoters of *pbp1a*, encoding aPBP1a, *mgtE*, encoding a magnesium transporter and *murJ*, encoding the lipid II flippase exhibiting the highest degree of protection (**Fig. 3AB, Fig. S3)**. Surprisingly, we were again unable to construct a well-conserved binding site for VxrB based on this subset of promoters, despite three independent assays unequivocally demonstrating binding. This apparent binding promiscuity suggests that VxrB might have the capacity to bind to a wide range of promoter sequences *in vivo* and/or that it requires as yet unidentified accessory factors to modulate binding.

**Figure 3.**
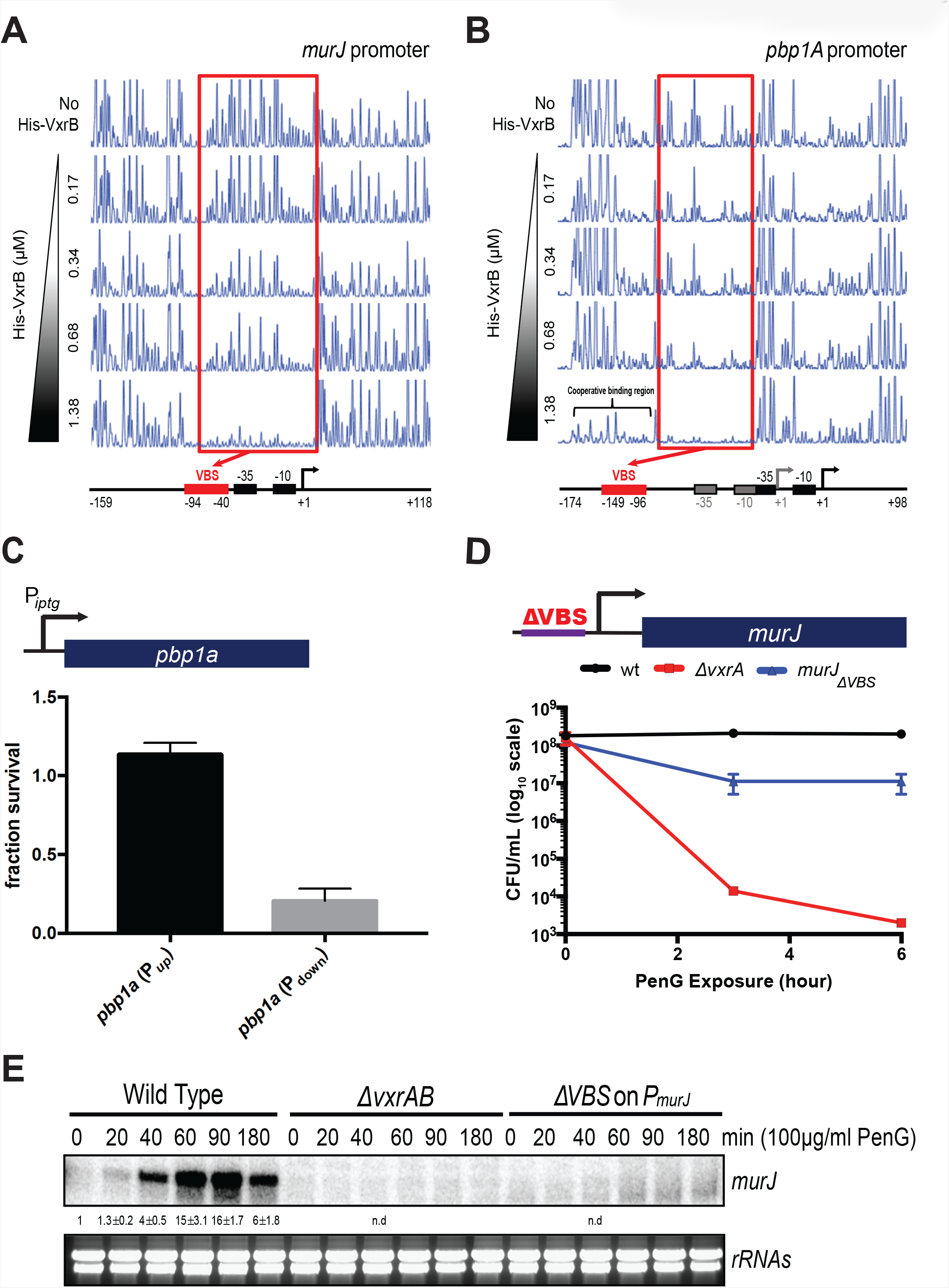
Peptidoglycan synthesis control only partially explains *Δvxr*AB lethality. **(A and B)** DNase I footprinting results for VxrB interaction with promoters controlling cell wall synthesis functions. Red boxes indicate core DNA regions protected by purified VxrB protein (final concentration 0.17 - 1.38 μM) on the *murJ* (A) or *pbp1A* (B) promoter. Identified transcriptional start site (TSS; +1) and promoter elements (−10 and −35) were derived from published datasets^89,90^, and are presented with the VxrB binding site. VBS, putative **V**xrB **B**inding **S**equence. **(C-E)** Reduction of cell wall synthesis gene expression causes a tolerance defect. **(C)** Survival of a Δ*pbp1b* Δ*pbp1a* P_*iptg*_:*pbp1a* strain after 6 hours of PenG exposure. P_up_ indicates conditions of high IPTG concentrations (100 µM), P_down_ low IPTG concentration (2 µM). Survival fraction is cfu/ml of PenG-treated cultures normalized to untreated cultures grown in and plated on medium containing the same IPTG concentration. (**D)** Time-dependent killing experiment in the presence of PenG (100 µg/ml, 10 × MIC). All data are means (+/− standard error) of 3 independent biological replicates. **(E)** Deleting the putative VxrB binding site from the *murJ* promoter abrogates induction of *murJ* by PenG exposure. Transcript levels were measured by S1 nuclease mapping. Numbers represent induction as measured by band intensity, normalized to pre-exposure WT. n.d., not determined.

### Control of cell wall synthesis only partially explains VxrAB’s requirement for β-lactam tolerance

To elucidate the mechanism of β-lactam tolerance in *V. cholerae*, we sought to dissect the individual contributions of VxrB-regulated genes to resistance to killing. Since the primary mechanism of action of β-lactam antibiotics is inhibition of cell wall synthesis, we first investigated the role of Vxr-induced PG biogenesis in antibiotic tolerance. We took two approaches to eliminate VxrB’s positive control over PG synthesis. For aPBP1a, we engineered a strain with a deletion in the second, functionally semi-redundant ^41,42^ aPBP1b, which also contains an IPTG-inducible PBP1a as its sole copy. As expected, growth of this strain was IPTG concentration-dependent (**Fig. S4**). We then grew this strain either in high concentrations of IPTG (high PBP1a levels) or very low concentrations that markedly reduced growth rate (**Fig. S4**). Addition of PenG to low aPBP concentration cultures resulted in only a modest (5 – 10-fold) decrease in viability (**Fig. 3C**). This observation suggests that upregulation of PBP1a is indeed critical for survival in the presence of PenG, but cannot account entirely for the steep, ~10,000-fold decrease in post-antibiotic plating efficiency (**Fig. 3D** and ^17^) of the Δ*vxrAB* mutant.

To corroborate these results, we also created a mutant with a partial deletion in the putative VxrB binding site of the *murJ* gene as determined from our DNA footprinting experiment. MurJ-mediated PG precursor translocation is a critical bottleneck for cell wall synthesis and upstream of all periplasmic cell wall assembly functions ^43,44^. We thus expected that introducing a mutation that removes VxrB’s positive control over this step would allow us to gauge the significance of VxrB control over cell wall synthesis as a whole. S1 nuclease assays confirmed that in the VxrB binding site deletion strain, *murJ* expression was not significantly induced by PenG, similar to a Δ*vxrAB* strain (**Fig. 3E**). Similar to our observations with low PBP1a expression, the Vxr-independent *murJ* strain exhibited 10-fold loss in viability upon exposure to PenG (**Fig. 3D**). We conclude that the ability to upregulate cell wall synthesis functions via VxrAB accounts for only some of its ability to promote tolerance.

### The VxrB regulon controls iron acquisition systems

Based on our previous RNA-Seq data, VxrB downregulates genes involved in iron acquisition ^17^. Indeed, examination of the RNA-Seq/ChIP-Seq datasets revealed a high overlap between genes regulated by VxrB with those regulated by the iron starvation response repressor Fur ^45^. 47 out of the putative 63 Fur-regulated genes were also downregulated by VxrB and 20 out of these 63 genes contained both a putative VxrB and a Fur binding site (**Fig. 4AB**). Thus, the VxrB system appears to control multiple functions required for iron homeostasis.

**Figure 4.**
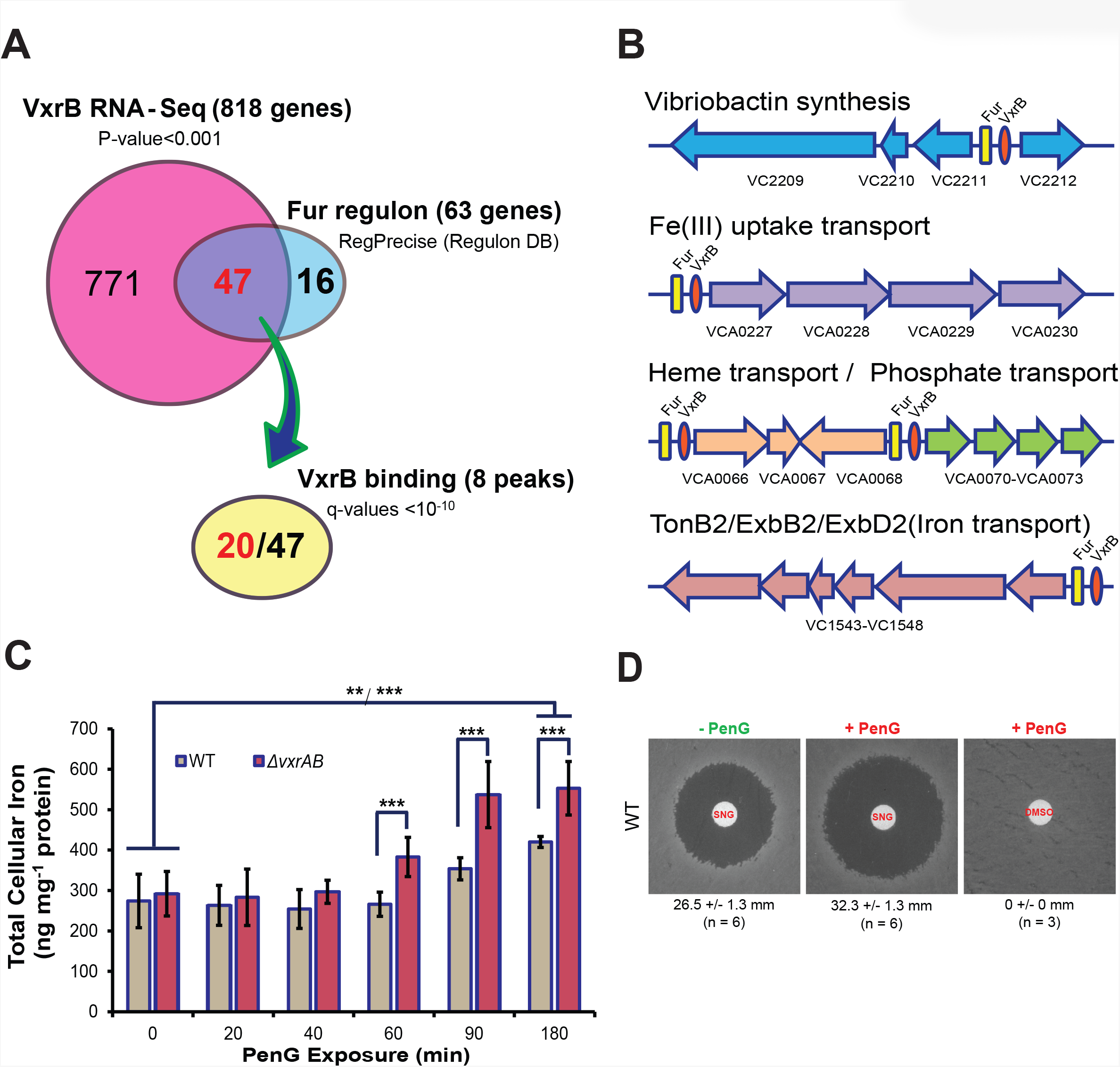
VxrAB controls genes involved in iron homeostasis. **(A)** Venn diagram showing overlap between the VxrB regulon and known *V. cholerae* Fur-controlled genes. (as per RegPrecise3.0 DB, http://regprecise.lbl.gov/RegPrecise/). 20 out of 47 Fur-regulated genes also have a VxrB binding site and are differentially regulated by VxrB^D78E^. (**B)** Genomic organization of iron uptake systems and their putative Fur/VxrB binding sites. **(C)** Time-course of iron accumulation in PenG-treated WT and Δ*vxrAB* cells as measured by ICP-MS. Data are average of 3 independent biological replicates, error bars represent standard deviation. **P=0.001; ***P<0.0003 (paired t-test). **(D)** Streptonigrin-sensitivity of PenG-treated cells. Cultures were grown to a density of ~2 × 10^8^ cfu/ml. Before (“-PenG”) and after addition of PenG (100 µg/ml, 10× MIC), ~ 5 × 10^7^ cells were spread on an LB agar plate. In the PenG-treated samples (“+ PenG”), beta lactamase solution was also added to remove the antibiotic. Streptonigrin (5 µL of 5 mg/ml, in DMSO) was then added on a filter disk, followed by incubation for 24 h at 37°C. Values represent average diameter of clearance zones (+/− standard deviation) of six biological replicates. SNG, streptonigrin.

Given VxrAB’s apparent role in downregulating iron acquisition in response to cell wall damage, we expected that surplus iron accumulates in a PenG-treated Δ*vxrAB* mutant and that this might be detrimental to cells under these conditions. To test this, we measured intracellular iron levels using ICP-MS during treatment with PenG. Before treatment, both the wild-type and Δ*vxrAB* mutant strains contained similar iron levels at ~280 ng/mg protein (**Fig. 4C)**. After 1 h of PenG exposure, wild-type iron levels remained relatively constant while the iron content of the Δ*vxrAB* strain started to increase significantly by ~60% compared to baseline. Iron levels increased slightly in the wild-type strain between 90 min and 3 h, while levels in the Δ*vxrAB* mutant remained significantly higher. Consistent with an increase in free intracellular iron, the wild-type strain also became more sensitive to the iron-dependent quinone antibiotic streptonigrin ^46,47^ upon treatment with PenG (**Fig. 4D).** Thus, PenG exposure causes enhanced iron influx.

### Penicillin exposure induces the Fur regulon

Why do PenG-treated Δ*vxrAB* cells increase their iron load? Exposure to β-lactam antibiotics has been shown to result in the accumulation of H_2_O_2_ in other bacteria ^31,48,49^ and we hypothesized that hydrogen peroxide might directly activate Fur via oxidation of its iron cofactor. Derepression of the Fur regulon in turn would result in the upregulation of iron acquisition systems. We first confirmed that PenG exposure resulted in H_2_O_2_ production in *V. cholerae* as well. We used a highly sensitive, fluorimetric assay (Amplex Red) to measure intracellular hydrogen peroxide concentrations. Upon exposure to PenG, both the wild-type strain and Δ*vxrAB* mutant experienced a transient increase in hydrogen peroxide concentrations, with a peak between 20 and 60 minutes, followed by a decrease back to baseline. However, the hydrogen peroxide concentration in the *vxrAB* mutant was significantly elevated and at peak (20 - 60 min) accumulated more than double the concentration measured in wild-type cells (**Fig. 5A**). As expected, and consistent with the accurate ability of Amplex Red to report hydrogen peroxide, a Δ*oxyR1* Δ*katB* Δ*katG* triple mutant, which lacks the ability to elaborate the primary hydrogen peroxide detoxification systems, exhibited both higher baseline and PenG-induced levels of hydrogen peroxide than the wild-type and Δ*vxrAB* mutant; additionally, external spiking of catalase into our assay reactions completely abolished the signal (**Fig. 5A**). We also measured hydrogen peroxide in the supernatant of PenG-treated cells, since hydrogen peroxide might diffuse through membranes to some degree. We did not, however, note a meaningful accumulation of peroxide in the supernatant of any of our mutants (**Fig. S5A**), demonstrating that at least for the duration of our measurements, there was no significant diffusion out of the cell.

**Figure 5.**
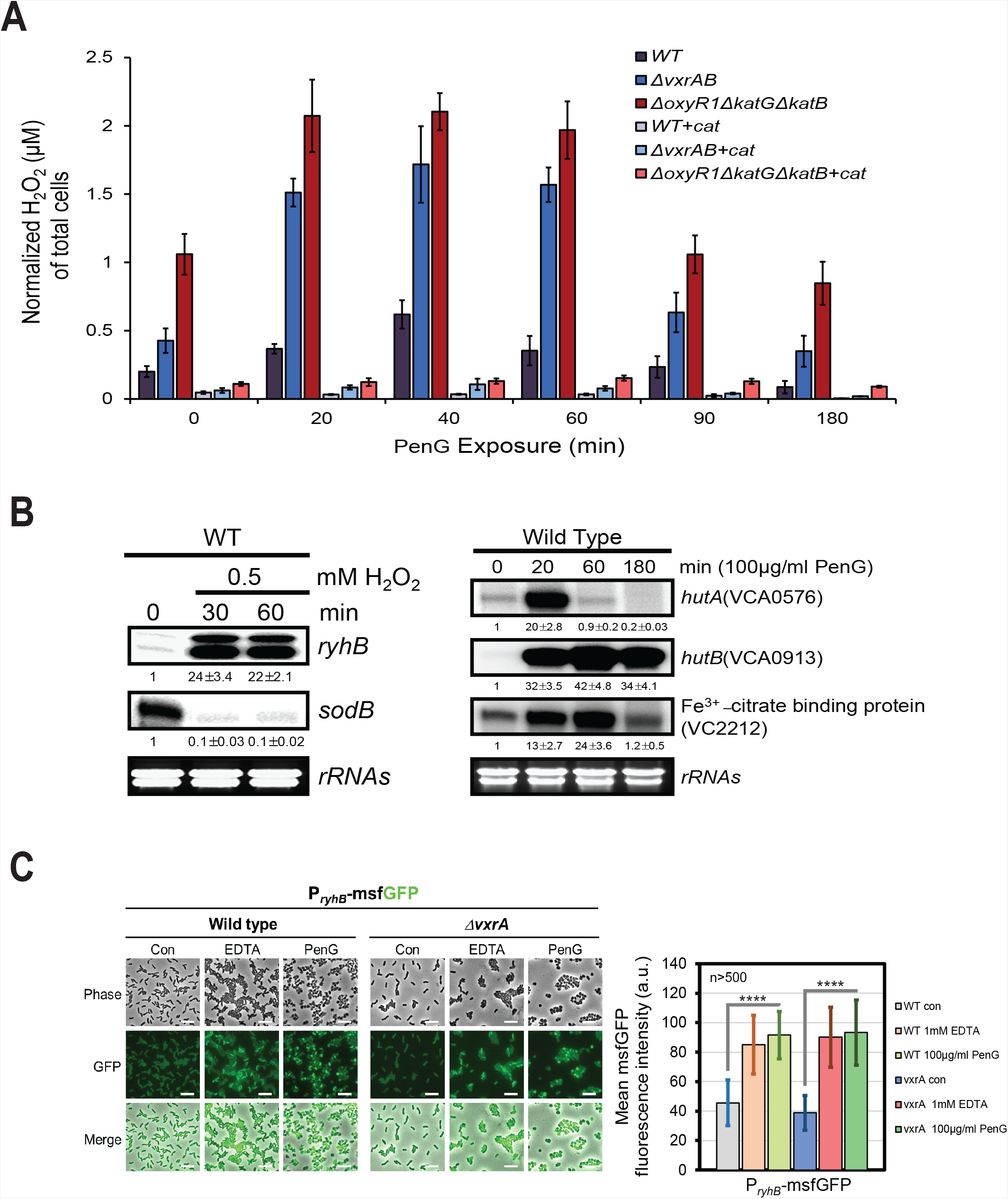
PenG induces hydrogen peroxide production and the Fur iron starvation response. **(A)** Total intracellular hydrogen peroxide (H_2_O_2_) levels were measured using Amplex™ Red Hydrogen Peroxide/Peroxidase Assay Kit and normalized to total cellular protein level (measured by Qubit™). Data are average values from three independent experiments, error bars represent standard deviation. Cat, catalase (1μg/ml). **(B)**. To test PenG-mediated induction of Fur-regulated genes, S1 nuclease mapping assays were performed on the small RNA *ryhB*, the RyhB target *sodB*, *hutA/B*, coding for heme transport systems and *vc2212*, which encodes a component of a ferric citrate uptake system). Cells were grown to mid-exponential phase followed by exposure to H_2_O_2_ or PenG for the indicated duration. Numbers represent expression levels normalized to untreated control (averages from three independent replicates). **(C)** To confirm the induction of *ryhB* transcript by PenG treatment, the native *ryhB* promoter was fused with *msfGFP* and introduced into WT and Δ*vxrA* mutant cells. Each strain was cultured until mid-exponential phase, when 1mM EDTA or 100μg/ml PenG were added for the indicated duration. Scale bar (5 µM). Quantified GFP fluorescence intensities are presented with average values from 3 independent biological replicates (cell, n>500), error bars represent standard deviation. **** P<0.0001 (paired t-test).

Next, we tested whether the observed accumulation of peroxide resulted in upregulation of the *fur* regulon. We used S1 nuclease mapping to monitor transcript levels of the canonical Fur regulon readout *ryhB*^50^, a small RNA ^51^. Consistent with our line of thinking, *ryhB* was highly induced by addition of external hydrogen peroxide **(Fig. 5B).** In *E. coli, sodB* transcript levels are negatively controlled by *ryhB*, and consistent with a similar regulatory mechanism in *V. cholerae, ryhB* induction (**Fig. 5B**) coincided with a disappearance of the *sodB* transcript during treatment with hydrogen peroxide (**Fig. 5B**). Consistent with a regulation via iron starvation sensing, the *ryhB* transcript also increased dramatically upon exposure of cells to the cation chelator EDTA (**Fig. S5B**). Upon PenG exposure, *ryhB* transcript was reproducibly induced ~2-fold in both the wild-type and Δ*vxrAB* strains after 20 min, followed by a return to baseline. Other Fur-regulated genes (*hutA, hutB* and *vc2212*), also exhibited a marked, and in some cases transient, upregulation upon PenG exposure (**Fig. 5B**). After prolonged PenG-treatment (180 min), *ryhB* remained uninduced in the wild-type, but increased dramatically in the Δ*vxrAB* mutant (**Fig. S5B**). Consistent with these transcriptional changes, a P_*ryhB*_-*msfGFP* transcriptional reporter construct also increased in fluorescence after exposure to PenG in both wild-type and Δ*vxrAB* strains (**Fig. 5C**), to similar degree as in the positive control, the metal chelator EDTA. These results suggest that the Fur regulon is indeed induced by exposure to PenG. Interestingly, we noticed in our ICP-MS dataset that concomitantly with iron uptake, manganese concentrations also increased in PenG-treated cells (**Fig. S5C**). Manganese has been proposed to be a cellular response to “rescue” iron-dependent enzymes that have lost their iron cofactor due to iron starvation or oxidative damage; biologically appropriately, manganese ion uptake is thus under control of Fur in Gram-negative model organisms ^52–54^. While a specific manganese ion importer has not been identified in *V. cholerae* ^55^, our data suggest that the cholera pathogen might possess a similar regulatory arrangement.

### Interfering with iron uptake partially restores tolerance to the ΔvxrAB mutant

If PenG-induced iron influx is detrimental, we would expect iron chelators to alleviate the Δ*vxrAB* β-lactam sensitivity defect. Indeed, addition of 250 µM (0.25 × MIC) of the intracellular iron chelator 2,2’ bipyridyl reproducibly increased survival of the Δ*vxrAB* mutant 50 – 100-fold (**Fig. S6**); the ability to partially rescue could be reverted by addition of iron to the growth medium. To probe the involvement of iron uptake systems even further, we sequentially deleted known iron uptake systems in the Δ*vxrAB* background and measured survival in the resulting mutants in the presence of PenG. Several combinations of Δ*vxrAB* with iron uptake system mutants reduced killing, particularly at early timepoints (**Fig. S7A-C**). Crucially, the Δ*vxrAB* Δ*vctPDGC* Δ*feoABC* Δ*fbpABC* mutant rescued the tolerance defect by ~100-fold (**Fig. 6A**). The Vct, Feo and Fbp systems encode uptake systems for the siderophore vibriobactin, ferrous and ferric iron, respectively ^56^; their importance is consistent with them likely being the main drivers of iron uptake under our experimental conditions (LB broth). In addition, we serendipitously found that the Δ*vxrAB* mutant is growth-defective on 10% sucrose plates, and this was exacerbated by addition of iron to the plate (**Fig. 6B**). Strikingly, deleting Δ*vctPDGC* Δ*feoABC* Δ*fbpABC* in the Δ*vxrAB* background completely restored growth on sucrose with or without 100 µM FeSO_4_ (but did not substantially increase plating efficiency at 1 mM FeSO_4_) (**Fig. 6BC**). Similar to our observations in PenG, combinations of Δ*vxrAB* with different combinations of iron uptake mutants had varying effects on restoration of growth on iron-supplemented sucrose (**Fig. S7 D-F**); most notably a mutation in the siderophore uptake system VctP itself was able to significantly improve plating efficiency of the Δ*vxrAB* mutant at lower iron concentrations (**Fig. S7D**). Thus, under cell envelope stress conditions, iron uptake significantly contributes to Δ*vxrAB* cell death or growth inhibition.

**Figure 6.**
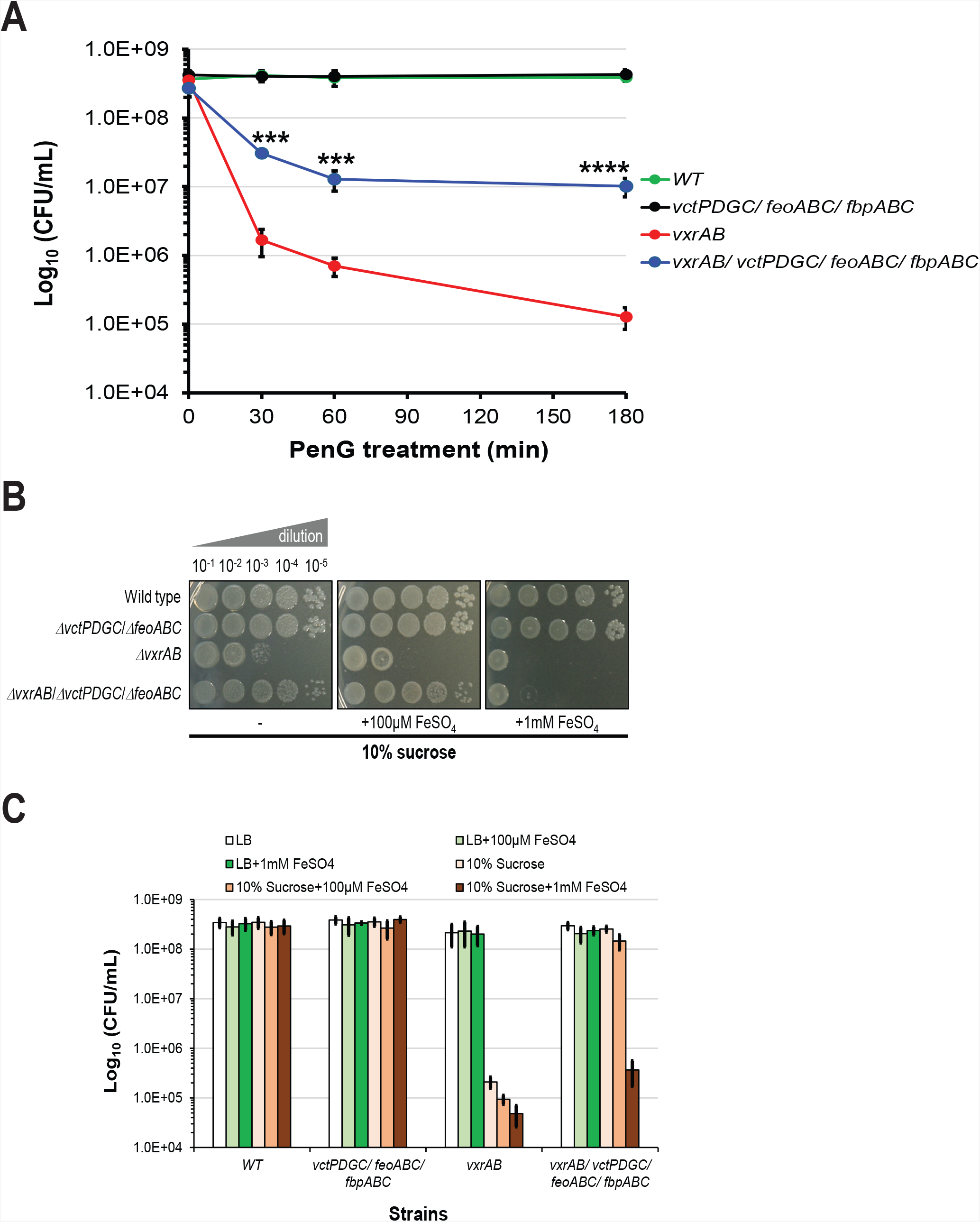
Disruption of iron influx systems partially restores tolerance of the *ΔvxrAB* mutant against PenG exposure. **(A)** Time-dependent killing experiment in the presence of PenG. **(B)** and **(C)** Iron uptake mutants restore ΔvxrAB growth on 10% sucrose. The indicated strains were grown to exponential phase (OD_600_~0.5) in LB medium and spot-plated on 10% Sucrose plates with or without additional iron sulfate. An example is shown in **(B)**, quantification of cfu is shown in **(C).** Data are average of 3 independent biological replicates, error bars represent standard deviation. *** P<0.0003; **** P<0.0001 (paired t-test).

### The iron chelator deferoxamine promotes β-lactam tolerance in a mouse model of cholera infection

Since our data thus far suggest iron influx is detrimental to tolerance in the absence of mitigating factors, we reasoned that the clinical use of iron chelators like deferoxamine^57–61^ may increase bacterial antibiotic tolerance as a collateral effect. We employed a mouse model of cholera infection (see Methods for details) to test this hypothesis *in vivo.* Streptomycin-treated mice were infected with *V. cholerae* and then treated with either deferoxamine, the oral β-lactam antibiotic amoxicillin, or both (**Fig. 7A**). *In vivo* pathogen survival under antibiotic pressure was quantified by assessing time-resolved viable cell counts from stool samples. As expected, viable counts from amoxicillin-treated mice drastically decreased over time (**Fig. 7B**), while deferoxamine alone did not affect pathogen survival compared to vehicle control. Strikingly, however, addition of deferoxamine to amoxicillin treatment completely undermined the latter antibiotic’s lethal effect and induced high-level amoxicillin tolerance (**Fig. 7B**). Thus, limiting available iron reduces antibiotic effectiveness *in vivo*; these results are also consistent with a model where detrimental iron accumulation results from β-lactam exposure.

**Figure 7.**
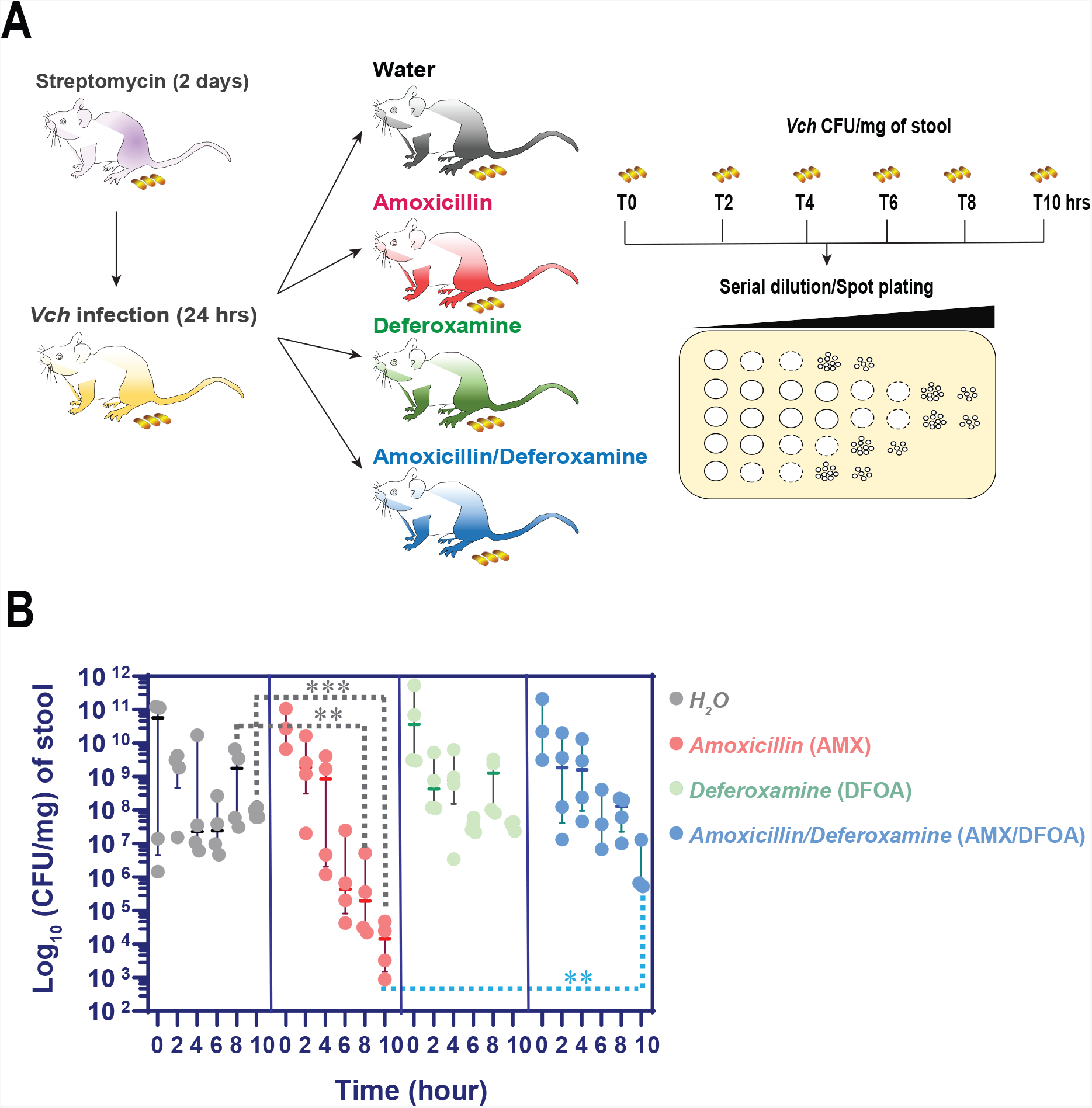
The iron chelator deferoxamine (DFOA) promotes amoxicillin (AMX) tolerance *in vivo*. **(A)** Schema of experimental design. Briefly, mice were pretreated with streptomycin (5 mg/mL) for 2 days and then intragastrically inoculated with *V. cholerae* (~1.97×10^10^ cells in 200 µl PBS). After 24 h, mice in each group received water (H_2_O), amoxicillin (50 mg/kg), deferoxamine (200 µl of a 100 µM solution), or both amoxicillin and deferoxamine. **(B)** Excretion of *V. cholerae*. Fecal samples were collected every two hours and assessed for their viable cell content (Colony Forming Units, CFU) to monitor the level of *V. cholerae* excretion. Each symbol represents fecal samples from an individual mouse (n=4). The number of bacterial CFU was normalized per milligram of feces (CFU/mg). ** P<0.005; *** P<0.001 (two-way ANOVA).

### Penicillin exposure induces ROS defense systems and oxidative damage

Iron is thought to be toxic to cells due to its ability to convert hydrogen peroxide into oxygen radicals that pleiotropically oxidize and damage macromolecules such as proteins and DNA ^62^ and generally disturb metal homeostasis ^63^. Indeed, antibiotics have been proposed to induce the increased production of ROS through metabolic perturbations ^29,31,37^, however, the extent to which this contributes to killing by antibiotics remains highly controversial ^32–34^. To interrogate the potential significance of ROS to lethality of the Δ*vxrAB* mutant, we first assessed whether oxidative stress response systems were induced by PenG. We found that PenG rapidly (within ~20 min) induced members of the hydrogen peroxide responsive OxyR1 regulon (*prxA* and *dps*) ^64–66^ in the wild-type strain (**Fig. 8A**). Transcript levels of both OxyR1 readouts remained high initially and then decreased to below baseline after ~ 60 min (*prxA*) or 180 min (*dps)*, suggesting that the wild-type strain induces a response that detoxifies (*prxA*) or protects against (*dps*) hydrogen peroxide (**Fig. 8A, S8A**). Both transcripts are induced by hydrogen peroxide in an OxyR1-dependent manner ^64,65^ and we validated responsiveness to hydrogen peroxide for the *prxA* transcript in our *V. cholerae* wild-type strain **(Fig. S8B).** Thus, *prxA* and *dps* are *bona fide* readouts for hydrogen peroxide production. Interestingly, the transcript for the response regulator OxyR1 itself was upregulated upon PenG exposure, with dynamics similar to those observed for members of its regulon (**Fig. 8A**). In the model organism *E. coli*, *oxyR* transcript levels do not change under hydrogen peroxide exposure ^67^ and our results suggest that *V. cholerae* might employ a different mode of regulation of its oxidative stress responses. A return to baseline coincided with the upregulation of *V. cholerae*’s two catalases (*katB* and *katG*; only *katG* appears to be under OxyR1 control in the cholera pathogen ^65,66^) at later stages of antibiotic exposure (**Fig. 8A**, **Fig. S8B**), likely revealing the detoxification mechanism that enables the repression of the OxyR1 regulon. Consistent with its hydrogen peroxide sensing function, KatG was also induced by hydrogen peroxide (**Fig. S8A**).

**Figure 8.**
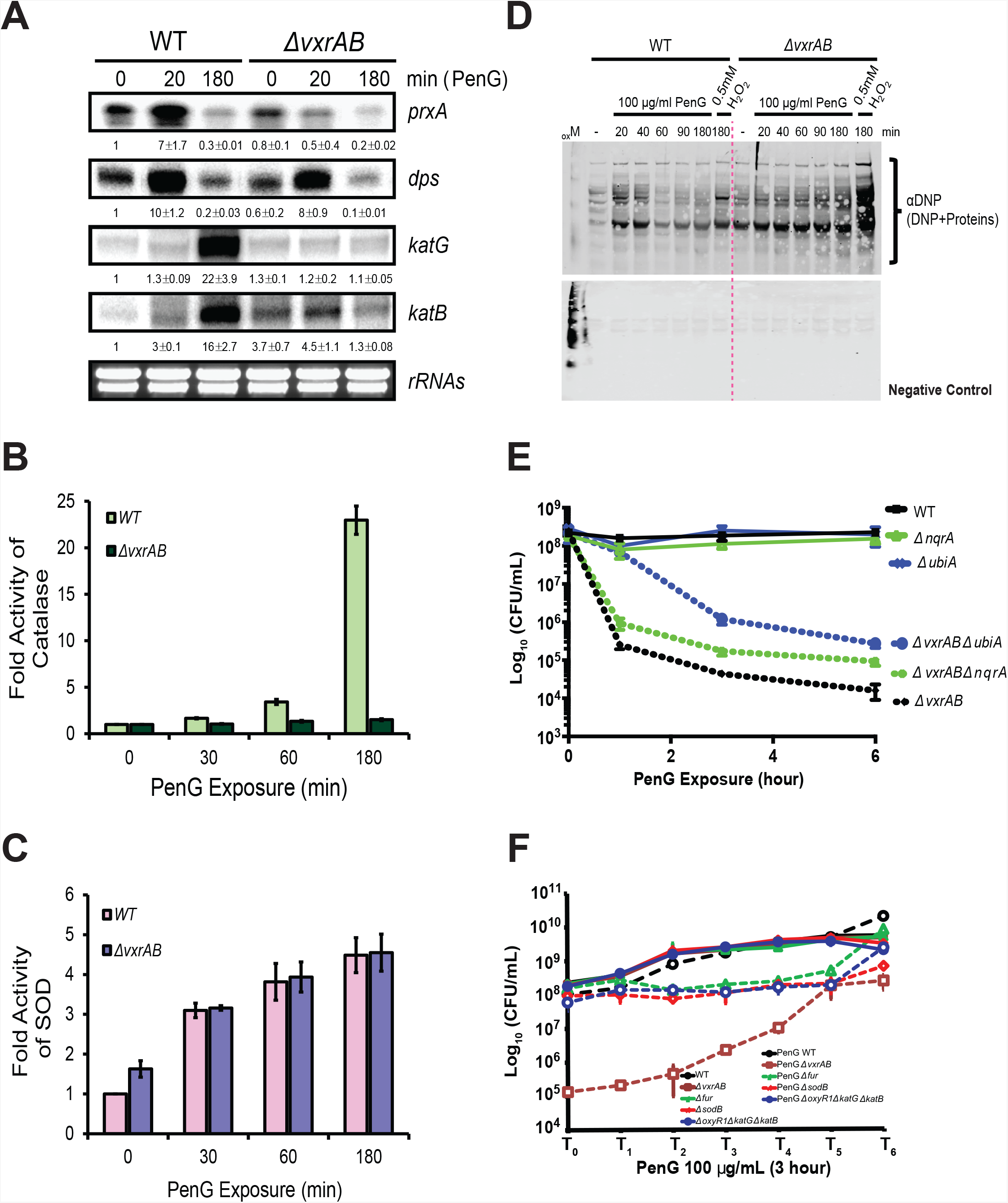
PenG exposure is associated with ROS production. **(A)** RNA expression profiles of first line ROS defense systems in *V. cholerae* monitored by S1 nuclease mapping assay. Quantified average values (normalized to WT before PenG exposure) from three independent experiments are presented under the signals of each gene. **(B and C)** Quantification of the SOD and catalase activity values from the gel shown in **Fig. S9**. Data are average values from three independent experiments normalized to untreated wild type. **(D)** Detection of protein oxidation after PenG treatment. To measure total cellular protein oxidation, OxyBlot Protein Oxidation Detection Kit (Sigma-Aldrich) was used. For positive protein oxidation control, oxidized standard marker (_OX_M) was used. Both DNP-derivatized or nonderivatized (negative control) protein samples were separated by polyacrylamide gel electrophoresis followed by Western blotting with-DNP specific antibody. **(E)** Time-dependent killing of respiratory chain mutants (Δ*nqrA*, encoding NADH dehydrogenase and Δ*ubiA*, encoding an early step of ubiquinone synthesis) in the presence of PenG (100 µg/ml, 10xMIC). All strains were grown in LB 0.2 % glucose. **(F)** Growth-phase dependent killing assay of ROS detoxification systems (Fur, SodB, OxyR1, KatG, and KatB). At designated time-points, a 5 ml aliquot was withdrawn and exposed to 100 µg/ml PenG for 3 h. CFU/mL for each time point was assessed before (solid lines) and after addition of PenG (dotted lines). Data are mean of three independent biological replicates and error bars represent standard deviation.

In the Δ*vxrAB* strain, *oxyR1* baseline levels were slightly increased compared to the wild-type, while *prxA* and *dps* levels were similar. After exposure to PenG, *oxyR1* levels decreased at first but the transcript remained detectable and above the wild-type baseline for longer than the wild-type background (**Fig. 8A, S8**). The *prxA* and *dps* transcripts exhibited a dynamic pattern with strong induction (*dps)* or transient downregulation (*prxA*) in the mutant. We observed a pattern similar to *oxyR1* with a gene coding for another oxidative stress mitigation factor, the alkyl hydroperoxide reductase C, *ahpC* (**Fig. S8**). Interestingly, the transcript for *katG* remained uninduced in the *vxrAB* mutant.

We confirmed that transcriptional regulation also resulted in increased catalase protein activity using a gel-based activity assay for catalases (**Fig. S9A**, see Methods for details) after exposure to PenG. We also included an assay for superoxide dismutase (SOD) activity, since SODs are broadly conserved, important ROS scavenging enzymes^68^. Consistent with the observed transcript levels (**Fig. 8B**), catalases were only induced in the wild-type, but not the mutant (**Fig. 8B**). In contrast, SOD activity increased similarly in both the wild-type and the *vxrAB* mutant (**Fig. 8C**). Taken together, these results suggest that both the wild type and Δ*vxrAB* induce responses upon exposure to PenG that are associated with oxidative stress, albeit with different, complex dynamics.

We next assessed the cellular consequences of oxidative damage using a protein oxidation assay via Western Blot. Consistent with our results showing enhanced iron sensitivity of the mutant, protein oxidation levels were increased in the Δ*vxrAB* mutant vs. wild-type even prior to addition of antibiotic (**Fig. 8D**). Addition of PenG resulted in a rapid (within 20 min) increase in protein oxidation levels in the wild-type, which decreased after 40 min but stayed above baseline throughout β-lactam exposure. In contrast, the *vxrAB* mutant steadily accumulated oxidized proteins throughout the exposure time, corroborating that oxidative stress has a more profound effect on the *vxrAB* mutant than the wild-type strain. Both wild-type and the *vxrAB* mutant exhibited increased protein oxidation when exposed to hydrogen peroxide as a positive control (**Fig. 8D, Fig. S9B**). Taken together, these data suggest that both wild-type and Δ*vxrAB* experience oxidative stress in the presence of PenG, but the Δ*vxrAB* mutant is unable to mount a mitigating response.

### The electron transport chain is required for both maximal killing and hydrogen peroxide production

How does PenG treatment result in production of ROS? A possible origin is the electron transport chain (ETC), which in *V. cholerae* is a major generator of ROS under normal growth conditions ^69^. To test the ETC’s involvement in PenG-induced ROS production, we created mutants in two ETC components, namely *V. cholerae’s* principal sodium translocating NADH dehydrogenase NqrA ^70^ and *ubiA*, which encodes a key initial step towards ubiquinone biosynthesis. NqrA has been shown to be a significant source of ROS under normal growth conditions ^69^, but for both mutants, we expect a drastic reduction in ETC electron flow. Crucially, neither mutation resulted in a significantly decreased growth rate when grown in medium containing the fermentable carbon source glucose (albeit, both mutations caused a slight increase in lag phase) (**Fig. S10A**). Reduced growth rates can affect β-lactam susceptibility non-specifically ^71^ and mutants conferring slower growth can thus confound the interpretation of experiments conducted with such strains. Consistent with the expectation of reduced ETC activity in these mutants, disruption of *nqrA* and *ubiA* in both the WT and the Δ*vxrAB* mutant background resulted in a marked reduction in PenG-induced accumulation of hydrogen peroxide, as measured by Amplex Red (**Fig. S10B**). To test the hypothesis that the reduction in hydrogen peroxide also results in lack of killing in the Δ*vxrAB* mutant, we conducted time-dependent killing experiments with these strains. As observed previously, wild-type viability remained stable for the entire duration of the experiment while the Δ*vxrAB* mutant displayed a ~10,000-fold decrease in viability. The Δ*nqrA* and Δ*ubiA* mutants were as tolerant as the wild-type. However, a combination of ETC mutations with Δ*vxrAB* reproducibly increased survival by 10-(Δ*nqrA*) or 100-fold (Δ*ubiA*) (**Fig. 8E)**. Thus, Δ*vxrAB* lethality after PenG exposure partially depends on the presence of the ETC.

### ROS detoxification systems promote tolerance at high cell densities

We next asked whether ROS damage mitigation systems were required for wild-type tolerance. We created knockout mutants deficient in the iron starvation repressor Fur (expected to result in an increase in intracellular iron and thus enhanced ROS), SodB and a triple mutant defective in both catalases and OxyR1. We grew all strains from late exponential into stationary phase and periodically withdrew aliquots that were then exposed to PenG for 3 h, and measured survival. As a comparison, we subjected the Δ*vxrAB* mutant to the same treatment. While all mutants except for Δ*vxrAB* exhibited wild-type survival patterns in exponential phase, entry into stationary phase coincided with significant killing in all mutants that are expected to experience enhanced ROS production (**Fig. 8F**). Consistent with these results, *sodB* had previously answered a Transposon Insertion Sequencing (TnSeq) screen for factors that are required for PenG tolerance ^72^. We thus propose that ROS contribute significantly to β-lactam-mediated killing in stationary phase (see Discussion for details).

## Discussion

Our overall model of VxrAB-mediated tolerance is depicted in **Fig. 9;** based on the evidence presented in this study, we propose the following: PenG induces cell wall degradation, resulting in perturbations of the ETC. Our work does not directly address the nature of this perturbation, but it may simply result from topological alterations of the cytoplasmic membrane brought about by loss of the cell wall. Interestingly, and consistent with this line of thinking, other cell wall deficient cells (L-Forms) also appear to accumulate ROS^73,74^. We hypothesize that partial reduction of molecular oxygen through the ETC (particularly NqrA) results in the production of superoxide ion. Superoxide ions are removed by SOD, which catalyzes the formation of hydrogen peroxide. Hydrogen peroxide induces the *oxyR1* regulon, but we propose that it also inappropriately inactivates the Fur repressor via oxidation of its ferrous ion cofactor. Inactivation of Fur (i.e., induction of the Fur regulon) results in the upregulation of iron uptake systems, which increases the intracellular iron pool. Intracellular iron reacts with hydrogen peroxide to form hydroxyl radical, which damages proteins and other macromolecules. The wild-type strain mounts effective responses to all these types of damage. SOD, AhpC and catalases detoxify superoxide ions and hydrogen peroxide. VxrAB induces cell wall synthesis functions to prepare cells for speedy recovery once the antibiotic is removed. At the same time VxrAB also downregulates some iron acquisition genes and other Fur functions to dampen detrimental iron influx and potentially concomitant oxidative damage. Interestingly, some of our data suggest that ROS production might contribute significantly to killing in later growth phases (early stationary phase), and stationary phase-specific oxidative damage is not without precedent. Periplasmic SODs are under control of stationary phase regulators in diverse bacteria and yeast ^75–77^ and SOD is required for antibiotic tolerance in stationary phase in *P. aeruginosa* as well ^78^. We hypothesize that during rapid growth in exponential phase, the primary mechanism of action (cell wall degradation) is likely the more important contributor to cell death, while in stationary phase (slower/no growth, less cell wall damage), the harmful effects of ROS accumulation may become more visible. This is noteworthy, since in the natural environment, bacterial growth can be expected to resemble stationary phase more than rapid growth in exponential phase ^79^. On a separate note, β-lactam-mediated killing in stationary phase may seem counterintuitive at first, since at least in the well-characterized model organism *E. coli*, cessation of growth upon entry into this growth phase is generally assumed to result in complete protection from β-lactam-induced damage ^71,80^. However, the *vxrAB* mutant still exhibited significant PenG-dependent mortality even when PenG exposure only occurred several hours into stationary phase, i.e., far into a period of non-growth, suggesting that this mutant experiences PenG-induced cell envelope damage even when populations stop growing. We thus propose that in *V. cholerae*, cell wall turnover (and thus PenG’s ability to inflict damage) continues at least for some time after cells stop growing. A similar phenomenon could explain the unusual observation that the Lyme disease pathogen *Borrelia burgdorferi* remains susceptible to killing by cell wall acting antibiotics in stationary phase ^81^.

**Figure 9.**
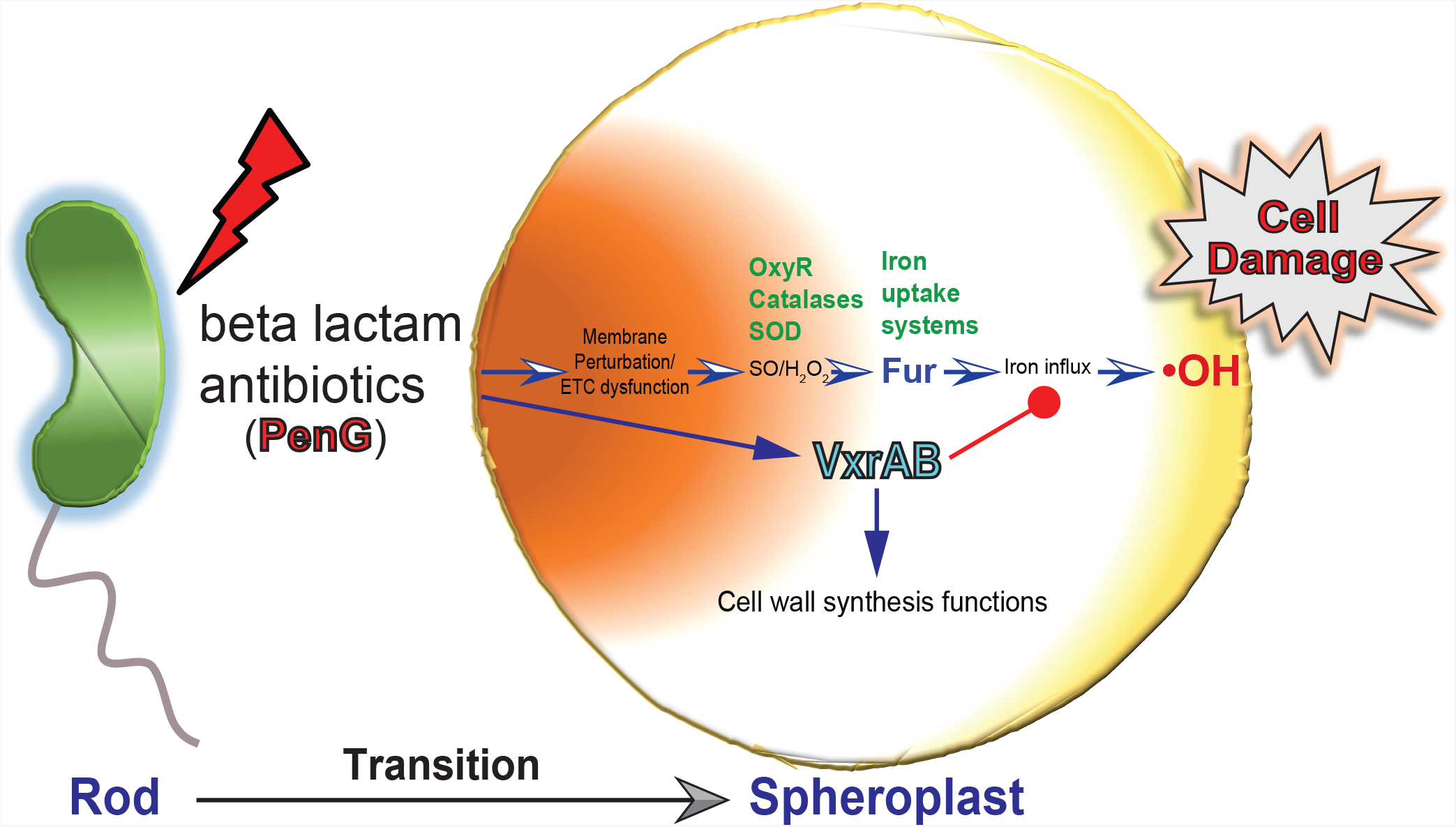
Proposed model for VxrAB-mediated antibiotic tolerance. PenG induces ETC dysfunction during transition of cell shape from rod to spheroplast, resulting in the production of ROS. Hydrogen peroxide illegitimately induces the Fur regulon, resulting in the upregulation of iron acquisition systems, resulting in enhanced iron influx and lethal oxygen radical generation. VxrAB dampens this response by downregulating iron acquisition genes, in addition to its primary role as a positive regulator of cell wall synthesis genes. SO, superoxide.

The connection between antibiotic exposure and ROS production and particularly the extent of the contribution of ROS to killing is still actively debated in the field. We agree with a previous hypothesis^34^ that some of the interpretive difficulties in other studies may have arisen from the low concentrations of antibiotics used (often around the MIC). High concentrations of bactericidal antibiotics might kill cells too quickly through their primary mechanism of action to measure a response to the proposed secondary mechanism. Lower concentrations around the MIC might result in day-to-day variation of killing dynamics that are difficult to reproduce under different conditions. In contrast, bacteria like *V. cholerae* (as well as *P. aeruginosa, Klebsiella pneumoniae, Klebsiella* (formerly *Enterobacter*) *aerogenes* and *Enterobacter cloacae*), exhibiting almost population-wide tolerance to high concentrations of β-lactams ^11,13,14,72^, may be more appropriate model organisms to dissect the layers of physiological changes occurring as a consequence of antibiotic exposure. In these tolerant bacteria, the primary mechanism of action (degradation of the cell wall) is recapitulated; however, unlike in other bacteria, this does not result in lysis and cell death, allowing us to examine a large population of live cells after treatment with high concentrations (10 × MIC or more) of antibiotic.

One of the main critiques of the idea that ROS contribute to killing was that some groups failed to measure meaningful ROS levels (and the expected responses to them) in antibiotic-treated cells in the model organism *E. coli* ^33,34^. This has been challenged by other groups^35,37,82^. Here, we have shown that *V. cholerae* accumulates ROS and induces defense systems against them after exposure to PenG. We propose that in the *vxrAB* mutant, antibiotic-induced lethality is multifactorial. Both, cell wall defects and ROS may contribute to killing. Importantly, other detrimental consequences of iron influx may also play a role, for example mismetalation of key enzymes ^63^. While ROS production and iron influx are measurable in the wild-type as well, albeit to a lower degree than the mutant, it is likely that its multiple, highly effective defense systems prevent antibiotic killing. This thinking is consistent with a previous observation that compared to *E. coli* and *P. aeruginosa*, exposure of *V. cholerae* to sub-MIC of β-lactam antibiotics only caused a moderate increase in mutagenesis (a possible consequence of ROS accumulation) ^83^. Thus, the cholera pathogen’s high tolerance to β-lactam exposure might be the consequence of an exceptional ability to repair cell damage.

Since we demonstrate here that PenG-treatment induces toxic iron uptake, it is important to note that iron withdrawal is a common antibacterial strategy in the human host (and deferoxamine is clinically used to treat iron homeostasis disorders), which might significantly affect antibiotic susceptibility *in vivo*^84^. The extent of the host’s ability to control iron levels in a specific infection locale might significantly and divergently influence antibiotic susceptibility in different host tissues. In summary, β-lactams (and likely other antibiotics) elicit more complex cellular responses than just those associated with their primary mechanism of action. A more context-, metabolism- and physiology-centered view of antibiotic action ^85–88^ might therefore yet yield critical insight to enable the development of novel adjuvants that increase the efficacy of antibiotics in the complex infection environment. Lastly, with or without ROS, our data suggest that targeting bacterial stress responses could be a powerful strategy to enhance the activity of bactericidal drugs against tolerant pathogens.

## Supporting information

Supplementary Informations

## Acknowledgments

Research in the Dörr laboratory is supported by NIH grants R01AI143704 and R01GM130971. This work was partially supported by the Korea Bio Grand Challenge Project (2018M3A9H3024759 to B.K.C.) through the National Research Foundation of Korea (NRF) funded by the Ministry of Science and ICT, Republic of Korea. Research in the Brito Laboratory is funded by the Packard Foundation, Sloan Foundation, Pew Foundation and NIH grant 1DP2HL141007. We thank the Helmann lab (Cornell University) for sharing equipment and reagents. We further thank Stavroula Hatzios (Yale University), John Helmann (Cornell University), Joseph Peters (Cornell University), James Imlay (University of Illinois) and members of the Dörr lab for helpful comments.

## MATERIALS AND METHODS

### Bacterial strains and culture conditions

All *V. cholerae* strains used in this study were derivatives of *V. cholerae* WT El Tor strain N16961 or E7946 (for chitin-induced transformation) listed in **Table S1**. *V. cholerae* was grown on Luria-Bertani (LB) medium with or without 0.2% glucose or in M9 medium containing 0.2% glucose at 37°C unless otherwise indicated. All media contained 200 µg/ml streptomycin (to which *V. cholerae* is resistant). Where applicable, growth media were supplemented with 100 μg/ml Benzylpenicillin potassium salt (PenG) (Fisher BioReagents, cat# BP914-100), isopropyl-β-D-1-thiogalactopyranoside (IPTG; final 0.1~1mM) (GOLDBIO, cat# 12481C50), or catalase 1μg/ml (Sigma-Aldrich, Cat# c9322). *Escherichia coli* DH5α and DH5α λpir were used for routine DNA cloning. Unless indicated otherwise, LB liquid media were inoculated from an overnight culture and incubated at 37°C with shaking at 200 rpm until reaching mid-exponential phase (OD_600_ ~0.4). Where applicable, trimethoprim (Sigma, cat #T7883, dissolved in DMSO) was used at 15 µg/ml final concentration.

### Strain construction

Deletion mutants were generated via replacement of the open reading frame/binding site with the scar sequence TAATGCGGCCGCACTCGAGTAATAATGATGA or, where indicated, a trimethoprim resistance cassette. Knockouts were generated either by homologous recombination using the suicide vector pCVD442 ^91^ or by chitin-induced natural transformation, both as described before ^92^ ^17^. The following primer combinations were used (see supplemental material for sequence information).

*murJ* Δ*vbs*: TDP1062/1064 + TDP1066/1067, fused via stitching PCR with nesting primers TDP1078/1079, followed by chitin transformation using *vc1807*::trim [generated using primers TDP597/598 + TDP599/600 (on trimR from the *V. cholerae* HAITI strain 93)+TDP601/602, fused via stitching PCR with nesting primers TDP603/604] as primary selection. lacZ::P_ryhB_:*msfGFP*: TDP1152/1235 + TDP1237/38, cloned into *SmaI*-digested pTD101, a suicide vector for chromosomal integration into the *lacZ* locus ^95^. Δ*nqrA*: TDP1388/1389 + TDP1390/1391, fused via stitching PCR with nesting primers TDP1392/1393, followed by chitin transformation using *vc1807*::trim (see above) as primary selection. Δ*ubiA:*:*trim* TDP1399/1401 + TDP1404/1403 + TDP636/637, fused via stitching PCR with nesting primers TDP1405/1406, followed by chitin transformation. Δ*oxyR1*: TDP1057/58 + TDP1059/60, Δ*katG:* TDP1100/1101 + TDP1102/1103, Δ*katB*: TDP1103/1104 + TDP1105/1106, Δ*sodB*: TDP828/829 + TDP830/831, Δ*fur*: TDP834/835 + TDP836/837 cloned into *XbaI*-digested pCVD442 using isothermal assembly ^94^ and Δ*vctPDGC:* TD-JHS394/395 (by amplification from gene block; *vctPDGC* deletion), Δ*feoABC:* TD-JHS397/398 (*feoABC* deletion), Δ*fbpABC:* TD-JHS400/401(*fbpABC* deletion), Δ*vc2211:* TD-JHS403/404 (*vc2211* deletion), Δ*vc2212:* TD-JHS406/407 (*vc2212* deletion), Δ*tonB2exbB2:* TD-JHS409/410 (*tonB2exbB2* deletion), and Δ*hutA:* TD-JHS412/413 (*hutA* deletion) cloned into *SphI/XmaI*-digested pCVD442 followed by knockout via homologous recombination.

#### Pulldown of VxrB binding DNA fragments and ChIP-Seq library construction

Cells carrying either 6xHis+VxrB or the native gene were grown in LB medium to mid-logarithmic phase (at an OD_600_ ~0.5). Following treatment with PenG (100 µg/ml, 3 hours), 30 ml aliquots were spun down and the pellets were saved at −80°C. For cross-linking, pellets were resuspended in 1ml buffer CJ1 [100 mM Na_2_HPO_4_, 600 mM NaCl, and 5 mM imidazole (pH 8.0)] and 30 μl of 34% formaldehyde was added, followed by rocking at room temperature for 10 min. To quench cross-linking reaction, 133 μl of 1 M Glycine pH7.5 (final molar concentration 0.133 M) was added, and the cells were shaken gently at 4°C for 30 min. Cells were collected by centrifugation, and the pellets were resuspended in 1 ml buffer CJ2 [100 mM Na_2_HPO_4_, 600 mM NaCl, 0.05% tween™, and 30 mM imidazole (pH 8.0)] after washing twice with same CJ1 buffer. Cells were then lysed by sonication, followed by addition of Micrococcal nuclease (100LULml^−1^, NEB cat# M0247S) and RNase A (1LmgLml^−1^) for fragmentation of genomic DNA and removal of RNA. 950 μl of supernatant was collected after centrifugation.

For DNA-VxrB complex purification, HisPur™ Ni-NTA magnetic beads (cat#88831-Thermo Scientific) were equilibrated with six bed volumes of CJ2 buffer [100 mM Na_2_HPO_4_, 600 mM NaCl, 0.05% tween™, and 30 mM imidazole (pH 8.0)]. 500 μg of crude lysate (protein concentration determined by Bradford assay) was mixed with the Ni-NTA resin and incubated at 4°C for 4 hours on rotation mixer. The bead slurry was recovered via magnetic stand for 5 min standing and washed twice with 400 μl buffer CJ3 [100 mM Na_2_HPO_4_, 600 mM NaCl, 0.05% tween™, and 50 mM imidazole (pH 8.0)]. The VxrB-DNA complexes were then eluted from the beads by addition of CJ4 elution buffer [100 mM Na_2_HPO_4_, 600 mM NaCl, 0.05% tween™, and 500 mM imidazole (pH 8.0)]. Enriched target DNAs were purified by using PCR Clean-Up Kits (Thermo Fisher Scientific) after reversal of crosslinking at 65°C overnight. The desired DNA fragment size (200~300 bps) was then purified by gel extraction. For illumina HiSeq, each DNA library was constructed with specific barcode sequences, according to the manufacturer’s instruction of TruSeq ChIP Library Preparation Kit (IP-202-1012, Illumina). The constructed libraries were sequenced using 100-bp paired-end read protocol on an Illumina HiSeq 2500 platform (Illumina) at the Biotechnology Resource Center (BRC) at Cornell University.

#### ChIP-Seq, binding peak detection, and motif analysis

Sequencing data was processed by CLC Genomics Workbench (CLC Bio). First, raw sequencing reads were quality trimmed using a Trim Sequence Tool with a quality limit of 0.05. Any reads with more than two ambiguous bases were discarded at this step. Then, quality trimmed reads were mapped onto the *V. cholerae* reference genome (NCBI accession NC_002505.1 and NC_002506.1) with mismatch cost of 2, insertion/deletion cost of 3, length fraction of 0.9, and similarity fraction of 0.9. Non-specific matches were mapped randomly. Mapping files (.bam) were submitted to the Model-based Analysis of ChIP-Seq 2 (MACS2) software with a mapping file of a mock sample as a control. Peaks were detected with shift size of 50 and q-value cutoff of 10^−10^. For motif analysis, peaks with q-value lower than 10^−40^ were selected and 100 bp DNA sequences centered at peak summits were submitted to MEME Suite software v4.11.4.

### RNA-Seq analysis

Sequencing data (obtained previously) were also processed on a CLC Genomics Workbench (CLC Bio). Raw sequencing reads were quality trimmed with the same method with ChIP-Seq reads. Quality trimmed reads were mapped strand specifically on the reference genome sequence by Transcriptome Analysis Tool with the following parameters: maximum number of mismatches of 2, length fraction, and similarity fraction of both 0.9. Number of mapped reads on each gene was normalized by DESeq2 package in Bioconductor v3.4. Genes with p-value (padj, DESeq2) lower than 0.001 were scored as differentially expressed genes (DEGs).

#### Purification of VxrB protein

Wild-type VxrB protein was purified from *E. coli* BL21 (DE3/pLysS) cells containing pET15b based recombinant plasmid. The coding sequence for WT VxrB was cloned into pET15b (Novagen) into the *Nde*I and *Bam*HI restriction sites. For purification of VxrB from this construct, an overnight culture from a single colony was used to inoculate 1 liter of LB medium. Cells were grown with vigorous shaking at 37°C to an optical density at 600 nm (OD_600_) of 0.5 and were induced with 1 mM (final concentration) isopropyl-β-D-thiogalactopyranoside (IPTG) for 6 h at 30°C. Harvested cells were re-suspended in binding buffer [20 mM Tris-HCl (pH 7.9), 0.5 M NaCl, 1 mM TCEP (tris(2-carboxyethyl)-phosphine), and 5 mM imidazole] and cell extracts were prepared by sonication and centrifugation at 16,000 rpm for 45 min. Cell extracts were loaded onto a cobalt-charged NTA column and then washed with 6 volumes of binding buffer followed by 6 volumes of washing buffer [20 mM Tris-HCl (pH 8.0), 0.5 M NaCl, and 50 mM imidazole]. VxrB protein was eluted with 10 volumes of elution buffer [20 mM Tris-HCl (8.0) and 0.5 M NaCl] containing linear imidazole gradients from 100 to 500 mM. Fractions containing VxrB protein were pooled and dialyzed against buffer J1 [20 mM Tris-HCl (pH 8.0), 250 mM NaCl, 5% (vol/vol) glycerol, and 4 mM EDTA] to remove imidazole and cobalt and then transferred to two extra dialysis buffer J2 [20 mM Tris-HCl (pH 8.0), 150 mM NaCl, 10% glycerol, and 0.2 mM DTT] and buffer J3 [20 mM Tris-HCl (pH 8.0), 150 mM NaCl, 30% glycerol, and 1 mM DTT]. The purified VxrB protein was concentrated by centrifugal filter devices (Millipore, 3,000 MW CO) before injection onto High load TM (16/60) pg Superdex G75 column in FPLC system (Pharmacia). The column was equilibrated with buffer SG [20 mM Tris-HCl (pH 8.0), 150 mM NaCl, and 1 mM DTT]. Eluted fractions were monitored through UV detector. Concentration of purified VxrB was estimated in triplicate by Bradford assay (Bio-Rad) using BSA (bovine serum albumin, Sigma-Aldrich) as the calibration standard at A595. Measurement of UV absorbance at 280 nm, combined with calculated molar extinction coefficient (ε_280_ = 46470 M^−1^ cm^−1^, http://expasy.org/cgi-bin/protparam), gave nearly identical values. The purity of protein was confirmed through Coomassie blue staining of loaded protein samples in the SDS-PAGE gel. The protein was stored in final storage buffer JHS [20 mM Tris-HCl (pH 8.0), 150 mM NaCl, 1 mM DTT, 30% Glycerol, 1 mM MgCl_2_, and 1 mM ATP] at −80°C.

#### DNase I foot printing assay

The probe DNAs were prepared by using the “crush and soaking method” (Sambrook *et al*., 1989) after PCR with the primer pairs shown in Table S2. The 5′ end of the probes was labeled with 6-FAM (Fluorescein). Binding reactions were performed as described below for EMSA, except using 250◻ng probe DNA in a 40◻μl reaction mixture. DNA probes were incubated with increasing concentrations of VxrB (0.17, 0.34, 0.68, and 1.38◻μM). After 20◻min incubation at room temperature, 40 μl of 5LmM CaCl_2_ and 10◻mM MgCl_2_ was added, followed by adding 200◻U of RQI RNase-free DNase I (Promega) for 1◻min. The cleavage reaction was stopped by adding 90◻μl stop solution (200◻mM NaCl, 30◻mM EDTA, 1% sodium dodecyl sulfate, 125◻μgLml^−1^ of glycogen), followed by DNA extraction and precipitation. Samples were analyzed by capillary electrophoresis on an ABI 3730×1 DNA analyzer (Life Technologies). The DNA probe only with no added VxrB was analyzed in parallel. The core protected regions were indicated with red lines and its corresponding DNA sequences were shown at the bottom of peak profiles.

#### Preparation of total RNA and S1 nuclease mapping analysis

Wild type and mutant strains were cultured to mid-logarithmic phase (at an OD_600_ of 0.4 to 0. 5) in LB medium. Following application of a stress (PenG or H_2_O_2_), total RNA was extracted using the “hot phenol method”, albeit without lysozyme pretreatment. RNA was quantified and quality-controlled by absorbance spectroscopy and confirmed by resolving RNA samples on 1.3% formaldehyde-agarose gels. Gene-specific DNA oligonucleotide probes for *vxrA*, *vxrB*, *pbp1A*, *zapB*, *flaA*, *murJ*, *oxyR1*, *sodB*, *sodA*, *katG*, *katB*, *prxA*, *oxyR2*, *ahpC*, *dps*, *fur*, and *ryhB* transcripts were generated by PCR amplification using *V. cholerae* wild-type genomic DNA as template. The appropriate primer pairs are listed in **Table S2**. Each specific DNA probe was radiolabeled with (γ-^32^P) ATP and T4 polynucleotide kinase and 30,000-40,000 cpm of labeled probe was used in each reaction. For S1 assays, 100 μg of total RNA was pelleted and lyophilized. The total RNA pellet was then carefully resuspended in 20 μl hybridization buffer [40 mM PIPES (pH 6.4), 400 mM NaCl, 1 mM EDTA, 80% (v/v) formamide]. Individual samples were incubated at 95°C for 25 min and cooled to 42°C. Following overnight incubation, 300 μl of S1 nuclease mix containing 100 units of S1 nuclease in S1 nuclease buffer [280 mM NaCl, 30 mM NaOAc (pH 4.4), 4.5 mM ZnOAc] was added and incubated at 37°C for 45 min. The reaction was terminated by addition of 75 μl of S1 nuclease termination solution (2.5 M NH_4_OAc, 0.05 M EDTA). The DNA-RNA hybrid was precipitated by adding 400 μl of isopropanol (100 % v/v) and the pellet was washed with 70% (v/v) ethanol, vacuum dried, and re-suspended in 13 μl alkaline loading dye. The protected DNA fragments were then resolved by 6% (w/v) polyacrylamide gels containing 7 M urea. Gels were dried using a Hoefer Scientific Drygel Slab Gel Dryer 01385 and then imaged on a phosphor imaging screen (Typhoon FLA 7000; GE); bands were quantified using Multi Gauge V3.0 (Fuji). Quantified average fold intensity (compared to non-treated sample) from three independent experiments were presented under each signal with standard deviation values.

#### Killing Assay

For most time-dependent killing experiments, overnight cultures of wild type and each mutant strains were grown at 37°C in LB medium in the presence or absence of additional supplement such as IPTG,0.2% glucose, or 10% sucrose. At each time-point, samples were collected, serially diluted (sterile PBS) and spot-plated for CFU/mL determination (plates were incubated at 30 °C for 20 – 24 h). Where indicated, PenG (100 μg/ml), 250 μM Bip, 250 µM FeCl_2_, 100 µM, or 1mM FeSO_4_ were added. For the growth phase dependent experiment, wild type, Δ*vxrAB*, Δ*fur*, Δ*sodB* and Δ*oxyR1*Δ*katG*Δ*katB* mutants were cultured in the presence of 1μg/ml catalase for overnight seed (catalase is required for Δ*oxyR1* growth) in 100 mL LB in flasks. At designated time points, 5ml cell cultures were transferred to new tubes and cultured with or without PenG for 3 hours; CFU/mL was determined before and after addition of the antibiotic. Where applicable, cells were stained with 5 µM of the membrane dye FM4-64 (Invitrogen, cat# LST3166) for 5 min. Collected images were processed in ImageJ by subtracting background and adjusting brightness/contrast uniformly across all fluorescent images.

### Streptonigrin sensitivity assay

Cells were diluted 100 fold from overnight cultures and grown to a density of ~2 × 10^8^ CFU/mL. At each time point (before addition of 100 µg/ml PenG and 3 h after), 200 µL of culture were spread on 25 ml LB agar plates using glass beads. At the 3 h time point, 1 mg/ml (final concentration) β-lactamase (Sigma, catalog # P4524) was added prior to plating. Following plating, a sterile filter disk containing 5 µL of 5 mg/ml streptonigrin solution (Sigma, catalog # S1014, dissolved in DMSO) was placed in the center of the plate. Plates were then incubated 24 h at 37 °C. Measurements are averages of six biological replicates.

#### ICP-MS analysis for intracellular metal concentration measurement

A single colony of the indicated *V. cholerae* strains from LB plates was inoculated into 5 mL LB with streptomycin (200 µg/mL) and grown overnight. The next morning, 0.1% of overnight cultured seeds were inoculated in 200 ml LB supplemented with antibiotics. These cells were grown shaking at 37°C for 2~3 hours until they reached an OD_600_ of ~0.5. Samples were collected before and at indicated time points after PenG treatment. At the indicated time points, cells were pelleted by centrifugation (10 min, 20000 rcf) and washed three times with phosphate buffered saline (PBS) buffer containing 0.1 M EDTA followed by two chelex (BioRad, cat # 1422822)-treated PBS buffer washes to remove all external metal ions. Cells were then resuspended in 400 μl of chelex-treated PBS buffer with 1% Triton X-100 containing 75 mM NaN3 to induce lysis and incubated at 37°C for 90 min. 10 μl of each supernatant was used to measure total cell protein concentration by Bradford assay. 600 μl of 5% HNO3 with 0.1% (v/v) Triton X-100 were then added to the rest of the supernatant, which was boiled at 95°C for 30 min. After centrifugation (10 min, 20000 rcf), the supernatant was then diluted 1000fold with 1% HNO3. Metal levels were measured by ICP-MS (Perkin-Elmer ELAN DRC II using ammonia as the reaction gas and gallium as an internal standard) and normalized against total cell protein concentration. Data represent with average values from biological triplicates. Significant differences between wild type and Δ*vxr*AB after PenG exposure were determined by paired R t-test as indicated: **P=0.001; ***P<0.0003.

#### Catalase and SOD activity assay

Catalase activity staining was performed. Briefly, cell extracts (30 μg per well, obtained by 20 cycles of sonication on ice at 20 % power, 3 s pulse, followed by 10 s cooling) were separated on a 5% native polyacrylamide gel and electrophoresed at 4°C with 80 volts. The gel was washed in distilled water and then treated with 1 mM H_2_O_2_ for 20 min. After treatment, the gel was rinsed and transferred to 1% (w/v) ferric chloride and 1% (w/v) potassium ferricyanide solution. The reaction was stopped by washing twice in distilled water. Superoxide dismutase activity staining was performed as described in ^96^. The cell extracts (30 μg/well) were electrophoresed on an 8% native polyacrylamide gel under the same conditions as described above for catalase activity, after which the gel was incubated in a solution containing 2.5 mM nitroblue tetrazolium (NBT) for 25 min and then in 50 mM phosphate buffer (pH 7.8) containing 28 μM riboflavin and 28 mM tetramethyl ethylene diamine for 20 min in the dark. The gel was placed in distilled water and exposed on a light box for 15 min until the dark blue background color appeared. To quantify SOD and catalase activity, stained gels were scanned and analyzed by Odyssey Imaging Systems (LI-COR Biosciences) and Image Studio Lite software. Quantified average values from three independent experiments are presented as a fold activity against wild type non-treated sample for SODs and Catalases.

#### Measurement of total cellular H_2_O_2_ level

Total cellular hydrogen peroxide (H_2_O_2_) level was measured using the Amplex™ Red Hydrogen Peroxide/Peroxidase Assay Kit (Invitrogen™ cat#A22188) and normalized to total cellular proteins. Wild type and indicated mutant stains were cultured in LB medium (in the absence or presence of additional 0.2% glucose when it needed) until OD_600_ ~0.5. PenG (100 μg/ml) was then added and 1ml cell culture was withdrawn and pelleted at each indicated time point. Pellets were resuspended in 100 μl cold 1xPBS buffer and lysed by sonication, followed by centrifugation at 4°C (10 min, 20000 rcf). 3 μl of each supernatant was used to measure total cell protein concentration by Bradford assay. 20μl extracts from cleaned supernatant were reacted with kit components following manufacturer’s instruction. For H_2_O_2_ quantification, the reaction product resorufin was read by fluorometer (QUBIT 3, Invitrogen) at Ex/Em of: ~571/585 nm. Obtained values were normalized to total protein concentration.

#### Detection of oxidized proteins

Oxidized proteins containing carbonyl groups were detected using the OxyBlot Protein Oxidation Detection Kit (Sigma-Aldrich: cat#S7150). Briefly, wild type and Δ*vxr*AB mutant were grown to exponential growth phase and harvested at different times after PenG treatment. 0.5 mM H_2_O_2_ treated cells were used as a positive protein oxidation control with oxidized standard marker (_OX_M). The protein sample (10µg/well) was incubated with dinitrophenylhydrazine (DNPH) (derivatization reaction) for 15 min, followed by neutralization with a solution containing glycerol and β-mercaptoethanol. Both DNP-derivatized or nonderivatized (for negative control) protein samples were electrophoresed on a 10% SDS-PAGE and then transferred to a nitrocellulose membrane. After blocking, the membrane was incubated for 1 h with a rabbit antibody against DNP (cat# D9656), followed by incubation with secondary antibody: goat anti-rabbit IgG (Li-COR, IRDye® 800CW cat# 926-32211) for 1 h at room temperature as recommended by the manufacturer. The signals were scanned on an Odyssey CLx imaging device (LI-COR Biosciences) and visualized using image studio lite software.

#### Microscopy

All images were taken on a Leica MDi8 microscope (Leica Microsystems, GmbH, Wetzlar, Germany) with a PECON Temp-Controller 2000-1 (Erbach, Germany), heated stage at 37 °C for growth experiments, or room temperature. Time-lapse microscopy was performed by imaging frames five minutes apart and data were analyzed by ImageJ program. The msfGFP expressing cells were imaged at 485_Ex_/530_Em_ for two second exposure time. Collected images were processed for quantification in ImageJ by subtracting background and adjusting brightness/contrast uniformly across all fluorescent images. Representative images were taken at least three independent experiments.

### *In vivo* mouse experiments with *V. cholerae*

All mouse-related protocols, sample, and data collection were reviewed and approved by the Cornell University Institutional Animal Care and Use Committee (IACUC), protocol number 2016-0088. Female 7-week-old BALB/c mice were acquired from the Jackson Lab. All mice were housed in groups of up to five animals (upon receipt) or individually (upon completion of Sm pre-treatment) under standard barrier conditions in individually ventilated cages with autoclaved food and autoclaved water *ad libitum* and monitored in accordance with the rules of the Weill Hall Barrier Animal Facility. For experiments, animals between 8-9 weeks of age were given Streptomycin (Sm: Sigma, cat# 1623003) in their drinking water (5 mg/ml) for 48 hours and then orally inoculated with ca 10^10^ Sm-resistant *V. cholerae* EI Tor strain N16961. 24 h p.i., mice in AMX group received oral gavage of Amoxicillin 50 mg/kg (AMX: Sigma, cat# 1031503), mice in DFOA group received oral gavage of Deferoxamine 100 µM 200ul (DFOA: Sigma, cat# 1166003), mice in AMX+DFOA received both AMX and DFOA, mice in the Vc group received water only. Mice were euthanized 24 hrs post treatment. Fecal samples were collected at 6 time points (e.g. 0, 2, 4, 6, 8, and 10 hrs) and processed to monitor the level of *V. cholerae* excretion. For the challenge of mice, bacteria were grown overnight and harvested by centrifugation (x 2717g for 20 min), and washed in phosphate-buffered saline (PBS: Thomas Scientific, cat# C999G06) at the same speed and immediately administered to individual mice in PBS (ca 10^10^ *V. cholerae*). To enumerate *V*. *cholerae* cells in mouse feces, fresh fecal pellets were collected from an individual mouse at the indicated time points p.i. and weighed before resuspending into 1 mL PBS, homogenized by vortexing and pipetting. The sample homogenates were centrifuged at low speed (x 480g for 30 sec) and the resulting supernatant was plated in serial dilutions onto agar plates containing Sm (200 μg/mL) for CFU counting. The number of bacterial CFU was normalized per gram of feces (CFU/mg).

## Notes

### Competing Interest Statement

The authors have declared no competing interest.

## References

1. Spellberg, B. The future of antibiotics. Crit Care 18, 228 (2014).

2. Fair, R.J. & Tor, Y. Antibiotics and bacterial resistance in the 21st century. Perspect Medicin Chem 6, 25–64 (2014).

3. Lewis, K. Persister cells: molecular mechanisms related to antibiotic tolerance. Handb Exp Pharmacol, 121–133 (2012).

4. Lewis, K. Persister cells. Annu Rev Microbiol 64, 357–372 (2010).

5. Lewis, K. Persister cells, dormancy and infectious disease. Nat Rev Microbiol 5, 48–56 (2007).

6. El-Halfawy, O.M. & Valvano, M.A. Antimicrobial heteroresistance: an emerging field in need of clarity. Clin Microbiol Rev 28, 191–207 (2015).

7. Wilmaerts, D., Windels, E.M., Verstraeten, N. & Michiels, J. General Mechanisms Leading to Persister Formation and Awakening. Trends Genet 35, 401–411 (2019).

8. Balaban, N.Q., et al. Definitions and guidelines for research on antibiotic persistence. Nat Rev Microbiol (2019).

9. Kitano, K., Tuomanen, E. & Tomasz, A. Transglycosylase and endopeptidase participate in the degradation of murein during autolysis of Escherichia coli. J Bacteriol 167, 759–765 (1986).

10. Flores-Kim, J., Dobihal, G.S., Fenton, A., Rudner, D.Z. & Bernhardt, T.G. A switch in surface polymer biogenesis triggers growth-phase-dependent and antibiotic-induced bacteriolysis. Elife 8(2019).

11. Dörr, T., Davis, B.M. & Waldor, M.K. Endopeptidase-mediated beta lactam tolerance. PLoS Pathog 11, e1004850 (2015).

12. Cho, H., Uehara, T. & Bernhardt, T.G. Beta-lactam antibiotics induce a lethal malfunctioning of the bacterial cell wall synthesis machinery. Cell 159, 1300–1311 (2014).

13. Monahan, L.G., et al. Rapid conversion of Pseudomonas aeruginosa to a spherical cell morphotype facilitates tolerance to carbapenems and penicillins but increases susceptibility to antimicrobial peptides. Antimicrob Agents Chemother 58, 1956–1962 (2014).

14. Cross, T., et al. Spheroplast-mediated carbapenem tolerance in Gram-negative pathogens Antimicrob Agents Chemother (2019).

15. Errington, J., Mickiewicz, K., Kawai, Y. & Wu, L.J. L-form bacteria, chronic diseases and the origins of life. Philos Trans R Soc Lond B Biol Sci 371(2016).

16. Errington, J. Cell wall-deficient, L-form bacteria in the 21st century: a personal perspective. Biochem Soc Trans 45, 287–295 (2017).

17. Dörr, T., et al. A cell wall damage response mediated by a sensor kinase/response regulator pair enables beta-lactam tolerance. Proc Natl Acad Sci U S A 113, 404–409 (2016).

18. Dörr, T., Lewis, K. & Vulic, M. SOS response induces persistence to fluoroquinolones in Escherichia coli. PLoS Genet 5, e1000760 (2009).

19. Raivio, T.L. Everything old is new again: an update on current research on the Cpx envelope stress response. Biochim Biophys Acta 1843, 1529–1541 (2014).

20. Helmann, J.D. Bacillus subtilis extracytoplasmic function (ECF) sigma factors and defense of the cell envelope. Curr Opin Microbiol 30, 122–132 (2016).

21. Radeck, J., Fritz, G. & Mascher, T. The cell envelope stress response of Bacillus subtilis: from static signaling devices to dynamic regulatory network. Curr Genet 63, 79–90 (2017).

22. Wall, E., Majdalani, N. & Gottesman, S. The Complex Rcs Regulatory Cascade. Annu Rev Microbiol 72, 111–139 (2018).

23. Guest, R.L. & Raivio, T.L. Role of the Gram-Negative Envelope Stress Response in the Presence of Antimicrobial Agents. Trends Microbiol 24, 377–390 (2016).

24. Bhagirath, A.Y., et al. Two Component Regulatory Systems and Antibiotic Resistance in Gram-Negative Pathogens. Int J Mol Sci 20(2019).

25. Bury-Mone, S., et al. Global analysis of extracytoplasmic stress signaling in Escherichia coli. PLoS Genet 5, e1000651 (2009).

26. Cheng, A.T., Ottemann, K.M. & Yildiz, F.H. Vibrio cholerae Response Regulator VxrB Controls Colonization and Regulates the Type VI Secretion System. PLoS Pathog 11, e1004933 (2015).

27. Teschler, J.K., Cheng, A.T. & Yildiz, F.H. The Two-Component Signal Transduction System VxrAB Positively Regulates Vibrio cholerae Biofilm Formation. J Bacteriol 199(2017).

28. Dwyer, D.J., Kohanski, M.A., Hayete, B. & Collins, J.J. Gyrase inhibitors induce an oxidative damage cellular death pathway in Escherichia coli. Mol Syst Biol 3, 91 (2007).

29. Kohanski, M.A., Dwyer, D.J., Hayete, B., Lawrence, C.A. & Collins, J.J. A common mechanism of cellular death induced by bactericidal antibiotics. Cell 130, 797–810 (2007).

30. Kohanski, M.A., Dwyer, D.J., Wierzbowski, J., Cottarel, G. & Collins, J.J. Mistranslation of membrane proteins and two-component system activation trigger antibiotic-mediated cell death. Cell 135, 679–690 (2008).

31. Dwyer, D.J., Kohanski, M.A. & Collins, J.J. Role of reactive oxygen species in antibiotic action and resistance. Curr Opin Microbiol 12, 482–489 (2009).

32. Imlay, J.A. Diagnosing oxidative stress in bacteria: not as easy as you might think. Curr Opin Microbiol 24, 124–131 (2015).

33. Liu, Y. & Imlay, J.A. Cell death from antibiotics without the involvement of reactive oxygen species. Science 339, 1210–1213 (2013).

34. Keren, I., Wu, Y., Inocencio, J., Mulcahy, L.R. & Lewis, K. Killing by bactericidal antibiotics does not depend on reactive oxygen species. Science 339, 1213–1216 (2013).

35. Hong, Y., Zeng, J., Wang, X., Drlica, K. & Zhao, X. Post-stress bacterial cell death mediated by reactive oxygen species. Proc Natl Acad Sci U S A 116, 10064–10071 (2019).

36. Dwyer, D.J., Collins, J.J. & Walker, G.C. Unraveling the physiological complexities of antibiotic lethality. Annu Rev Pharmacol Toxicol 55, 313–332 (2015).

37. Dwyer, D.J., et al. Antibiotics induce redox-related physiological alterations as part of their lethality. Proc Natl Acad Sci U S A 111, E2100–2109 (2014).

38. Jacoby, G.A. AmpC beta-lactamases. Clin Microbiol Rev 22, 161–182, Table of Contents (2009).

39. Kanehisa, M. & Goto, S. KEGG: kyoto encyclopedia of genes and genomes. Nucleic Acids Res 28, 27–30 (2000).

40. Bailey, T.L., et al. MEME SUITE: tools for motif discovery and searching. Nucleic Acids Res 37, W202–208 (2009).

41. Dörr, T., et al. Differential requirement for PBP1a and PBP1b in in vivo and in vitro fitness of Vibrio cholerae. Infect Immun 82, 2115–2124 (2014).

42. Dörr, T., et al. A novel peptidoglycan binding protein crucial for PBP1A-mediated cell wall biogenesis in Vibrio cholerae. PLoS Genet 10, e1004433 (2014).

43. Ruiz, N. Lipid Flippases for Bacterial Peptidoglycan Biosynthesis. Lipid Insights 8, 21–31 (2015).

44. Teo, A.C. & Roper, D.I. Core Steps of Membrane-Bound Peptidoglycan Biosynthesis: Recent Advances, Insight and Opportunities. Antibiotics (Basel) 4, 495–520 (2015).

45. Davies, B.W., Bogard, R.W. & Mekalanos, J.J. Mapping the regulon of Vibrio cholerae ferric uptake regulator expands its known network of gene regulation. Proc Natl Acad Sci U S A 108, 12467–12472 (2011).

46. White, J.R. & Yeowell, H.N. Iron enhances the bactericidal action of streptonigrin. Biochem Biophys Res Commun 106, 407–411 (1982).

47. Yeowell, H.N. & White, J.R. Iron requirement in the bactericidal mechanism of streptonigrin. Antimicrob Agents Chemother 22, 961–968 (1982).

48. Yeom, J., Imlay, J.A. & Park, W. Iron homeostasis affects antibiotic-mediated cell death in Pseudomonas species. J Biol Chem 285, 22689–22695 (2010).

49. Leger, L., et al. beta-Lactam Exposure Triggers Reactive Oxygen Species Formation in Enterococcus faecalis via the Respiratory Chain Component DMK. Cell Rep 29, 2184–2191 e2183 (2019).

50. Seo, S.W., et al. Deciphering Fur transcriptional regulatory network highlights its complex role beyond iron metabolism in Escherichia coli. Nat Commun 5, 4910 (2014).

51. Masse, E. & Gottesman, S. A small RNA regulates the expression of genes involved in iron metabolism in Escherichia coli. Proc Natl Acad Sci U S A 99, 4620–4625 (2002).

52. Kehres, D.G., Janakiraman, A., Slauch, J.M. & Maguire, M.E. Regulation of Salmonella enterica serovar Typhimurium mntH transcription by H(2)O(2), Fe(2+), and Mn(2+). J Bacteriol 184, 3151–3158 (2002).

53. Martin, J.E., Waters, L.S., Storz, G. & Imlay, J.A. The Escherichia coli small protein MntS and exporter MntP optimize the intracellular concentration of manganese. PLoS Genet 11, e1004977 (2015).

54. Ikeda, J.S., Janakiraman, A., Kehres, D.G., Maguire, M.E. & Slauch, J.M. Transcriptional regulation of sitABCD of Salmonella enterica serovar Typhimurium by MntR and Fur. J Bacteriol 187, 912–922 (2005).

55. Fisher, C.R., Wyckoff, E.E., Peng, E.D. & Payne, S.M. Identification and Characterization of a Putative Manganese Export Protein in Vibrio cholerae. J Bacteriol 198, 2810–2817 (2016).

56. Wyckoff, E.E. & Payne, S.M. The Vibrio cholerae VctPDGC system transports catechol siderophores and a siderophore-free iron ligand. Mol Microbiol 81, 1446–1458 (2011).

57. Zeng, L., et al. Deferoxamine therapy for intracerebral hemorrhage: A systematic review. PLoS One 13, e0193615 (2018).

58. Fisher, S.A., et al. Desferrioxamine mesylate for managing transfusional iron overload in people with transfusion-dependent thalassaemia. Cochrane Database Syst Rev, CD004450 (2013).

59. Goodwin, J.F. & Whitten, C.F. Chelation of Ferrous Sulphate Solutions by Desferrioxamine B. Nature 205, 281–283 (1965).

60. Miyajima, H., et al. Use of desferrioxamine in the treatment of aceruloplasminemia. Ann Neurol 41, 404–407 (1997).

61. Richardson, D., Ponka, P. & Baker, E. The effect of the iron(III) chelator, desferrioxamine, on iron and transferrin uptake by the human malignant melanoma cell. Cancer Res 54, 685–689 (1994).

62. Imlay, J.A. The molecular mechanisms and physiological consequences of oxidative stress: lessons from a model bacterium. Nat Rev Microbiol 11, 443–454 (2013).

63. Imlay, J.A. The mismetallation of enzymes during oxidative stress. J Biol Chem 289, 28121–28128 (2014).

64. Xia, X., et al. OxyR-activated expression of Dps is important for Vibrio cholerae oxidative stress resistance and pathogenesis. PLoS One 12, e0171201 (2017).

65. Wang, H., et al. Catalases promote resistance of oxidative stress in Vibrio cholerae. PLoS One 7, e53383 (2012).

66. Wang, H., et al. OxyR2 Modulates OxyR1 Activity and Vibrio cholerae Oxidative Stress Response. Infect Immun 85(2017).

67. Storz, G., Tartaglia, L.A. & Ames, B.N. The OxyR regulon. Antonie Van Leeuwenhoek 58, 157–161 (1990).

68. Johnson, L.A. & Hug, L.A. Distribution of reactive oxygen species defense mechanisms across domain bacteria. Free Radic Biol Med (2019).

69. Muras, V., Dogaru-Kinn, P., Minato, Y., Hase, C.C. & Steuber, J. The Na+-Translocating NADH:Quinone Oxidoreductase Enhances Oxidative Stress in the Cytoplasm of Vibrio cholerae. J Bacteriol 198, 2307–2317 (2016).

70. Steuber, J., et al. The structure of Na(+)-translocating of NADH:ubiquinone oxidoreductase of Vibrio cholerae: implications on coupling between electron transfer and Na(+) transport. Biol Chem 396, 1015–1030 (2015).

71. Tuomanen, E., Cozens, R., Tosch, W., Zak, O. & Tomasz, A. The rate of killing of Escherichia coli by beta-lactam antibiotics is strictly proportional to the rate of bacterial growth. J Gen Microbiol 132, 1297–1304 (1986).

72. Weaver, A.I., et al. Genetic Determinants of Penicillin Tolerance in Vibrio cholerae. Antimicrob Agents Chemother 62(2018).

73. Kawai, Y., et al. Cell growth of wall-free L-form bacteria is limited by oxidative damage. Curr Biol 25, 1613–1618 (2015).

74. Kawai, Y., et al. Crucial role for central carbon metabolism in the bacterial L-form switch and killing by beta-lactam antibiotics. Nat Microbiol (2019).

75. Schnell, S. & Steinman, H.M. Function and stationary-phase induction of periplasmic copper-zinc superoxide dismutase and catalase/peroxidase in Caulobacter crescentus. J Bacteriol 177, 5924–5929 (1995).

76. Longo, V.D., Gralla, E.B. & Valentine, J.S. Superoxide dismutase activity is essential for stationary phase survival in Saccharomyces cerevisiae. Mitochondrial production of toxic oxygen species in vivo. J Biol Chem 271, 12275–12280 (1996).

77. Gort, A.S., Ferber, D.M. & Imlay, J.A. The regulation and role of the periplasmic copper, zinc superoxide dismutase of Escherichia coli. Mol Microbiol 32, 179–191 (1999).

78. Martins, D., et al. Superoxide dismutase activity confers (p)ppGpp-mediated antibiotic tolerance to stationary-phase Pseudomonas aeruginosa. Proc Natl Acad Sci U S A 115, 9797–9802 (2018).

79. Haugan, M.S., Charbon, G., Frimodt-Moller, N. & Lobner-Olesen, A. Chromosome replication as a measure of bacterial growth rate during Escherichia coli infection in the mouse peritonitis model. Sci Rep 8, 14961 (2018).

80. Keren, I., Kaldalu, N., Spoering, A., Wang, Y. & Lewis, K. Persister cells and tolerance to antimicrobials. FEMS Microbiol Lett 230, 13–18 (2004).

81. Sharma, B., Brown, A.V., Matluck, N.E., Hu, L.T. & Lewis, K. Borrelia burgdorferi, the Causative Agent of Lyme Disease, Forms Drug-Tolerant Persister Cells. Antimicrob Agents Chemother 59, 4616–4624 (2015).

82. Brynildsen, M.P., Winkler, J.A., Spina, C.S., MacDonald, I.C. & Collins, J.J. Potentiating antibacterial activity by predictably enhancing endogenous microbial ROS production. Nat Biotechnol 31, 160–165 (2013).

83. Gutierrez, A., et al. beta-Lactam antibiotics promote bacterial mutagenesis via an RpoS-mediated reduction in replication fidelity. Nat Commun 4, 1610 (2013).

84. Sheldon, J.R. & Skaar, E.P. Metals as phagocyte antimicrobial effectors. Curr Opin Immunol 60, 1–9 (2019).

85. Yang, J.H., et al. A White-Box Machine Learning Approach for Revealing Antibiotic Mechanisms of Action. Cell 177, 1649–1661 e1649 (2019).

86. Radlinski, L. & Conlon, B.P. Antibiotic efficacy in the complex infection environment. Curr Opin Microbiol 42, 19–24 (2018).

87. Yang, J.H., Bening, S.C. & Collins, J.J. Antibiotic efficacy-context matters. Curr Opin Microbiol 39, 73–80 (2017).

88. Stokes, J.M., Lopatkin, A.J., Lobritz, M.A. & Collins, J.J. Bacterial Metabolism and Antibiotic Efficacy. Cell Metab (2019).

89. Papenfort, K., Forstner, K.U., Cong, J.P., Sharma, C.M. & Bassler, B.L. Differential RNA-seq of Vibrio cholerae identifies the VqmR small RNA as a regulator of biofilm formation. Proc Natl Acad Sci U S A 112, E766–775 (2015).

90. Krin, E., et al. Expansion of the SOS regulon of Vibrio cholerae through extensive transcriptome analysis and experimental validation. BMC Genomics 19, 373 (2018).

91. Donnenberg, M.S. & Kaper, J.B. Construction of an eae deletion mutant of enteropathogenic Escherichia coli by using a positive-selection suicide vector. Infect Immun 59, 4310–4317 (1991).

92. Dalia, A.B. Natural Cotransformation and Multiplex Genome Editing by Natural Transformation (MuGENT) of Vibrio cholerae. Methods Mol Biol 1839, 53–64 (2018).

93. Hubbard, T.P., et al. A live vaccine rapidly protects against cholera in an infant rabbit model. Sci Transl Med 10(2018).

94. Gibson, D.G., et al. Enzymatic assembly of DNA molecules up to several hundred kilobases. Nat Methods 6, 343–345 (2009).

95. Dörr, T., Cava, F., Lam, H., Davis, B.M. & Waldor, M.K. Substrate specificity of an elongation-specific peptidoglycan endopeptidase and its implications for cell wall architecture and growth of Vibrio cholerae. Mol Microbiol 89, 949–962 (2013).

96. Beauchamp, C. & Fridovich, I. Superoxide dismutase: improved assays and an assay applicable to acrylamide gels. Anal Biochem 44, 276–287 (1971).

