## Supplementary Informations for "A multifaceted cellular damage repair and prevention pathway promotes high level tolerance to β-lactam antibiotics"


Jung-Ho Shin[1, 2], Donghui Choe[4, 5], Brett Ransegnola[1, 2], Hye-Rim Hong[1, 2], Ikenna Onyekwere[1, 2], Trever Cross[1,2], Hector Loyola Irizarry[9], Qian Juan Shi[9], Byung-Kwan Cho[4, 5, 6], Lars F. Westblade[7,8], Ilana Lauren Brito[9], and Tobias Dörr[1, 2, 3a]

[1] Weill Institute for Cell and Molecular Biology, Cornell, University, Ithaca, NY 14853, USA

[2] Department of Microbiology, Cornell University, Ithaca NY 14853, USA

[3] Cornell Institute of Host-Microbe Interactions and Disease, Cornell University, Ithaca NY 14853, USA

[4] Department of Biological Sciences, Korea Advanced Institute of Science and Technology, Daejeon 34141, Republic of Korea.

[5] KI for the BioCentury, Korea Advanced Institute of Science and Technology, Daejeon 34141, Republic of Korea.

[6] Intelligent Synthetic Biology Center, Daejeon 34141, Republic of Korea.

[7] Department of Pathology and Laboratory Medicine, Weill Cornell Medicine, New York, NY, USA.

[8] Division of Infectious Diseases, Department of Medicine, Weill Cornell Medicine, New York, NY, USA

[9] Meinig School of Biomedical Engineering, Cornell University, Ithaca, NY, 14850, USA

### Supplementary Tables

**Supplementary Table 1** Strains and plasmids used in this study.

| **Strain /plasmid** | | **Relevant description** | **Reference /source** |
|---|---|---|---|
| ***V. cholerae* strains** | | | |
| N16961 | | Wild-type O1 Inaba; $Str^r$ | Lab stock |
| | TDW933 | *ΔvxrAB* | Dorr et al. 2016 (PNAS) |
| | TDW943 | *ΔvxrA* | Dorr et al. 2016 (PNAS) |
| | TDW929 | vxrB::vxrB-6xHis | Dorr et al. 2016 (PNAS) |
| | DL722 | *murJ ΔVBS* (vxrB binding site) | This study |
| | TDW453 | $P_{IPTG}$-pbp1a *Δpbp1a Δpbp1B::kan* | Dorr et al. 2014 (Plos Genetics) |
| | DL800 | $P_{rhyB}$+msfGFP | This study |
| | DL632 | *wigK::3xFlag-WigK Δfur* | This study |
| | DL636 | *wigK::3xFlag-WigK ΔsodB* | This study |
| | DL744 | *ΔoxyR1ΔkatGΔkatB* | This study |
| | JHF900 | *ΔvctPDGC* | This study |
| | JHF901 | *ΔfeoABC* | This study |
| | JHF902 | *ΔfbpABC* | This study |
| | JHF903 | *Δvc2211* | This study |
| | JHF904 | *Δvc2212* | This study |
| | JHF905 | *ΔtonB2exbB2* | This study |
| | JHF906 | *ΔhutA* | This study |
| | JHF907 | *ΔvxrAB ΔvctPDGC* | This study |
| | JHF908 | *ΔvxrAB ΔfeoABC* | This study |
| | JHF909 | *ΔvxrAB ΔfbpABC* | This study |
| | JHF910 | *ΔvxrAB Δvc2211* | This study |
| | JHF911 | *ΔvxrAB Δvc2212* | This study |
| | JHF912 | *ΔvxrAB ΔtonB2exbB2* | This study |
| | JHF913 | *ΔvxrAB ΔhutA* | This study |
| | JHF914 | *ΔvctPDGC ΔfeoABC* | This study |
| | JHF915 | *ΔvctPDGC ΔfbpABC* | This study |
| | JHF916 | *ΔvctPDGC Δvc2211* | This study |
| | JHF917 | *ΔfeoABC ΔfbpABC* | This study |
| | JHF918 | *ΔtonB2exbB2 ΔhutA* | This study |
| | JHF919 | *ΔvxrAB ΔvctPDGC ΔfeoABC* | This study |
| | JHF920 | *ΔvxrAB ΔvctPDGC ΔfbpABC* | This study |
| | JHF921 | *ΔvxrAB ΔvctPDGC Δvc2211* | This study |
| | JHF922 | *ΔvxrAB ΔfeoABC ΔfbpABC* | This study |

| | | | |
|---|---|---|---|
| | JHF923 | *ΔvxrAB ΔtonB2exbB2 ΔhutA* | This study |
| | JHF924 | *ΔvctPDGC ΔfeoABC ΔfbpABC* | This study |
| | JHF925 | *ΔvctPDGC ΔtonB2exbB2 ΔhutA* | This study |
| | JHF926 | *Δvc2211 ΔtonB2exbB2 ΔhutA* | This study |
| | JHF927 | *ΔvctPDGC Δvc2211 ΔhutA* | This study |
| | JHF928 | *ΔvxrAB ΔvctPDGC ΔfeoABC ΔfbpABC* | This study |
| | JHF929 | *ΔvxrAB ΔvctPDGC ΔtonB2exbB2 ΔhutA* | This study |
| | JHF930 | *ΔvxrAB Δvc2211 ΔtonB2exbB2 ΔhutA* | This study |
| | JHF931 | *ΔvxrAB ΔvctPDGC Δvc2211 ΔhutA* | This study |
| E7946 | SAD30 | Wild-type O1 Inaba; Str$^r$ | Ankur Dalia |
| | DL801 | *ΔnqrA* | This study |
| | DL802 | *ΔubiA* | This study |
| | DL803 | *ΔvxrAB::chl ΔnqrA* | This study |
| | DL804 | *ΔvxrAB:: chl ΔubiA* | This study |
| ***E. coli* strains** | | | |
| DH5α λpir | | F- endA1 glnV44 thi-1 recA1 relA1 gyrA96 deoR nupG Φ80d*lacZ*ΔM15 Δ(*lacZYA-argF*)U169, hsdR17(rK- mK+), λpir | Lab stock |
| | DJ-ED2 | pET15b | This study |
| | DJ-ED3 | pET15b::VxrB | This study |
| SM10 λpir | | Δ(ara-leu)7697 ΔlacX74 ΔphoA PvuII phoR araD139 ahpC galE galK rpsL (DE3) F'[lac+ lacIq pro] gor522::Tn10 trxB pLysSRARE (CamR, StrR, TetR) | Lab stock |
| | DL711 | pCVD442 ΔoxyR1 | This study |
| | DL730 | pCVD442 ΔkatB | This study |
| | DL736 | pCVD442 ΔkatG | This study |
| | DL620 | pCVD442 Δvc2106 (fur) | This study |
| | DL622 | pCVD442 Δvc2045 (sodB) | This study |
| | DL805 | pTD101 PryhB::msfGFP | This study |
| | DJES900 | pCVD442 *ΔvctPDGC* | This study |
| | DJES901 | pCVD442 *ΔfeoABC* | This study |

| | | | |
|---|---|---|---|
| | DJES902 | pCVD442 *ΔfbpABC* | This study |
| | DJES903 | pCVD442 *Δvc2211* | This study |
| | DJES904 | pCVD442 *Δvc2212* | This study |
| | DJES905 | pCVD442 *ΔtonB2exbB2* | This study |
| | DJES906 | pCVD442 *ΔhutA* | This study |
| BL21(DE3/pLysS) | DJ-EB1 | B F– ompT gal dcm lon hsdSB(rB– mB–) λ(DE3 [lacI lacUV5-T7p07 ind1 sam7 nin5]) [malB+]K-12(λS) pLysS[T7p20 orip15A](CmR) | Lab stock |
| | DJ-EB2 | pET15b | This study |
| | DJ-EB3 | pET15b::VxrB | This study |
| **Plasmids** | | | |
| pET15b | pDJ1 | Over expression vector in *E.coli* | Novagen |
| pCVD442 | | $carb^R$, ori R6K gamma, *sacB* | PMID: 15466024 |

**Supplementary Table 2** Oligonucleotides used in this study.

| | | Primer sequence (5’ to 3’) | Description |
|---|---|---|---|
| EMSA | | | |
| | TD-JHS201 | AAACCCTTTTTTAGTTCTTTTAGTTTCTTTACAATTTTCT GATAATCGAG | VC2635 FP1P50 For |
| | TD-JHS202 | CTCGATTATCAGAAAATTGTAAAGAAACTAAAAGAACTA AAAAAGGGTTT | VC2635 FP1P50 Rev |
| | TD-JHS203 | TCGAGCGTGTGATTATCGGTGAAAATATCAGCGGTAAC GAGCAACTCCTC | VC2635 S2P50 For |
| | TD-JHS204 | GAGGAGTTGCTCGTTACCGCTGATATTTTCACCGATAA TCACACGCTCGA | VC2635 S2P50 Rev |
| | TD-JHS205 | ATTCTATCAGCATTTTTTTACCAATGCTAATACTTATCTG ATCTCACTAT | VC0680 MP50 For |
| | TD-JHS206 | ATAGTGAGATCAGATAAGTATTAGCATTGGTAAAAAAAT GCTGATAGAAT | VC0680 MP50 Rev |
| | TD-JHS207 | CAATATCATGCAAATGCATGGTTCTGACCAAAACTTTAC GTAACTATTAC | VC1655 M50 For |
| | TD-JHS208 | GTAATAGTTACGTAAAGTTTTGGTCAGAACCATGCATTT GCATGATATTG | VC1655 M50 Rev |
| | TD-JHS209 | ACTGAGTTTGGGTCAACCCTAGGTCAATTCTGGTTAGG CGGCGCACAAAC | VC0370 M50 For |
| | TD-JHS210 | GTTTGTGCGCCGCCTAACCAGAATTGACCTAGGGTTGA CCCAAACTCAGT | VC0370 M50 Rev |
| | TD-JHS211 | AACGCGTCATCTTTTCAACTTCTCACATTAAGTTGTTCT AAATTCAACGA | VC0969 M50 For |
| | TD-JHS212 | TCGTTGAATTTAGAACAACTTAATGTGAGAAGTTGAAAA GATGACGCGTT | VC0969 M50 Rev |
| | TD-JHS213 | TAAAACCGCCCTATTACTCACTGTTGTTGGTGCCGTTCT GCTCACTGGCT | VCA0594 FP1P50 For |
| | TD-JHS214 | AGCCAGTGAGCAGAACGGCACCAACAACAGTGAGTAA TAGGGCGGTTTTA | VCA0594 FP1P50 Rev |
| | TD-JHS215 | TTTGACAAATCTGACATATAGCAAAAAATTAATTTTGTA GAATACTGATA | VCA0594 S2P50 For |
| | TD-JHS216 | TATCAGTATTCTACAAAATTAATTTTTTGCTATATGTCAG ATTTGTCAAA | VCA0594 S2P50 Rev |
| | TD-JHS217 | AGAGCCTTTCGGTTATTATTTTGTCACGTCTTTGCTAAA AATGAGAAAAG | VC2213 M50 For |
| | TD-JHS218 | CTTTTCTCATTTTTAGCAAAGACGTGACAAAATAATAAC CGAAAGGCTCT | VC2213 M50 Rev |
| | TD-JHS219 | GCTATTCGGTTAAAAAAACCGCGCAATTTGACCCAAGC TTAACTTAACGA | VC2686 FP1P50 For |
| | TD-JHS220 | TCGTTAAGTTAAGCTTGGGTCAAATTGCGCGGTTTTTTT AACCGAATAGC | VC2686 FP1P50 Rev |
| | TD-JHS221 | TTAACCAAGTTAGAGTATTGACGTCAGATTTTTACCAAA AAACTTATTTC | VC2037 M50 For |
| | TD-JHS222 | GAAATAAGTTTTTTGGTAAAAATCTGACGTCAATACTCT AACTTGGTTAA | VC2037 M50 Rev |
| | TD-JHS223 | AAAGATCATTTGTCGCTGAGAACTTGACCTCATCCCTA CAAACGTCGAAC | VC1272 FP1P50 For |
| | TD-JHS224 | GTTCGACGTTTGTAGGGATGAGGTCAAGTTCTCAGCGA CAAATGATCTTT | VC1272 FP1P50 Rev |
| | TD-JHS039 | CCATGGCGGCCGCGGGAATTCGCAGCACTTCTGTGCT CGCTCAAGCG | VC2143 T22S1 BGM For |
| | TD-JHS040 | GAAGTTTGTTGTGCATTGGTTGCACC | VC2143 S1 BGM Rev |
| | TD-JHS041 | CCATGGCGGCCGCGGGAATTCGGTTGCTAATACAACA TTGAGCCTTGG | VC2188 T22S1 BGM For |
| | TD-JHS042 | GAGAGGCGTTCCATGGAGGTGTTAAGC | VC2188 S1 BGM Rev |
| | TD-JHS043 | CCATGGCGGCCGCGGGAATTCGGGTGCGTTGTAACCC TGGTATCATC | VC2200 T22S1 BGM For |
| | TD-JHS044 | GGTGGCAATCACCTCAGAGTTGCGCTC | VC2200 S1 BGM Rev |
| | TD-JHS045 | CCATGGCGGCCGCGGGAATTCGGAGGTCAATTTAGGA CATTGCAGGAC | VC2203 T22S1 BGM For |
| | TD-JHS046 | CACAGAAAGCTCTACACTTAGTTATG | VC2203 S1 BGM Rev |

| | | | |
|---|---|---|---|
| | TD-JHS047 | CCATGGCGGCCGCGGGAATTCGGGCTGATGGAGACTTAACAGTGTCATG | VCA0139 T22S1 BGM For |
| | TD-JHS048 | GCTTTGTGGTTGGTATCGATAGACGC | VCA0139 S1 BGM Rev |
| | TD-JHS049 | CCATGGCGGCCGCGGGAATTCGCTCGGTGCTTGAGCTCAACGTCAAAC | VC1962 T22S1 BGM For |
| | TD-JHS050 | CTACTTCCACTTTTGCGTCTTGTTGG | VC1962 S1 BGM Rev |
| | TD-JHS051 | CCATGGCGGCCGCGGGAATTCGCCTATTGGAAGCGGTAGCCATGGTG | VC0600 T22S1 BGM For |
| | TD-JHS052 | CGGGTAGACCAAATTGACAACGCGAG | VC0600 S1 BGM Rev |
| | TD-JHS053 | CCATGGCGGCCGCGGGAATTCGGGTTGCTTGAGCCAGTGAGTGATTGC | VC2409 T22S1 BGM For |
| | TD-JHS054 | GATACCATCCGGCTTGATCGCCAATCC | VC2409 S1 BGM Rev |
| | TD-JHS055 | CCATGGCGGCCGCGGGAATTCGGCAGATTTGTTATTTGCCAAGGGTC | VC0680 T22S1 BGM For |
| | TD-JHS056 | CACGAGAAATCAAAGTCATAGCAC | VC0680 S1 BGM Rev |
| Disruption/fusion | | | |
| | TDP1062 | CTGGTTGTGGTCGAGGATGAT | murJfwLarge |
| | TDP1064 | TTATTACTCGAGTGCGGCCGCATTAATTAGCATTGGTAAAAAAATGCTGATAGAAT | murJrev2 |
| | TDP1066 | TAATGCGGCCGCACTCGAGTAATAACTCACTATGACATTCTTATCCTTTCGAATCA | murJfwDown3 |
| | TDP1067 | GCCTAGTGCACGGATGATGTT | murJrevLarge |
| | TDP1078 | ACGTGCACGGCCAACCA | murJfwNEST |
| | TDP1079 | CCAAGACGCTTAAAGCTCTCTTTTT | murJrevNEST |
| | TDP597 | TACTCGTATGCCGATTTTAAAGGGGAT | vc1807fw |
| | TDP598 | TAGTCACCTCTATTGTTAACTTGTTCATAGAAG | vc1807rev |
| | TDP599 | CTTCTATGAACAAGTTAACAATAGAGGTGACTAGGTGGAGACATTACTGAGAAGTCTCAG | trimfw_vc1807 |
| | TDP600 | GTTTTAAATGTAACCATTAAAAAGAGGCGAGCCTCTTAACCCTTTTGCCAGATTTGGTAAC | trimrev_vc1807 |
| | TDP601 | GAGGCTCGCCTCTTTTTAATGGTTAC | vc1807downFW |
| | TDP602 | ATGTGGACAGGATTCTGGAAATTTTCAG | vc1807_rev |
| | TDP603 | AAACCCAATGCCAAGAGGGG | vc1807fw_INT |
| | TDP604 | AGACAGGACATAACGTTTTGTTGCT | vc1807rev_INT |
| | TDP1056 | ATATGTGATGGGTTAAAAAGGATCGATCCTACTTTGAACGGGTCGCCCG | oxyRfwpCVD |
| | TDP1057 | TCATCATTATTACTCGAGTGCGGCCGCATTAATCACGAATGTTCATAGCTTCCCC | oxyRrev |
| | TDP1058 | TAATGCGGCCGCACTCGAGTAATAATGACAGCAAAGCGAGTAAGACGGC | oxyRfw |
| | TDP1059 | AGAGCTCGATATCGCATGCGGTACCTCTAGAAAACGCTTTCTTAACGCGTGAAATG | oxyRrevpCVD |
| | TDP1097 | ATATGTGATGGGTTAAAAAGGATCGATCCTTGGGCTCGGCTTCATCGC | katGfwpCVD |
| | TDP1098 | TCATCATTATTACTCGAGTGCGGCCGCATTATGATTACTCCTTGCTTCGCATTATTCC | katGrev |
| | TDP1099 | TAATGCGGCCGCACTCGAGTAATAATGATAATCCATCTCACCATCCCTTTCAAGG | katGfw |
| | TDP1100 | AGAGCTCGATATCGCATGCGGTACCTCTAGCAACAACGTGCCCCTAATTTGAAGTTA | katGrevpCVD |
| | TDP1103 | ATATGTGATGGGTTAAAAAGGATCGATCCTGCCAGAGTAATAAGCTACTTTAGAAGAGC | katBfwpCVD |
| | TDP1104 | TCATCATTATTACTCGAGTGCGGCCGCATTAGGTTGCTCTCCAATGCGACC | katBrev |
| | TDP1105 | TAATGCGGCCGCACTCGAGTAATAATGATAACCGATAAAAATTTGCCCCGCAAGA | katBfw |

| | | | |
|---|---|---|---|
| | TDP1106 | AGAGCTCGATATCGCATGCGGTACCTCTAGAGATTGAC CGTACAACTCAGCA | katBrevpCVD |
| | TDP1152 | CAGGAAACAGACCATGGAATTCGAGCTCGGTACCCCC GAATCCACTTTGTTAAAGTGCTCTAA | ryhBfwpHL |
| | TDP1235 | AACAGTTCTTCACCTTTATGAATTCCCATGATCATGTGG TTAATAATAATAGTTCTCATTTAATAGTCAA | rhyBrevmsfGFP |
| | TDP1237 | ATGATCATGGGAATTCATAAAGGTGAAGAAC | msfGFPfw |
| | TDP1238 | CGCTCGGCAATGAATCGGGGATTGGTACCGCGGCCTT ATTTGTAGAGTTCATCCATGCCGTG | msfGFPrevpJL1 |
| | TDP1388 | CGCGTATACCGCAAACACACTG | vc2295fw_up |
| | TDP1389 | TCATTATTACTCGAGTGCGGCCGCATTAATTCGAGGCG CACTCCGTTT | vc2295rev_up |
| | TDP1390 | TAATGCGGCCGCACTCGAGTAATAATGATAATTTCCAT GGGCCTTAAAAAGTTTCTTGAA | vc2295fw_down |
| | TDP1391 | AGATACGAATACACCAGCGCCT | vc2295rev_down |
| | TDP1392 | ATGGGACCGGGGTTGCTG | vc2295_NEST_fw |
| | TDP1393 | CAGAACCAAAGCCAGTATAATCAGGGTA | vc2295_NEST_rev |
| | TDP1399 | TTAAGCCAGACCCGCTGATGC | vc0094fw_up |
| | TDP1401 | CGACATTACTGAGACTTCTCAGTAATGTCTCCACCGCC AGTAAGCGCGCGC | vc0094revTrim |
| | TDP1404 | TAAATTATAGTTACCAAATCTGGCAAAAGGGTTAATAAT CCCAGCTTAAGGCAAAAGAAAAGG | vc0094fw_trim |
| | TDP1403 | GTGTGCTTCCACATGCTCTTGAG | vc0094_rev_down |
| | TDP1405 | CCACGCCAGAGTCAGACCT | vc0094_fw_NEST |
| | TDP1406 | GTTATCGACGAGGATGCGGATCC | vc0094_rev_NEST |
| | TDP636 | GGTGGAGACATTACTGAGAAGTCTCAG | trimFW |
| | TDP637 | TTAACCCTTTTGCCAGATTTGGTAAC | trimREV |
| | TDP828 | ATATGTGATGGGTTAAAAAGGATCGATCCTGTATCGCT CATCATAAGCGTCTGG | vc2045fwpCVD |
| | TDP829 | TCATCATTATTACTCGAGTGCGGCCGCATTAGGGTAAT TCAAATGCCATTGCTCGA | vc2045REV |
| | TDP830 | TAATGCGGCCGCACTCGAGTAATAATGAGGGACTTCGT TGCTCAGAACCT | vc2045fw |
| | TDP831 | AGAGCTCGATATCGCATGCGGTACCTCTAGAGCGCTA GGTTGGTTCCC | vc2045revpCVD |
| | TDP834 | ATATGTGATGGGTTAAAAAGGATCGATCCTGTTTCAAAA ACCGCCGAAGCTT | vc2106fwpCVD |
| | TDP835 | TCATCATTATTACTCGAGTGCGGCCGCATTAGGTTATT GTCTGACATATACTTTCCTGTTG | vc2106rev |
| | TDP836 | TAATGCGGCCGCACTCGAGTAATAATGAGAAGAAATAA CCATAGGCTTTACGCTCT | vc2106fw |
| | TDP837 | AGAGCTCGATATCGCATGCGGTACCTCTAGGTAAAACC GTGCGTGTGCAAAATG | vc2106revpCVD |
| | TD-JHS394 | CCGGTGGCAGGCATGCGCCAAACCATTGCGGAAATAG AAGCCTTGATG | VCA0227 SphI F |
| | TD-JHS395 | CAACACTCCTCCCGGGTGCAACATTTTCTGCAACAAGA CAAAGCAC | VCA0227 XmaI R |
| | TD-JHS396 | GCCATGATTCTGGCCAAGCTTATC | VCA0227 Ver STOP 43 R |
| | TD-JHS397 | GGTAGTCGATGCATGCGCGCTACAAATGTGATGAATTT TGTTCAG | VC2078 SphI F |
| | TD-JHS398 | TCTGCCGTGCCCCGGGTTATTTTAGTCCCGCAAAGAGC GGATTTACG | VC2078 XmaI R |
| | TD-JHS399 | CGAGCATAGATACACTATGCGATTGGGATGAACAAACG TTGCCAAC | VC2078 Ver STOP 73R |
| | TD-JHS400 | ATTTTCCTCAATACGCATGCGGCTGCGGCGGCAGGGG CTATCGCTGCGC | VC0608 SphI F |
| | TD-JHS401 | GCTTTAAACACCCGGGTTCACAAATGGAAGCGGACGA CTCTCTGATTG | VC0608 XmaI R |
| | TD-JHS402 | CAGGTAACATAGTTTGGCAATCCAAGACGCTTGGTGAG AGC | VC0608 Ver STOP 42R |

| | | | |
|---|---|---|---|
| | TD-JHS403 | TAAATAAAACGCATGCTAGTAATAATAGAGGCTTTTGGC<br>TAGCACGAG | VC2211 SphI F |
| | TD-JHS404 | CACAACAAAACCCGGGTCAAGCTCATTGCTCTGTGCTC<br>GGTATTGTC | VC2211 XmaI R |
| | TD-JHS405 | CGCCTCTCAGCTGACTTAGAGGCGTTGTTTTGTTTCAG<br>TCCATTTG | VC2211 Ver STOP 55R |
| | TD-JHS406 | GTTGTGTTTTGCATGCAATCACGACCAACACGGGCATT<br>TCTTGCTCTTG | VC2212 SphI F |
| | TD-JHS407 | GACATCGTACCCCGGGGCGCAGAAACCGCTACACTCG<br>GTATCG | VC2212 XmaI R |
| | TD-JHS408 | CGGTAATCATCAAGATTCAATCTACAAAGGCCCATTCG<br>GGC | VC2212 Ver STOP 83R |
| | TD-JHS409 | TGCGGAAATCGCATGCTTGAACGAAGAAGTCAAAAATC<br>TTCAGGTG | VC1547 SphI F |
| | TD-JHS410 | CCAACACCAACCCGGGTTTTAGGGTGAGAATTCCCGG<br>CTTCCACTGTTC | VC1547 XmaI R |
| | TD-JHS411 | CGTAAGTCGTCACGGATAGGCAAGAGAGCAAGAGATA<br>ACTTAAGATTC | VC1547 Ver STOP 54R |
| | TD-JHS412 | TGAATGCGCAGCATGCTTTTACTGATGTTACCCAGTGC<br>TTCAACCATAG | VCA0576 SphI F |
| | TD-JHS413 | ACTCAATTACCCCGGGATTGAGTTTTTGAGCAAAGGAG<br>AGGATTTC | VCA0576 XmaI R |
| | TD-JHS414 | CTGGCTTTTTGTTATCGTTCCTAAACCACGGTAAAATGG<br>TGC | VCA0576 Ver STOP 55R |
| Gene block | | CCGGTGGCAGGCATGCGCCAAACCATTGCGGAAATAG<br>AAGCCTTGATGGTCGTAAAAAGCGTCGTGTTACCGTTT<br>AAAGCCTCTATTTGTCTCTCTGATGGTCAACAGTTTTGG<br>CTGGTTCGTTACAGTACCGATGGTCAACCGCCGACGG<br>TTTACCGCAAACCTTGGCAAGATGGAATCATCTTAGCC<br>TCCGAGCCCTTGGATGCTTGTCCTAATTGGTTATTGGT<br>AGAGCCACAAACCATTACGCATGTGCTTGGGGCTGAGT<br>GTCAGTCATACCTTATTTAAGCTATCCCCTTCCTACTTG<br>AAGCTGCAGCGGTGTTGGCTACGTTAGTTCTCCCCAAC<br>CACATAGTCTATCTATGTTCATGGGGATTGTGAGAGCC<br>ATCCTTGGCTCTCACCAAAGGCCAGCCGATGGCTGTG<br>CAAATTTGTTCCAGACAAATTTGTCACTCACTTGCCGCC<br>TACCTGCAACTCCAAGTGGTTTGGGTATGAATCAAATT<br>CATTAAATGAATCTTGTAGACCCTTGCTATTGATAAGAA<br>TTGGCGAGGGTTTGTGCATTTCTAAATTGTTCAACGATA<br>TCTTCCCTCACACAGTTGCATTGCCAAGGGCAGCAGTA<br>CAAGTAGGGGATAAGCTTGGCCAGAATCATGGCAATTA<br>ACCCGCCTGTTTTAATTCCACAAAAGCCTGCGGAGAGA<br>TACCAAAGGCTTTTTTAAACGCTGCGGTAAAGTTGGAA<br>GGGTGCAGGTAGCCTGCTTCATACGCGGCCTCTGTGA<br>TGGTGACCAAACCTCGCTCTAATTGCTGACGCGCCAGT<br>TGTAACCGTCGGTGCCGGATGTAGTGTGCAATCGTCAT<br>ATTCAGGCGAGTTTTAAATTTGCGTTGTAAGTTGGAAAC<br>GCTCATCGCATGGCGTTCTGCCAGTTGCTCTAAGCAAA<br>TCTCTTCATGCAAATGCTGATCGATATAGTGCAGCAACT<br>GATTAATCCATTGCTCAGATTGTCCCTCTTGGTGCTGA<br>GATAACGAGGTGGCGTGTTTAGGGCATTGCAACACGG<br>CGTCATGCACCAAGCGCCAAACCGTGGCTTCCCGAGC<br>AATATGCGCACTGATGGTGTGGATCACACGACGGTTGA<br>GCAGAGACTCAACGGCTTGCCAACTTTCCTCGTTCAGT<br>GAAAGTGGAACCAGTGCTTTGTCTTGTTGCAGAAAATG<br>TTGCACCCGGGAGGAGTGTTG | *vctPDGC* deletion |
| | | GGTAGTCGATGCATGCGCGCTACAAATGTGATGAATTT<br>TGTTCAGACAAACATAAGCTAACAAGCTTGTCGCTGAA<br>TCGGCTCAAGAAACAAATGCTATTGAAGTGAGTGATGG<br>TAATAGAGGTAAAACAGTAGGAAAAAAATATCGTGGTAT<br>AGCCGGGGGGACTATACCACGGGTGTCAACGACACCT<br>TCGCTTGCTGCCGGAGTGGTATGACATGGTCATACCAC<br>CTCTGGAATTCTTGTCTCTAGGAGGGATGTCCTTTCAA<br>CGAAATCACTATAATCCTATAGACTTATTTTACTTATAAA<br>ATTAACGACAACTTTACATAGAAAAACGACAGCAGCTTA<br>ATCATGTTTTTAAGGTGTTGATTTTAAATATGTAAATGTA<br>ATTTGTCGTTTTTTCTATCATGTTGATTTTATGCAGGAG<br>ACTTTTCTGACTTGGTCTTTGACGTTTTTGACGAAAAGT<br>TTTTGACAGAATTGACGTTGACTATTACCTAAGCAAACA | *feoABC* deletion |

| | | |
|---|---|---|
| | TTGTTTTCGCGCAAGAAAACCTTACCCTGCGTATGGTA<br>AGTTCGTTTAATATTAATAGTAATATTTCTTATTAACACT<br>TCCTTTTCAAAGTTACCTCTAACCTTCCAACTTGAAGGT<br>GCCGCGGTGTTGGCAACGTTTGTTCATCCCAATCGCAT<br>AGTGTATCTATGCTCGTGGGGATGAATTCTCTTGCTGC<br>CTACCTGCAACGCCAAGTAGTTTGGGTATAGTTGCTTG<br>TGGTATCACCGACTCATAAAAAAGCCGCGTAATGCGGT<br>TTTTTTATGGTCATTCCTAAACCACTCTGCTTTTAGCAG<br>CGATGCTTGATGCCCAGTGCTTCATCTAAGCTGGCGGT<br>GATGACGCAGACGCTATGCCGTAAACAGGGCATGTTTT<br>GCGAGGGCATCTCTTAACTGGGTCAGTAATGCCGCTTG<br>CTCTTGCTGAGTACGAAACACGCCTTGTGTTCTGCCCC<br>AAACCGGGCCCGGCCATGCCATATCATGGGTAAAGCG<br>CGCAATATGATGGATGTGCAGTTGTGGCACCATATTGC<br>CTAGAGCACCTAAGTTGAGTTTGTCTGGGGAAAATAGA<br>GTTTCCAGAGTTTGGCAAACCAGCTGCGATTCGACTAA<br>AAACTGCTGCTGCTCATCCATTGGCAGATGGTGTAGTT<br>CACGTAAATCCGCTCTTTGCGGGACTAAAATAACCCGG<br>GGCACGGCAGA | |
| | ATTTTCCTCAATACGCATGCGGCTGCGGCGGCAGGGG<br>CTATCGCTGCGCTTTTAGTGTGTAAAACCACTTGGGGT<br>AAAGCGGATCTCACCATGGTACTGAACGGTGCATTAGC<br>GGGCTTAGTAGCGATTACGGCCGATCCTCTCTCTCCTT<br>CACCGCTTTATTCGGTAGCGATTGGCGCAGTATCTGGC<br>TCTCTGGTGGTGTTTAGCATTATTGGTCTCGATAAGCT<br>GAAAATTGATGATCCAGTGGGTGCGATTTCCGTCCATG<br>GCGTGTGTGGTTTCTTTGGTCTGATGGTGGTACCACTG<br>AGCAACGCTGATGCAACCTTTGGCGCACAATTACTCGG<br>TGCTGCGGTGATCTTCGCATGGGTGTTTGGTGCAAGTC<br>TGGTGGTTTGGGGCGTGCTTAAAGCGACGATGGGGAT<br>CCGCGTCTCTGAAGAAGAAGAGATGGAAGGGATGGAC<br>ATGCACGATTGTGGTGTCGGTGCTTACCCAGAGTTTGT<br>TACTGTTAAGTGAAATATCATTTTACAAATTGAACCGAA<br>CAAAACCTGCCGTAAAGGCGGGTTTTGTTTTTTTGCCA<br>ATTGTTTTCTTATTGAACATTCCTATAATACATGCTCTCA<br>CCAAGCGTCTTGGATTGCCAAACTATGTTACCTGCATA<br>AAAAAGCCCGACGTCGAGCGTCGGGCAAGTGTTCTTC<br>TCCGCTGAGGGGACTTAACCTCAGGGGATGATTTGACA<br>AGGCGATGATTACAACGCCAAGAAAGCTTTCATCTGTT<br>GAAGAACACGCTCCGCAGTATCTTTGTCGGCCATTTCG<br>GCGAAAATACGTAATAAAGGTTCAGTACCGGAGAATCG<br>GGCAATTACCCAGCCACCGTTCTTGAAGTAGACTTTCG<br>CCCCATCTTCATAGCTGACTTTATCGATGTCGAATTCAA<br>ACTCAGGCAGTTTTTTCTCAATGTAGACCTTGTTGTAGA<br>TCTCATCACGCTGAGCGGGCTTAAATTTGCAATCGCCT<br>TCCGCAGTGTAGGCATAGCCGTAACGATCATAGATCTC<br>GGTTAACAGCTCAGAGAGTTTCTTGCCCGTCACGCTGA<br>TCATTTCCACCAATAGGCTAGAAGCGAACACGCCATCT<br>TTCCCCTTGATGTGACCACGAATCGTGAGGCCGCCAG<br>AGCTCTCCCCGCCAATCAGAGAGTCGTCCGCTTCCATT<br>TGTGAACCCGGGTGTTTAAAGC | *fbpABC* deletion |
| | TAAATAAAACGCATGCTAGTAATAATAGAGGCTTTTGGC<br>TAGCACGAGAAGAAATGTGCTGTAGGAATGACCAAAGA<br>TTGACATAGTAAGCTTTACTCTGACCTTGTATCCTCTTC<br>CTACTTGAAGCAGTAGCGGTGTTGGCTACGTTCGCCCC<br>CCAATCACAGTGTTTATCTATATTCATGGAGATGAACTC<br>TCTTGTCGCCTACCTGCAACTCCAAGTCGTTTGGGTAT<br>AGTGACGCTATTTACGTTTTCGCTTCTCTCTACTTCCCC<br>TGCAACGCTCTTTCTATCTAAGCAATGTGCTCATAAATG<br>CAAATGAGAATGCTTTACATTTGATTTGTGAATTATTAA<br>GATTCTCAATGATGTACGTTCTATGCAACTCAGTGTCTT<br>GGTTAAACCGTGGCTCAGTGCATTGTTACTGTATGTTC<br>GTCGGAATACTTGCAATCAAGCGCATGGACTTGAGTCA<br>TGGTTTAGCAGTGCTGTTTTGTTTTGCCGCGTACTTTTT<br>CGTTCTGGTATTTGTCATTGAAAACGAAGGAAGCCTCT<br>CATCTCCATACTGGCCTGAGCGCAGTGTAGTGGTTAAT<br>TTAGAGTTAAGGAGAAATTCAAATGATATTTCAAATGGA<br>CTGAAACAAAACAACGCCTCTAAGTCAGCTGAGAGGCG<br>TTTTTTTATGTTCGCATTTAAGTCTTGAGCTGAGCTCTC<br>GAGGTTAAACCATATCCTAAGAATCTGGGCTCCCTTGG | VC2211 deletion |

| | | |
|---|---|---|
| | TTAAGTTGATCGACAGTTGTGTAAAATTGGTTAATGATA<br>TACATTCTCATTTGAATTGATTGTTTACAGGGCCGAACC<br>ATTGTTTTTCAACTGGGGCAGAACGTTGGGAGGATTGA<br>GTGAATGACACTCCCAGCAAGGTTGGCTAAGGCATAAG<br>ATAAATTAGGTGAAAGAATGAGTAACGAAGTCGAGCGC<br>GTTTATCCGAGATTACTGGATTTTGTGCGTAAAAAATAT<br>GTATCAAAAAACCTACTTCGAGTGACACTGACTGGTGA<br>GGATTTAATTGGATTCCCCGAAGATCAAAATGGGTCAC<br>ACATTAAGGTTTTTTTCCCTAATCAGGCGAGTGGCATAC<br>TCCAATTACCGATTCGTGAGGGTGATAAAGTCATTTGG<br>CCAGAGCATAAACCGGTTCCGCGAGCGTATACCGTTA<br>GACAATACCGAGCACAGAGCAATGAGCTTGACCCGGG<br>TTTTGTTGTG | |
| | GTTGTGTTTTGCATGCAATCACGACCAACACGGGCATT<br>TCTTGCTCTTGATTACTCGTATTCTGCGTTTCCGCATAG<br>GCAAAACTGCCCGTGAAAAGCCCCATCATCATCGCCGA<br>CAAGGTGTGCAATTTAAACTTTTTGTTCTCTGCAACACT<br>GACTCGCGCTGGGCATAAAACTGCCATTTGAATTTCTC<br>CTTAACTCTAAATTAACCACTACACTGCGCTCAGGCCA<br>GTATGGAGATGAGAGGCTTCCTTCGTTTTCAATGACAA<br>ATACCAGAACGAAAAAGTACGCGGCAAAACAAAACAGC<br>ACTGCTAAACCATGACTCAAGTCCATGCGCTTGATTGC<br>AAGTATTCCGACGAACATACAGTAACAATGCACTGAGC<br>CACGGTTTAACCAAGACACTGAGTTGCATAGAACGTAC<br>ATCATTGAGAATCTTAATAATTCACAAATCAAATGTAAA<br>GCATTCTCATTTGCATTTATGAGCACATTGCTTAGATAG<br>AAAGAGCGTTGCAGGGGAAGTAGAGAGAAGCGAAAAC<br>GTAAATAGCGTCACTATACCCAAACGACTTGGAGTTGC<br>AGGTAGGCGACAAGAGAGTTCATCTCCAACCTTTTTCA<br>CTCGATTTTTGTTATCACCAGAAAAAACAAAGCCCGAAT<br>GGGCCTTTGTAGATTGAATCTTGATGATTACCGTAAATT<br>ATTCAGTAACTTGGTACTGGAATTCAGGGATCACTAGC<br>TCAACACGACGGTTCTTCTCACGACCTTCTGCTGTCGC<br>GTTAGAAGCGATTGGGCTGCTTTCGCCTAGACCTTTAG<br>CTGAAATACGAGATGCATCAATACCTTGAGCTTCTAGT<br>GCTTTCGCTACTGCTTGAGCACGACGCTCAGACAGTTT<br>TTGGTTGTAAGCTTCAGAACCTGTTGAGTCTGTGTGAC<br>CAACTACTTCTACTTTTGCTTGTGGGTATTGGTTTAGGT<br>AACCCACAATTTTATCCAGCTTCTGAACCGTTGCTGGTT<br>TTAGCTCAGCACTTGCAGTAGCAAAGGTGCTGCTATCC<br>AGATGTTGGAAAGTAAAGGTTTTAGTGACTGCCACTTT<br>CTCGACTGGTGCGGCAACAGGTGCAGGCTCAGCTGGA<br>CGCTGTTCTACCACTGGTGCTGGCTCTTCACTGCCACC<br>GAACTTATAAGCGATACCGAGTGTAGCGGTTTCTGCGC<br>CCCGGGGTACGATGTC | VC2212 deletion |
| | TGCGGAAATCGCATGCTTGAACGAAGAAGTCAAAAATC<br>TTCAGGTGTACCGTAATCACTTACAATCTTTGGTGGCC<br>AATCAACAGCAAGAGATGGCCAGTCTTGAACAGCAAAC<br>TGAAGAGATCAAACGCACGCGCCAAGGCATAGTTCCTT<br>TGATGTATGACATGATTGAAGGACTGGAGGAGTGGGTT<br>GCGCAAGATAAACCGATTCGTCTTGCGGCACGTCAAGA<br>ACGCATTGAGAAACTCAAAGAACTGATGCCTCGGGCAG<br>ATGTCAGTGATGCTGAAAAGTACCGTCGTATTCTCGAG<br>GCATATCAGATTGAACTCGACTACGGCAATAAGCTTGG<br>CACCTATCAAGCCAAAATCACTCTCCCTTCTGCACAAG<br>AAGTGGAAGCGGATGTGCTCTATTTAGGTCGTCTGTCG<br>CTATTAGCACGCAGTTTGGATGGCGAGCAGTTCTGGAC<br>TTGGAATTCCAAGCAGAACGCATGGCAAGCAATCACGG<br>ATGCCAACAAAAGCGACCTCGCTGCCGCTTACCAATTG<br>GCTCAGCAACAGATTGCGCCGACACTGTTGAACTTGCC<br>TGTTTCGTTGACCGCTGCGGAGGCTAAATAAACGACGA<br>ATCTTAAGTTATCTCTTGCTCTCTTGCCTATCCGTGACG<br>ACTTACGCCGCGGAGCTGACACCGTACACGGCCAGTA<br>AAGTGGTGAAAGCGAATCAGCTAGCACAAGAGAATAAG<br>GTGAAAGAAGCGATTCAGATCCTAAAACAAGCAGAACT<br>CAGCCGCAGTTATGATCGCGCTTATGTGGCGCGTATG<br>CTTGGGGTGTTTTATTGGCAGAATGAGCAAATCCCTGC<br>GGCGATAGCATCCTTGAAGTTGGCGGTAGAGAGCAAC<br>GAGTTACAAGATGATCAAGCGTGGACAACACGCAAAAT<br>GTTGGCGGATCTTTTGATAAGCCAACAAGAGTATCGTT | *exbB2tonB2* deletion |

| | | | |
|---|---|---|---|
| | | CAGCACTCAAACATTACTACACGCTAGCGCAAGCCATT<br>CCCGCCGATCAAAAAGGGAATGAACTCTGGCTGCGCA<br>TTAGTCAGTTGCACTTCCAAATGCAAGAGTGGAAGGCG<br>GTACTGAATGGGATGCAACAGTATGCTCAGTTCCAAAC<br>CAAGGATGCGGTATTGCCACTTTCACTACAATTAGGCG<br>CACAGCTTAATTTGGAACAGTGGAAGCCGGGAATTCTC<br>ACCCTAAAACCCGGGTTGGTGTTGG | |
| | | TGAATGCGCAGCATGCTTTTACTGATGTTACCCAGTGC<br>TTCAACCATAGGAAAAAGATTCGCTGCGGGCACAATGC<br>CATCCAGTGCAATCGTGATATCGAGCTCCCAGCCATTA<br>GCCAGTAACGTAGCATCATTGACCAATTTTTCGGTCGC<br>AGCCAAGATGGCCCTGCCCCGCTCTAAGATAAGTTTGC<br>CCGCTTCAGTAAAATTGGCTCGATGACCGGAGCGATCA<br>AAGATCATCAAATCCAAATCCTGTTCAAGCTTTTGGATT<br>TGATAGCTCAATGAACTCGGTGCTCGATTAAGCTCATT<br>AGCTGCGGCTGCAAAGCTGCCTCTACGCTCAATTGCAT<br>CCAAAATATGCAGCGCTTCAAGAGTGATTGGACTTAAC<br>AATGCTCTCTCCTATTGATAGTTCACACCGCCAACTGT<br>GATGAGAGTCACAATTAGCAACATCATAAATGCCATTAT<br>GAATAATATAACCCTTTGATTATAAAATTCATTTAGAAAA<br>CTCAAGTCCTGTTCTTTTCACATTAAGCAAACAATTAAA<br>CACAAATGATAGCAATTATCATTAATATTTATTAGAATTC<br>CTGCGTTTTACTCAACATGGAAAATCCTTCTAAAAAGCA<br>CCATTTTACCGTGGTTTAGGAACGATAACAAAAAGCCA<br>GCTTACGCTGGCTTTTTTCTTAAAGAACACTTCATTCAA<br>AAGGAATTTAAAAGGTGATGAAATGAATTATTTGTTCTC<br>TGCCATCTCTTTTTTCACCATTACTGCAGCAGCAACGAT<br>AAAAGCAATGATCAATGCCAGTTCCACTTATCCTCCTAA<br>GCGATTCTACTGAGTTCGCACATAGTCTAGCACTGGGC<br>TTACCGACAAGCAGTGTTTATTTTGCGAAACTGGCTAAT<br>TAGCGATCGAGATCAAATCCCACGAGTGAGTAAAAGCT<br>AGTGTTCTGACAAGTCTGGCTTTACCCAGACTTGGCTA<br>AAATCGAACCAACCTAGCGCATTGCATTTCGCATTTTGT<br>AACGCACCGCACTGATCTTTACTGATGCCTAACCAGCA<br>GTGGAACATCGGGATCAGTTGGTGTTTTTCCACCAGTG<br>ATTTCCCCAGTTCTTTCGCAGGGAAGAGCGCAGAAGAG<br>TCAGCACGCCAAGCATCAATCAAACTCACCCACTGATT<br>GAAATCCTCTCCTTTGCTCAAAAACTCAATCCCGGGGT<br>AATTGAGT | *hutA* deletion |
| S1 mapping | | | |
| | TD-JHS065 | CCATGGCGGCCGCGGGAATTCGCGATAGCCAAAGCTC<br>TATAAAATCG | VxrA S1T22F280 |
| | TD-JHS066 | CAAGTCGATGCGCTCAGGAAGCGAATCG | VxrA S1R280 |
| | TD-JHS067 | CCATGGCGGCCGCGGGAATTCGCCTGTCACTCTCGAA<br>GTGATAACAC | VxrB S1T22F309 |
| | TD-JHS068 | CTAAACTGACGAGTAGACCGTCAGCGAG | VxrB S1R309 |
| | TD-JHS069 | CCATGGCGGCCGCGGGAATTCGCACCACATTTCGGCT<br>TGCACTGGC | MgtE S1T22F291 |
| | TD-JHS070 | CGCTCACGCGAAAGCCACTTGGGTTAAC | MgtE S1R291 |
| | TD-JHS071 | CCATGGCGGCCGCGGGAATTCGGCTCTTTAATGTCAAA<br>TGATTTGTGC | MurJ S1T22F344 |
| | TD-JHS072 | TACGTCAGCACTGGCACCTGCGCCCATC | MurJ S1R344 |
| | TD-JHS073 | CCATGGCGGCCGCGGGAATTCGGACACAAGCTTTGCT<br>GAGCTATGTACG | Pbp1B S1T22F384 |
| | TD-JHS074 | CCGAGAGCGTAGGCTCCATCCCTAACTG | Pbp1B S1R384 |
| | TD-JHS075 | CCATGGCGGCCGCGGGAATTCGGTGCCTCTTCCTCCA<br>AGCTGCAAATC | Pbp1B S2T22F145 |
| | TD-JHS076 | GTTTGATCATCGAATCGAGATAGATACC | Pbp1B S2R145 |
| | TD-JHS077 | CCATGGCGGCCGCGGGAATTCGGGTGGAAGGCAGTTT<br>GAATCGCTTTGTG | VC0213 S1T22F280 |
| | TD-JHS078 | CGATTTTTAGCAGCACAGGATAAGGCAG | VC0213 R280 |

| | | | |
|---|---|---|---|
| | TD-JHS079 | CCATGGCGGCCGCGGGAATTCGCGATTATCAGTACGT GGATCTGCCATG | VC0370 S1T22F349 |
| | TD-JHS080 | CTGATAGCCAAACTCGACTTGATGTTTC | VC0370 R349 |
| | TD-JHS081 | CCATGGCGGCCGCGGGAATTCGGAACAAGTAAAATCC GCAGGCTATGAC | VC0947 S1T22F282 |
| | TD-JHS082 | GGTGCGTCTGGTGTCACGATAGGAGAG | VC0947 R282 |
| | TD-JHS083 | CCATGGCGGCCGCGGGAATTCGGTATGATGGCTGGCA GTATCGTATTGAG | VC0948 S1T22F318 |
| | TD-JHS084 | GGTGCTTGATCGTCAGACATATCATAGC | VC0948 R318 |
| | TD-JHS085 | CCATGGCGGCCGCGGGAATTCGGTATCCCGTTTGATG TTCTGAGCTCTAG | VC0969 S1T22F343 |
| | TD-JHS086 | GTTAGCCAGTAGCGGAAGAACAACGAAC | VC0969 R343 |
| | TD-JHS087 | CCATGGCGGCCGCGGGAATTCGCAGCAGATATTTTTC CTTTCTGCTCAAG | VC1269 S1T22F289 |
| | TD-JHS088 | GTTGCTTAATGCCAATTCTCTTGGCTTG | VC1269 R289 |
| | TD-JHS089 | CCATGGCGGCCGCGGGAATTCGCGGTGTTGGTTACTA TCCGAAAAGCCAA | VC1270 S1T22F280 |
| | TD-JHS090 | GAGCTGTTTCACATCACCACCCGGATCC | VC1270 R280 |
| | TD-JHS091 | CCATGGCGGCCGCGGGAATTCGCTGAAACAACTCAAT GAATTTGCTCGC | VC1655 S1T22F118 |
| | TD-JHS092 | GAATTTCTTGTTCGGTGAGTTTGAACAG | VC1655 R118 |
| | TD-JHS093 | CCATGGCGGCCGCGGGAATTCGCAGATGTTTGGTAAA CCTATTTTTGTC | VC1956 S1T22F333 |
| | TD-JHS094 | CTTGCTGCTTTAGCCTTTCGACATATTG | VC1956 R333 |
| | TD-JHS095 | CCATGGCGGCCGCGGGAATTCGGAGCACATTGCTTAG ATAGAAAGAGCG | VC2212 S1T22F358 |
| | TD-JHS096 | CGTCTCTAGGTTACTTTTCGACATGACG | VC2212 R358 |
| | TD-JHS097 | CCATGGCGGCCGCGGGAATTCGGCAAAGACGTGACAA AATAATAACCG | VC2213 S1T22F338 |
| | TD-JHS098 | CCTGATCATCATCTTCACAAGATTGACC | VC2213 R338 |
| | TD-JHS099 | CCATGGCGGCCGCGGGAATTCGGCTAACTATCAGCCG GTTTATCAATTG | VC2312 S1T22F380 |
| | TD-JHS100 | GTTCATACAGCGGTCGATAGATCTTGG | VC2312 R380 |
| | TD-JHS101 | CCATGGCGGCCGCGGGAATTCGGTTCTAACCGCATTG ATGTTTGGGAAC | VC2518 S1T22F278 |
| | TD-JHS102 | GTTCAGCGCGATCGACTCAACGCGTCC | VC2518 R278 |
| | TD-JHS103 | CCATGGCGGCCGCGGGAATTCGGGTTTGAGCACCGCA GCTTTAATGCTG | VC2635 S1T22F405 |
| | TD-JHS104 | GATCAGTTTGCCGTCTTGACTGAAGAC | VC2635 R405 |
| | TD-JHS105 | CCATGGCGGCCGCGGGAATTCGCTAGCAGCCAGAGTC ACATTATCCAATTG | VCA0140 S1T22F298 |
| | TD-JHS106 | CTGTGGAGCATTTTCTCCTTGCATGTG | VCA0140 R298 |
| | TD-JHS107 | CCATGGCGGCCGCGGGAATTCGCAACTAAAACCTGCA CAAGTCATTAAAG | VCA0235 S1T22F179 |
| | TD-JHS108 | GACAGCGGTTTTCTTAACTGCTAGAG | VCA0235 R179 |
| | TD-JHS109 | CCATGGCGGCCGCGGGAATTCGCACCCTGTTGCACCA TATACATTGACG | VCA0594 S1T22F310 |
| | TD-JHS110 | GTATTCCTTAACGGTGTACTCATTCTC | VCA0594 R310 |
| | TD-JHS111 | CCATGGCGGCCGCGGGAATTCGGATATCATGATGCAG CTCTTGGAGCTTG | VC0049 S1T22F260 |
| | TD-JHS112 | CTCTTCCAACCAGTGGAGGGCCTTGTAG | VC0049 R260 |
| | TD-JHS113 | CCATGGCGGCCGCGGGAATTCGCGACTGGGAAGCCT GTCATTTCAATCG | VC0271 S1T22F286 |

| | | | |
|---|---|---|---|
| | TD-JHS114 | GATAGCTTGGGGTAATACTTAAGAGTAC | VC0271 R286 |
| | TD-JHS115 | CCATGGCGGCCGCGGGAATTCGCGATTTGAATGGCGC GTATTCTAACC | VC0277 S1T22F283 |
| | TD-JHS116 | GGTGCGCGTTGCCGGAGCAATCAGCTC | VC0277 R283 |
| | TD-JHS117 | CCATGGCGGCCGCGGGAATTCGGTCATCTAAGCGGGC TAAAGATCATTTG | VC1272 S1T22F284 |
| | TD-JHS118 | GATTGCGCAGGTTGCTGCACAGAGGTGC | VC1272 R284 |
| | TD-JHS119 | CCATGGCGGCCGCGGGAATTCGGAGCATTTCACCGCG AATAAGACGATGG | VC1548 S1T22F369 |
| | TD-JHS120 | CGATTTCCGCACGCAGTAACAAACTCTG | VC1548 R369 |
| | TD-JHS121 | CCATGGCGGCCGCGGGAATTCGCAGCAAATGCTACAA ATTAACCACTTG | VC2037 S1T22F328 |
| | TD-JHS122 | GAATACCTAACGTCATGATCACGAACCAG | VC2037 R328 |
| | TD-JHS123 | CCATGGCGGCCGCGGGAATTCGCAGCTCGGATCGGG CTTAGCGGCTCG | VC2686 S1T22F329 |
| | TD-JHS124 | CTCTTGCGCTTCATTTTGCAGCTGTTGC | VC2686 R329 |
| | TD-JHS125 | CCATGGCGGCCGCGGGAATTCGCCATCAAATTGTATTA AAAGGTGATCC | VC2698 S1T22F313 |
| | TD-JHS126 | GAAGGGTATGAATACCGTAGTAAGCATC | VC2698 R313 |
| | TD-JHS127 | CCATGGCGGCCGCGGGAATTCGGTTCCAGACAAATTT GTCACTCACTTGC | VCA0227 S1T22F346 |
| | TD-JHS128 | GCTTGTGCAGCAAACGCAGCCAATAATC | VCA0227 R346 |
| | TD-JHS129 | CCATGGCGGCCGCGGGAATTCGGTGCAGCAGCCACAC ACATGCGCCTCCAG | VCA0657 S1T22F362 |
| | TD-JHS130 | GTTGCACTGGCAAAATCTTTGGCATCG | VCA0657 R362 |
| | TD-JHS131 | CCATGGCGGCCGCGGGAATTCGCCAACAATCTAACCA AAAGACAGATC | VCA0665 S1T22F321 |
| | TD-JHS132 | GCAAAATCCCTGCGGTGAGTAGGATAC | VCA0665 R321 |
| | TD-JHS133 | CCATGGCGGCCGCGGGAATTCGCAAACCATTGGAACA TGAACAAGGTGCC | VC0200 S1T22F265 |
| | TD-JHS134 | CACTCGGTGCGGCAACCAAAACAACAG | VC0200 R265 |
| | TD-JHS135 | CCATGGCGGCCGCGGGAATTCGGTAATCGGGATGAAT GCCTGCTTTCATC | VC0876 S1T22F347 |
| | TD-JHS136 | GACATGGCTAAATTGCTCTATGCCACG | VC0876 R347 |
| | TD-JHS137 | CCATGGCGGCCGCGGGAATTCGCAAGCTGATTTTCTG CTTTCGAGTGATG | VC1560 S1T22F345 |
| | TD-JHS138 | CAAGTGGATTGGTTTTGCTGTCATGCTG | VC1560 R345 |
| | TD-JHS139 | CCATGGCGGCCGCGGGAATTCGCTGTGAACCAATATC ACAAATATTGGC | VC1585 S1T22F403 |
| | TD-JHS140 | GCAGCAATACACTGCCATGCTCACCTG | VC1585 R403 |
| | TD-JHS141 | CCATGGCGGCCGCGGGAATTCGGTTGCTAAAATACTT GGCAAGTCAGTCG | VC2045 S1T22F221 |
| | TD-JHS142 | GGTGCTTACCGTGGTGGAAATCCAGAG | VC2045 R221 |
| | TD-JHS143 | CCATGGCGGCCGCGGGAATTCGGTAACTGGTAAGAGC ACTTTGTTCAGTG | VC2106 S1T22F351 |
| | TD-JHS144 | GTGGGAGGGTAACTTTAAGACCAGCATCC | VC2106 R351 |
| | TD-JHS145 | CCATGGCGGCCGCGGGAATTCGCAATCAAGCGCATGG ACTTGAGTCATGG | VC2211 S1T22F303 |
| | TD-JHS146 | CTGCGTTTCCGCATAGGCAAAACTGC | VC2211 R303 |
| | TD-JHS147 | CCATGGCGGCCGCGGGAATTCGGATGTAAACATTGTG TTCCTCATTTGTG | VC2636 S1T22F263 |
| | TD-JHS148 | GTGGGTTGGCTGACAAAGCACGCTTCTG | VC2636 R263 |

| | | | |
|---|---|---|---|
| | TD-JHS149 | CCATGGCGGCCGCGGGAATTCGCTTTACCTCTGGCGAGCGCTTAACGCAG | VC2694 S1T22F222 |
| | TD-JHS150 | CTAACGCATCGTAAGCGTAAGGTAGATC | VC2694 R222 |
| | TD-JHS151 | CCATGGCGGCCGCGGGAATTCGGGCATGGATAAATCCATACGCGATTAG | VCA0909 S1T22F334 |
| | TD-JHS152 | CACGTTCGCCTGCTCATGACTCGGTAC | VCA0909 R334 |
| | TD-JHS153 | CCATGGCGGCCGCGGGAATTCGGTATGGATTTATCCATGCCTCGTTAGTC | VCA0910 S1T22F227 |
| | TD-JHS154 | GTGCTTCATCTGTCGTGATAAGCAATAG | VCA0910 R227 |
| | TD-JHS155 | CCATGGCGGCCGCGGGAATTCGGTATGGATTTATCCATGCCTCGTTAGTC | VCA0084 S1T22F232 |
| | TD-JHS156 | CACCTGTGCTTCATCTGTCGTGATAAGC | VCA0084 R232 |
| | TD-JHS157 | CCATGGCGGCCGCGGGAATTCGGTTGTTATTCATGGTGGAGGTTTGTGC | VC1825 S1T22F295 |
| | TD-JHS158 | GGAAAATTCACCTGATAACTGAATTCAG | VC1825 R295 |
| | TD-JHS159 | CCATGGCGGCCGCGGGAATTCGCACAAACAACTAAGTTGCTGCCAGCAC | VCA0079 S1T22F264 |
| | TD-JHS160 | CACAGCCAAAAGCGGCAAACCTACTCG | VCA0079 R264 |
| | TD-JHS161 | CCATGGCGGCCGCGGGAATTCGGATACCGTTGAGTAACAGAGGCGTGTTC | VC0503 S1T22F232 |
| | TD-JHS162 | CTTGATCTGGCTCATCTGCGGTAGGTAG | VC0503 R232 |
| | TD-JHS163 | CCATGGCGGCCGCGGGAATTCGGCTTCAATACTCGCCATTCTTGTTCAC | VC0630 S1T22F316 |
| | TD-JHS164 | GATAGTGGCGTCCTACTTCTCGCTGAGC | VC0630 R316 |
| | TD-JHS165 | CCATGGCGGCCGCGGGAATTCGGTGAAGGTTCACCAGGACGTTAAGTCAC | RhyB S1T22F138 |
| | TD-JHS166 | CACTGGAAGCAATGTGAGCAATGTCGTG | RhyB R138 |
| | TD-JHS167 | GAGAAAATAGGTCTATCGAGCTGAAC | PBP1A(VC2635)R272 |
| | TD-JHS168 | GATAGTGCTTCAGCGCTTGCAACACG | PBP1B(VC0602)R289 |
| | TD-JHS169 | CGTCGCAGAATTATAATCAAAGAGCG | VxrB(VCA0565)R289 |
| | TD-JHS170 | CCATCAAATTGGCCACAACCACATC | MurJ(VC0680)R277 |
| | TD-JHS171 | GCCTAACGAGATGCTGTAATTACAC | YkoK(VC1655)R336 |
| | TD-JHS172 | GCCATGACTTCGCTTTAAGTAACGT | SIP(VC0370)R299 |
| | TD-JHS173 | CTTGTCAGGAAAAGGCTGGTTACAC | RlpA(VC0948)R262 |
| | TD-JHS174 | CTTGTCGGGGTACTTTACCAAAACTC | CysZ(VC0969)R271 |
| | TD-JHS175 | GCCATAGAAGTGTATTCCTTAACGG | Hemolysin(VCA0594)R274 |
| | TD-JHS176 | GATTGCATAAAAAGCCAAAGCCGCAG | OpmA(VC2213)R339 |
| | TD-JHS177 | GAGTAAAAAGTGGCACGGAAGTTGC | Flagellin(VC2188)R379 |
| | TD-JHS178 | CGCCTTTGCCTTTGATCAAAAACGC | ZapB(VC2686)R379 |
| | TD-JHS179 | GGAGTCAGGATATCCTTATTGAGTCG | NaHPump(VC2037)R325 |
| | TD-JHS180 | CCAAGAGCTGCATCATGATATCTTGC | SMG(VC0049)R266 |
| | TD-JHS181 | CAAGGTTAGCAACGATCGCTTTAATG | Hypo(VC1272)R275 |
| | TD-JHS233 | CCACCAGTCCATGTTGGACATGCTC | KatG(VC1560)R297 |
| | TD-JHS234 | CCGTTGTCTCGCGTAAGAGTTTGTGC | KatB(VC1585)R283 |
| | TD-JHS235 | GCTGAAGGCACGAGTGCTAGCATG | HutA(VCA0576)R310 |

| | | | |
|---|---|---|---|
| | TD-JHS236 | GCTTGTGGTCAGCTAATGCGACCAAG | prxA S1F305 |
| | TD-JHS237 | CACGATAACGGTTTTGCCTTTGAACAG | prxA S1R305 |
| | TD-JHS238 | GTAAGTACTGAACTATGTGCGTTGAG | OxyR2 S1F304 |
| | TD-JHS239 | GGTCGATTGACTGACAAAGCAACGCTC | OxyR2 S1R304 |
| | TD-JHS240 | CATCATGGTACCTAGGCACTACTGAC | AhpC S1F265 |
| | TD-JHS241 | GTTGAAGTTATCAACGATTTCACCG | AhpC S1R265 |
| | TD-JHS242 | CTGGTTCTTGAGAACAGCCATTAAG | dps S1F301 |
| | TD-JHS243 | CCACTGGAATATTCAAGGTAAGGAG | dps S1R301 |
| | TD-JHS244 | CGTACTAACTTACCAAGGACAACGC | dps S1F386 |
| | TD-JHS245 | CTCTTCAAACTTAGCGTGCAGTTCG | dps S1R386 |
| Footprinting | | | |
| | TD-JHS5'FAM1 | GCTATCGCGACTTTATGACTAAATAG | 6FAM-PBP1A(VC2635)F272 |
| | TD-JHS5'FAM2 | GCTTCGTCGTGAAAAGCTGCCAGAAC | 6FAM-PBP1B(VC0602)F289 |
| | TD-JHS5'FAM3 | GATAGCCAAAGCTCTATAAAATCGAC | 6FAM-WigK(VCA0565)F289 |
| | TD-JHS5'FAM4 | GTGCAAAAACCCAACTGGACCAAAC | 6FAM-MurJ(VC0680)F277 |
| | TD-JHS5'FAM5 | GGCAACGTTATTAATAGCTTTAGCG | 6FAM-YkoK(VC1655)F336 |
| | TD-JHS5'FAM6 | CAAACTCGACTTGATGTTTCCAAGG | 6FAM-SIP(VC0370)F299 |
| | TD-JHS5'FAM7 | GCTCACCATGCCTATGTTCACAAAC | 6FAM-RlpA(VC0948)F262 |
| | TD-JHS5'FAM8 | GACTAAGATGATGTTAGCCAGTAGC | 6FAM-CysZ(VC0969)F271 |
| | TD-JHS5'FAM9 | GTCTTAACGGCCAATCATGGGATAC | 6FAM-Hemolysin(VCA0594)F274 |
| | TD-JHS5'FAM10 | GAAGAAGCAAAAAGTAACGTCGCTG | 6FAM-OpmA(VC2213)F339 |
| | TD-JHS5'FAM11 | CGTTAATGGTCATAGTTTGCTCTCC | 6FAM-Flagellin(VC2188)F379 |
| | TD-JHS5'FAM12 | CTGCAGTTTGAATTTTCGCTTCCAG | 6FAM-ZapB(VC2686)F283 |
| | TD-JHS5'FAM13 | CTCAAGCCTTTCTTAGCAAATCTGAC | 6FAM-NaHPump(VC2037)F325 |
| | TD-JHS5'FAM14 | CAGCCAAGGTAGCAACCAATTGATTC | 6FAM-SMG(VC0049)F266 |
| | TD-JHS5'FAM15 | CTGATGATTGTACAGTTGCAAAGTGG | 6FAM-Hypo(VC1272)F275 |
| | TD-JHS5'FAM16 | CGTCAACACAGTATCTCAACTCATTCG | 6FAM-KatG(VC1560)F297 |
| | TD-JHS5'FAM17 | GACGGTTGATAAGTAAAAATATGAG | 6FAM-KatB(VC1585)F282 |
| | TD-JHS5'FAM18 | CAAGAGTGATTGGACTTAACAATGCTC | 6FAM-HutA(VCA0576)F310 |
| | TD-JHS167 | GAGAAAATAGGTCTATCGAGCTGAAC | PBP1A(VC2635)R272 |
| | TD-JHS168 | GATAGTGCTTCAGCGCTTGCAACACG | PBP1B(VC0602)R289 |
| | TD-JHS169 | CGTCGCAGAATTATAATCAAAGAGCG | VxrA(VCA0565)R289 |
| | TD-JHS170 | CCATCAAATTGGCCACAACCACATC | MurJ(VC0680)R277 |
| | TD-JHS171 | GCCTAACGAGATGCTGTAATTACAC | YkoK(VC1655)R336 |
| | TD-JHS172 | GCCATGACTTCGCTTTAAGTAACGT | SIP(VC0370)R299 |
| | TD-JHS173 | CTTGTCAGGAAAAGGCTGGTTACAC | RlpA(VC0948)R262 |
| | TD-JHS174 | CTTGTCGGGGTACTTTACCAAAACTC | CysZ(VC0969)R271 |

| | | | |
|---|---|---|---|
| | TD-JHS175 | GCCATAGAAGTGTATTCCTTAACGG | Hemolysin(VCA0594)R274 |
| | TD-JHS176 | GATTGCATAAAAAGCCAAAGCCGCAG | OpmA(VC2213)R339 |
| | TD-JHS177 | GAGTAAAAAGTGGCACGGAAGTTGC | Flagellin(VC2188)R379 |
| | TD-JHS178 | CGCCTTTGCCTTTGATCAAAAACGC | ZapB(VC2686)R379 |
| | TD-JHS179 | GGAGTCAGGATATCCTTATTGAGTCG | NaHPump(VC2037)R325 |
| | TD-JHS180 | CCAAGAGCTGCATCATGATATCTTGC | SMG(VC0049)R266 |
| | TD-JHS181 | CAAGGTTAGCAACGATCGCTTTAATG | Hypo(VC1272)R275 |
| | TD-JHS233 | CCACCAGTCCATGTTGGACATGCTC | KatG(VC1560)R297 |
| | TD-JHS234 | CCGTTGTCTCGCGTAAGAGTTTGTGC | KatB(VC1585)R283 |
| | TD-JHS235 | GCTGAAGGCACGAGTGCTAGCATG | HutA(VCA0576)R310 |
| Purification | | | |
| | TD-JHS001 | CCTTGGATTGACATATGTCGAATCAATGGTGGGACGAATG | For *Nde*I VxrB |
| | TD-JHS002 | CCATAAAAGAGGATCCGGCTTAACCATGATCACGCTTTC | Rev *Bam*HI VxrB |

**Supplementary Table S3** List of genes with VxrB binding peaks.

**Supplementary Table S4** List of genes differentially regulated by overexpression of VxrB$^{D78E}$

**Supplementary Table S5** List of genes that are regulated by overexpression of VxrB$^{D78E}$ (RNA-Seq) and exhibit VxrB binding (ChIP-Seq).

**Supplementary Table S6** List of genes that are regulated by overexpression of VxrB$^{D78E}$ (RNA-Seq) and exhibit VxrB binding (ChIP-Seq) summary, including validation results.

#### Supplementary Figure legends

**Supplementary Figure S1 | Specific binding of VxrB protein to target DNA fragments. (A)** To confirm VxrB's binding ability and specificity for these DNA fragments, electro mobility shift assay (EMSA) was carried out in two different binding conditions. Each labeled promoter probe of 50 bp (P*pbp1a* site I, P*pbp1a* site II, P*murJ*, P*vc1655*, P*vc0370*, P*vc0969*, P*vca0594* site I, P*vca0594* site II, P*ompA*, P*zapB*, P*vc2037*, and P*vc1272*) was incubated with increasing amounts of purified VxrB. **(B)** For VxrB specific binding activity test, a x50 or x300 molar excess of cold (non-labeled) self-probes or non-specific DNA mixtures were incubated with each labeled DNA fragment. After electrophoresis, the dried gels were exposed and visualized by a phosphor image analyzer (Typhoon-FLA 7000) and Multi Gauge V3.0 software.

**Supplementary Figure S2 | Overview of Vxr-regulated functions.** Functional classification of each positively or negatively controlled target genes was analyzed by KEGG mapper (https:www.genome.jp/kegg/mapper.html).

**Supplementary Figure S3 | VxrB core binding sites mapping by capillary electrophoresis based DNase I footprinting.** To validate the ChIP-Seq results via DNase I footprinting assay, 15 target DNA fragments (VC2635 PBP1A, VC0602 PBP1B, VCA0565 VxrB, VC0680 MurJ, VC1655 MgtE, VC0370 salt induced membrane protein, VC0948 lipo protein RlpA, VC0969 CysZ, VCA0594 hemolysin, VC2213 OpmA, VC2188 flagellin, VC2686 ZapB, VC2037 Na+/H+ pump, VC0049 SMG protein, VC1272 hypothetical) were chosen for the assay. The 5′ forward end was labeled with 6-FAM (Fluorescein). Each DNA probe was incubated with increasing amounts of VxrB (0.17, 0.34, 0.68, and 1.38 μM) and digested with DNAseI. Digested samples were analyzed by ABI 3730x1 DNA analyzer (Life Technologies). Mapping profiles of 10 target DNAs [VC1655 MgtE **(A)**, VC0370 salt induced membrane protein **(B)**, VC0969 CysZ **(C)**, VCA0594 hemolysin **(D)**, VC2213 OmpA **(E)**, VC2686 ZapB **(F)**, VC2037 Na+/H+ pump **(G)**, and VC1272 hypothetical **(H)**] are presented with no added VxrB was analyzed in parallel. P*murJ* **(A)** and P*pbp1a* **(B)** profiles are shown in **Figure 3**. The core protected regions were indicated in red as VxrB binding site; VBS and promoter elements (TSS; + 1 as bent arrow and -10 -35 as black rectangles) were shown on the DNA probe used for the DNase I foot printing assay.

**Supplementary Figure S4 | IPTG-dependence of a ∆*pbp1AB* mutant.** Wild type and the ∆*pbp1AB* mutant (∆*pbp1A* ∆*pbp1B* $P_{IPTG}$::*pbp1a*) were grown overnight, washed 2 x with fresh growth medium and then diluted 10,000 fold into fresh medium with the indicated IPTG concentrations. Data are presented starting from the 200 min time point, where the wild type cultures reached OD readings above background.

**Supplementary Figure S5 | Manganese uptake, hydrogen peroxide production and *ryhB* induction by PenG-treated cells (A)** Total supernatant hydrogen peroxide ($H_2O_2$) levels were measured using Amplex™ Red Hydrogen Peroxide/Peroxidase Assay Kit and normalized to total cellular protein level (measured by Qubit™). Data are average values from three independent experiments, error bars represent standard deviation. Cat, catalase (1µg/ml). **(B)** Expression levels of the Fur-regulated small RNA *ryhB* was quantified using S1 nuclease mapping. Cells were grown to mid-exponential phase followed by exposure to $H_2O_2$, PenG or the positive control 2 mM EDTA (a strong chelator of divalent cations) for the indicated duration. Numbers represent expression levels normalized to untreated control (averages from three independent replicates). **(C)** Time-course of manganese accumulation in PenG-treated WT and ∆*vxrAB* cells as measured by ICP-MS. Data are average of 3 independent biological replicates, error bars represent standard deviation. ∗∗ P=0.001; ∗∗∗P <0.0003 (paired t-test).

**Supplementary Figure S6 | Iron chelation reduces ∆*vxrAB* death in the presence of PenG.** Time-dependent killing experiment in the absence or presence of 250 µM bipyridyl (Bip) with or without addition of 250 µM $Fe^{2+}$. Data are average of 3 independent biological replicates, error bars represent standard deviation.

**Supplementary Figure S7 | Several combinations of ∆*vxrAB* with iron uptake system mutants restore ∆*vxrAB* tolerance (A-C)** Time-dependent killing experiment in the presence of PenG for the indicated time points. **(A)** Iron influx systems' single mutation. **(B)** Double disruption. **(C)** Triple deletion with

Δ*vxrAB*. (**D-F**) Combinations of Δ*vxrAB* with different combinations of iron uptake mutants effect on restoration of growth on iron-supplemented sucrose. Wild type and all combination mutant strains of iron uptake systems in Δ*vxrAB* were grown to exponential phase ($OD_{600}$~0.5) in LB medium and exposed to PenG **(A-C)** or spot-plated on 10% Sucrose plates with or without additional iron sulfate **(D-F)**.

**Supplementary Figure S8 | RNA induction profiles of known ROS defense systems after exposure to PenG.** mRNA expression profiles of ROS defense systems (*oxyR1*, *katG*, *prxA*, *dps* and *ahpC*), and *vxrAB* system were detected by S1 nuclease mapping assay. Wild type and *ΔvxrAB* mutant strains were grown until $O.D_{600}$~0.5 in 300 ml LB medium and total RNA samples isolated from each 40 ml cell culture after PenG **(A)** or $H_2O_2$ **(B)** shock for indicated time points. Representative images and average signal intensities are shown from three independent experiments.

**Supplementary Figure S9 | Gel based catalase and superoxide dismutase (SOD) activity staining and detection of protein oxidation triggered by $H_2O_2$. (A)** Wild type and *Δvxr*AB cells were grown to exponential phase ($OD_{600}$~0.5) in LB medium and exposed to PenG. Samples were collected at the indicated time points. 30 µg/well of crude proteins were separated by 5% (catalases) or 8% (SODs) native PAGE at 4 °C followed by staining (see Methods for details). To quantify the SOD or catalases activity, stained gels were scanned and analyzed by Odyssey Imaging Systems (LI-COR Biosciences) and Image Studio Lite software. Rpol $\alpha$ subunit (RNA polymerase I, $\alpha$ subunit) was visualized as internal control. Coomassie staining (loading control) was used to ensure consistent protein loading between samples. **(B)** Detection of protein carbonylation. For the positive control of protein damage by PenG shock, Wild type and *Δvxr*AB mutant were grown to logarithmic growth (log) and harvested at different times (0, 30, 60, and 180 min) after 0.5 mM $H_2O_2$ treatment. Both DNP-derivatized or nonderivatized (negative control) protein samples were separated by polyacrylamide gel electrophoresis followed by Western blotting with-DNP specific antibody (cat# D9656). The signals were scanned on an Odyssey CLx imaging device (LI-COR Biosciences) and visualized by image studio lite software.

**Supplementary Figure S10 | Growth rate and total cellular $H_2O_2$ level in electron transport chain mutants. (A)** Overnight cultures were diluted 100fold into fresh medium, growth was monitored via OD600 readings in a Bioscreen growth analyzer plate reader. **(B)** To measure total intracellular hydrogen peroxide ($H_2O_2$) levels, wild type and each single (*Δvxr*AB, *Δnqr*A or *Δubi*A) or double mutant (*Δvxr*AB*Δnqr*A and *Δvxr*AB*Δubi*A) strains were grown in medium containing 0.2% glucose to $O.D_{600}$ ~0.5. After that, PenG (100 μg/ml) was added. At indicated time points, 1ml cell culture was pelleted. $H_2O_2$ level was measured by Amplex$^{TM}$ Red Hydrogen Peroxide/Peroxidase Assay Kit (Invitrogen™ cat#A22188) and normalized with total cellular proteins. Average values from three independent experiments are presented with standard deviation (see Materials and Methods for detail).

#### Supplementary Methods

**Electrophoretic mobility shift assay (EMSA) for VxrB-DNA binding activity.** Each VxrB target promoter DNA probes of ~50 bp containing VxrB binding sites were isolated by using crush and soaking method from the polyacrylamide gel (Sambrook J and Russell DW, 2006) after annealing with each primer pairs in Table S2. The purified DNAs were labeled at 5′ ends with ($\gamma$-$^{32}P$) ATP using T4 polynucleotide kinase. Binding reactions were performed with approximately 1 fmol of labeled DNA fragments and 43 to 344 nM of purified VxrB protein in 20 μl of the reaction buffer [20 mM Tris-HCl (pH 6.4), 50 mM KCl, 1 mM DTT, 0.1 mg of bovine serum albumin/ml, 5% glycerol, 0.1 μg of poly(dI-dC), 0.1 mM ATP, and 0.1 mM $MgCl_2$]. For VxrB specific binding activity test, x50 or x300 molar excess of cold (non-labeled) self-probes or non-specifc DNA mixtures which is digested pGEM vector by *Hpa*II. Following incubation at room temperature for 20 min, the binding mixture was subjected to electrophoresis at 4°C on a 5% polyacrylamide gel in TA (pH6.4) buffer. After electrophoresis, the dried gels were exposed and visualized by a phosphor image analyzer (Typhoon-FLA 7000) and Multi Gauge V3.0 software.

**Western blot analysis to detect alpha subunit of RNA polymerase.** For the internal control of SOD and catalase activity, the same crude extracts were resolved on two independent 12 % polyacrylamide gels for each western blotting and coomassie staining. After electrophoresis the proteins were transferred to a membrane by using a semi-drying transfer system (I Blot 2-Invitrogen). The membrane was then blocked with blocking solution (dry skim milk dissolved in 20 mM Tris-HCl (pH7.8), 150 mM NaCl, 0.1% Triton X-100) overnight. The membrane was incubated with monoclonal anti-αRpoI antibody (BioLegend-cat#663104) for two hours, washed with TBST (20 mM Tris-HCl (pH7.8), 150 mM NaCl, 0.1% Triton X-100), and incubated with anti-mouse secondary antibody (IRDye 800CW, Li-Cor cat# 926-32211). All images were scanned on an Odyssey CLx imaging device (LI-COR Biosciences) and visualized using image studio lite software.

#### Supplementary Table S3. List of genes with VxrB binding peaks.

### This file is generated by MACS version 2.0.10.20120913 (tag:beta)
### ARGUMENTS LIST:
### name = peaks
### format = AUTO
### ChIP-seq file = ['CHIPWigRHis100PG3H.bam']
### control file = ['InpWigRHis100PG3H.bam']
### effective genome size = 2.70e+09
### band width = 300
### model fold = [5, 50]
### qvalue cutoff = 1.00e-10
### Larger dataset will be scaled towards smaller dataset.
### Range for calculating regional lambda is: 1000 bps and 10000 bps
### Broad region calling is off

### tag size is determined as 150 bps
### total tags in treatment: 6618387
### tags after filtering in treatment: 2812499
### maximum duplicate tags at the same position in treatment = 1
### Redundant rate in treatment: 0.58
### total tags in control: 9724075
### tags after filtering in control: 4734726
### maximum duplicate tags at the same position in control = 1
### Redundant rate in control: 0.51
# d = 200

| chr | start | end | length | abs_summit (summit position) | pileup (stacked bases) | -LOG10(pvalue) of summit | fold enrichment | -LOG10(qvalue) | name | ID | Rank |
|---|---|---|---|---|---|---|---|---|---|---|---|
| NC_002505 | 20807 | 21475 | 669 | 21301 | 266 | 14.49967 | 1.67129 | 12.77599 | peaks_peak_6 | Peak_chr1_1 | 345 |
| NC_002505 | 44654 | 45468 | 815 | 44885 | 339 | 34.27369 | 2.10523 | 32.06459 | peaks_peak_10 | Peak_chr1_2 | 156 |
| NC_002505 | 46815 | 47221 | 407 | 46981 | 323 | 25.01373 | 1.88941 | 23.01955 | peaks_peak_11 | Peak_chr1_3 | 219 |
| NC_002505 | 90658 | 91009 | 352 | 90836 | 342 | 36.61581 | 2.15971 | 34.35271 | peaks_peak_19 | Peak_chr1_4 | 144 |
| NC_002505 | 94532 | 94854 | 323 | 94699 | 290 | 20.73194 | 1.83064 | 18.84114 | peaks_peak_20 | Peak_chr1_5 | 256 |
| NC_002505 | 105834 | 106133 | 300 | 105986 | 263 | 14.81652 | 1.68766 | 13.08221 | peaks_peak_24 | Peak_chr1_6 | 335 |
| NC_002505 | 107057 | 107654 | 598 | 107259 | 285 | 18.64637 | 1.77478 | 16.80893 | peaks_peak_25 | Peak_chr1_7 | 279 |
| NC_002505 | 109874 | 110220 | 347 | 110052 | 280 | 17.74968 | 1.7554 | 15.93566 | peaks_peak_26 | Peak_chr1_8 | 289 |
| NC_002505 | 134439 | 135147 | 709 | 134779 | 248 | 12.11572 | 1.61507 | 10.47253 | peaks_peak_29 | Peak_chr1_9 | 402 |
| NC_002505 | 144823 | 145272 | 450 | 145036 | 342 | 35.85617 | 2.13954 | 33.6104 | peaks_peak_30 | Peak_chr1_10 | 148 |
| NC_002505 | 148899 | 149306 | 408 | 149063 | 332 | 34.08725 | 2.11845 | 31.88239 | peaks_peak_32 | Peak_chr1_11 | 159 |
| NC_002505 | 166827 | 167265 | 439 | 167038 | 280 | 19.36474 | 1.80786 | 17.50901 | peaks_peak_40 | Peak_chr1_12 | 274 |
| NC_002505 | 167995 | 168563 | 569 | 168415 | 301 | 24.73361 | 1.92882 | 22.74656 | peaks_peak_41 | Peak_chr1_13 | 222 |
| NC_002505 | 176732 | 177439 | 708 | 177245 | 240 | 12.52953 | 1.64481 | 10.87126 | peaks_peak_43 | Peak_chr1_14 | 390 |
| NC_002505 | 178100 | 178964 | 865 | 178744 | 248 | 14.59802 | 1.70799 | 12.87071 | peaks_peak_44 | Peak_chr1_15 | 340 |
| NC_002505 | 200455 | 200811 | 357 | 200646 | 334 | 30.24522 | 2.00923 | 28.13066 | peaks_peak_50 | Peak_chr1_16 | 184 |
| NC_002505 | 221440 | 221779 | 340 | 221598 | 340 | 31.56861 | 2.03075 | 29.42199 | peaks_peak_51 | Peak_chr1_17 | 176 |
| NC_002505 | 222028 | 222331 | 304 | 222199 | 325 | 31.36217 | 2.06161 | 29.22062 | peaks_peak_52 | Peak_chr1_18 | 178 |
| NC_002505 | 231691 | 232116 | 426 | 231929 | 259 | 12.94695 | 1.62784 | 11.27414 | peaks_peak_53 | Peak_chr1_19 | 375 |
| NC_002505 | 234082 | 235308 | 1227 | 234218 | 252 | 13.30765 | 1.65286 | 11.62344 | peaks_peak_54 | Peak_chr1_20 | 368 |
| NC_002505 | 254215 | 254887 | 673 | 254465 | 242 | 14.75966 | 1.7262 | 13.02708 | peaks_peak_61 | Peak_chr1_21 | 337 |
| NC_002505 | 257131 | 257694 | 564 | 257299 | 314 | 32.87756 | 2.1341 | 30.70063 | peaks_peak_62 | Peak_chr1_22 | 167 |
| NC_002505 | 278124 | 278605 | 482 | 278322 | 335 | 37.61663 | 2.20623 | 35.32866 | peaks_peak_69 | Peak_chr1_23 | 140 |
| NC_002505 | 282520 | 282820 | 301 | 282661 | 303 | 25.71098 | 1.9533 | 23.69982 | peaks_peak_70 | Peak_chr1_24 | 214 |
| NC_002505 | 299628 | 300102 | 475 | 299865 | 363 | 47.27649 | 2.37526 | 44.72988 | peaks_peak_72 | Peak_chr1_25 | 88 |
| NC_002505 | 329636 | 330077 | 442 | 329861 | 329 | 30.51612 | 2.02839 | 28.39545 | peaks_peak_79 | Peak_chr1_26 | 181 |
| NC_002505 | 330913 | 331812 | 900 | 331127 | 378 | 51.33161 | 2.42908 | 48.66179 | peaks_peak_80 | Peak_chr1_27 | 72 |
| NC_002505 | 360326 | 360666 | 341 | 360516 | 312 | 28.78175 | 2.02131 | 26.70035 | peaks_peak_85 | Peak_chr1_28 | 193 |
| NC_002505 | 375201 | 375689 | 489 | 375416 | 377 | 45.47874 | 2.28833 | 42.98847 | peaks_peak_87 | Peak_chr1_29 | 98 |
| NC_002505 | 386245 | 386718 | 474 | 386488 | 368 | 48.25588 | 2.38423 | 45.68023 | peaks_peak_91 | Peak_chr1_30 | 82 |
| NC_002505 | 391449 | 391892 | 444 | 391683 | 327 | 32.37654 | 2.08459 | 30.21124 | peaks_peak_92 | Peak_chr1_31 | 171 |
| NC_002505 | 421820 | 422526 | 707 | 422057 | 298 | 24.80823 | 1.93804 | 22.81907 | peaks_peak_98 | Peak_chr1_32 | 221 |
| NC_002505 | 444943 | 445442 | 500 | 445160 | 315 | 28.11467 | 1.99477 | 26.04833 | peaks_peak_103 | Peak_chr1_33 | 198 |
| NC_002505 | 461809 | 462249 | 441 | 462099 | 292 | 21.9732 | 1.86497 | 20.05072 | peaks_peak_105 | Peak_chr1_34 | 242 |
| NC_002505 | 495119 | 495405 | 287 | 495269 | 252 | 13.23831 | 1.6503 | 11.55661 | peaks_peak_106 | Peak_chr1_35 | 369 |
| NC_002505 | 510481 | 510957 | 477 | 510718 | 322 | 35.10947 | 2.17465 | 32.88123 | peaks_peak_108 | Peak_chr1_36 | 151 |
| NC_002505 | 513176 | 513778 | 603 | 513417 | 330 | 35.12903 | 2.15242 | 32.90027 | peaks_peak_109 | Peak_chr1_37 | 150 |
| NC_002505 | 532424 | 532873 | 450 | 532656 | 314 | 32.52176 | 2.12385 | 30.3532 | peaks_peak_115 | Peak_chr1_38 | 170 |
| NC_002505 | 535096 | 535478 | 383 | 535240 | 253 | 15.91523 | 1.74599 | 14.14931 | peaks_peak_116 | Peak_chr1_39 | 319 |
| NC_002505 | 539946 | 540528 | 583 | 540179 | 261 | 20.92893 | 1.90374 | 19.03273 | peaks_peak_117 | Peak_chr1_40 | 253 |
| NC_002505 | 544022 | 546240 | 2219 | 544242 | 277 | 31.57719 | 2.21148 | 29.43033 | peaks_peak_120 | Peak_chr1_41 | 175 |
| NC_002505 | 548496 | 549009 | 514 | 548676 | 213 | 11.97412 | 1.67545 | 10.33628 | peaks_peak_122 | Peak_chr1_42 | 410 |
| NC_002505 | 563077 | 564843 | 1767 | 564169 | 397 | 63.44891 | 2.65981 | 60.24549 | peaks_peak_125 | Peak_chr1_43 | 32 |
| NC_002505 | 580292 | 580694 | 403 | 580481 | 263 | 16.64312 | 1.75137 | 14.85751 | peaks_peak_127 | Peak_chr1_44 | 304 |
| NC_002505 | 594734 | 595026 | 293 | 594899 | 254 | 14.55521 | 1.69487 | 12.8293 | peaks_peak_130 | Peak_chr1_45 | 343 |
| NC_002505 | 597378 | 597707 | 330 | 597544 | 288 | 22.66558 | 1.89541 | 20.72677 | peaks_peak_131 | Peak_chr1_46 | 237 |
| NC_002505 | 601440 | 601755 | 316 | 601591 | 284 | 23.08668 | 1.91804 | 21.13806 | peaks_peak_132 | Peak_chr1_47 | 230 |
| NC_002505 | 604838 | 605114 | 277 | 604981 | 250 | 12.93047 | 1.64234 | 11.2583 | peaks_peak_134 | Peak_chr1_48 | 378 |
| NC_002505 | 611443 | 611969 | 527 | 611803 | 358 | 42.75315 | 2.27474 | 40.33649 | peaks_peak_136 | Peak_chr1_49 | 113 |
| NC_002505 | 622413 | 622744 | 332 | 622582 | 265 | 17.57383 | 1.77936 | 15.76401 | peaks_peak_139 | Peak_chr1_50 | 292 |
| NC_002505 | 623676 | 624061 | 386 | 623874 | 305 | 28.52184 | 2.0316 | 26.44569 | peaks_peak_140 | Peak_chr1_51 | 194 |

| | | | | | | | | | | | |
|---|---|---|---|---|---|---|---|---|---|---|---|
| NC_002505 | 628061 | 628402 | 342 | 628239 | 303 | 22.27448 | 1.851 | 20.34448 | peaks_peak_141 | Peak_chr1_52 | 239 |
| NC_002505 | 630404 | 630675 | 272 | 630558 | 259 | 12.32331 | 1.60516 | 10.67227 | peaks_peak_142 | Peak_chr1_53 | 394 |
| NC_002505 | 636584 | 636984 | 401 | 636772 | 334 | 40.10639 | 2.2773 | 37.7582 | peaks_peak_143 | Peak_chr1_54 | 122 |
| NC_002505 | 663446 | 664103 | 658 | 663590 | 257 | 16.04509 | 1.74246 | 14.27558 | peaks_peak_147 | Peak_chr1_55 | 316 |
| NC_002505 | 670926 | 673059 | 2134 | 671328 | 391 | 61.32818 | 2.63037 | 58.25627 | peaks_peak_148 | Peak_chr1_56 | 38 |
| NC_002505 | 684383 | 684609 | 227 | 684469 | 264 | 16.15863 | 1.7326 | 14.38593 | peaks_peak_149 | Peak_chr1_57 | 312 |
| NC_002505 | 694082 | 694533 | 452 | 694375 | 281 | 20.72178 | 1.8495 | 18.83117 | peaks_peak_150 | Peak_chr1_58 | 258 |
| NC_002505 | 695034 | 695758 | 725 | 695527 | 364 | 41.6158 | 2.22887 | 39.22874 | peaks_peak_151 | Peak_chr1_59 | 118 |
| NC_002505 | 707646 | 707941 | 296 | 707807 | 250 | 15.22877 | 1.7272 | 13.48215 | peaks_peak_152 | Peak_chr1_60 | 329 |
| NC_002505 | 713890 | 714295 | 406 | 714034 | 241 | 13.14983 | 1.6669 | 11.47079 | peaks_peak_153 | Peak_chr1_61 | 371 |
| NC_002505 | 716634 | 717027 | 394 | 716786 | 267 | 19.57212 | 1.84298 | 17.71095 | peaks_peak_154 | Peak_chr1_62 | 272 |
| NC_002505 | 723568 | 724154 | 587 | 723755 | 366 | 53.47551 | 2.5244 | 50.73292 | peaks_peak_155 | Peak_chr1_63 | 66 |
| NC_002505 | 725780 | 726246 | 467 | 726005 | 352 | 47.12893 | 2.40764 | 44.58707 | peaks_peak_156 | Peak_chr1_64 | 90 |
| NC_002505 | 739071 | 740155 | 1085 | 739806 | 392 | 60.14623 | 2.59773 | 57.13753 | peaks_peak_159 | Peak_chr1_65 | 45 |
| NC_002505 | 753655 | 753990 | 336 | 753807 | 274 | 21.07038 | 1.8767 | 19.17101 | peaks_peak_160 | Peak_chr1_66 | 251 |
| NC_002505 | 755918 | 756924 | 1007 | 756599 | 258 | 16.85218 | 1.76895 | 15.06098 | peaks_peak_161 | Peak_chr1_67 | 298 |
| NC_002505 | 778869 | 779260 | 392 | 779065 | 326 | 37.61255 | 2.23302 | 35.32477 | peaks_peak_165 | Peak_chr1_68 | 141 |
| NC_002505 | 788318 | 789169 | 852 | 788569 | 265 | 19.87583 | 1.85791 | 18.00656 | peaks_peak_166 | Peak_chr1_69 | 268 |
| NC_002505 | 799421 | 800013 | 593 | 799747 | 375 | 57.04038 | 2.58271 | 54.15988 | peaks_peak_167 | Peak_chr1_70 | 52 |
| NC_002505 | 809556 | 810043 | 488 | 809824 | 372 | 56.08322 | 2.56945 | 53.23766 | peaks_peak_170 | Peak_chr1_71 | 54 |
| NC_002505 | 819478 | 820020 | 543 | 819598 | 255 | 13.98841 | 1.67246 | 12.28215 | peaks_peak_172 | Peak_chr1_72 | 352 |
| NC_002505 | 841988 | 842432 | 445 | 842242 | 308 | 32.75872 | 2.14809 | 30.58434 | peaks_peak_174 | Peak_chr1_73 | 168 |
| NC_002505 | 893481 | 894476 | 996 | 894065 | 276 | 30.27129 | 2.17212 | 28.1561 | peaks_peak_186 | Peak_chr1_74 | 183 |
| NC_002505 | 898815 | 902363 | 3549 | 900213 | 226 | 22.77344 | 2.07955 | 20.83233 | peaks_peak_190 | Peak_chr1_75 | 234 |
| NC_002505 | 911578 | 911925 | 348 | 911756 | 225 | 13.77408 | 1.72433 | 12.07463 | peaks_peak_194 | Peak_chr1_76 | 358 |
| NC_002505 | 922998 | 923287 | 290 | 923135 | 298 | 28.20002 | 2.04049 | 26.13175 | peaks_peak_197 | Peak_chr1_77 | 197 |
| NC_002505 | 925000 | 925301 | 302 | 925128 | 255 | 18.43201 | 1.83137 | 16.60048 | peaks_peak_198 | Peak_chr1_78 | 281 |
| NC_002505 | 928351 | 928654 | 304 | 928500 | 279 | 22.14977 | 1.90004 | 20.22357 | peaks_peak_199 | Peak_chr1_79 | 241 |
| NC_002505 | 940191 | 941659 | 1469 | 940776 | 392 | 68.56988 | 2.80668 | 65.03608 | peaks_peak_202 | Peak_chr1_80 | 16 |
| NC_002505 | 941956 | 942705 | 750 | 942350 | 374 | 57.53819 | 2.59918 | 54.63897 | peaks_peak_203 | Peak_chr1_81 | 49 |
| NC_002505 | 962070 | 962304 | 235 | 962196 | 262 | 13.47029 | 1.64174 | 11.78094 | peaks_peak_208 | Peak_chr1_82 | 365 |
| NC_002505 | 980401 | 980774 | 374 | 980582 | 341 | 43.00624 | 2.33393 | 40.58389 | peaks_peak_211 | Peak_chr1_83 | 111 |
| NC_002505 | 982618 | 983065 | 448 | 982814 | 266 | 21.6414 | 1.91528 | 19.72722 | peaks_peak_212 | Peak_chr1_84 | 243 |
| NC_002505 | 1000738 | 1000982 | 245 | 1000863 | 244 | 13.79156 | 1.68564 | 12.0915 | peaks_peak_213 | Peak_chr1_85 | 356 |
| NC_002505 | 1001676 | 1002056 | 381 | 1001862 | 322 | 35.09714 | 2.1743 | 32.86915 | peaks_peak_214 | Peak_chr1_86 | 152 |
| NC_002505 | 1007451 | 1007692 | 242 | 1007605 | 243 | 11.83999 | 1.61284 | 10.20728 | peaks_peak_216 | Peak_chr1_87 | 416 |
| NC_002505 | 1012163 | 1012597 | 435 | 1012381 | 321 | 38.7357 | 2.28065 | 36.41997 | peaks_peak_217 | Peak_chr1_88 | 131 |
| NC_002505 | 1013181 | 1013593 | 413 | 1013406 | 375 | 61.14527 | 2.688 | 58.08503 | peaks_peak_218 | Peak_chr1_89 | 40 |
| NC_002505 | 1031364 | 1032184 | 821 | 1031990 | 350 | 52.35248 | 2.55485 | 49.64951 | peaks_peak_219 | Peak_chr1_90 | 69 |
| NC_002505 | 1033848 | 1034264 | 417 | 1034054 | 327 | 38.50333 | 2.25485 | 36.19363 | peaks_peak_220 | Peak_chr1_91 | 133 |
| NC_002505 | 1039043 | 1039454 | 412 | 1039229 | 276 | 26.54198 | 2.05026 | 24.51179 | peaks_peak_223 | Peak_chr1_92 | 210 |
| NC_002505 | 1040761 | 1041819 | 1059 | 1041312 | 381 | 63.22756 | 2.71705 | 60.03635 | peaks_peak_224 | Peak_chr1_93 | 34 |
| NC_002505 | 1043254 | 1043698 | 445 | 1043465 | 257 | 20.55158 | 1.90094 | 18.6651 | peaks_peak_225 | Peak_chr1_94 | 261 |
| NC_002505 | 1047978 | 1048312 | 335 | 1048127 | 261 | 21.13986 | 1.911 | 19.2382 | peaks_peak_226 | Peak_chr1_95 | 250 |
| NC_002505 | 1061042 | 1061553 | 512 | 1061246 | 339 | 47.28175 | 2.45778 | 44.73499 | peaks_peak_227 | Peak_chr1_96 | 87 |
| NC_002505 | 1079426 | 1079721 | 296 | 1079572 | 252 | 17.13282 | 1.79194 | 15.33441 | peaks_peak_228 | Peak_chr1_97 | 294 |
| NC_002505 | 1127484 | 1127990 | 507 | 1127752 | 377 | 56.38986 | 2.55886 | 53.5333 | peaks_peak_233 | Peak_chr1_98 | 53 |
| NC_002505 | 1130019 | 1130926 | 908 | 1130436 | 381 | 65.26976 | 2.76947 | 61.969 | peaks_peak_234 | Peak_chr1_99 | 26 |
| NC_002505 | 1131282 | 1131669 | 388 | 1131456 | 330 | 43.9232 | 2.39684 | 41.47543 | peaks_peak_235 | Peak_chr1_100 | 109 |
| NC_002505 | 1136930 | 1137428 | 499 | 1137205 | 352 | 51.79832 | 2.53218 | 49.11334 | peaks_peak_239 | Peak_chr1_101 | 71 |
| NC_002505 | 1158175 | 1158702 | 528 | 1158508 | 232 | 14.50461 | 1.73788 | 12.78068 | peaks_peak_241 | Peak_chr1_102 | 344 |
| NC_002505 | 1160441 | 1161270 | 830 | 1160644 | 306 | 36.19617 | 2.25624 | 33.94256 | peaks_peak_242 | Peak_chr1_103 | 146 |
| NC_002505 | 1163290 | 1163733 | 444 | 1163514 | 359 | 52.09127 | 2.51384 | 49.39691 | peaks_peak_243 | Peak_chr1_104 | 70 |
| NC_002505 | 1168930 | 1169238 | 309 | 1169052 | 229 | 11.87188 | 1.63897 | 10.23802 | peaks_peak_244 | Peak_chr1_105 | 414 |
| NC_002505 | 1170128 | 1171137 | 1010 | 1170581 | 380 | 63.5905 | 2.73047 | 60.37962 | peaks_peak_245 | Peak_chr1_106 | 31 |
| NC_002505 | 1176898 | 1177146 | 249 | 1177003 | 247 | 15.45983 | 1.74187 | 13.7065 | peaks_peak_247 | Peak_chr1_107 | 325 |
| NC_002505 | 1190461 | 1190704 | 244 | 1190564 | 234 | 12.92274 | 1.67148 | 11.25088 | peaks_peak_249 | Peak_chr1_108 | 379 |
| NC_002505 | 1200348 | 1201617 | 1270 | 1200981 | 380 | 67.2246 | 2.8245 | 63.77274 | peaks_peak_251 | Peak_chr1_109 | 21 |
| NC_002505 | 1206486 | 1206803 | 318 | 1206631 | 268 | 22.8204 | 1.94973 | 20.87807 | peaks_peak_252 | Peak_chr1_110 | 233 |
| NC_002505 | 1221427 | 1221812 | 386 | 1221587 | 329 | 47.6357 | 2.50601 | 45.0792 | peaks_peak_253 | Peak_chr1_111 | 85 |
| NC_002505 | 1226195 | 1226531 | 337 | 1226360 | 304 | 36.5589 | 2.2739 | 34.29718 | peaks_peak_256 | Peak_chr1_112 | 145 |
| NC_002505 | 1228464 | 1229387 | 924 | 1229018 | 366 | 60.42415 | 2.70641 | 57.40019 | peaks_peak_257 | Peak_chr1_113 | 43 |
| NC_002505 | 1232282 | 1232549 | 268 | 1232405 | 250 | 18.10946 | 1.83172 | 16.28624 | peaks_peak_258 | Peak_chr1_114 | 286 |
| NC_002505 | 1234761 | 1235199 | 439 | 1234975 | 380 | 60.15955 | 2.64294 | 57.1497 | peaks_peak_259 | Peak_chr1_115 | 44 |
| NC_002505 | 1237653 | 1238266 | 614 | 1237934 | 384 | 67.91038 | 2.82455 | 64.4163 | peaks_peak_260 | Peak_chr1_116 | 19 |
| NC_002505 | 1241832 | 1242131 | 300 | 1241951 | 224 | 11.87577 | 1.6487 | 10.24178 | peaks_peak_261 | Peak_chr1_117 | 413 |
| NC_002505 | 1247167 | 1247475 | 309 | 1247338 | 284 | 26.64048 | 2.03055 | 24.60815 | peaks_peak_262 | Peak_chr1_118 | 207 |
| NC_002505 | 1252942 | 1253314 | 373 | 1253149 | 301 | 29.43087 | 2.06935 | 27.33464 | peaks_peak_264 | Peak_chr1_119 | 188 |
| NC_002505 | 1265964 | 1266311 | 348 | 1266155 | 248 | 19.47701 | 1.88614 | 17.61835 | peaks_peak_267 | Peak_chr1_120 | 273 |
| NC_002505 | 1319097 | 1319419 | 323 | 1319271 | 323 | 39.46406 | 2.29482 | 37.1312 | peaks_peak_270 | Peak_chr1_121 | 126 |
| NC_002505 | 1344451 | 1344770 | 320 | 1344599 | 307 | 37.91693 | 2.30423 | 35.6218 | peaks_peak_272 | Peak_chr1_122 | 138 |
| NC_002505 | 1346546 | 1346920 | 375 | 1346762 | 298 | 34.84215 | 2.24228 | 32.61919 | peaks_peak_274 | Peak_chr1_123 | 154 |
| NC_002505 | 1353242 | 1353692 | 451 | 1353468 | 306 | 39.11659 | 2.34383 | 36.79183 | peaks_peak_275 | Peak_chr1_124 | 129 |
| NC_002505 | 1359521 | 1359832 | 312 | 1359665 | 237 | 15.92963 | 1.78165 | 14.1633 | peaks_peak_276 | Peak_chr1_125 | 317 |
| NC_002505 | 1371320 | 1371683 | 364 | 1371486 | 298 | 34.25888 | 2.22444 | 32.05008 | peaks_peak_279 | Peak_chr1_126 | 157 |
| NC_002505 | 1378973 | 1379438 | 466 | 1379264 | 278 | 28.20484 | 2.09827 | 26.13639 | peaks_peak_280 | Peak_chr1_127 | 196 |
| NC_002505 | 1381460 | 1381675 | 216 | 1381584 | 224 | 12.45057 | 1.67243 | 10.79516 | peaks_peak_281 | Peak_chr1_128 | 392 |
| NC_002505 | 1408733 | 1409052 | 320 | 1408889 | 255 | 22.90015 | 1.98873 | 20.95604 | peaks_peak_286 | Peak_chr1_129 | 232 |
| NC_002505 | 1410101 | 1410445 | 345 | 1410283 | 298 | 39.15545 | 2.37517 | 36.82971 | peaks_peak_287 | Peak_chr1_130 | 128 |
| NC_002505 | 1418508 | 1418763 | 256 | 1418643 | 232 | 16.46606 | 1.81441 | 14.68497 | peaks_peak_288 | Peak_chr1_131 | 307 |
| NC_002505 | 1430890 | 1431498 | 609 | 1431270 | 275 | 27.84762 | 2.09594 | 25.78715 | peaks_peak_290 | Peak_chr1_132 | 201 |

| | | | | | | | | | | | |
|---|---|---|---|---|---|---|---|---|---|---|---|
| NC_002505 | 1436288 | 1436877 | 590 | 1436548 | 348 | 55.25502 | 2.64243 | 52.44188 | peaks_peak_291 | Peak_chr1_133 | 60 |
| NC_002505 | 1438533 | 1439028 | 496 | 1438795 | 370 | 55.54283 | 2.56301 | 52.71869 | peaks_peak_292 | Peak_chr1_134 | 59 |
| NC_002505 | 1452230 | 1452741 | 512 | 1452526 | 290 | 33.28684 | 2.22127 | 31.09996 | peaks_peak_294 | Peak_chr1_135 | 164 |
| NC_002505 | 1456251 | 1458124 | 1874 | 1456815 | 388 | 74.12511 | 2.96744 | 70.05345 | peaks_peak_295 | Peak_chr1_136 | 7 |
| NC_002505 | 1470438 | 1470712 | 275 | 1470572 | 225 | 12.28747 | 1.6637 | 10.63777 | peaks_peak_297 | Peak_chr1_137 | 397 |
| NC_002505 | 1489948 | 1490786 | 839 | 1490522 | 285 | 33.10244 | 2.23271 | 30.91968 | peaks_peak_299 | Peak_chr1_138 | 165 |
| NC_002505 | 1493377 | 1493853 | 477 | 1493524 | 279 | 25.5172 | 2.00853 | 23.51076 | peaks_peak_300 | Peak_chr1_139 | 216 |
| NC_002505 | 1495201 | 1495767 | 567 | 1495437 | 376 | 72.76059 | 2.99055 | 68.84422 | peaks_peak_301 | Peak_chr1_140 | 9 |
| NC_002505 | 1496486 | 1497581 | 1096 | 1496692 | 323 | 44.10071 | 2.4276 | 41.64748 | peaks_peak_302 | Peak_chr1_141 | 108 |
| NC_002505 | 1499226 | 1499830 | 605 | 1499591 | 384 | 71.32064 | 2.91317 | 67.55944 | peaks_peak_303 | Peak_chr1_142 | 11 |
| NC_002505 | 1503483 | 1503777 | 295 | 1503624 | 231 | 15.09689 | 1.76339 | 13.35428 | peaks_peak_305 | Peak_chr1_143 | 331 |
| NC_002505 | 1506967 | 1507449 | 483 | 1507168 | 334 | 48.86776 | 2.52124 | 46.27251 | peaks_peak_307 | Peak_chr1_144 | 79 |
| NC_002505 | 1514270 | 1514662 | 393 | 1514480 | 347 | 49.77359 | 2.49631 | 47.15183 | peaks_peak_308 | Peak_chr1_145 | 76 |
| NC_002505 | 1520497 | 1521037 | 541 | 1520739 | 298 | 41.98685 | 2.46352 | 39.59053 | peaks_peak_310 | Peak_chr1_146 | 116 |
| NC_002505 | 1529918 | 1531283 | 1366 | 1530487 | 382 | 72.80769 | 2.96194 | 68.88506 | peaks_peak_312 | Peak_chr1_147 | 8 |
| NC_002505 | 1539519 | 1540471 | 953 | 1540146 | 350 | 59.66706 | 2.75625 | 56.68484 | peaks_peak_316 | Peak_chr1_148 | 47 |
| NC_002505 | 1541125 | 1541427 | 303 | 1541276 | 229 | 15.8136 | 1.79637 | 14.05062 | peaks_peak_317 | Peak_chr1_149 | 322 |
| NC_002505 | 1545158 | 1545733 | 576 | 1545545 | 309 | 38.08369 | 2.30221 | 35.78411 | peaks_peak_319 | Peak_chr1_150 | 136 |
| NC_002505 | 1559023 | 1559395 | 373 | 1559212 | 330 | 37.36329 | 2.21399 | 35.08251 | peaks_peak_321 | Peak_chr1_151 | 142 |
| NC_002505 | 1560639 | 1560964 | 326 | 1560798 | 266 | 21.6414 | 1.91528 | 19.72722 | peaks_peak_322 | Peak_chr1_152 | 243 |
| NC_002505 | 1588411 | 1588657 | 247 | 1588548 | 212 | 11.62994 | 1.66267 | 10.00567 | peaks_peak_335 | Peak_chr1_153 | 423 |
| NC_002505 | 1589110 | 1589595 | 486 | 1589352 | 323 | 44.25067 | 2.43193 | 41.79332 | peaks_peak_336 | Peak_chr1_154 | 107 |
| NC_002505 | 1602504 | 1602840 | 337 | 1602689 | 261 | 24.52824 | 2.02749 | 22.54644 | peaks_peak_339 | Peak_chr1_155 | 225 |
| NC_002505 | 1605819 | 1606174 | 356 | 1605989 | 249 | 21.35905 | 1.95131 | 19.45225 | peaks_peak_340 | Peak_chr1_156 | 248 |
| NC_002505 | 1608918 | 1609261 | 344 | 1609102 | 219 | 12.93398 | 1.70309 | 11.26161 | peaks_peak_341 | Peak_chr1_157 | 377 |
| NC_002505 | 1630659 | 1630920 | 262 | 1630790 | 253 | 20.56675 | 1.91197 | 18.67993 | peaks_peak_343 | Peak_chr1_158 | 260 |
| NC_002505 | 1649234 | 1649609 | 376 | 1649419 | 307 | 40.71469 | 2.38829 | 38.35043 | peaks_peak_344 | Peak_chr1_159 | 120 |
| NC_002505 | 1660617 | 1662098 | 1482 | 1661145 | 391 | 69.56259 | 2.83611 | 65.96381 | peaks_peak_347 | Peak_chr1_160 | 14 |
| NC_002505 | 1675395 | 1675715 | 321 | 1675546 | 263 | 23.72265 | 1.99402 | 21.76017 | peaks_peak_349 | Peak_chr1_161 | 227 |
| NC_002505 | 1680451 | 1680780 | 330 | 1680629 | 310 | 38.31422 | 2.30556 | 36.00933 | peaks_peak_350 | Peak_chr1_162 | 135 |
| NC_002505 | 1703465 | 1703730 | 266 | 1703612 | 232 | 16.08478 | 1.79959 | 14.31404 | peaks_peak_352 | Peak_chr1_163 | 315 |
| NC_002505 | 1710937 | 1711727 | 791 | 1711531 | 351 | 57.26861 | 2.68519 | 54.37827 | peaks_peak_353 | Peak_chr1_164 | 50 |
| NC_002505 | 1731810 | 1732947 | 1138 | 1732605 | 393 | 69.04184 | 2.81429 | 65.47992 | peaks_peak_357 | Peak_chr1_165 | 15 |
| NC_002505 | 1741242 | 1741781 | 540 | 1741499 | 327 | 45.9857 | 2.46657 | 43.48005 | peaks_peak_358 | Peak_chr1_166 | 96 |
| NC_002505 | 1751033 | 1751571 | 539 | 1751417 | 279 | 28.8311 | 2.11543 | 26.74864 | peaks_peak_361 | Peak_chr1_167 | 192 |
| NC_002505 | 1768082 | 1768578 | 497 | 1768273 | 252 | 20.87522 | 1.92562 | 18.98046 | peaks_peak_363 | Peak_chr1_168 | 254 |
| NC_002505 | 1770147 | 1770445 | 299 | 1770304 | 246 | 16.83594 | 1.79475 | 15.04515 | peaks_peak_364 | Peak_chr1_169 | 299 |
| NC_002505 | 1775883 | 1776219 | 337 | 1776017 | 243 | 19.18969 | 1.88889 | 17.33858 | peaks_peak_365 | Peak_chr1_170 | 275 |
| NC_002505 | 1777424 | 1778206 | 783 | 1777847 | 385 | 72.19291 | 2.93127 | 68.33851 | peaks_peak_366 | Peak_chr1_171 | 10 |
| NC_002505 | 1783846 | 1784106 | 261 | 1783981 | 240 | 12.6295 | 1.64868 | 10.9676 | peaks_peak_368 | Peak_chr1_172 | 386 |
| NC_002505 | 1786141 | 1786893 | 753 | 1786535 | 388 | 75.28632 | 2.99787 | 71.048 | peaks_peak_370 | Peak_chr1_173 | 4 |
| NC_002505 | 1793963 | 1794261 | 299 | 1794082 | 229 | 13.62516 | 1.7096 | 11.93066 | peaks_peak_371 | Peak_chr1_174 | 362 |
| NC_002505 | 1794468 | 1795203 | 736 | 1794757 | 374 | 66.29622 | 2.82708 | 62.91714 | peaks_peak_372 | Peak_chr1_175 | 23 |
| NC_002505 | 1801774 | 1802087 | 314 | 1801931 | 230 | 14.49445 | 1.74195 | 12.77098 | peaks_peak_374 | Peak_chr1_176 | 346 |
| NC_002505 | 1818115 | 1818376 | 262 | 1818232 | 237 | 14.9176 | 1.7429 | 13.18049 | peaks_peak_375 | Peak_chr1_177 | 333 |
| NC_002505 | 1848438 | 1848701 | 264 | 1848582 | 229 | 12.84147 | 1.67818 | 11.17226 | peaks_peak_377 | Peak_chr1_178 | 382 |
| NC_002505 | 1856281 | 1856625 | 345 | 1856448 | 291 | 32.11894 | 2.18143 | 29.9595 | peaks_peak_378 | Peak_chr1_179 | 173 |
| NC_002505 | 1860133 | 1861124 | 992 | 1860800 | 392 | 75.10676 | 2.9737 | 70.89553 | peaks_peak_379 | Peak_chr1_180 | 5 |
| NC_002505 | 1866617 | 1867533 | 917 | 1866772 | 249 | 19.62275 | 1.88878 | 17.76006 | peaks_peak_380 | Peak_chr1_181 | 271 |
| NC_002505 | 1871146 | 1871403 | 258 | 1871288 | 232 | 15.1064 | 1.76145 | 13.36355 | peaks_peak_381 | Peak_chr1_182 | 330 |
| NC_002505 | 1884634 | 1886001 | 1368 | 1885471 | 395 | 77.61683 | 3.02413 | 73.12509 | peaks_peak_385 | Peak_chr1_183 | 3 |
| NC_002505 | 1902135 | 1903009 | 875 | 1902587 | 268 | 28.26749 | 2.13243 | 26.1974 | peaks_peak_387 | Peak_chr1_184 | 195 |
| NC_002505 | 1904036 | 1905934 | 1899 | 1904429 | 371 | 67.54165 | 2.87423 | 64.07333 | peaks_peak_388 | Peak_chr1_185 | 20 |
| NC_002505 | 1907479 | 1907775 | 297 | 1907618 | 232 | 14.66618 | 1.74422 | 12.93692 | peaks_peak_389 | Peak_chr1_186 | 339 |
| NC_002505 | 1919382 | 1919980 | 599 | 1919665 | 221 | 13.60292 | 1.72636 | 11.90901 | peaks_peak_392 | Peak_chr1_187 | 363 |
| NC_002505 | 1924876 | 1925272 | 397 | 1925049 | 245 | 22.21598 | 1.99424 | 20.28794 | peaks_peak_393 | Peak_chr1_188 | 240 |
| NC_002505 | 1926525 | 1927012 | 488 | 1926646 | 223 | 14.25346 | 1.74837 | 12.53788 | peaks_peak_394 | Peak_chr1_189 | 350 |
| NC_002505 | 1934823 | 1935255 | 433 | 1934950 | 256 | 20.72513 | 1.90961 | 18.83448 | peaks_peak_397 | Peak_chr1_190 | 257 |
| NC_002505 | 1935561 | 1935943 | 383 | 1935674 | 223 | 12.27462 | 1.66721 | 10.62531 | peaks_peak_398 | Peak_chr1_191 | 398 |
| NC_002505 | 1940285 | 1940580 | 296 | 1940423 | 220 | 11.81862 | 1.65424 | 10.18675 | peaks_peak_401 | Peak_chr1_192 | 417 |
| NC_002505 | 1940817 | 1942085 | 1269 | 1941333 | 374 | 64.14154 | 2.77026 | 60.90097 | peaks_peak_402 | Peak_chr1_193 | 30 |
| NC_002505 | 1946678 | 1948278 | 1601 | 1947249 | 325 | 48.4058 | 2.5445 | 45.82543 | peaks_peak_403 | Peak_chr1_194 | 81 |
| NC_002505 | 1950284 | 1950600 | 317 | 1950429 | 223 | 13.85928 | 1.73231 | 12.15692 | peaks_peak_405 | Peak_chr1_195 | 354 |
| NC_002505 | 1952639 | 1953348 | 710 | 1952854 | 362 | 68.41977 | 2.94244 | 64.8958 | peaks_peak_406 | Peak_chr1_196 | 17 |
| NC_002505 | 1958958 | 1959516 | 559 | 1959334 | 234 | 14.02983 | 1.71496 | 12.32189 | peaks_peak_407 | Peak_chr1_197 | 351 |
| NC_002505 | 1966906 | 1967560 | 655 | 1967065 | 239 | 15.49718 | 1.76061 | 13.74299 | peaks_peak_408 | Peak_chr1_198 | 323 |
| NC_002505 | 1970976 | 1971512 | 537 | 1971352 | 253 | 20.17913 | 1.89822 | 18.30221 | peaks_peak_409 | Peak_chr1_199 | 265 |
| NC_002505 | 1977073 | 1978060 | 988 | 1977293 | 329 | 44.92846 | 2.42888 | 42.45257 | peaks_peak_410 | Peak_chr1_200 | 102 |
| NC_002505 | 1980655 | 1981227 | 573 | 1980811 | 293 | 32.05616 | 2.17303 | 29.89841 | peaks_peak_411 | Peak_chr1_201 | 174 |
| NC_002505 | 1984013 | 1984938 | 926 | 1984283 | 365 | 61.15433 | 2.73013 | 58.09334 | peaks_peak_412 | Peak_chr1_202 | 39 |
| NC_002505 | 2003225 | 2003449 | 225 | 2003345 | 228 | 13.69969 | 1.71473 | 12.00307 | peaks_peak_416 | Peak_chr1_203 | 360 |
| NC_002505 | 2004154 | 2005117 | 964 | 2004306 | 263 | 23.59181 | 1.98955 | 21.63222 | peaks_peak_417 | Peak_chr1_204 | 228 |
| NC_002505 | 2009528 | 2009970 | 443 | 2009796 | 314 | 34.02568 | 2.16725 | 31.82214 | peaks_peak_418 | Peak_chr1_205 | 160 |
| NC_002505 | 2014072 | 2014464 | 393 | 2014220 | 243 | 16.81041 | 1.80078 | 15.02028 | peaks_peak_419 | Peak_chr1_206 | 300 |
| NC_002505 | 2025471 | 2026004 | 534 | 2025618 | 232 | 13.07351 | 1.68141 | 11.39682 | peaks_peak_420 | Peak_chr1_207 | 373 |
| NC_002505 | 2029629 | 2029960 | 332 | 2029796 | 296 | 32.25659 | 2.16967 | 30.0943 | peaks_peak_421 | Peak_chr1_208 | 172 |
| NC_002505 | 2044258 | 2044513 | 256 | 2044386 | 224 | 12.04996 | 1.65591 | 10.4093 | peaks_peak_423 | Peak_chr1_209 | 405 |
| NC_002505 | 2048877 | 2049556 | 680 | 2049272 | 382 | 66.09913 | 2.78656 | 62.73773 | peaks_peak_424 | Peak_chr1_210 | 25 |
| NC_002505 | 2054694 | 2055066 | 373 | 2054821 | 251 | 17.78704 | 1.81775 | 15.97208 | peaks_peak_425 | Peak_chr1_211 | 287 |
| NC_002505 | 2056865 | 2057197 | 333 | 2057030 | 262 | 19.9925 | 1.86906 | 18.12009 | peaks_peak_426 | Peak_chr1_212 | 266 |
| NC_002505 | 2081505 | 2082065 | 561 | 2081776 | 323 | 41.88313 | 2.36384 | 39.4894 | peaks_peak_430 | Peak_chr1_213 | 117 |

| | | | | | | | | | | | |
|---|---|---|---|---|---|---|---|---|---|---|---|
| NC_002505 | 2083971 | 2084520 | 550 | 2084160 | 320 | 41.00801 | 2.34928 | 38.63639 | peaks_peak_431 | Peak_chr1_214 | 119 |
| NC_002505 | 2088752 | 2089286 | 535 | 2088930 | 245 | 16.74436 | 1.79368 | 14.95598 | peaks_peak_433 | Peak_chr1_215 | 302 |
| NC_002505 | 2093453 | 2093772 | 320 | 2093614 | 319 | 39.02094 | 2.29546 | 36.69834 | peaks_peak_434 | Peak_chr1_216 | 130 |
| NC_002505 | 2097286 | 2097620 | 335 | 2097446 | 280 | 26.2572 | 2.02954 | 24.23336 | peaks_peak_435 | Peak_chr1_217 | 212 |
| NC_002505 | 2098708 | 2098946 | 239 | 2098837 | 227 | 12.93994 | 1.68624 | 11.26735 | peaks_peak_437 | Peak_chr1_218 | 376 |
| NC_002505 | 2110710 | 2111106 | 397 | 2110857 | 266 | 21.45356 | 1.90893 | 19.54412 | peaks_peak_438 | Peak_chr1_219 | 246 |
| NC_002505 | 2112731 | 2113460 | 730 | 2113103 | 399 | 64.40525 | 2.67552 | 61.15176 | peaks_peak_439 | Peak_chr1_220 | 28 |
| NC_002505 | 2150420 | 2150680 | 261 | 2150572 | 237 | 12.03379 | 1.63081 | 10.39366 | peaks_peak_442 | Peak_chr1_221 | 406 |
| NC_002505 | 2151626 | 2152376 | 751 | 2152020 | 280 | 25.82623 | 2.0157 | 23.81227 | peaks_peak_443 | Peak_chr1_222 | 213 |
| NC_002505 | 2163389 | 2163841 | 453 | 2163682 | 232 | 13.5519 | 1.70036 | 11.8597 | peaks_peak_448 | Peak_chr1_223 | 364 |
| NC_002505 | 2167590 | 2169159 | 1570 | 2168572 | 386 | 70.45755 | 2.88142 | 66.79388 | peaks_peak_449 | Peak_chr1_224 | 13 |
| NC_002505 | 2178714 | 2179002 | 289 | 2178857 | 225 | 11.6279 | 1.63651 | 10.00375 | peaks_peak_450 | Peak_chr1_225 | 424 |
| NC_002505 | 2179714 | 2180002 | 289 | 2179857 | 230 | 12.48497 | 1.66185 | 10.82839 | peaks_peak_451 | Peak_chr1_226 | 391 |
| NC_002505 | 2190152 | 2191083 | 932 | 2190742 | 379 | 62.88959 | 2.7166 | 59.7188 | peaks_peak_453 | Peak_chr1_227 | 36 |
| NC_002505 | 2192242 | 2193699 | 1458 | 2192947 | 389 | 68.01542 | 2.80548 | 64.51865 | peaks_peak_454 | Peak_chr1_228 | 18 |
| NC_002505 | 2228853 | 2229429 | 577 | 2229140 | 341 | 45.22181 | 2.39411 | 42.73884 | peaks_peak_459 | Peak_chr1_229 | 101 |
| NC_002505 | 2255133 | 2255592 | 460 | 2255340 | 306 | 32.89452 | 2.15807 | 30.71711 | peaks_peak_464 | Peak_chr1_230 | 166 |
| NC_002505 | 2258349 | 2258668 | 320 | 2258525 | 252 | 15.47674 | 1.73218 | 13.72293 | peaks_peak_465 | Peak_chr1_231 | 324 |
| NC_002505 | 2262885 | 2263482 | 598 | 2263274 | 353 | 44.33174 | 2.33079 | 41.87215 | peaks_peak_466 | Peak_chr1_232 | 106 |
| NC_002505 | 2307372 | 2307875 | 504 | 2307741 | 230 | 11.76643 | 1.63282 | 10.13671 | peaks_peak_467 | Peak_chr1_233 | 418 |
| NC_002505 | 2311812 | 2312244 | 433 | 2311927 | 250 | 16.1283 | 1.76001 | 14.35619 | peaks_peak_468 | Peak_chr1_234 | 313 |
| NC_002505 | 2316422 | 2316843 | 422 | 2316566 | 288 | 26.5921 | 2.01805 | 24.56085 | peaks_peak_469 | Peak_chr1_235 | 208 |
| NC_002505 | 2341810 | 2342025 | 216 | 2341913 | 237 | 12.03379 | 1.63081 | 10.39366 | peaks_peak_470 | Peak_chr1_236 | 406 |
| NC_002505 | 2350313 | 2350657 | 345 | 2350498 | 256 | 16.99026 | 1.77808 | 15.19549 | peaks_peak_471 | Peak_chr1_237 | 296 |
| NC_002505 | 2359620 | 2359826 | 207 | 2359706 | 249 | 12.81151 | 1.63963 | 11.14335 | peaks_peak_472 | Peak_chr1_238 | 383 |
| NC_002505 | 2364278 | 2364957 | 680 | 2364620 | 379 | 60.64729 | 2.65923 | 57.61192 | peaks_peak_473 | Peak_chr1_239 | 42 |
| NC_002505 | 2365783 | 2366768 | 986 | 2366407 | 356 | 50.36884 | 2.47932 | 47.72803 | peaks_peak_475 | Peak_chr1_240 | 75 |
| NC_002505 | 2375759 | 2376218 | 460 | 2375948 | 279 | 22.34035 | 1.90619 | 20.40896 | peaks_peak_476 | Peak_chr1_241 | 238 |
| NC_002505 | 2389003 | 2389297 | 295 | 2389145 | 258 | 13.78647 | 1.65985 | 12.08658 | peaks_peak_477 | Peak_chr1_242 | 357 |
| NC_002505 | 2397666 | 2398537 | 872 | 2398059 | 390 | 64.22018 | 2.70584 | 60.97736 | peaks_peak_478 | Peak_chr1_243 | 29 |
| NC_002505 | 2399980 | 2400326 | 347 | 2400153 | 328 | 37.68598 | 2.22895 | 35.39604 | peaks_peak_479 | Peak_chr1_244 | 139 |
| NC_002505 | 2409899 | 2410204 | 306 | 2410063 | 288 | 21.05943 | 1.84509 | 19.16049 | peaks_peak_480 | Peak_chr1_245 | 252 |
| NC_002505 | 2414410 | 2414661 | 252 | 2414536 | 263 | 16.50026 | 1.74641 | 14.71828 | peaks_peak_481 | Peak_chr1_246 | 306 |
| NC_002505 | 2429431 | 2430257 | 827 | 2430080 | 314 | 34.20999 | 2.17257 | 32.00211 | peaks_peak_482 | Peak_chr1_247 | 158 |
| NC_002505 | 2432558 | 2433519 | 962 | 2433268 | 392 | 66.52154 | 2.75525 | 63.12645 | peaks_peak_483 | Peak_chr1_248 | 22 |
| NC_002505 | 2439507 | 2439925 | 419 | 2439751 | 349 | 48.61334 | 2.45767 | 46.02553 | peaks_peak_484 | Peak_chr1_249 | 80 |
| NC_002505 | 2441165 | 2442264 | 1100 | 2441834 | 301 | 30.7536 | 2.10901 | 28.62682 | peaks_peak_485 | Peak_chr1_250 | 180 |
| NC_002505 | 2444672 | 2444958 | 287 | 2444812 | 244 | 13.3911 | 1.67048 | 11.704 | peaks_peak_486 | Peak_chr1_251 | 367 |
| NC_002505 | 2447236 | 2447614 | 379 | 2447431 | 307 | 27.771 | 2.0044 | 25.71236 | peaks_peak_488 | Peak_chr1_252 | 202 |
| NC_002505 | 2450804 | 2451408 | 605 | 2450947 | 264 | 16.70915 | 1.75165 | 14.9219 | peaks_peak_489 | Peak_chr1_253 | 303 |
| NC_002505 | 2463857 | 2464193 | 337 | 2464037 | 282 | 22.93151 | 1.91792 | 20.98681 | peaks_peak_491 | Peak_chr1_254 | 231 |
| NC_002505 | 2466074 | 2466329 | 256 | 2466183 | 241 | 12.59403 | 1.64549 | 10.93357 | peaks_peak_492 | Peak_chr1_255 | 388 |
| NC_002505 | 2469735 | 2469958 | 224 | 2469850 | 266 | 16.22054 | 1.73089 | 14.44592 | peaks_peak_493 | Peak_chr1_256 | 310 |
| NC_002505 | 2493348 | 2494176 | 829 | 2494013 | 267 | 17.26072 | 1.76455 | 15.45893 | peaks_peak_496 | Peak_chr1_257 | 293 |
| NC_002505 | 2506747 | 2507075 | 329 | 2506934 | 255 | 16.09884 | 1.74846 | 14.3277 | peaks_peak_499 | Peak_chr1_258 | 314 |
| NC_002505 | 2513015 | 2514934 | 1920 | 2513763 | 391 | 63.29185 | 2.67884 | 60.09526 | peaks_peak_501 | Peak_chr1_259 | 33 |
| NC_002505 | 2516932 | 2517455 | 524 | 2517313 | 246 | 12.32225 | 1.62629 | 10.6713 | peaks_peak_502 | Peak_chr1_260 | 395 |
| NC_002505 | 2547537 | 2547916 | 380 | 2547705 | 351 | 42.22856 | 2.28192 | 39.82602 | peaks_peak_505 | Peak_chr1_261 | 115 |
| NC_002505 | 2549158 | 2550239 | 1082 | 2549465 | 226 | 13.64807 | 1.71702 | 11.95295 | peaks_peak_506 | Peak_chr1_262 | 361 |
| NC_002505 | 2551522 | 2552640 | 1119 | 2552287 | 252 | 19.78759 | 1.88689 | 17.92083 | peaks_peak_508 | Peak_chr1_263 | 269 |
| NC_002505 | 2558398 | 2558842 | 445 | 2558585 | 322 | 33.66742 | 2.13403 | 31.47216 | peaks_peak_509 | Peak_chr1_264 | 161 |
| NC_002505 | 2560731 | 2561108 | 378 | 2560933 | 362 | 44.89297 | 2.31753 | 42.41769 | peaks_peak_510 | Peak_chr1_265 | 103 |
| NC_002505 | 2564882 | 2565175 | 294 | 2565034 | 263 | 16.88326 | 1.75969 | 15.09127 | peaks_peak_511 | Peak_chr1_266 | 297 |
| NC_002505 | 2572267 | 2572771 | 505 | 2572637 | 273 | 14.90072 | 1.67318 | 13.16413 | peaks_peak_513 | Peak_chr1_267 | 334 |
| NC_002505 | 2577059 | 2577416 | 358 | 2577232 | 316 | 31.51037 | 2.0893 | 29.36487 | peaks_peak_514 | Peak_chr1_268 | 177 |
| NC_002505 | 2580487 | 2580826 | 340 | 2580693 | 290 | 21.60489 | 1.85786 | 19.69174 | peaks_peak_515 | Peak_chr1_269 | 245 |
| NC_002505 | 2588413 | 2588649 | 237 | 2588529 | 248 | 13.43775 | 1.66485 | 11.74939 | peaks_peak_516 | Peak_chr1_270 | 366 |
| NC_002505 | 2590329 | 2590821 | 493 | 2590597 | 347 | 42.80388 | 2.30935 | 40.38606 | peaks_peak_517 | Peak_chr1_271 | 112 |
| NC_002505 | 2607350 | 2607804 | 455 | 2607543 | 314 | 31.13592 | 2.08395 | 28.9999 | peaks_peak_519 | Peak_chr1_272 | 179 |
| NC_002505 | 2633970 | 2634237 | 268 | 2634115 | 268 | 15.44893 | 1.70063 | 13.69605 | peaks_peak_525 | Peak_chr1_273 | 326 |
| NC_002505 | 2649097 | 2649669 | 573 | 2649303 | 274 | 17.6103 | 1.76239 | 15.79966 | peaks_peak_526 | Peak_chr1_274 | 291 |
| NC_002505 | 2653899 | 2654509 | 611 | 2654073 | 311 | 29.79268 | 2.05313 | 27.68839 | peaks_peak_527 | Peak_chr1_275 | 186 |
| NC_002505 | 2664714 | 2665004 | 291 | 2664854 | 259 | 16.78023 | 1.76432 | 14.99078 | peaks_peak_528 | Peak_chr1_276 | 301 |
| NC_002505 | 2686261 | 2686551 | 291 | 2686397 | 275 | 19.89759 | 1.83589 | 18.02765 | peaks_peak_534 | Peak_chr1_277 | 267 |
| NC_002505 | 2697164 | 2697449 | 286 | 2697326 | 258 | 14.35616 | 1.68032 | 12.6374 | peaks_peak_535 | Peak_chr1_278 | 349 |
| NC_002505 | 2701459 | 2702002 | 544 | 2701644 | 344 | 40.16289 | 2.24832 | 37.81338 | peaks_peak_538 | Peak_chr1_279 | 121 |
| NC_002505 | 2704958 | 2705447 | 490 | 2705260 | 371 | 46.49415 | 2.3308 | 43.97294 | peaks_peak_540 | Peak_chr1_280 | 93 |
| NC_002505 | 2706835 | 2707379 | 545 | 2707059 | 285 | 20.74975 | 1.84172 | 18.8584 | peaks_peak_541 | Peak_chr1_281 | 255 |
| NC_002505 | 2721640 | 2721864 | 225 | 2721777 | 246 | 11.92308 | 1.61104 | 10.28713 | peaks_peak_542 | Peak_chr1_282 | 412 |
| NC_002505 | 2749395 | 2749721 | 327 | 2749586 | 282 | 18.91072 | 1.78914 | 17.06665 | peaks_peak_543 | Peak_chr1_283 | 277 |
| NC_002505 | 2755117 | 2755448 | 332 | 2755296 | 281 | 19.11284 | 1.79767 | 17.26369 | peaks_peak_544 | Peak_chr1_284 | 276 |
| NC_002505 | 2805248 | 2805740 | 493 | 2805484 | 362 | 47.01187 | 2.37168 | 44.47338 | peaks_peak_547 | Peak_chr1_285 | 92 |
| NC_002505 | 2818278 | 2818581 | 304 | 2818443 | 304 | 26.87741 | 1.98549 | 24.83968 | peaks_peak_548 | Peak_chr1_286 | 205 |
| NC_002505 | 2826479 | 2826900 | 422 | 2826738 | 272 | 15.39293 | 1.69162 | 13.6417 | peaks_peak_550 | Peak_chr1_287 | 327 |
| NC_002505 | 2830424 | 2830761 | 338 | 2830609 | 347 | 38.72068 | 2.20156 | 36.40562 | peaks_peak_551 | Peak_chr1_288 | 132 |
| NC_002505 | 2832131 | 2832745 | 615 | 2832496 | 393 | 55.9684 | 2.49314 | 53.12749 | peaks_peak_552 | Peak_chr1_289 | 57 |
| NC_002505 | 2836481 | 2836846 | 366 | 2836654 | 294 | 22.76004 | 1.88483 | 20.8193 | peaks_peak_554 | Peak_chr1_290 | 235 |
| NC_002505 | 2845054 | 2845381 | 328 | 2845232 | 294 | 22.69947 | 1.88297 | 20.75995 | peaks_peak_555 | Peak_chr1_291 | 236 |
| NC_002505 | 2855130 | 2855952 | 823 | 2855511 | 388 | 54.50028 | 2.47415 | 51.71504 | peaks_peak_556 | Peak_chr1_292 | 62 |
| NC_002505 | 2864607 | 2865242 | 636 | 2864824 | 315 | 29.25192 | 2.02731 | 27.15967 | peaks_peak_558 | Peak_chr1_293 | 190 |
| NC_002505 | 2892820 | 2893190 | 371 | 2893002 | 339 | 36.97895 | 2.17763 | 34.70722 | peaks_peak_561 | Peak_chr1_294 | 143 |

| | | | | | | | | | | | |
|---|---|---|---|---|---|---|---|---|---|---|---|
| NC_002505 | 2909856 | 2910160 | 305 | 2909995 | 288 | 19.75394 | 1.80404 | 17.8879 | peaks_peak_562 | Peak_chr1_295 | 270 |
| NC_002505 | 2918258 | 2919188 | 931 | 2918675 | 393 | 56.04652 | 2.49501 | 53.20255 | peaks_peak_564 | Peak_chr1_296 | 55 |
| NC_002505 | 2931481 | 2932034 | 554 | 2931611 | 264 | 11.63325 | 1.57259 | 10.00887 | peaks_peak_567 | Peak_chr1_297 | 422 |
| NC_002505 | 2933226 | 2934008 | 783 | 2933453 | 310 | 12.87404 | 1.55504 | 11.2038 | peaks_peak_568 | Peak_chr1_298 | 381 |
| NC_002505 | 2940101 | 2940918 | 818 | 2940551 | 394 | 52.70198 | 2.41206 | 49.98661 | peaks_peak_571 | Peak_chr1_299 | 68 |
| NC_002505 | 2941937 | 2942457 | 521 | 2942239 | 370 | 47.48534 | 2.35859 | 44.93339 | peaks_peak_573 | Peak_chr1_300 | 86 |
| NC_002505 | 2949398 | 2949829 | 432 | 2949561 | 268 | 14.7944 | 1.67807 | 13.06071 | peaks_peak_574 | Peak_chr1_301 | 336 |
| NC_002505 | 2954151 | 2954488 | 338 | 2954354 | 307 | 24.69427 | 1.91413 | 22.70812 | peaks_peak_575 | Peak_chr1_302 | 223 |
| NC_002505 | 2960414 | 2961069 | 656 | 2960692 | 384 | 54.45431 | 2.48628 | 51.67185 | peaks_peak_578 | Peak_chr1_303 | 64 |
| NC_002506 | 73 | 906 | 834 | 598 | 281 | 24.5601 | 1.97245 | 22.57747 | peaks_peak_579 | Peak_chr2_1 | 224 |
| NC_002506 | 2889 | 3565 | 677 | 3223 | 395 | 62.51577 | 2.64456 | 59.3657 | peaks_peak_581 | Peak_chr2_2 | 37 |
| NC_002506 | 32517 | 32901 | 385 | 32696 | 306 | 34.35072 | 2.20127 | 32.13993 | peaks_peak_585 | Peak_chr2_3 | 155 |
| NC_002506 | 40382 | 40714 | 333 | 40572 | 324 | 39.21965 | 2.2846 | 36.8925 | peaks_peak_586 | Peak_chr2_4 | 127 |
| NC_002506 | 43050 | 43425 | 376 | 43238 | 338 | 45.32961 | 2.40754 | 42.84315 | peaks_peak_587 | Peak_chr2_5 | 100 |
| NC_002506 | 45037 | 45400 | 364 | 45217 | 232 | 12.74975 | 1.66854 | 11.0836 | peaks_peak_589 | Peak_chr2_6 | 384 |
| NC_002506 | 48167 | 48546 | 380 | 48302 | 233 | 13.12791 | 1.68157 | 11.44959 | peaks_peak_590 | Peak_chr2_7 | 372 |
| NC_002506 | 60697 | 61026 | 330 | 60884 | 322 | 32.52867 | 2.10202 | 30.35982 | peaks_peak_592 | Peak_chr2_8 | 169 |
| NC_002506 | 75330 | 75598 | 269 | 75479 | 230 | 12.07005 | 1.64512 | 10.42854 | peaks_peak_594 | Peak_chr2_9 | 404 |
| NC_002506 | 79135 | 79507 | 373 | 79340 | 335 | 44.70268 | 2.40082 | 42.23217 | peaks_peak_595 | Peak_chr2_10 | 105 |
| NC_002506 | 80197 | 80568 | 372 | 80388 | 309 | 33.42539 | 2.1647 | 31.23545 | peaks_peak_596 | Peak_chr2_11 | 162 |
| NC_002506 | 83962 | 84318 | 357 | 84121 | 235 | 14.48789 | 1.73066 | 12.76454 | peaks_peak_597 | Peak_chr2_12 | 347 |
| NC_002506 | 86222 | 86769 | 548 | 86414 | 348 | 53.08484 | 2.58272 | 50.35648 | peaks_peak_599 | Peak_chr2_13 | 67 |
| NC_002506 | 88942 | 89224 | 283 | 89058 | 229 | 13.79219 | 1.71627 | 12.09208 | peaks_peak_600 | Peak_chr2_14 | 355 |
| NC_002506 | 92145 | 92558 | 414 | 92389 | 334 | 47.8334 | 2.49205 | 45.27089 | peaks_peak_602 | Peak_chr2_15 | 84 |
| NC_002506 | 101796 | 102378 | 583 | 102202 | 257 | 20.20156 | 1.8887 | 18.3242 | peaks_peak_603 | Peak_chr2_16 | 264 |
| NC_002506 | 113127 | 113455 | 329 | 113293 | 296 | 29.51802 | 2.08603 | 27.41978 | peaks_peak_606 | Peak_chr2_17 | 187 |
| NC_002506 | 114472 | 115460 | 989 | 114985 | 274 | 25.58486 | 2.02477 | 23.57684 | peaks_peak_607 | Peak_chr2_18 | 215 |
| NC_002506 | 153218 | 154043 | 826 | 153646 | 399 | 71.20307 | 2.84268 | 67.47878 | peaks_peak_612 | Peak_chr2_19 | 12 |
| NC_002506 | 185471 | 185759 | 289 | 185628 | 304 | 27.9092 | 2.01605 | 25.84709 | peaks_peak_613 | Peak_chr2_20 | 200 |
| NC_002506 | 212515 | 213323 | 809 | 212704 | 325 | 49.0576 | 2.56352 | 46.45624 | peaks_peak_615 | Peak_chr2_21 | 78 |
| NC_002506 | 219869 | 220356 | 488 | 219989 | 236 | 14.70996 | 1.73709 | 12.97906 | peaks_peak_617 | Peak_chr2_22 | 338 |
| NC_002506 | 236497 | 236797 | 301 | 236676 | 222 | 12.92199 | 1.69606 | 11.25017 | peaks_peak_619 | Peak_chr2_23 | 380 |
| NC_002506 | 237693 | 238080 | 388 | 237908 | 279 | 26.87406 | 2.05225 | 24.83637 | peaks_peak_620 | Peak_chr2_24 | 206 |
| NC_002506 | 241776 | 242067 | 292 | 241936 | 221 | 11.85549 | 1.65378 | 10.2222 | peaks_peak_621 | Peak_chr2_25 | 415 |
| NC_002506 | 244139 | 245125 | 987 | 244353 | 319 | 38.34626 | 2.27606 | 36.04035 | peaks_peak_622 | Peak_chr2_26 | 134 |
| NC_002506 | 248673 | 248959 | 287 | 248796 | 241 | 14.57455 | 1.72127 | 12.84786 | peaks_peak_624 | Peak_chr2_27 | 341 |
| NC_002506 | 253335 | 253642 | 308 | 253451 | 248 | 18.12274 | 1.83704 | 16.29917 | peaks_peak_625 | Peak_chr2_28 | 285 |
| NC_002506 | 255593 | 256030 | 438 | 255839 | 366 | 60.82727 | 2.71713 | 57.78638 | peaks_peak_626 | Peak_chr2_29 | 41 |
| NC_002506 | 275460 | 275783 | 324 | 275655 | 271 | 23.35266 | 1.95949 | 21.39851 | peaks_peak_627 | Peak_chr2_30 | 229 |
| NC_002506 | 278891 | 279307 | 417 | 279130 | 366 | 58.57586 | 2.65752 | 55.63623 | peaks_peak_628 | Peak_chr2_31 | 48 |
| NC_002506 | 289860 | 290363 | 504 | 290109 | 396 | 54.60437 | 2.45095 | 51.81646 | peaks_peak_630 | Peak_chr2_32 | 61 |
| NC_002506 | 292292 | 292652 | 361 | 292428 | 228 | 11.74622 | 1.63572 | 10.11734 | peaks_peak_631 | Peak_chr2_33 | 419 |
| NC_002506 | 310117 | 310437 | 321 | 310260 | 298 | 36.10144 | 2.28089 | 33.84974 | peaks_peak_634 | Peak_chr2_34 | 147 |
| NC_002506 | 322819 | 323710 | 892 | 323139 | 234 | 14.57007 | 1.73603 | 12.84357 | peaks_peak_641 | Peak_chr2_35 | 342 |
| NC_002506 | 334843 | 335321 | 479 | 335044 | 342 | 54.47573 | 2.64627 | 51.69189 | peaks_peak_649 | Peak_chr2_36 | 63 |
| NC_002506 | 344826 | 345123 | 298 | 345005 | 216 | 12.1809 | 1.67787 | 10.53529 | peaks_peak_657 | Peak_chr2_37 | 400 |
| NC_002506 | 351921 | 353042 | 1122 | 352252 | 236 | 20.27205 | 1.94925 | 18.39255 | peaks_peak_660 | Peak_chr2_38 | 262 |
| NC_002506 | 365576 | 366112 | 537 | 365926 | 236 | 11.98309 | 1.6306 | 10.34495 | peaks_peak_663 | Peak_chr2_39 | 409 |
| NC_002506 | 373927 | 375021 | 1095 | 374096 | 248 | 21.41506 | 1.9562 | 19.50654 | peaks_peak_666 | Peak_chr2_40 | 247 |
| NC_002506 | 379893 | 380359 | 467 | 380136 | 217 | 11.70457 | 1.65549 | 10.07729 | peaks_peak_668 | Peak_chr2_41 | 420 |
| NC_002506 | 399891 | 400666 | 776 | 400418 | 318 | 51.13597 | 2.65638 | 48.47186 | peaks_peak_674 | Peak_chr2_42 | 73 |
| NC_002506 | 403548 | 405172 | 1625 | 404753 | 263 | 30.23884 | 2.21725 | 28.12459 | peaks_peak_676 | Peak_chr2_43 | 185 |
| NC_002506 | 409978 | 410361 | 384 | 410142 | 230 | 15.91808 | 1.79801 | 14.1521 | peaks_peak_678 | Peak_chr2_44 | 318 |
| NC_002506 | 410832 | 411408 | 577 | 411146 | 277 | 24.38416 | 1.97733 | 22.40575 | peaks_peak_679 | Peak_chr2_45 | 226 |
| NC_002506 | 412175 | 412655 | 481 | 412378 | 234 | 12.22156 | 1.64371 | 10.57436 | peaks_peak_680 | Peak_chr2_46 | 399 |
| NC_002506 | 420832 | 421183 | 352 | 421029 | 220 | 13.04854 | 1.70566 | 11.37273 | peaks_peak_681 | Peak_chr2_47 | 374 |
| NC_002506 | 427864 | 428746 | 883 | 428229 | 321 | 38.06873 | 2.26162 | 35.76943 | peaks_peak_683 | Peak_chr2_48 | 137 |
| NC_002506 | 431223 | 431972 | 750 | 431668 | 347 | 53.8356 | 2.60742 | 51.08102 | peaks_peak_685 | Peak_chr2_49 | 65 |
| NC_002506 | 438968 | 439300 | 333 | 439138 | 314 | 39.58649 | 2.32906 | 37.25007 | peaks_peak_688 | Peak_chr2_50 | 125 |
| NC_002506 | 443572 | 444200 | 629 | 443906 | 350 | 57.17034 | 2.68682 | 54.28672 | peaks_peak_689 | Peak_chr2_51 | 51 |
| NC_002506 | 472019 | 472326 | 308 | 472165 | 273 | 26.5208 | 2.05846 | 24.49101 | peaks_peak_692 | Peak_chr2_52 | 211 |
| NC_002506 | 473816 | 474116 | 301 | 473990 | 232 | 15.83577 | 1.7899 | 14.07216 | peaks_peak_694 | Peak_chr2_53 | 321 |
| NC_002506 | 485302 | 485776 | 475 | 485497 | 312 | 42.58051 | 2.42505 | 40.16882 | peaks_peak_695 | Peak_chr2_54 | 114 |
| NC_002506 | 491965 | 493769 | 1805 | 492436 | 392 | 78.44988 | 3.06093 | 73.86951 | peaks_peak_696 | Peak_chr2_55 | 2 |
| NC_002506 | 503142 | 503409 | 268 | 503262 | 216 | 12.74059 | 1.70162 | 11.07488 | peaks_peak_697 | Peak_chr2_56 | 385 |
| NC_002506 | 509601 | 510294 | 694 | 509747 | 277 | 27.92431 | 2.09225 | 25.86183 | peaks_peak_700 | Peak_chr2_57 | 199 |
| NC_002506 | 510930 | 511268 | 339 | 511082 | 237 | 17.03689 | 1.82382 | 15.24093 | peaks_peak_701 | Peak_chr2_58 | 295 |
| NC_002506 | 512263 | 512810 | 548 | 512392 | 214 | 12.08645 | 1.67812 | 10.44449 | peaks_peak_702 | Peak_chr2_59 | 403 |
| NC_002506 | 517158 | 517769 | 612 | 517396 | 348 | 50.96916 | 2.52501 | 48.30945 | peaks_peak_703 | Peak_chr2_60 | 74 |
| NC_002506 | 520842 | 521969 | 1128 | 521410 | 345 | 55.9913 | 2.6758 | 53.14935 | peaks_peak_704 | Peak_chr2_61 | 56 |
| NC_002506 | 528860 | 530078 | 1219 | 529512 | 391 | 74.32182 | 2.95816 | 70.20187 | peaks_peak_705 | Peak_chr2_62 | 6 |
| NC_002506 | 532096 | 532665 | 570 | 532338 | 321 | 45.41952 | 2.4736 | 42.93078 | peaks_peak_706 | Peak_chr2_63 | 99 |
| NC_002506 | 539914 | 540318 | 405 | 540090 | 329 | 47.24244 | 2.49476 | 44.69712 | peaks_peak_708 | Peak_chr2_64 | 89 |
| NC_002506 | 564922 | 565224 | 303 | 565081 | 237 | 20.60973 | 1.95901 | 18.72177 | peaks_peak_712 | Peak_chr2_65 | 259 |
| NC_002506 | 591822 | 592134 | 313 | 591970 | 237 | 11.70062 | 1.61764 | 10.0735 | peaks_peak_715 | Peak_chr2_66 | 421 |
| NC_002506 | 592351 | 592789 | 439 | 592641 | 269 | 27.16223 | 2.09209 | 25.11757 | peaks_peak_716 | Peak_chr2_67 | 203 |
| NC_002506 | 598182 | 598989 | 808 | 598364 | 238 | 18.63022 | 1.8815 | 16.79317 | peaks_peak_717 | Peak_chr2_68 | 280 |
| NC_002506 | 600066 | 601115 | 1050 | 600671 | 395 | 85.00883 | 3.21892 | 78.50765 | peaks_peak_719 | Peak_chr2_69 | 1 |
| NC_002506 | 601927 | 602550 | 624 | 602146 | 308 | 39.80518 | 2.35717 | 37.46393 | peaks_peak_720 | Peak_chr2_70 | 124 |
| NC_002506 | 606691 | 607009 | 319 | 606836 | 293 | 35.49644 | 2.27994 | 33.25877 | peaks_peak_722 | Peak_chr2_71 | 149 |
| NC_002506 | 611001 | 611334 | 334 | 611160 | 335 | 48.08622 | 2.4953 | 45.51633 | peaks_peak_724 | Peak_chr2_72 | 83 |

| | | | | | | | | | | | |
|---|---|---|---|---|---|---|---|---|---|---|---|
| NC_002506 | 621877 | 622829 | 953 | 622375 | 379 | 64.61762 | 2.76116 | 61.35179 | peaks_peak_727 | Peak_chr2_73 | 27 |
| NC_002506 | 634670 | 635098 | 429 | 634853 | 302 | 40.09607 | 2.38868 | 37.74814 | peaks_peak_728 | Peak_chr2_74 | 123 |
| NC_002506 | 649341 | 650010 | 670 | 649596 | 302 | 34.90301 | 2.23074 | 32.67854 | peaks_peak_730 | Peak_chr2_75 | 153 |
| NC_002506 | 658660 | 658936 | 277 | 658787 | 220 | 12.60279 | 1.6871 | 10.94198 | peaks_peak_731 | Peak_chr2_76 | 387 |
| NC_002506 | 665620 | 666163 | 544 | 665856 | 368 | 63.08979 | 2.76871 | 59.9067 | peaks_peak_734 | Peak_chr2_77 | 35 |
| NC_002506 | 673489 | 673883 | 395 | 673678 | 318 | 46.07541 | 2.50497 | 43.56647 | peaks_peak_735 | Peak_chr2_78 | 95 |
| NC_002506 | 683255 | 683514 | 260 | 683380 | 244 | 15.30901 | 1.74261 | 13.56002 | peaks_peak_738 | Peak_chr2_79 | 328 |
| NC_002506 | 703420 | 703676 | 257 | 703558 | 218 | 12.57471 | 1.69022 | 10.91489 | peaks_peak_740 | Peak_chr2_80 | 389 |
| NC_002506 | 717822 | 718211 | 390 | 718055 | 272 | 25.22093 | 2.01852 | 23.22158 | peaks_peak_741 | Peak_chr2_81 | 217 |
| NC_002506 | 733266 | 733624 | 359 | 733437 | 339 | 47.05308 | 2.45146 | 44.51347 | peaks_peak_742 | Peak_chr2_82 | 91 |
| NC_002506 | 735567 | 736889 | 1323 | 736198 | 243 | 13.2017 | 1.66515 | 11.52116 | peaks_peak_743 | Peak_chr2_83 | 370 |
| NC_002506 | 757837 | 758354 | 518 | 758078 | 330 | 44.88443 | 2.42394 | 42.40934 | peaks_peak_745 | Peak_chr2_84 | 104 |
| NC_002506 | 776812 | 777195 | 384 | 777026 | 277 | 27.0724 | 2.06456 | 25.0301 | peaks_peak_746 | Peak_chr2_85 | 204 |
| NC_002506 | 787029 | 788437 | 1409 | 787470 | 372 | 55.55176 | 2.55585 | 52.72683 | peaks_peak_750 | Peak_chr2_86 | 58 |
| NC_002506 | 808798 | 809083 | 286 | 808944 | 289 | 30.3577 | 2.13279 | 28.24091 | peaks_peak_754 | Peak_chr2_87 | 182 |
| NC_002506 | 811361 | 811681 | 321 | 811541 | 249 | 18.19344 | 1.83717 | 16.36813 | peaks_peak_755 | Peak_chr2_88 | 283 |
| NC_002506 | 819916 | 820312 | 397 | 820067 | 220 | 12.30839 | 1.67479 | 10.65793 | peaks_peak_756 | Peak_chr2_89 | 396 |
| NC_002506 | 823846 | 824237 | 392 | 824086 | 256 | 16.50905 | 1.761 | 14.72679 | peaks_peak_757 | Peak_chr2_90 | 305 |
| NC_002506 | 826794 | 827131 | 338 | 826958 | 321 | 45.59758 | 2.47881 | 43.10437 | peaks_peak_758 | Peak_chr2_91 | 97 |
| NC_002506 | 829092 | 829602 | 511 | 829432 | 211 | 12.03206 | 1.68232 | 10.39208 | peaks_peak_759 | Peak_chr2_92 | 408 |
| NC_002506 | 830091 | 830368 | 278 | 830248 | 237 | 17.70211 | 1.84907 | 15.88927 | peaks_peak_760 | Peak_chr2_93 | 290 |
| NC_002506 | 870731 | 871263 | 533 | 871071 | 284 | 28.90216 | 2.10226 | 26.81796 | peaks_peak_765 | Peak_chr2_94 | 191 |
| NC_002506 | 881845 | 882203 | 359 | 882059 | 270 | 24.86878 | 2.01257 | 22.87787 | peaks_peak_767 | Peak_chr2_95 | 220 |
| NC_002506 | 894453 | 894807 | 355 | 894671 | 244 | 14.91848 | 1.72801 | 13.18128 | peaks_peak_768 | Peak_chr2_96 | 332 |
| NC_002506 | 899626 | 899887 | 262 | 899732 | 244 | 17.77796 | 1.83421 | 15.96319 | peaks_peak_769 | Peak_chr2_97 | 288 |
| NC_002506 | 901578 | 902158 | 581 | 901906 | 271 | 26.55833 | 2.06575 | 24.52762 | peaks_peak_770 | Peak_chr2_98 | 209 |
| NC_002506 | 902582 | 903156 | 575 | 902722 | 243 | 18.37082 | 1.85863 | 16.54076 | peaks_peak_771 | Peak_chr2_99 | 282 |
| NC_002506 | 906318 | 906741 | 424 | 906437 | 232 | 13.73932 | 1.70777 | 12.0412 | peaks_peak_772 | Peak_chr2_100 | 359 |
| NC_002506 | 908228 | 908535 | 308 | 908391 | 249 | 18.18457 | 1.83685 | 16.3594 | peaks_peak_773 | Peak_chr2_101 | 284 |
| NC_002506 | 910786 | 911257 | 472 | 911031 | 391 | 60.1454 | 2.60134 | 57.13753 | peaks_peak_775 | Peak_chr2_102 | 45 |
| NC_002506 | 913971 | 914268 | 298 | 914133 | 276 | 25.06067 | 2.00202 | 23.0656 | peaks_peak_776 | Peak_chr2_103 | 218 |
| NC_002506 | 918610 | 918989 | 380 | 918776 | 330 | 43.63111 | 2.38862 | 41.19163 | peaks_peak_777 | Peak_chr2_104 | 110 |
| NC_002506 | 919332 | 919937 | 606 | 919500 | 243 | 16.41068 | 1.7859 | 14.63103 | peaks_peak_778 | Peak_chr2_105 | 308 |
| NC_002506 | 942238 | 942540 | 303 | 942395 | 232 | 12.17506 | 1.64558 | 10.52957 | peaks_peak_780 | Peak_chr2_106 | 401 |
| NC_002506 | 949426 | 949704 | 279 | 949541 | 228 | 12.42931 | 1.66353 | 10.7747 | peaks_peak_781 | Peak_chr2_107 | 393 |
| NC_002506 | 954872 | 955200 | 329 | 955066 | 259 | 20.23023 | 1.88466 | 18.3518 | peaks_peak_782 | Peak_chr2_108 | 263 |
| NC_002506 | 957417 | 957793 | 377 | 957557 | 287 | 21.2352 | 1.85276 | 19.33132 | peaks_peak_783 | Peak_chr2_109 | 249 |
| NC_002506 | 958930 | 959228 | 299 | 959079 | 256 | 18.84393 | 1.84354 | 17.00179 | peaks_peak_784 | Peak_chr2_110 | 278 |
| NC_002506 | 962782 | 963037 | 256 | 962896 | 237 | 13.91362 | 1.7042 | 12.20993 | peaks_peak_785 | Peak_chr2_111 | 353 |
| NC_002506 | 963455 | 963760 | 306 | 963601 | 247 | 16.3119 | 1.77324 | 14.53477 | peaks_peak_786 | Peak_chr2_112 | 309 |
| NC_002506 | 965164 | 965427 | 264 | 965278 | 253 | 14.43087 | 1.69223 | 12.70965 | peaks_peak_787 | Peak_chr2_113 | 348 |
| NC_002506 | 976203 | 976854 | 652 | 976629 | 390 | 66.29358 | 2.75776 | 62.91476 | peaks_peak_789 | Peak_chr2_114 | 24 |
| NC_002506 | 1013415 | 1014018 | 604 | 1013567 | 248 | 15.90752 | 1.75622 | 14.14206 | peaks_peak_793 | Peak_chr2_115 | 320 |
| NC_002506 | 1015396 | 1016124 | 729 | 1015893 | 304 | 33.36004 | 2.17802 | 31.17187 | peaks_peak_794 | Peak_chr2_116 | 163 |
| NC_002506 | 1026643 | 1026985 | 343 | 1026828 | 295 | 29.27958 | 2.08161 | 27.18674 | peaks_peak_796 | Peak_chr2_117 | 189 |
| NC_002506 | 1029255 | 1029538 | 284 | 1029347 | 233 | 11.93217 | 1.634 | 10.29596 | peaks_peak_798 | Peak_chr2_118 | 411 |
| NC_002506 | 1037451 | 1037860 | 410 | 1037652 | 343 | 46.28771 | 2.41614 | 43.77232 | peaks_peak_800 | Peak_chr2_119 | 94 |
| NC_002506 | 1039473 | 1039846 | 374 | 1039662 | 348 | 49.39413 | 2.48233 | 46.78255 | peaks_peak_801 | Peak_chr2_120 | 77 |
| NC_002506 | 1069967 | 1070202 | 236 | 1070059 | 246 | 16.17197 | 1.7703 | 14.39874 | peaks_peak_803 | Peak_chr2_121 | 311 |

#### Supplementary Table S4. List of genes differentially regulated by overexpression of VxrB(D78E)

### $C_{average}$ = Average value (C1+C2+C3/3) from triplicate control samples
### $E_{average}$ = Average value (E1+E2+E3/3) from triplicate samples of VxrB overexpression
### FC = Fold change ($E_{average}/C_{average}$)

| DEG, padj<0.001 | Control | | | VxrB overexpression | | | | | | | | | |
|---|---|---|---|---|---|---|---|---|---|---|---|---|---|
| Gene ID | C1 | C2 | C3 | E1 | E2 | E3 | *p*-value | $p_{adj}$ | $C_{average}$ | $E_{average}$ | FC | Log2FC | 0.5Log2(C*E) |
| VC0030 | 132.909 | 145.7183 | 167.8967 | 285.39227 | 256.0695 | 289.52969 | 1.42E-07 | 1.04E-06 | 148.8414 | 276.9972 | 1.861023 | 0.896096 | 2.307599 |
| VC0031 | 1741.4219 | 2295.064 | 2250.866 | 4119.7093 | 4012.082 | 3752.4194 | 1.17E-08 | 9.47E-08 | 2095.784 | 3961.404 | 1.890178 | 0.918522 | 3.459598 |
| VC0038 | 2695.855 | 2947.271 | 2957.431 | 5958.3336 | 5857.346 | 5698.8616 | 1.75E-28 | 4.26E-27 | 2866.852 | 5838.181 | 2.036443 | 1.026051 | 3.611841 |
| VC0048 | 3370.8653 | 3316.267 | 3349.19 | 1207.2709 | 1133.743 | 1101.7417 | 2.07E-64 | 1.25E-62 | 3345.441 | 1147.585 | 0.34303 | -1.5436 | 3.292119 |
| VC0049 | 7835.352 | 7269.465 | 7308.754 | 3158.8202 | 3627.978 | 3278.4698 | 7.52E-26 | 1.64E-24 | 7471.19 | 3355.089 | 0.44907 | -1.15499 | 3.699547 |
| VC0050 | 2103.5204 | 1681.637 | 2284.095 | 1111.7979 | 1221.706 | 961.27678 | 2.66E-06 | 1.64E-05 | 2023.084 | 1098.26 | 0.542864 | -0.88134 | 3.17336 |
| VC0054 | 882.22275 | 956.5704 | 871.8387 | 642.6459 | 637.2418 | 551.34861 | 1.11E-05 | 6.25E-05 | 903.544 | 610.4121 | 0.675575 | -0.56581 | 2.870786 |
| VC0088 | 1233.856 | 1049.407 | 1138.55 | 827.43227 | 820.009 | 857.12254 | 0.000138 | 0.000651 | 1140.604 | 834.8546 | 0.731941 | -0.4502 | 2.989373 |
| VC0091 | 3084.1168 | 4035.458 | 2585.785 | 1703.1143 | 1630.244 | 1645.4459 | 3.02E-06 | 1.84E-05 | 3235.12 | 1659.602 | 0.512995 | -0.96298 | 3.364947 |
| VC0095 | 1093.6213 | 1287.962 | 1014.376 | 669.33727 | 710.5441 | 638.30307 | 5.9E-07 | 3.96E-06 | 1131.986 | 672.7282 | 0.59429 | -0.75076 | 2.94084 |
| VC0099 | 2420.6183 | 2657.009 | 2086.467 | 1329.4352 | 1272.529 | 1206.8515 | 3.13E-10 | 2.97E-09 | 2388.031 | 1269.605 | 0.531653 | -0.91144 | 3.240854 |
| VC0100 | 1414.9053 | 1621.704 | 1296.827 | 745.305 | 633.3323 | 688.94688 | 2.53E-12 | 2.87E-11 | 1444.479 | 689.1947 | 0.477123 | -1.06757 | 2.999027 |
| VC0103 | 758.73249 | 752.0947 | 724.0546 | 480.44454 | 390.9459 | 418.52806 | 5.08E-09 | 4.2E-08 | 744.9606 | 429.9728 | 0.577175 | -0.79292 | 2.752787 |
| VC0130 | 3285.0501 | 2068.26 | 3430.515 | 1000.9261 | 1476.798 | 1445.7373 | 3.18E-05 | 0.00017 | 2927.942 | 1307.82 | 0.446669 | -1.16272 | 3.291555 |
| VC0131 | 308.72564 | 457.1325 | 319.1787 | 1503.9557 | 1450.409 | 1475.3592 | 3.25E-24 | 6.59E-23 | 361.6789 | 1476.575 | 4.082556 | 2.029473 | 2.863789 |
| VC0132 | 2148.5211 | 2727.518 | 2324.32 | 9793.6772 | 9780.488 | 9425.4813 | 8.21E-49 | 3.12E-47 | 2400.12 | 9666.549 | 4.027528 | 2.009894 | 3.682752 |
| VC0135 | 2036.5427 | 1550.02 | 1890.587 | 1092.2927 | 941.2022 | 1031.0315 | 2.77E-07 | 1.95E-06 | 1825.717 | 1021.509 | 0.559511 | -0.83776 | 3.135338 |
| VC0136 | 1575.024 | 1059.983 | 1517.192 | 572.83772 | 552.211 | 652.63623 | 6.92E-09 | 5.67E-08 | 1384.066 | 592.5617 | 0.428131 | -1.22388 | 2.956945 |
| VC0137 | 2921.905 | 2096.464 | 2765.924 | 1245.2548 | 1301.85 | 1225.0068 | 2.52E-09 | 2.15E-08 | 2594.764 | 1257.37 | 0.48458 | -1.04519 | 3.256781 |
| VC0139 | 752.45333 | 713.3148 | 775.6479 | 436.30113 | 486.7276 | 504.52698 | 1.52E-07 | 1.11E-06 | 747.1387 | 475.8519 | 0.636899 | -0.65086 | 2.775437 |
| VC0140 | 122.44373 | 133.9669 | 103.1865 | 46.196591 | 45.93614 | 37.266197 | 3.69E-10 | 3.49E-09 | 119.8657 | 43.13298 | 0.359844 | -1.47456 | 1.856752 |
| VC0147 | 3549.8215 | 3983.751 | 3927.209 | 8687.0122 | 9054.306 | 8091.5426 | 2.74E-26 | 6.05E-25 | 3820.261 | 8610.954 | 2.254022 | 1.172502 | 3.758572 |
| VC0148 | 2068.985 | 2312.691 | 2151.177 | 4937.9022 | 4658.12 | 4711.7851 | 1.31E-30 | 3.49E-29 | 2177.618 | 4769.269 | 2.190132 | 1.131018 | 3.508217 |
| VC0149 | 3080.9772 | 3476.087 | 3299.346 | 8847.1604 | 7651.788 | 7655.8147 | 2E-25 | 4.31E-24 | 3285.47 | 8051.588 | 2.450665 | 1.293173 | 3.71124 |
| VC0157 | 718.96445 | 910.7396 | 782.6436 | 50658.155 | 51820.85 | 52343.719 | 0 | 0 | 804.1159 | 51607.58 | 64.17928 | 6.004036 | 3.809016 |
| VC0161 | 591.28808 | 428.929 | 596.3832 | 273.07318 | 319.5982 | 338.26241 | 9.33E-05 | 0.000459 | 538.8667 | 310.3113 | 0.575859 | -0.79621 | 2.611639 |
| VC0164 | 23693.384 | 30294.14 | 22053.93 | 43361.147 | 41974.88 | 41486.833 | 1.08E-05 | 6.12E-05 | 25347.15 | 42274.29 | 1.667812 | 0.737957 | 4.515003 |
| VC0182 | 193.6076 | 115.1645 | 178.3903 | 52.356136 | 80.1439 | 53.510437 | 6.6E-06 | 3.87E-05 | 162.3875 | 62.00349 | 0.381824 | -1.38902 | 2.001484 |
| VC0183 | 232.32912 | 216.2272 | 297.3171 | 84.180454 | 101.6459 | 108.93196 | 1.06E-09 | 9.37E-09 | 248.6245 | 98.25278 | 0.395185 | -1.3394 | 2.193944 |
| VC0191 | 118.25762 | 101.0627 | 100.5631 | 166.30773 | 175.9256 | 170.08675 | 6.37E-05 | 0.000326 | 106.6278 | 170.7734 | 1.601584 | 0.679499 | 2.130145 |
| VC0192 | 288.84161 | 305.5385 | 326.1744 | 541.01341 | 513.1165 | 499.74926 | 2.52E-09 | 2.15E-08 | 306.8515 | 517.9597 | 1.687982 | 0.755299 | 2.600612 |
| VC0199 | 4987.7504 | 7069.69 | 3972.681 | 2340.6273 | 1926.386 | 1717.1117 | 1.54E-07 | 1.12E-06 | 5343.374 | 1994.708 | 0.373305 | -1.42157 | 3.513847 |
| VC0200 | 22126.733 | 32237.83 | 19235.54 | 13371.347 | 10228.12 | 9545.8798 | 1.55E-05 | 8.65E-05 | 24533.37 | 11048.45 | 0.450344 | -1.1509 | 4.216529 |
| VC0202 | 1638.8622 | 2561.822 | 1412.256 | 843.85772 | 747.684 | 607.72568 | 2.43E-06 | 1.51E-05 | 1870.98 | 733.0891 | 0.391821 | -1.35173 | 3.068613 |
| VC0203 | 2163.1725 | 3651.185 | 1712.197 | 1197.005 | 1083.897 | 948.85472 | 0.000129 | 0.000613 | 2508.851 | 1076.586 | 0.429115 | -1.22056 | 3.215762 |
| VC0212 | 2448.8745 | 3994.328 | 2760.677 | 8933.394 | 7662.539 | 6900.9353 | 3.08E-08 | 2.4E-07 | 3067.96 | 7832.289 | 2.552931 | 1.352155 | 3.690369 |
| VC0213 | 3428.4243 | 4450.285 | 3865.997 | 14064.295 | 11944.37 | 11104.371 | 1.3E-21 | 2.33E-20 | 3914.902 | 12371.01 | 3.15998 | 1.659916 | 3.842563 |
| VC0215 | 2696.9016 | 3447.884 | 2860.366 | 2019.3043 | 2131.632 | 1800.244 | 0.00018 | 0.000825 | 3001.717 | 1983.727 | 0.660864 | -0.59757 | 3.387426 |
| VC0223 | 3275.6313 | 3833.332 | 3176.921 | 2469.9777 | 2139.451 | 1958.8642 | 4.82E-05 | 0.000251 | 3428.628 | 2189.431 | 0.638573 | -0.64708 | 3.437726 |
| VC0224 | 2534.6898 | 2585.325 | 2526.321 | 1542.9661 | 1436.726 | 1407.5156 | 4.11E-18 | 6.48E-17 | 2548.779 | 1462.403 | 0.573766 | -0.80147 | 3.2857 |
| VC0225 | 2862.2529 | 3078.887 | 2723.95 | 2072.687 | 2053.443 | 1935.9312 | 8.75E-07 | 5.73E-06 | 2888.363 | 2020.687 | 0.699596 | -0.51541 | 3.383075 |
| VC0228 | 5360.3143 | 4812.231 | 4654.762 | 2816.9654 | 2558.741 | 2646.8556 | 1.08E-14 | 1.46E-13 | 4942.436 | 2674.187 | 0.541067 | -0.88612 | 3.560566 |

| | | | | | | | | | | | | | |
|---|---|---|---|---|---|---|---|---|---|---|---|---|---|
| VC0235 | 4776.3519 | 4727.62 | 5014.166 | 3868.1945 | 3486.26 | 3381.6685 | 4.2E-05 | 0.000221 | 4839.379 | 3578.708 | 0.739497 | -0.43538 | 3.619258 |
| VC0237 | 2500.1544 | 2793.327 | 2703.837 | 1921.7782 | 1679.113 | 1668.379 | 8.05E-07 | 5.3E-06 | 2665.773 | 1756.423 | 0.65888 | -0.60191 | 3.335226 |
| VC0238 | 3812.5 | 4164.724 | 3698.975 | 3030.4963 | 2601.745 | 2649.7222 | 7.25E-05 | 0.000365 | 3892.066 | 2760.654 | 0.709303 | -0.49553 | 3.515596 |
| VC0242 | 36627.419 | 40937.45 | 34627.83 | 23635.202 | 25058.65 | 25556.967 | 3.18E-07 | 2.22E-06 | 37397.57 | 24750.27 | 0.661815 | -0.5955 | 4.483212 |
| VC0243 | 62729.91 | 68243.19 | 58921.26 | 39928.227 | 40338.77 | 40981.351 | 2.74E-10 | 2.63E-09 | 63298.12 | 40416.12 | 0.638504 | -0.64723 | 4.703973 |
| VC0244 | 51091.476 | 52903.98 | 49123.78 | 35682.247 | 36835.9 | 37375.129 | 3.29E-09 | 2.75E-08 | 51039.75 | 36631.09 | 0.717697 | -0.47855 | 4.635879 |
| VC0245 | 57922.162 | 63933.92 | 52665.36 | 39176.762 | 36805.6 | 38918.332 | 7.82E-07 | 5.16E-06 | 58173.81 | 38300.23 | 0.658376 | -0.60302 | 4.673964 |
| VC0247 | 36090.55 | 37717.55 | 34899.78 | 23981.163 | 21810.87 | 23309.529 | 3.05E-12 | 3.42E-11 | 36235.96 | 23033.85 | 0.635663 | -0.65367 | 4.460753 |
| VC0248 | 21751.029 | 22540.51 | 20446.67 | 13091.087 | 12232.7 | 13873.536 | 1E-11 | 1.07E-10 | 21579.4 | 13065.77 | 0.605474 | -0.72386 | 4.225087 |
| VC0249 | 50385.07 | 54275.38 | 54288.36 | 35157.659 | 31991.1 | 32604.101 | 2.76E-12 | 3.12E-11 | 52982.94 | 33250.95 | 0.627579 | -0.67213 | 4.62297 |
| VC0250 | 41766.916 | 48119.96 | 45135.36 | 30690.962 | 28519.5 | 28670.128 | 8.1E-09 | 6.61E-08 | 45007.41 | 29293.53 | 0.65086 | -0.61958 | 4.560028 |
| VC0269 | 3839.7097 | 4646.535 | 3645.633 | 2259.5266 | 2189.297 | 2007.5969 | 1.75E-09 | 1.51E-08 | 4043.959 | 2152.14 | 0.532186 | -0.91 | 3.469839 |
| VC0270 | 3033.8834 | 3779.276 | 2882.227 | 2094.2454 | 2078.855 | 2139.4619 | 5.73E-05 | 0.000295 | 3231.795 | 2104.187 | 0.651089 | -0.61907 | 3.416264 |
| VC0275 | 936.64218 | 1051.757 | 914.6874 | 665.2309 | 731.0688 | 688.94688 | 0.000103 | 0.0005 | 967.6956 | 695.0822 | 0.718286 | -0.47737 | 2.913887 |
| VC0277 | 717.91792 | 511.1893 | 675.0848 | 312.08363 | 366.5118 | 322.97371 | 1.61E-06 | 1.02E-05 | 634.7307 | 333.8564 | 0.525981 | -0.92692 | 2.663075 |
| VC0283 | 109.8854 | 266.7586 | 87.44621 | 46.196591 | 30.2983 | 20.066414 | 1.57E-05 | 8.78E-05 | 154.6967 | 32.1871 | 0.208066 | -2.26489 | 1.848582 |
| VC0284 | 7926.3999 | 14393.21 | 5340.34 | 1632.2795 | 1260.8 | 1098.8751 | 1.28E-12 | 1.49E-11 | 9219.984 | 1330.652 | 0.144323 | -2.79263 | 3.544397 |
| VC0315 | 2066.892 | 2857.96 | 2410.892 | 5488.155 | 5384.302 | 5393.0876 | 1.71E-12 | 1.95E-11 | 2445.248 | 5421.848 | 2.2173 | 1.148804 | 3.561235 |
| VC0317 | 317.09786 | 253.8319 | 267.5854 | 162.20136 | 161.2652 | 135.68718 | 1.27E-06 | 8.19E-06 | 279.5051 | 153.0512 | 0.547579 | -0.86886 | 2.315613 |
| VC0318 | 3438.8896 | 3896.79 | 3169.051 | 13631.074 | 11010.01 | 11004.039 | 4.39E-27 | 9.86E-26 | 3501.577 | 11881.71 | 3.393245 | 1.762666 | 3.809571 |
| VC0319 | 2671.7849 | 2916.717 | 2726.573 | 9623.2631 | 7953.794 | 7807.7461 | 7.67E-31 | 2.06E-29 | 2771.692 | 8461.601 | 3.052865 | 1.610164 | 3.685099 |
| VC0335 | 847.68734 | 938.9432 | 897.1981 | 3841.5032 | 3432.505 | 3329.1136 | 3.72E-61 | 2.05E-59 | 894.6095 | 3534.374 | 3.950745 | 1.982125 | 3.249973 |
| VC0340 | 5875.2058 | 6106.068 | 5493.371 | 4556.0104 | 4176.279 | 4158.5254 | 2.42E-05 | 0.000131 | 5824.882 | 4296.938 | 0.737687 | -0.43892 | 3.699223 |
| VC0343 | 2014.5656 | 2005.977 | 1769.037 | 5337.2461 | 4427.462 | 4602.8531 | 2.73E-21 | 4.82E-20 | 1929.86 | 4789.187 | 2.481624 | 1.311285 | 3.482894 |
| VC0344 | 5924.3926 | 5985.028 | 5746.091 | 20110.916 | 18058.77 | 18240.37 | 1.56E-68 | 9.72E-67 | 5885.17 | 18803.35 | 3.195039 | 1.675834 | 4.021997 |
| VC0345 | 4778.4449 | 5749.999 | 5220.539 | 17694.321 | 15595.81 | 14603.572 | 1.23E-28 | 3.01E-27 | 5249.661 | 15964.57 | 3.041066 | 1.604577 | 3.961644 |
| VC0346 | 9953.5238 | 10436.49 | 10420.09 | 18932.389 | 16651.36 | 17895.419 | 1.28E-15 | 1.81E-14 | 10270.03 | 17826.39 | 1.735767 | 0.795574 | 4.131318 |
| VC0347 | 6347.1898 | 6973.328 | 6589.072 | 9171.5631 | 8795.305 | 9260.1723 | 4.84E-07 | 3.32E-06 | 6636.53 | 9075.68 | 1.367534 | 0.451577 | 3.88991 |
| VC0348 | 24654.097 | 23884.88 | 28274.86 | 39049.465 | 34408.12 | 35744.017 | 7.41E-05 | 0.000372 | 25604.61 | 36400.53 | 1.42164 | 0.507556 | 4.484713 |
| VC0350 | 24425.954 | 23493.56 | 27267.48 | 36561.008 | 36418.56 | 37300.597 | 1.52E-07 | 1.1E-06 | 25062.33 | 36760.06 | 1.466745 | 0.552618 | 4.482199 |
| VC0351 | 214.53815 | 270.284 | 241.3515 | 379.83863 | 337.1908 | 396.55056 | 0.000158 | 0.000738 | 242.0579 | 371.1933 | 1.53349 | 0.616819 | 2.47676 |
| VC0352 | 751.4068 | 693.3373 | 711.8122 | 353.14727 | 346.9645 | 315.32936 | 3.91E-19 | 6.4E-18 | 718.8521 | 338.4804 | 0.470862 | -1.08662 | 2.693087 |
| VC0353 | 1208.7394 | 936.5929 | 1126.307 | 304.8975 | 399.7422 | 353.5511 | 1.07E-18 | 1.72E-17 | 1090.546 | 352.7303 | 0.323444 | -1.62841 | 2.792543 |
| VC0354 | 42986.12 | 37799.81 | 45310.26 | 149560.95 | 148936.7 | 140742.96 | 1.12E-51 | 4.88E-50 | 42032.06 | 146413.5 | 3.483377 | 1.800487 | 4.894581 |
| VC0364 | 1083.156 | 1397.251 | 970.653 | 569.75795 | 473.0445 | 589.57035 | 1.57E-07 | 1.13E-06 | 1150.353 | 544.1243 | 0.473006 | -1.08007 | 2.898265 |
| VC0365 | 2327.4773 | 2921.418 | 1914.198 | 1446.4666 | 1147.426 | 1155.2521 | 2.42E-05 | 0.000131 | 2387.698 | 1249.715 | 0.523398 | -0.93402 | 3.237395 |
| VC0370 | 834.08248 | 1219.804 | 910.3151 | 13361.081 | 10820.4 | 10037.029 | 7.65E-58 | 3.97E-56 | 988.067 | 11406.17 | 11.54392 | 3.529062 | 3.525963 |
| VC0371 | 9651.0773 | 10227.31 | 9132.008 | 18754.789 | 17080.43 | 16797.5 | 1.99E-15 | 2.79E-14 | 9670.132 | 17544.24 | 1.814271 | 0.85939 | 4.114783 |
| VC0373 | 6182.8849 | 6251.787 | 6308.37 | 10601.604 | 10160.68 | 10075.251 | 3.16E-22 | 5.81E-21 | 6247.68 | 10279.18 | 1.645279 | 0.718333 | 3.903839 |
| VC0389 | 6591.0307 | 6641.936 | 6373.08 | 4271.6447 | 4083.43 | 4282.7461 | 6.79E-16 | 9.71E-15 | 6535.349 | 4212.607 | 0.644588 | -0.63355 | 3.71991 |
| VC0395 | 22291.037 | 19577.96 | 22093.29 | 9157.1908 | 11968.81 | 12916.082 | 5.87E-07 | 3.94E-06 | 21320.76 | 11347.36 | 0.532221 | -0.9099 | 4.191849 |
| VC0399 | 3856.4541 | 3563.048 | 3836.265 | 2182.5323 | 2244.029 | 2227.372 | 9.15E-18 | 1.41E-16 | 3751.923 | 2217.978 | 0.591158 | -0.75839 | 3.460106 |
| VC0400 | 1932.9364 | 1842.632 | 2092.588 | 1327.382 | 1377.107 | 1302.4058 | 2.18E-07 | 1.56E-06 | 1956.052 | 1335.632 | 0.68282 | -0.55042 | 3.208534 |
| VC0401 | 1323.8574 | 1239.781 | 1424.499 | 900.32022 | 975.4099 | 884.8333 | 7.69E-06 | 4.44E-05 | 1329.379 | 920.1878 | 0.692194 | -0.53075 | 3.043763 |
| VC0402 | 8745.831 | 8278.917 | 10065.06 | 6252.9652 | 6218.971 | 5926.2809 | 7.79E-06 | 4.5E-05 | 9029.936 | 6132.739 | 0.679156 | -0.55818 | 3.87167 |
| VC0417 | 1434.7893 | 1749.795 | 1330.057 | 2940.1563 | 2616.405 | 2518.8127 | 3.72E-07 | 2.58E-06 | 1504.88 | 2691.791 | 1.788708 | 0.838918 | 3.303772 |
| VC0418 | 3878.4312 | 4433.833 | 3824.897 | 6725.197 | 5969.743 | 6120.2563 | 3.16E-07 | 2.21E-06 | 4045.721 | 6271.732 | 1.550214 | 0.632467 | 3.702192 |
| VC0419 | 5859.5079 | 5871.039 | 5684.878 | 7270.3168 | 7172.879 | 7023.2449 | 3.23E-05 | 0.000172 | 5805.142 | 7155.48 | 1.232611 | 0.301717 | 3.809226 |
| VC0420 | 3710.9868 | 4390.353 | 4189.548 | 7245.6786 | 6585.483 | 6363.9199 | 2.01E-08 | 1.59E-07 | 4096.962 | 6731.694 | 1.643094 | 0.716415 | 3.720293 |

| | | | | | | | | | | | | | |
|---|---|---|---|---|---|---|---|---|---|---|---|---|---|
| VC0426 | 824.66373 | 990.6497 | 766.0288 | 482.49772 | 363.5797 | 338.26241 | 1.09E-07 | 8.04E-07 | 860.4474 | 394.7799 | 0.458808 | -1.12404 | 2.76554 |
| VC0428 | 1328.0435 | 1727.467 | 1379.901 | 728.87954 | 831.7373 | 827.50069 | 8.03E-08 | 5.97E-07 | 1478.471 | 796.0392 | 0.538421 | -0.89319 | 3.035374 |
| VC0434 | 7232.5521 | 7733.648 | 6590.821 | 5045.6943 | 4474.375 | 4513.0321 | 6.49E-07 | 4.31E-06 | 7185.674 | 4677.701 | 0.650976 | -0.61932 | 3.76325 |
| VC0461 | 3397.0285 | 3136.47 | 3567.805 | 2454.5788 | 2503.031 | 2366.8813 | 7.02E-06 | 4.11E-05 | 3367.101 | 2441.497 | 0.725104 | -0.46374 | 3.457456 |
| VC0470 | 1355.2532 | 1515.941 | 1113.19 | 10521.53 | 9268.349 | 9137.8627 | 2.65E-63 | 1.57E-61 | 1328.128 | 9642.581 | 7.260279 | 2.860025 | 3.553717 |
| VC0473 | 10033.06 | 11512.92 | 10070.31 | 7094.7697 | 8097.466 | 7680.6588 | 0.000145 | 0.000682 | 10538.76 | 7624.298 | 0.723453 | -0.46703 | 3.952495 |
| VC0474 | 618.4978 | 996.5254 | 557.9068 | 317.21659 | 274.6395 | 241.75251 | 1.13E-06 | 7.32E-06 | 724.31 | 277.8695 | 0.383633 | -1.3822 | 2.651883 |
| VC0475 | 14897.32 | 20142.03 | 11611.11 | 2522.3338 | 2193.206 | 1837.5102 | 3.79E-27 | 8.58E-26 | 15550.15 | 2184.35 | 0.140471 | -2.83165 | 3.765528 |
| VC0476 | 5292.29 | 4976.751 | 5010.668 | 4120.7359 | 3721.805 | 3579.466 | 0.000104 | 0.000506 | 5093.236 | 3807.336 | 0.747528 | -0.4198 | 3.643807 |
| VC0479 | 170.584 | 222.103 | 172.269 | 326.45591 | 306.8925 | 300.99621 | 2.44E-05 | 0.000132 | 188.3187 | 311.4482 | 1.653836 | 0.725816 | 2.38414 |
| VC0480 | 12940.313 | 18134.88 | 10582.74 | 45434.86 | 45123.95 | 47417.892 | 2.21E-13 | 2.71E-12 | 13885.98 | 45992.23 | 3.312135 | 1.727761 | 4.402631 |
| VC0481 | 190.46802 | 212.7018 | 215.9921 | 470.17863 | 419.2894 | 422.35024 | 3.32E-14 | 4.31E-13 | 206.3873 | 437.2728 | 2.1187 | 1.083179 | 2.477718 |
| VC0483 | 2705.2738 | 2788.626 | 2760.677 | 59632.612 | 54473.42 | 56332.157 | 0 | 0 | 2751.526 | 56812.73 | 20.64772 | 4.367911 | 4.09701 |
| VC0500 | 58.605544 | 31.72899 | 59.46342 | 127.29727 | 170.0615 | 119.44294 | 2.5E-06 | 1.55E-05 | 49.93265 | 138.9339 | 2.782425 | 1.476343 | 1.920596 |
| VC0501 | 150.69997 | 145.7183 | 166.1478 | 101.6325 | 92.84964 | 84.08783 | 5.57E-05 | 0.000287 | 154.1887 | 92.85666 | 0.602227 | -0.73162 | 2.077933 |
| VC0503 | 782.80263 | 673.3597 | 711.8122 | 370.59932 | 440.7915 | 374.57306 | 3.01E-09 | 2.54E-08 | 722.6582 | 395.3213 | 0.547038 | -0.87029 | 2.727942 |
| VC0512 | 2299.2211 | 2117.617 | 2417.013 | 1612.7743 | 1630.244 | 1473.4481 | 3.27E-06 | 1.97E-05 | 2277.95 | 1572.156 | 0.690162 | -0.53499 | 3.27702 |
| VC0513 | 244.88745 | 230.329 | 199.3774 | 150.90886 | 115.329 | 124.22066 | 4.38E-05 | 0.00023 | 224.8646 | 130.1529 | 0.578805 | -0.78885 | 2.233187 |
| VC0514 | 1279.9032 | 1106.989 | 1252.23 | 862.33636 | 917.7454 | 864.76689 | 9.57E-05 | 0.00047 | 1213.041 | 881.6162 | 0.726782 | -0.46041 | 3.014577 |
| VC0515 | 755.59291 | 674.5349 | 750.2885 | 460.93932 | 438.8367 | 424.26132 | 2.38E-09 | 2.05E-08 | 726.8054 | 441.3458 | 0.607241 | -0.71966 | 2.753099 |
| VC0527 | 824.66373 | 1062.334 | 775.6479 | 2361.1591 | 2230.346 | 2112.7067 | 1.61E-14 | 2.14E-13 | 887.5484 | 2234.737 | 2.517876 | 1.332207 | 3.148709 |
| VC0528 | 1109.3192 | 1403.127 | 1170.905 | 3834.317 | 3367.999 | 3331.9803 | 5.43E-23 | 1.04E-21 | 1227.784 | 3511.432 | 2.859976 | 1.516003 | 3.317303 |
| VC0529 | 1814.6788 | 2260.985 | 1724.439 | 4869.1206 | 4263.265 | 4229.2356 | 3.61E-13 | 4.36E-12 | 1933.368 | 4453.874 | 2.303687 | 1.203945 | 3.467526 |
| VC0530 | 4255.1811 | 5490.291 | 4109.098 | 10777.151 | 9813.719 | 9818.2097 | 1.22E-11 | 1.29E-10 | 4618.19 | 10136.36 | 2.194877 | 1.13414 | 3.835177 |
| VC0531 | 2356.7801 | 3080.063 | 2315.576 | 5820.7704 | 5348.139 | 5135.0909 | 3.5E-10 | 3.31E-09 | 2584.139 | 5434.667 | 2.103086 | 1.072508 | 3.573744 |
| VC0532 | 1784.3295 | 2047.108 | 1649.236 | 3657.7434 | 3270.262 | 3252.6702 | 2.56E-10 | 2.46E-09 | 1826.891 | 3393.559 | 1.85756 | 0.893408 | 3.396184 |
| VC0554 | 9444.9114 | 10988.81 | 10155.13 | 15649.352 | 15807.9 | 15444.45 | 3.14E-09 | 2.63E-08 | 10196.28 | 15633.9 | 1.533294 | 0.616634 | 4.101255 |
| VC0567 | 9428.1669 | 8194.306 | 10714.78 | 16745.751 | 18402.8 | 16783.166 | 6.74E-09 | 5.54E-08 | 9445.753 | 17310.57 | 1.83263 | 0.873916 | 4.106774 |
| VC0578 | 3135.3966 | 2766.298 | 3232.886 | 6538.3575 | 5727.357 | 5921.5032 | 4E-15 | 5.5E-14 | 3044.86 | 6062.406 | 1.991029 | 0.993514 | 3.633106 |
| VC0579 | 2681.2037 | 2356.172 | 2739.69 | 4800.3391 | 4334.612 | 4049.5934 | 1.38E-08 | 1.11E-07 | 2592.355 | 4394.848 | 1.695311 | 0.76155 | 3.528319 |
| VC0580 | 1251.647 | 1198.651 | 1273.217 | 3690.5943 | 3329.881 | 3283.2475 | 2.65E-43 | 8.97E-42 | 1241.172 | 3434.574 | 2.767204 | 1.468429 | 3.314852 |
| VC0581 | 8474.7803 | 8473.991 | 9323.515 | 34753.182 | 32901.03 | 30237.219 | 3.19E-63 | 1.87E-61 | 8757.429 | 32630.48 | 3.726034 | 1.897641 | 4.228 |
| VC0582 | 1537.349 | 1940.169 | 1345.797 | 872.60227 | 868.8772 | 799.78993 | 7.13E-07 | 4.72E-06 | 1607.772 | 847.0898 | 0.526872 | -0.92448 | 3.067077 |
| VC0583 | 1838.749 | 1816.779 | 1918.57 | 4490.3086 | 4169.438 | 4298.9903 | 2.62E-42 | 8.55E-41 | 1858.032 | 4319.579 | 2.324813 | 1.217115 | 3.452247 |
| VC0587 | 1291.415 | 1196.301 | 1135.926 | 2450.4725 | 2420.932 | 2409.8808 | 4.68E-23 | 8.99E-22 | 1207.881 | 2427.095 | 2.009383 | 1.006753 | 3.233555 |
| VC0601 | 1865.9587 | 2288.013 | 1894.085 | 7970.4518 | 7220.77 | 6380.1641 | 8.61E-30 | 2.18E-28 | 2016.019 | 7190.462 | 3.566664 | 1.834575 | 3.580626 |
| VC0602 | 3025.5112 | 3453.76 | 3078.981 | 30748.451 | 27918.42 | 26284.136 | 9E-138 | 1.7E-135 | 3186.084 | 28317 | 8.887714 | 3.151812 | 3.977652 |
| VC0603 | 3525.7514 | 3157.622 | 4007.66 | 18744.523 | 16114.79 | 15701.491 | 3.42E-46 | 1.25E-44 | 3563.678 | 16853.6 | 4.729272 | 2.241618 | 3.889296 |
| VC0608 | 13523.229 | 14944.36 | 12822.24 | 10601.604 | 10012.12 | 10217.627 | 0.00011 | 0.000532 | 13763.27 | 10277.12 | 0.746706 | -0.42139 | 4.075297 |
| VC0610 | 3431.5639 | 3419.68 | 2897.967 | 2324.2018 | 2377.928 | 2248.3939 | 7.04E-05 | 0.000356 | 3249.737 | 2316.841 | 0.712932 | -0.48816 | 3.438372 |
| VC0626 | 13911.491 | 15875.07 | 12266.95 | 6684.1334 | 6966.655 | 7059.5555 | 8.3E-13 | 9.69E-12 | 14017.84 | 6903.448 | 0.492476 | -1.02187 | 3.992874 |
| VC0629 | 797.45401 | 829.6544 | 862.2197 | 1282.212 | 1160.132 | 1210.6736 | 2.23E-07 | 1.59E-06 | 829.776 | 1217.673 | 1.467471 | 0.553332 | 3.002246 |
| VC0638 | 2491.7822 | 2515.992 | 2546.434 | 2100.405 | 1906.838 | 1925.4202 | 0.000211 | 0.000954 | 2518.069 | 1977.555 | 0.785346 | -0.3486 | 3.348598 |
| VC0648 | 2511.6662 | 2345.595 | 2386.407 | 13050.024 | 11712.74 | 11327.968 | 1.06E-97 | 1.15E-95 | 2414.556 | 12030.24 | 4.982383 | 2.316836 | 3.731556 |
| VC0649 | 198.84024 | 245.6059 | 193.2561 | 341.85477 | 322.5303 | 350.68447 | 3.3E-05 | 0.000175 | 212.5674 | 338.3565 | 1.591761 | 0.670624 | 2.428436 |
| VC0666 | 90.001372 | 97.53728 | 89.19514 | 215.58409 | 208.1787 | 233.15262 | 7.01E-14 | 8.92E-13 | 92.24459 | 218.9718 | 2.373817 | 1.247209 | 2.152665 |
| VC0673 | 878.03664 | 985.9491 | 1010.004 | 1338.6745 | 1313.578 | 1241.251 | 0.000169 | 0.000781 | 957.9965 | 1297.835 | 1.354738 | 0.438014 | 3.047292 |
| VC0676 | 186.28191 | 168.0461 | 130.2949 | 421.92886 | 347.9418 | 370.75089 | 8.08E-10 | 7.34E-09 | 161.541 | 380.2072 | 2.353627 | 1.234886 | 2.394152 |
| VC0680 | 2010.3795 | 3041.283 | 1987.652 | 29259.894 | 23949.34 | 24424.648 | 3E-50 | 1.24E-48 | 2346.438 | 25877.96 | 11.02861 | 3.46318 | 3.89167 |

| | | | | | | | | | | | | | |
|---|---|---|---|---|---|---|---|---|---|---|---|---|---|
| VC0681 | 3557.1472 | 5048.435 | 3705.97 | 11324.324 | 9446.23 | 9809.6098 | 3.73E-11 | 3.83E-10 | 4103.851 | 10193.39 | 2.483859 | 1.312583 | 3.810755 |
| VC0682 | 40447.244 | 51826.37 | 37932.42 | 78825.756 | 75680.28 | 77593.001 | 5.39E-07 | 3.66E-06 | 43402.01 | 77366.35 | 1.782552 | 0.833944 | 4.763031 |
| VC0683 | 3888.8965 | 5507.918 | 3363.181 | 7722.0168 | 7404.515 | 8133.5865 | 0.000131 | 0.000622 | 4253.332 | 7753.373 | 1.822894 | 0.866231 | 3.75911 |
| VC0687 | 25646.205 | 12035.86 | 25556.16 | 828.45886 | 2443.412 | 5162.8017 | 2.85E-06 | 1.75E-05 | 21079.41 | 2811.557 | 0.133379 | -2.90639 | 3.886403 |
| VC0695 | 6872.5466 | 4727.62 | 6191.192 | 2987.3795 | 3105.088 | 3159.9824 | 2.68E-07 | 1.89E-06 | 5930.453 | 3084.15 | 0.520053 | -0.94327 | 3.631112 |
| VC0700 | 1330.1366 | 1182.199 | 1432.369 | 5592.8672 | 5146.802 | 5020.4257 | 3.24E-53 | 1.46E-51 | 1314.901 | 5253.365 | 3.995254 | 1.998287 | 3.419665 |
| VC0702 | 601.75336 | 584.0485 | 586.7641 | 950.62318 | 881.5829 | 862.8558 | 1.67E-08 | 1.34E-07 | 590.8553 | 898.354 | 1.52043 | 0.604479 | 2.862464 |
| VC0703 | 845.59428 | 842.581 | 835.9858 | 3219.3891 | 2775.716 | 2640.1668 | 2.48E-42 | 8.16E-41 | 841.387 | 2878.424 | 3.421046 | 1.774438 | 3.192075 |
| VC0706 | 39005.129 | 15506.08 | 61088.18 | 7361.6834 | 12289.38 | 11856.384 | 0.000131 | 0.000621 | 38533.13 | 10502.48 | 0.272557 | -1.87537 | 4.303563 |
| VC0708 | 3083.0702 | 3279.838 | 3162.055 | 4200.81 | 3909.459 | 3963.5945 | 0.000115 | 0.00055 | 3174.988 | 4024.621 | 1.267602 | 0.342102 | 3.553234 |
| VC0718 | 4716.6998 | 5648.936 | 4855.888 | 14301.438 | 13727.09 | 13205.611 | 5.55E-31 | 1.51E-29 | 5073.841 | 13744.71 | 2.708936 | 1.437726 | 3.921736 |
| VC0743 | 19944.723 | 23452.43 | 23133.9 | 15079.594 | 15114.94 | 16591.102 | 3.91E-05 | 0.000206 | 22177.01 | 15595.21 | 0.703215 | -0.50796 | 4.269447 |
| VC0744 | 14223.356 | 16340.43 | 15343.31 | 11674.392 | 11275.86 | 11618.454 | 5.02E-05 | 0.00026 | 15302.37 | 11522.9 | 0.753014 | -0.40925 | 4.12316 |
| VC0746 | 2313.8725 | 3964.949 | 2413.515 | 1087.1598 | 1062.395 | 931.65493 | 1.33E-08 | 1.07E-07 | 2897.446 | 1027.07 | 0.354474 | -1.49625 | 3.236808 |
| VC0755 | 14236.961 | 12925.45 | 15463.99 | 34112.589 | 37824.99 | 36711.027 | 5.43E-28 | 1.29E-26 | 14208.8 | 36216.2 | 2.548857 | 1.349851 | 4.35573 |
| VC0756 | 6621.38 | 5519.67 | 8583.72 | 13359.027 | 13293.14 | 13535.274 | 2.34E-06 | 1.46E-05 | 6908.257 | 13395.81 | 1.939102 | 0.955389 | 3.983169 |
| VC0765 | 281.51592 | 230.329 | 296.4427 | 428.08841 | 422.2215 | 454.83872 | 1.18E-05 | 6.65E-05 | 269.4292 | 435.0496 | 1.614708 | 0.691274 | 2.534492 |
| VC0771 | 2051.1941 | 2371.448 | 1740.18 | 732.9859 | 652.8796 | 565.68177 | 4.42E-18 | 6.94E-17 | 2054.274 | 650.5158 | 0.316665 | -1.65897 | 3.062958 |
| VC0772 | 189.42149 | 192.7243 | 156.5287 | 45.17 | 63.5287 | 72.621308 | 1.14E-10 | 1.13E-09 | 179.5582 | 60.44 | 0.336604 | -1.57088 | 2.017765 |
| VC0773 | 198.84024 | 341.968 | 204.6241 | 51.329545 | 46.9135 | 43.955002 | 3.02E-15 | 4.18E-14 | 248.4775 | 47.39935 | 0.190759 | -2.39018 | 2.03553 |
| VC0774 | 316.05133 | 386.6237 | 288.5725 | 80.07409 | 76.23444 | 81.221199 | 5.72E-25 | 1.21E-23 | 330.4158 | 79.17658 | 0.239627 | -2.06114 | 2.208829 |
| VC0775 | 63.838182 | 129.2663 | 69.08251 | 18.478636 | 22.47939 | 21.021958 | 8.97E-08 | 6.64E-07 | 87.39565 | 20.65999 | 0.236396 | -2.08072 | 1.62831 |
| VC0776 | 379.88951 | 364.2958 | 374.2698 | 149.88227 | 129.9895 | 146.19816 | 2.48E-22 | 4.6E-21 | 372.8184 | 142.0233 | 0.380945 | -1.39235 | 2.361928 |
| VC0791 | 53.372906 | 48.18106 | 66.45912 | 97.526136 | 108.4875 | 102.24316 | 9.54E-05 | 0.000468 | 56.00436 | 102.7523 | 1.834719 | 0.875559 | 1.880007 |
| VC0792 | 58.605544 | 59.93254 | 63.83574 | 137.56318 | 121.1932 | 124.22066 | 4.9E-08 | 3.75E-07 | 60.79127 | 127.659 | 2.099956 | 1.070359 | 1.944946 |
| VC0793m | 68.024293 | 43.48047 | 60.33789 | 187.86614 | 178.8577 | 164.35349 | 3.45E-13 | 4.2E-12 | 57.28088 | 177.0258 | 3.090486 | 1.627834 | 2.003023 |
| VC0794 | 0 | 7.050887 | 3.497849 | 25.664773 | 19.54729 | 28.666306 | 3.87E-06 | 2.3E-05 | 3.516245 | 24.62612 | 7.003528 | 2.808082 | 0.968738 |
| VC0795 | 9.4187482 | 11.75148 | 11.36801 | 42.090227 | 31.27567 | 41.088371 | 1.14E-06 | 7.42E-06 | 10.84608 | 38.15142 | 3.517532 | 1.814563 | 1.308392 |
| VC0812 | 2764.9259 | 2410.228 | 2818.391 | 1453.6527 | 1821.808 | 1834.6436 | 5.2E-05 | 0.000269 | 2664.515 | 1703.368 | 0.639279 | -0.64548 | 3.328463 |
| VC0813 | 795.36096 | 938.9432 | 812.3753 | 428.08841 | 509.207 | 556.12633 | 3.82E-06 | 2.27E-05 | 848.8931 | 497.8072 | 0.586419 | -0.77 | 2.812957 |
| VC0815 | 1772.8177 | 1483.037 | 1810.137 | 1143.6223 | 1187.498 | 1199.2071 | 8.24E-05 | 0.00041 | 1688.664 | 1176.776 | 0.696868 | -0.52104 | 3.149118 |
| VC0819 | 69.07082 | 61.10769 | 61.21235 | 129.35045 | 103.6007 | 123.26511 | 1.74E-05 | 9.62E-05 | 63.79695 | 118.7387 | 1.861198 | 0.896231 | 1.939696 |
| VC0821 | 3140.6293 | 2520.692 | 3479.485 | 1859.1561 | 1666.407 | 1618.6907 | 2.2E-06 | 1.37E-05 | 3046.935 | 1714.751 | 0.562779 | -0.82936 | 3.359032 |
| VC0824 | 5969.3933 | 6129.571 | 5742.593 | 4562.17 | 4545.723 | 4761.4734 | 1.11E-05 | 6.25E-05 | 5947.186 | 4623.122 | 0.777363 | -0.36334 | 3.719623 |
| VC0826 | 9706.5433 | 7874.666 | 10112.28 | 5319.7941 | 5440.012 | 4802.5617 | 6.21E-08 | 4.69E-07 | 9231.163 | 5187.456 | 0.56195 | -0.83149 | 3.840105 |
| VC0827 | 3075.7446 | 2591.201 | 3612.403 | 1803.7202 | 1872.631 | 1737.1781 | 4.55E-06 | 2.69E-05 | 3093.116 | 1804.51 | 0.583395 | -0.77745 | 3.373378 |
| VC0828 | 560.93878 | 651.0319 | 704.8165 | 1128.2234 | 1092.694 | 1076.8976 | 1.43E-08 | 1.15E-07 | 638.9291 | 1099.272 | 1.720491 | 0.78282 | 2.923279 |
| VC0845 | 2940.7425 | 2994.277 | 2739.69 | 2244.1277 | 1904.884 | 1909.176 | 4.21E-05 | 0.000221 | 2891.57 | 2019.396 | 0.698374 | -0.51793 | 3.383178 |
| VC0854 | 19333.55 | 17989.16 | 24590.75 | 11330.484 | 11410.73 | 9924.275 | 1.73E-07 | 1.24E-06 | 20637.82 | 10888.5 | 0.527599 | -0.92249 | 4.175816 |
| VC0858 | 14.651386 | 23.50296 | 22.73602 | 177.60023 | 176.903 | 147.1537 | 4.52E-29 | 1.13E-27 | 20.29679 | 167.219 | 8.238692 | 3.042415 | 1.765356 |
| VC0859 | 15.697914 | 27.0284 | 23.61048 | 96.499545 | 113.3743 | 110.84305 | 1.84E-16 | 2.66E-15 | 22.11226 | 106.9056 | 4.834676 | 2.273419 | 1.686817 |
| VC0860 | 75.349986 | 54.0568 | 71.70589 | 160.14818 | 154.4236 | 150.97588 | 2.48E-09 | 2.13E-08 | 67.03756 | 155.1826 | 2.31486 | 1.210925 | 2.008581 |
| VC0871 | 387.2152 | 475.9349 | 329.6722 | 254.59454 | 234.5675 | 221.6861 | 0.000173 | 0.000798 | 397.6074 | 236.9494 | 0.595938 | -0.74677 | 2.487055 |
| VC0872 | 594.42766 | 694.5124 | 483.5776 | 329.53568 | 327.4172 | 309.5961 | 3.14E-06 | 1.9E-05 | 590.8392 | 322.183 | 0.545297 | -0.87489 | 2.639786 |
| VC0878 | 372.56382 | 341.968 | 381.2655 | 191.9725 | 174.9483 | 168.17566 | 7.31E-13 | 8.58E-12 | 365.2658 | 178.3655 | 0.488317 | -1.03411 | 2.40696 |
| VC0879 | 115.11803 | 95.18698 | 105.8099 | 58.515681 | 56.68715 | 66.888047 | 0.000141 | 0.000663 | 105.3716 | 60.69696 | 0.576027 | -0.79579 | 1.902945 |
| VC0913 | 1313.3921 | 1079.961 | 1172.654 | 1943.3366 | 1859.925 | 1733.356 | 1.9E-06 | 1.19E-05 | 1188.669 | 1845.539 | 1.55261 | 0.634695 | 3.170592 |
| VC0914 | 2665.5057 | 2276.261 | 2738.815 | 4681.2545 | 4633.686 | 4529.2763 | 2.82E-12 | 3.18E-11 | 2560.194 | 4614.739 | 1.802496 | 0.849996 | 3.53621 |
| VC0916 | 18.837496 | 9.401183 | 13.11693 | 57.489091 | 39.09459 | 53.510437 | 6E-07 | 4.01E-06 | 13.7852 | 50.03137 | 3.629353 | 1.859712 | 1.419328 |

| | | | | | | | | | | | | | |
|---|---|---|---|---|---|---|---|---|---|---|---|---|---|
| VC0917 | 32.442355 | 45.83077 | 33.22956 | 93.419772 | 115.329 | 113.70968 | 8.18E-10 | 7.41E-09 | 37.16756 | 107.4862 | 2.891935 | 1.532035 | 1.800758 |
| VC0918 | 165.35136 | 165.6959 | 175.7669 | 356.22704 | 418.3121 | 383.17295 | 8.47E-16 | 1.21E-14 | 168.938 | 385.904 | 2.284293 | 1.191748 | 2.407103 |
| VC0919 | 50.233324 | 45.83077 | 46.34649 | 79.047499 | 101.6459 | 101.28761 | 2.05E-05 | 0.000112 | 47.47019 | 93.99368 | 1.980057 | 0.985542 | 1.82476 |
| VC0930 | 1297.6942 | 1304.414 | 1584.525 | 2144.5484 | 2192.229 | 2210.1722 | 1.56E-06 | 9.86E-06 | 1395.545 | 2182.317 | 1.563774 | 0.645032 | 3.241831 |
| VC0947 | 18031.67 | 22487.63 | 16482.74 | 51222.78 | 45919.52 | 46823.544 | 2.38E-15 | 3.33E-14 | 19000.68 | 47988.62 | 2.525626 | 1.336641 | 4.479954 |
| VC0948 | 3289.2362 | 3890.915 | 3358.809 | 22118.928 | 17210.41 | 16864.388 | 1.08E-43 | 3.7E-42 | 3512.987 | 18731.24 | 5.332 | 2.414677 | 3.909122 |
| VC0949 | 1631.5365 | 2110.566 | 1445.486 | 3203.9902 | 2804.059 | 2861.8529 | 6.15E-05 | 0.000315 | 1729.196 | 2956.634 | 1.709832 | 0.773854 | 3.354321 |
| VC0968 | 2300.2676 | 2605.303 | 2560.425 | 9201.3343 | 8912.588 | 8300.8066 | 3.55E-60 | 1.9E-58 | 2488.665 | 8804.91 | 3.538005 | 1.822936 | 3.670346 |
| VC0969 | 790.12832 | 730.942 | 790.5138 | 7173.8172 | 6069.435 | 5843.1486 | 3.3E-106 | 3.8E-104 | 770.528 | 6362.133 | 8.256849 | 3.045591 | 3.345196 |
| VC0970 | 6789.8709 | 7109.645 | 6486.76 | 15932.691 | 14111.19 | 14596.883 | 7.68E-27 | 1.71E-25 | 6795.425 | 14880.25 | 2.189746 | 1.130764 | 4.002414 |
| VC0971 | 3005.6272 | 4308.092 | 2893.595 | 6677.9738 | 6148.601 | 6369.6531 | 8.91E-06 | 5.12E-05 | 3402.438 | 6398.743 | 1.880635 | 0.91122 | 3.668942 |
| VC0975 | 1537.349 | 1504.189 | 1637.868 | 2887.8002 | 2862.701 | 2686.9884 | 3.43E-18 | 5.45E-17 | 1559.802 | 2812.497 | 1.803111 | 0.850488 | 3.321081 |
| VC0976 | 3194.0022 | 2780.4 | 3295.848 | 4914.2906 | 4836.978 | 4485.3213 | 4.84E-07 | 3.32E-06 | 3090.083 | 4745.53 | 1.535729 | 0.618923 | 3.583127 |
| VC0977 | 3527.8445 | 1685.162 | 6260.274 | 1124.117 | 1169.906 | 1100.7861 | 8.99E-05 | 0.000444 | 3824.427 | 1131.603 | 0.295888 | -1.75688 | 3.31813 |
| VC0980 | 1175.2505 | 746.2189 | 1170.03 | 520.48159 | 562.9621 | 521.72676 | 3.36E-05 | 0.000178 | 1030.5 | 535.0568 | 0.519221 | -0.94558 | 2.870724 |
| VC0982 | 952.3401 | 1048.232 | 1037.112 | 2088.0859 | 1859.925 | 1826.0437 | 8.32E-15 | 1.13E-13 | 1012.561 | 1924.685 | 1.900808 | 0.926613 | 3.14489 |
| VC0983 | 1569.7914 | 1566.472 | 1519.815 | 2188.6918 | 2320.264 | 2407.0141 | 1.27E-09 | 1.12E-08 | 1552.026 | 2305.323 | 1.485364 | 0.570816 | 3.276815 |
| VC0984 | 4537.7436 | 4613.631 | 5003.672 | 6445.9643 | 6302.047 | 6104.012 | 1.02E-05 | 5.82E-05 | 4718.349 | 6284.008 | 1.331824 | 0.413403 | 3.736013 |
| VC0985 | 113905.11 | 97520.82 | 152404.8 | 42062.509 | 49558.25 | 46651.546 | 8.55E-11 | 8.59E-10 | 121276.9 | 46090.77 | 0.380046 | -1.39575 | 4.873696 |
| VC1002 | 1798.9809 | 1956.621 | 1697.331 | 3438.0529 | 3319.13 | 3018.562 | 4.23E-12 | 4.67E-11 | 1817.644 | 3258.582 | 1.79275 | 0.842174 | 3.386269 |
| VC1003 | 1258.9727 | 1453.658 | 1153.416 | 2365.2654 | 2040.737 | 2288.5267 | 1.26E-07 | 9.22E-07 | 1288.682 | 2231.51 | 1.731622 | 0.792124 | 3.229372 |
| VC1004 | 1063.272 | 1357.296 | 1150.792 | 2522.3338 | 2216.663 | 2273.238 | 1.24E-10 | 1.23E-09 | 1190.453 | 2337.412 | 1.963463 | 0.973401 | 3.222224 |
| VC1008 | 1512.2323 | 1427.805 | 1602.015 | 402.42363 | 437.8594 | 375.5286 | 2.45E-50 | 1.02E-48 | 1514.017 | 405.2705 | 0.267679 | -1.90142 | 2.893938 |
| VC1009 | 1052.8067 | 1132.843 | 869.2154 | 599.52909 | 521.9127 | 536.05992 | 7.55E-08 | 5.65E-07 | 1018.288 | 552.5006 | 0.542578 | -0.8821 | 2.875102 |
| VC1033 | 13103.572 | 13473.07 | 14466.23 | 6799.1115 | 7725.09 | 6194.7887 | 3.95E-13 | 4.71E-12 | 13680.96 | 6906.33 | 0.504813 | -0.98618 | 3.987682 |
| VC1039 | 11637.387 | 10196.76 | 12703.31 | 102092.41 | 101215.9 | 96520.407 | 2.5E-127 | 4E-125 | 11512.49 | 99942.9 | 8.681262 | 3.117905 | 4.530461 |
| VC1042 | 298.26036 | 432.4544 | 233.4814 | 1053.2823 | 1005.708 | 983.25428 | 1.32E-10 | 1.29E-09 | 321.3987 | 1014.082 | 3.155214 | 1.657738 | 2.756559 |
| VC1050 | 860.24567 | 582.8734 | 837.7347 | 2477.1638 | 2862.701 | 2813.1201 | 1.99E-19 | 3.3E-18 | 760.2846 | 2717.662 | 3.574532 | 1.837754 | 3.157586 |
| VC1051 | 1308.1595 | 1331.443 | 1169.156 | 1881.7411 | 1936.159 | 2142.3286 | 1.57E-07 | 1.13E-06 | 1269.586 | 1986.743 | 1.564875 | 0.646047 | 3.200902 |
| VC1052 | 1240.1352 | 1131.667 | 1094.827 | 1497.7961 | 1801.283 | 1882.4207 | 9.73E-05 | 0.000476 | 1155.543 | 1727.167 | 1.49468 | 0.579836 | 3.15006 |
| VC1055 | 7505.6958 | 7707.795 | 8283.78 | 5846.4352 | 5602.254 | 5405.5097 | 7.33E-07 | 4.85E-06 | 7832.424 | 5618.066 | 0.717283 | -0.47939 | 3.821742 |
| VC1056 | 12716.357 | 13742.18 | 14412.01 | 9967.1711 | 9563.513 | 9436.9478 | 5.58E-07 | 3.78E-06 | 13623.52 | 9655.877 | 0.708765 | -0.49662 | 4.05954 |
| VC1059 | 4811.9338 | 4275.188 | 4539.333 | 3274.825 | 3385.591 | 3452.3787 | 1.19E-05 | 6.71E-05 | 4542.152 | 3370.932 | 0.742144 | -0.43023 | 3.592506 |
| VC1063 | 3500.6347 | 3271.612 | 3387.666 | 1556.3118 | 1485.594 | 1368.3383 | 3.15E-30 | 8.13E-29 | 3386.638 | 1470.081 | 0.434083 | -1.20396 | 3.348555 |
| VC1064 | 2232.2433 | 3103.566 | 2187.904 | 10889.05 | 8609.605 | 8182.3192 | 1.94E-17 | 2.95E-16 | 2507.904 | 9226.991 | 3.679164 | 1.879378 | 3.682186 |
| VC1067 | 1331.1831 | 2017.729 | 1504.949 | 6355.6243 | 4791.042 | 4575.1424 | 1.47E-12 | 1.69E-11 | 1617.954 | 5240.603 | 3.239031 | 1.695562 | 3.464174 |
| VC1073 | 551.52003 | 723.8911 | 494.9456 | 1116.9309 | 1570.625 | 1675.0678 | 1.37E-07 | 1E-06 | 590.1189 | 1454.208 | 2.464263 | 1.301156 | 2.966783 |
| VC1077 | 7365.4611 | 5131.871 | 11276.19 | 2827.2313 | 3251.692 | 3124.6273 | 1.11E-05 | 6.25E-05 | 7924.507 | 3067.85 | 0.387135 | -1.36909 | 3.692903 |
| VC1107 | 1927.7038 | 2168.148 | 1820.63 | 9278.3286 | 7964.545 | 7845.0123 | 6.78E-49 | 2.61E-47 | 1972.161 | 8362.629 | 4.240339 | 2.084179 | 3.608643 |
| VC1108 | 1666.0719 | 1922.542 | 1546.924 | 8837.9211 | 7975.296 | 7669.1923 | 2.77E-55 | 1.35E-53 | 1711.846 | 8160.803 | 4.767254 | 2.253158 | 3.572599 |
| VC1112 | 323.37702 | 373.697 | 328.7978 | 153.98864 | 181.7898 | 172.95338 | 1.28E-10 | 1.26E-09 | 341.9573 | 169.5773 | 0.495902 | -1.01187 | 2.38167 |
| VC1134 | 775.47694 | 790.8745 | 791.3882 | 1086.1332 | 1068.26 | 1072.1198 | 5.45E-07 | 3.7E-06 | 785.9132 | 1075.504 | 1.368477 | 0.452571 | 2.963493 |
| VC1138 | 329.65619 | 379.5728 | 385.6378 | 615.95454 | 582.5093 | 540.83763 | 1.69E-06 | 1.07E-05 | 364.9556 | 579.7672 | 1.588597 | 0.667753 | 2.662747 |
| VC1139 | 501.28671 | 548.7941 | 540.4176 | 809.98022 | 801.439 | 772.07917 | 6.85E-08 | 5.16E-07 | 530.1661 | 794.4995 | 1.498586 | 0.583602 | 2.812253 |
| VC1142 | 183.14233 | 157.4698 | 195.8795 | 340.82818 | 338.1682 | 355.46219 | 5.36E-10 | 4.97E-09 | 178.8306 | 344.8195 | 1.928191 | 0.947248 | 2.395017 |
| VC1143 | 5631.3649 | 4952.073 | 5812.55 | 3400.0691 | 3230.19 | 3304.2695 | 2.07E-10 | 2.02E-09 | 5465.329 | 3311.51 | 0.605912 | -0.72282 | 3.628821 |
| VC1147 | 399.77353 | 320.8154 | 402.2526 | 556.41227 | 723.2499 | 694.68014 | 1.26E-05 | 7.05E-05 | 374.2805 | 658.1141 | 1.758345 | 0.814218 | 2.695749 |
| VC1154 | 6723.9397 | 7645.512 | 6634.544 | 18904.671 | 17332.59 | 17152.962 | 1.74E-30 | 4.6E-29 | 7001.332 | 17796.74 | 2.541908 | 1.345912 | 4.047761 |
| VC1158 | 5481.7115 | 6068.464 | 5482.003 | 53989.442 | 48091.23 | 48277.881 | 6E-171 | 1.9E-168 | 5677.393 | 50119.52 | 8.827911 | 3.142072 | 4.227078 |

| | | | | | | | | | | | | | |
|---|---|---|---|---|---|---|---|---|---|---|---|---|---|
| VC1160 | 2554.5738 | 3080.063 | 2355.801 | 14448.24 | 13125.03 | 13003.036 | 7.01E-52 | 3.08E-50 | 2663.479 | 13525.44 | 5.078108 | 2.344291 | 3.7783 |
| VC1161 | 2779.5772 | 3720.518 | 2626.01 | 15768.436 | 14711.29 | 14726.837 | 1.97E-36 | 5.77E-35 | 3042.035 | 15068.86 | 4.953544 | 2.308461 | 3.830622 |
| VC1162 | 1154.3199 | 1559.421 | 1183.147 | 5213.0286 | 5051.998 | 4315.2345 | 1.2E-24 | 2.49E-23 | 1298.963 | 4860.087 | 3.741514 | 1.903622 | 3.40012 |
| VC1165 | 327.56313 | 304.3633 | 361.1529 | 218.66386 | 207.2013 | 213.08621 | 1.08E-05 | 6.13E-05 | 331.0264 | 212.9838 | 0.643404 | -0.6362 | 2.424105 |
| VC1185 | 446.86728 | 417.1775 | 508.937 | 283.33909 | 221.8618 | 212.13066 | 5.98E-07 | 4E-06 | 457.6606 | 239.1105 | 0.522463 | -0.9366 | 2.519571 |
| VC1186 | 801.64012 | 1020.028 | 752.0374 | 2871.3748 | 2274.328 | 2160.4839 | 1.64E-13 | 2.03E-12 | 857.902 | 2435.395 | 2.838781 | 1.505271 | 3.160004 |
| VC1189 | 256.39926 | 350.1941 | 172.269 | 628.27363 | 610.8529 | 473.94959 | 0.000161 | 0.000751 | 259.6208 | 571.0254 | 2.199459 | 1.137149 | 2.585497 |
| VC1190 | 492.91449 | 566.4213 | 568.4004 | 900.32022 | 890.3792 | 895.34428 | 1.01E-09 | 8.98E-09 | 542.5787 | 895.3479 | 1.650171 | 0.722616 | 2.843227 |
| VC1191 | 244.88745 | 238.555 | 181.8881 | 484.55091 | 528.7543 | 535.10437 | 3.75E-12 | 4.16E-11 | 221.7769 | 516.1365 | 2.327278 | 1.218644 | 2.52934 |
| VC1193 | 2070.0315 | 1178.673 | 2007.765 | 232.00954 | 507.2523 | 728.12416 | 3.64E-05 | 0.000193 | 1752.157 | 489.1287 | 0.279158 | -1.84085 | 2.966498 |
| VC1194 | 499.19365 | 696.8627 | 430.2354 | 1491.6366 | 1498.3 | 1307.1835 | 6.28E-10 | 5.78E-09 | 542.0972 | 1432.373 | 2.642281 | 1.401784 | 2.945067 |
| VC1195 | 5115.4268 | 7592.631 | 4577.809 | 19468.27 | 16330.79 | 16310.172 | 6.52E-11 | 6.59E-10 | 5761.956 | 17369.74 | 3.014557 | 1.591946 | 4.000182 |
| VC1198 | 2458.2933 | 2305.64 | 2733.569 | 4354.7986 | 3811.722 | 3838.4183 | 2.73E-07 | 1.92E-06 | 2499.167 | 4001.646 | 1.601192 | 0.679146 | 3.500017 |
| VC1207 | 1113.5053 | 1067.034 | 1000.385 | 3224.522 | 5105.753 | 4896.205 | 4.13E-20 | 6.96E-19 | 1060.308 | 4408.827 | 4.158062 | 2.055911 | 3.334878 |
| VC1210 | 1330.1366 | 1170.447 | 1093.078 | 797.66113 | 821.9637 | 796.9233 | 9.82E-06 | 5.62E-05 | 1197.887 | 805.516 | 0.672447 | -0.57251 | 2.992245 |
| VC1233 | 462.56519 | 465.3586 | 431.1098 | 309.00386 | 289.2999 | 302.9073 | 1.28E-06 | 8.26E-06 | 453.0112 | 300.4037 | 0.663126 | -0.59264 | 2.566907 |
| VC1253 | 130.81595 | 129.2663 | 111.9312 | 170.41409 | 251.1827 | 254.17458 | 0.000194 | 0.000885 | 124.0045 | 225.2571 | 1.816524 | 0.861181 | 2.223058 |
| VC1254 | 1368.8581 | 1667.535 | 1328.308 | 2390.9302 | 2505.963 | 2601.945 | 5.42E-08 | 4.11E-07 | 1454.9 | 2499.613 | 1.718065 | 0.780784 | 3.280353 |
| VC1255 | 10923.655 | 12706.87 | 10864.32 | 17006.505 | 21119.87 | 20947.425 | 5.6E-07 | 3.78E-06 | 11498.28 | 19691.27 | 1.71254 | 0.776138 | 4.177453 |
| VC1256 | 19187.037 | 26385.6 | 21662.18 | 35780.799 | 41197.88 | 41796.429 | 3.12E-06 | 1.9E-05 | 22411.6 | 39591.7 | 1.766572 | 0.820952 | 4.474039 |
| VC1257 | 1725.724 | 2024.78 | 1752.422 | 2902.1725 | 2605.654 | 2425.1695 | 0.000178 | 0.000819 | 1834.309 | 2644.332 | 1.441596 | 0.527667 | 3.342894 |
| VC1258 | 60201.499 | 66417.01 | 54793.8 | 79130.653 | 87238.59 | 86091.605 | 0.000169 | 0.000782 | 60470.77 | 84153.62 | 1.391641 | 0.476787 | 4.853309 |
| VC1262 | 961.75884 | 1367.872 | 865.7175 | 2486.4032 | 2691.662 | 3143.7382 | 3.05E-09 | 2.56E-08 | 1065.116 | 2773.935 | 2.604349 | 1.380923 | 3.235247 |
| VC1264 | 15805.706 | 16105.4 | 16999.54 | 3697.7804 | 3282.968 | 3227.826 | 1E-94 | 1.03E-92 | 16303.55 | 3402.858 | 0.208719 | -2.26037 | 3.872063 |
| VC1265 | 5244.1497 | 6936.898 | 5282.626 | 1813.9861 | 1711.366 | 1590.98 | 1.01E-24 | 2.13E-23 | 5821.224 | 1705.444 | 0.29297 | -1.77118 | 3.498426 |
| VC1266 | 3770.6389 | 4779.326 | 3594.914 | 1542.9661 | 1426.952 | 1398.9157 | 4.81E-19 | 7.84E-18 | 4048.293 | 1456.278 | 0.359726 | -1.47503 | 3.385258 |
| VC1267 | 2433.1766 | 3055.385 | 2248.242 | 808.95363 | 857.1488 | 730.99079 | 2.05E-21 | 3.66E-20 | 2578.934 | 799.0311 | 0.30983 | -1.69045 | 3.157002 |
| VC1269 | 968.03801 | 1029.43 | 1061.597 | 6106.1627 | 5634.507 | 5432.2649 | 1.1E-114 | 1.4E-112 | 1019.688 | 5724.312 | 5.613786 | 2.488974 | 3.383095 |
| VC1270 | 2365.1523 | 2720.467 | 2107.454 | 4730.5309 | 4440.168 | 4335.301 | 4.27E-10 | 4E-09 | 2397.691 | 4502 | 1.87764 | 0.90892 | 3.516599 |
| VC1271 | 19.884024 | 21.15266 | 14.86586 | 119.08454 | 101.6459 | 81.221199 | 3.6E-15 | 4.97E-14 | 18.63418 | 100.6506 | 5.401394 | 2.433332 | 1.636563 |
| VC1272 | 2628.8773 | 4726.445 | 1736.682 | 759.67727 | 727.1593 | 702.32449 | 4.27E-08 | 3.29E-07 | 3030.668 | 729.7204 | 0.240779 | -2.05422 | 3.172347 |
| VC1275 | 90.001372 | 96.36213 | 103.1865 | 175.54704 | 197.4277 | 173.90892 | 1.57E-07 | 1.13E-06 | 96.51668 | 182.2945 | 1.888736 | 0.917421 | 2.122688 |
| VC1293 | 4784.7241 | 5205.905 | 5259.89 | 8430.3645 | 7425.039 | 6997.4452 | 4.49E-06 | 2.65E-05 | 5083.506 | 7617.616 | 1.498496 | 0.583516 | 3.793991 |
| VC1301 | 13013.57 | 10380.08 | 14101.58 | 2407.3557 | 4464.602 | 6143.1893 | 2.42E-05 | 0.000131 | 12498.41 | 4338.382 | 0.347115 | -1.52652 | 3.867091 |
| VC1312 | 7339.2979 | 11155.68 | 5886.005 | 2708.1468 | 2375.974 | 2416.5696 | 4.98E-10 | 4.64E-09 | 8126.994 | 2500.23 | 0.307645 | -1.70066 | 3.653955 |
| VC1314 | 153.83955 | 142.1929 | 132.9182 | 363.41318 | 560.03 | 589.57035 | 1.86E-13 | 2.29E-12 | 142.9836 | 504.3378 | 3.527243 | 1.818541 | 2.429004 |
| VC1318 | 54339.898 | 78914.71 | 53018.64 | 122641.68 | 112094.9 | 102435.22 | 7.23E-05 | 0.000365 | 62091.08 | 112390.6 | 1.810093 | 0.856064 | 4.92188 |
| VC1319 | 1652.467 | 2460.76 | 1963.167 | 7307.274 | 6050.865 | 5555.53 | 3.83E-14 | 4.92E-13 | 2025.465 | 6304.556 | 3.112647 | 1.638142 | 3.55309 |
| VC1320 | 1901.5406 | 2230.431 | 2087.341 | 7253.8913 | 5779.157 | 5609.996 | 6.7E-22 | 1.22E-20 | 2073.104 | 6214.348 | 2.997605 | 1.58381 | 3.555008 |
| VC1329 | 1198.2741 | 2741.62 | 663.7168 | 385.99818 | 362.6023 | 340.17349 | 2.36E-05 | 0.000128 | 1534.537 | 362.9247 | 0.236504 | -2.08006 | 2.872897 |
| VC1339 | 74.303458 | 50.53136 | 69.08251 | 111.89841 | 130.9669 | 119.44294 | 6.74E-05 | 0.000341 | 64.63911 | 120.7694 | 1.868364 | 0.901776 | 1.946226 |
| VC1343 | 102.5597 | 115.1645 | 108.4333 | 216.61068 | 224.7939 | 205.44186 | 6.99E-10 | 6.4E-09 | 108.7192 | 215.6155 | 1.983233 | 0.987854 | 2.184993 |
| VC1344 | 275.23675 | 165.6959 | 306.0617 | 908.53295 | 814.1448 | 880.05558 | 5.06E-12 | 5.53E-11 | 248.9981 | 867.5778 | 3.484274 | 1.800858 | 2.667252 |
| VC1345 | 283.60897 | 227.9787 | 353.2827 | 712.45409 | 726.182 | 669.83601 | 2.91E-10 | 2.78E-09 | 288.2901 | 702.824 | 2.437905 | 1.285642 | 2.653338 |
| VC1346 | 254.3062 | 206.826 | 377.7676 | 631.3534 | 612.8076 | 573.32611 | 1.89E-05 | 0.000104 | 279.6333 | 605.8291 | 2.166513 | 1.115375 | 2.614469 |
| VC1347 | 188.37496 | 153.9444 | 197.6284 | 319.26977 | 313.7341 | 290.48523 | 3.08E-06 | 1.88E-05 | 179.9826 | 307.8297 | 1.71033 | 0.774275 | 2.371771 |
| VC1349 | 1003.6199 | 746.2189 | 1026.619 | 1470.0782 | 1460.183 | 1470.5815 | 0.000115 | 0.00055 | 925.4858 | 1466.947 | 1.585057 | 0.664534 | 3.066392 |
| VC1367 | 1823.051 | 1633.456 | 1645.738 | 545.11977 | 539.5053 | 462.48307 | 3.93E-38 | 1.2E-36 | 1700.748 | 515.7027 | 0.303221 | -1.72156 | 2.97152 |
| VC1373 | 2886.3231 | 2391.426 | 3448.879 | 1376.6584 | 1377.107 | 1318.6501 | 9.22E-10 | 8.25E-09 | 2908.876 | 1357.472 | 0.466665 | -1.09954 | 3.298228 |

| | | | | | | | | | | | | | |
|---|---|---|---|---|---|---|---|---|---|---|---|---|---|
| VC1374 | 5562.2941 | 7233.035 | 7794.081 | 3261.4793 | 3279.058 | 2928.7409 | 3.82E-10 | 3.59E-09 | 6863.137 | 3156.426 | 0.45991 | -1.12058 | 3.667859 |
| VC1375 | 2309.6864 | 2857.96 | 2992.409 | 1227.8027 | 1282.302 | 1196.3405 | 6.05E-14 | 7.75E-13 | 2720.018 | 1235.482 | 0.454218 | -1.13854 | 3.263204 |
| VC1377 | 1382.4629 | 1063.509 | 1408.758 | 759.67727 | 805.3485 | 751.05721 | 8.26E-06 | 4.76E-05 | 1284.91 | 772.0277 | 0.600842 | -0.73494 | 2.998253 |
| VC1379 | 476.17005 | 360.7704 | 616.4958 | 250.48818 | 261.9337 | 221.6861 | 6.44E-05 | 0.000329 | 484.4787 | 244.7027 | 0.505084 | -0.9854 | 2.536957 |
| VC1380 | 451.05339 | 591.0994 | 363.7762 | 250.48818 | 221.8618 | 214.04175 | 1.06E-05 | 6.02E-05 | 468.643 | 228.7972 | 0.488212 | -1.03442 | 2.515146 |
| VC1384 | 8857.8094 | 10090.99 | 8678.162 | 4511.867 | 4304.314 | 4138.459 | 7E-21 | 1.21E-19 | 9208.989 | 4318.213 | 0.468913 | -1.09261 | 3.799758 |
| VC1414 | 5987.1843 | 5742.948 | 6676.518 | 9955.8786 | 11179.1 | 12182.224 | 9.3E-10 | 8.31E-09 | 6135.55 | 11105.73 | 1.810063 | 0.85604 | 3.9167 |
| VC1415 | 101.51318 | 81.0852 | 77.82713 | 200.18523 | 181.7898 | 218.81947 | 1.17E-09 | 1.03E-08 | 86.8085 | 200.2648 | 2.306973 | 1.206001 | 2.120083 |
| VC1417 | 201.97982 | 237.3799 | 249.2217 | 550.25272 | 555.1431 | 552.30416 | 3.56E-19 | 5.85E-18 | 229.5271 | 552.5667 | 2.407413 | 1.267484 | 2.551609 |
| VC1418 | 1330.1366 | 1112.865 | 1397.39 | 13483.245 | 18140.87 | 18551.877 | 9.9E-80 | 7.93E-78 | 1280.131 | 16725.33 | 13.06533 | 3.707672 | 3.665314 |
| VC1419 | 431.16936 | 310.239 | 498.4434 | 3512.9941 | 4749.992 | 5539.2858 | 6.58E-38 | 1.99E-36 | 413.2839 | 4600.757 | 11.13219 | 3.476666 | 3.139539 |
| VC1420 | 100.46665 | 85.7858 | 98.81422 | 870.54909 | 1025.256 | 1074.0309 | 4.73E-82 | 3.87E-80 | 95.02222 | 989.9452 | 10.41804 | 3.381012 | 2.486718 |
| VC1421 | 141.28122 | 99.88757 | 178.3903 | 340.82818 | 406.5837 | 393.68393 | 1.22E-08 | 9.85E-08 | 139.853 | 380.3653 | 2.71975 | 1.443474 | 2.362936 |
| VC1422a | 121.3972 | 135.142 | 132.0438 | 474.285 | 548.3016 | 527.46002 | 2.11E-39 | 6.59E-38 | 129.5277 | 516.6822 | 3.988972 | 1.996017 | 2.412793 |
| VC1423 | 1460.9525 | 836.7053 | 1539.053 | 4462.5907 | 6068.457 | 5485.7754 | 6.88E-13 | 8.1E-12 | 1278.904 | 5338.941 | 4.174623 | 2.061646 | 3.417146 |
| VC1442 | 12707.984 | 11388.36 | 12640.35 | 5959.3602 | 7287.231 | 8505.2929 | 2.55E-05 | 0.000137 | 12245.56 | 7250.628 | 0.592102 | -0.75608 | 3.974177 |
| VC1443 | 868.61789 | 586.3988 | 811.5009 | 374.70568 | 439.8141 | 408.97263 | 1.64E-05 | 9.14E-05 | 755.5058 | 407.8308 | 0.539812 | -0.88947 | 2.744359 |
| VC1444 | 2850.7411 | 2276.261 | 2554.304 | 7858.5534 | 6531.728 | 6157.5225 | 4.49E-17 | 6.68E-16 | 2560.435 | 6849.268 | 2.67504 | 1.419561 | 3.621979 |
| VC1445 | 2277.244 | 1579.399 | 2236 | 5127.8216 | 4322.884 | 4126.0369 | 2.96E-08 | 2.31E-07 | 2030.881 | 4525.581 | 2.228383 | 1.155997 | 3.481679 |
| VC1451 | 32729.103 | 45202.06 | 28421.77 | 16331.008 | 17125.38 | 15361.318 | 2.39E-07 | 1.69E-06 | 35450.98 | 16272.57 | 0.459016 | -1.12338 | 4.380542 |
| VC1456 | 460.47213 | 400.7254 | 533.4219 | 760.70386 | 707.612 | 721.43536 | 4.37E-05 | 0.000229 | 464.8732 | 729.9171 | 1.570142 | 0.650895 | 2.765304 |
| VC1484 | 396.63395 | 324.3408 | 612.1235 | 4287.0436 | 5720.515 | 6499.607 | 1.05E-32 | 2.89E-31 | 444.3661 | 5502.389 | 12.38256 | 3.630237 | 3.194146 |
| VC1485 | 12936.127 | 14895 | 13193.88 | 20629.344 | 22358.19 | 22032.923 | 2.86E-09 | 2.42E-08 | 13675 | 21673.49 | 1.584898 | 0.66439 | 4.235928 |
| VC1496 | 11593.433 | 10408.28 | 15081.85 | 27617.348 | 28294.71 | 26041.428 | 7.29E-10 | 6.65E-09 | 12361.19 | 27317.83 | 2.209968 | 1.144025 | 4.264253 |
| VC1497 | 14711.038 | 15562.48 | 14899.09 | 24877.377 | 25318.63 | 23169.064 | 4.39E-15 | 5.99E-14 | 15057.54 | 24455.02 | 1.624105 | 0.699645 | 4.283061 |
| VC1498 | 4139.0166 | 3021.305 | 4514.848 | 7192.2959 | 7280.389 | 7329.0188 | 2.84E-06 | 1.74E-05 | 3891.723 | 7267.235 | 1.867356 | 0.900997 | 3.725756 |
| VC1505 | 2225.9642 | 2504.24 | 2324.32 | 18050.548 | 15721.89 | 14474.573 | 5.43E-87 | 5E-85 | 2351.508 | 16082.34 | 6.839158 | 2.773819 | 3.788848 |
| VC1510 | 63.838182 | 58.75739 | 78.70159 | 460.93932 | 485.7502 | 522.68231 | 8.85E-54 | 4.03E-52 | 67.09906 | 489.7906 | 7.299516 | 2.867801 | 2.258363 |
| VC1511 | 160.11872 | 153.9444 | 210.7454 | 959.86249 | 1058.486 | 1096.964 | 5.31E-45 | 1.88E-43 | 174.9362 | 1038.437 | 5.936094 | 2.569514 | 2.62963 |
| VC1512 | 70.117348 | 62.28284 | 90.94406 | 323.37613 | 485.7502 | 453.88317 | 1.12E-21 | 2.02E-20 | 74.44808 | 421.0032 | 5.65499 | 2.499525 | 2.248069 |
| VC1513 | 390.35479 | 323.1657 | 575.3961 | 2403.2493 | 3280.036 | 3113.1608 | 6.35E-24 | 1.26E-22 | 429.6388 | 2932.149 | 6.824682 | 2.770762 | 3.050145 |
| VC1514 | 33.488882 | 5.875739 | 61.21235 | 181.70659 | 211.1108 | 186.33099 | 6.69E-05 | 0.000339 | 33.52566 | 193.0494 | 5.75826 | 2.525633 | 1.905523 |
| VC1515 | 43.954158 | 16.45207 | 71.70589 | 333.64204 | 441.7688 | 391.77284 | 1.18E-11 | 1.26E-10 | 44.03737 | 389.0612 | 8.834796 | 3.143197 | 2.11692 |
| VC1516 | 135.00206 | 116.3396 | 291.1959 | 1807.8266 | 1916.612 | 2086.9071 | 3.56E-20 | 6.06E-19 | 180.8459 | 1937.115 | 10.71142 | 3.421077 | 2.772232 |
| VC1517 | 421.75061 | 302.013 | 446.8501 | 927.01158 | 1070.214 | 1042.498 | 7.89E-12 | 8.51E-11 | 390.2046 | 1013.241 | 2.596692 | 1.376675 | 2.798503 |
| VC1518 | 171.63052 | 108.1136 | 169.6457 | 323.37613 | 436.882 | 444.32774 | 4.61E-08 | 3.55E-07 | 149.7966 | 401.5286 | 2.680492 | 1.422498 | 2.389609 |
| VC1519 | 610.12558 | 505.3136 | 579.7684 | 1625.0934 | 1699.637 | 1586.2022 | 1.23E-33 | 3.42E-32 | 565.0692 | 1636.978 | 2.896951 | 1.534535 | 2.983072 |
| VC1523 | 1258.9727 | 862.5586 | 1453.356 | 3493.4888 | 4309.201 | 5127.4465 | 3.8E-12 | 4.21E-11 | 1191.629 | 4310.045 | 3.616935 | 1.854768 | 3.355311 |
| VC1524 | 190.46802 | 96.36213 | 210.7454 | 417.8225 | 607.9208 | 636.39199 | 5.58E-07 | 3.78E-06 | 165.8585 | 554.0451 | 3.340468 | 1.74005 | 2.481641 |
| VC1525 | 238.60829 | 231.5041 | 307.8107 | 554.35909 | 630.4002 | 688.94688 | 2.61E-11 | 2.71E-10 | 259.3077 | 624.5687 | 2.408601 | 1.268195 | 2.604698 |
| VC1527 | 1163.7387 | 1418.404 | 1247.857 | 1727.7525 | 1893.155 | 1996.1304 | 7.65E-05 | 0.000384 | 1276.667 | 1872.346 | 1.46659 | 0.552465 | 3.189232 |
| VC1537 | 1469.3247 | 1358.471 | 1330.057 | 504.05613 | 634.3097 | 616.32557 | 3.18E-17 | 4.8E-16 | 1385.951 | 584.8971 | 0.422019 | -1.24462 | 2.954414 |
| VC1538 | 2714.6925 | 2436.082 | 2500.962 | 1476.2377 | 1732.868 | 1894.8428 | 7.84E-05 | 0.000392 | 2550.579 | 1701.316 | 0.667031 | -0.58417 | 3.318712 |
| VC1547 | 5845.903 | 8197.832 | 4383.679 | 3328.2077 | 2880.294 | 2495.8797 | 0.000102 | 0.000498 | 6142.471 | 2901.46 | 0.47236 | -1.08204 | 3.62548 |
| VC1548 | 3007.7203 | 4234.058 | 2345.307 | 1666.157 | 1529.576 | 1198.2516 | 4.99E-05 | 0.000259 | 3195.695 | 1464.661 | 0.458323 | -1.12556 | 3.335151 |
| VC1572 | 101.51318 | 92.83668 | 79.57605 | 27.717954 | 22.47939 | 16.24424 | 2.21E-12 | 2.51E-11 | 91.30864 | 22.14719 | 0.242553 | -2.04363 | 1.652915 |
| VC1573 | 634.19571 | 678.0603 | 813.2498 | 237.1425 | 210.1334 | 183.46436 | 4.16E-21 | 7.27E-20 | 708.5019 | 210.2468 | 0.296748 | -1.75269 | 2.586535 |
| VC1575 | 605.93947 | 824.9538 | 622.617 | 1690.7952 | 1397.631 | 1409.4267 | 3.79E-09 | 3.16E-08 | 684.5034 | 1499.284 | 2.190324 | 1.131144 | 3.00563 |
| VC1577 | 12809.498 | 17057.27 | 11695.06 | 31903.365 | 28183.29 | 28131.201 | 2.71E-08 | 2.13E-07 | 13853.94 | 29405.95 | 2.122569 | 1.085812 | 4.305004 |

| | | | | | | | | | | | | | |
|---|---|---|---|---|---|---|---|---|---|---|---|---|---|
| VC1578 | 5558.108 | 6832.31 | 5108.608 | 11491.659 | 9815.673 | 10639.977 | 2.37E-07 | 1.68E-06 | 5833.009 | 10649.1 | 1.825662 | 0.86842 | 3.896603 |
| VC1579 | 17767.945 | 25934.34 | 17994.68 | 46346.473 | 38067.38 | 37474.506 | 7.34E-06 | 4.28E-05 | 20565.66 | 40629.45 | 1.975597 | 0.982289 | 4.460992 |
| VC1588 | 276.28328 | 244.4308 | 233.4814 | 385.99818 | 385.0817 | 362.151 | 2.81E-05 | 0.000151 | 251.3985 | 377.7436 | 1.502569 | 0.587431 | 2.48878 |
| VC1590 | 34.53541 | 34.07929 | 51.59327 | 217.63727 | 260.9564 | 212.13066 | 5.23E-24 | 1.04E-22 | 40.06932 | 230.2414 | 5.746078 | 2.522577 | 1.982498 |
| VC1591 | 75.349986 | 59.93254 | 63.83574 | 232.00954 | 272.6847 | 256.08566 | 3.23E-24 | 6.58E-23 | 66.37275 | 253.5933 | 3.820744 | 1.933854 | 2.113064 |
| VC1599 | 238.60829 | 224.4532 | 197.6284 | 113.95159 | 96.7591 | 90.776635 | 1.52E-09 | 1.32E-08 | 220.23 | 100.4958 | 0.456322 | -1.13188 | 2.172512 |
| VC1600 | 1110.3658 | 755.6201 | 908.5661 | 439.38091 | 548.3016 | 584.79264 | 0.000175 | 0.000806 | 924.8507 | 524.1584 | 0.566749 | -0.81922 | 2.842767 |
| VC1605 | 729.42972 | 452.4319 | 752.0374 | 241.24886 | 326.4398 | 336.35132 | 5.92E-05 | 0.000304 | 644.633 | 301.3467 | 0.46747 | -1.09705 | 2.644189 |
| VC1605a | 118.25762 | 135.142 | 111.9312 | 402.42363 | 464.2482 | 498.79372 | 2.62E-27 | 6.09E-26 | 121.7769 | 455.1552 | 3.737614 | 1.902118 | 2.371862 |
| VC1614 | 639.42835 | 823.7787 | 842.107 | 415.76932 | 477.9313 | 511.21578 | 9.08E-05 | 0.000448 | 768.438 | 468.3055 | 0.609425 | -0.71448 | 2.778069 |
| VC1641 | 365.23812 | 366.6461 | 327.0488 | 823.3259 | 769.186 | 788.32341 | 2.41E-22 | 4.49E-21 | 352.9777 | 793.6118 | 2.248334 | 1.168856 | 2.723678 |
| VC1642 | 218.72426 | 284.3858 | 212.4943 | 872.60227 | 818.0542 | 823.67852 | 2.71E-27 | 6.25E-26 | 238.5348 | 838.1117 | 3.513583 | 1.812943 | 2.650427 |
| VC1649 | 38360.468 | 51321.06 | 36284.93 | 15482.017 | 18707.74 | 21337.287 | 5.96E-08 | 4.52E-07 | 41988.82 | 18509.01 | 0.440808 | -1.18178 | 4.445258 |
| VC1652 | 1000.4804 | 1289.137 | 1000.385 | 2076.7934 | 1770.985 | 1622.5129 | 7.67E-05 | 0.000385 | 1096.667 | 1823.43 | 1.662701 | 0.733529 | 3.150482 |
| VC1653 | 875.94358 | 1188.075 | 911.1895 | 2260.5532 | 1823.762 | 1682.7121 | 3.9E-06 | 2.31E-05 | 991.7359 | 1922.343 | 1.938361 | 0.954838 | 3.140113 |
| VC1655 | 2555.6203 | 5482.065 | 2490.468 | 11723.668 | 10622 | 10825.353 | 2.13E-06 | 1.33E-05 | 3509.384 | 11057.01 | 3.150697 | 1.655671 | 3.794434 |
| VC1658 | 1526.8837 | 2712.241 | 858.7218 | 409.60977 | 405.6063 | 459.61643 | 1.32E-06 | 8.46E-06 | 1699.282 | 424.9442 | 0.250073 | -1.99958 | 2.929299 |
| VC1663 | 10242.365 | 13241.57 | 9872.677 | 64252.271 | 61535.86 | 58350.265 | 2.8E-49 | 1.09E-47 | 11118.87 | 61379.46 | 5.520297 | 2.464746 | 4.417042 |
| VC1664 | 992.10814 | 1047.057 | 1063.346 | 1699.0079 | 1854.061 | 1510.7143 | 8.49E-08 | 6.3E-07 | 1034.17 | 1687.928 | 1.632156 | 0.706779 | 3.120973 |
| VC1672 | 219.77079 | 209.1763 | 236.9792 | 1246.2814 | 1061.418 | 1054.92 | 1.62E-55 | 8.08E-54 | 221.9755 | 1120.873 | 5.049537 | 2.336151 | 2.697931 |
| VC1673 | 1964.3323 | 2110.566 | 2226.381 | 13056.183 | 12278.63 | 11827.718 | 1.8E-123 | 2.8E-121 | 2100.426 | 12387.51 | 5.897618 | 2.560132 | 3.707646 |
| VC1674 | 752.45333 | 740.3432 | 808.8775 | 5040.5613 | 4248.604 | 4349.6341 | 8.81E-87 | 7.93E-85 | 767.2247 | 4546.267 | 5.9256 | 2.566961 | 3.271289 |
| VC1675 | 853.9665 | 914.2651 | 932.1766 | 4733.6107 | 3792.175 | 3853.707 | 6.18E-50 | 2.48E-48 | 900.1361 | 4126.498 | 4.584304 | 2.196703 | 3.284945 |
| VC1676 | 731.52278 | 678.0603 | 771.2756 | 955.75613 | 1074.124 | 994.72081 | 0.000103 | 0.0005 | 726.9529 | 1008.2 | 1.386885 | 0.471848 | 2.932527 |
| VC1681 | 1241.1817 | 1255.058 | 1114.065 | 840.77795 | 907.9718 | 908.72189 | 0.000122 | 0.000581 | 1203.435 | 885.8239 | 0.73608 | -0.44207 | 3.013885 |
| VC1688 | 453.14644 | 486.5112 | 307.8107 | 107.79204 | 91.87228 | 101.28761 | 4.98E-17 | 7.37E-16 | 415.8228 | 100.3173 | 0.24125 | -2.0514 | 2.310142 |
| VC1695 | 968.03801 | 1872.011 | 755.5353 | 3394.9361 | 2990.736 | 2904.8523 | 0.000147 | 0.000693 | 1198.528 | 3096.841 | 2.583871 | 1.369534 | 3.284784 |
| VC1704 | 189.42149 | 156.2947 | 177.5158 | 332.61545 | 314.7114 | 321.06262 | 2.19E-09 | 1.89E-08 | 174.4107 | 322.7965 | 1.850784 | 0.888137 | 2.375251 |
| VC1709 | 4015.5263 | 4458.511 | 3664.871 | 2327.2816 | 2132.61 | 2075.4405 | 1.72E-11 | 1.81E-10 | 4046.303 | 2178.444 | 0.538379 | -0.89331 | 3.472602 |
| VC1712 | 115.11803 | 95.18698 | 134.6672 | 321.32295 | 303.9604 | 368.8398 | 2.98E-14 | 3.89E-13 | 114.9907 | 331.3744 | 2.881749 | 1.526945 | 2.290491 |
| VC1714 | 20004.375 | 21340.69 | 20127.49 | 28177.867 | 29816.46 | 29558.783 | 8.59E-10 | 7.72E-09 | 20490.85 | 29184.37 | 1.424263 | 0.510216 | 4.388355 |
| VC1722 | 8461.1755 | 9266.041 | 7580.712 | 26535.322 | 21757.11 | 25672.588 | 1.77E-24 | 3.63E-23 | 8435.976 | 24655.01 | 2.922603 | 1.547254 | 4.15902 |
| VC1723 | 194.65413 | 238.555 | 181.0137 | 531.77409 | 461.3161 | 418.52806 | 2.31E-10 | 2.23E-09 | 204.7409 | 470.5394 | 2.298219 | 1.200516 | 2.4919 |
| VC1724 | 30.3493 | 42.30532 | 48.96988 | 118.05795 | 129.9895 | 116.57631 | 1.21E-10 | 1.21E-09 | 40.5415 | 121.5413 | 2.997947 | 1.583975 | 1.846312 |
| VC1730 | 30962.565 | 27227 | 31108.99 | 17802.113 | 19878.62 | 20161.968 | 2.24E-07 | 1.6E-06 | 29766.19 | 19280.9 | 0.647745 | -0.6265 | 4.379425 |
| VC1748 | 280.46939 | 319.6402 | 258.8408 | 172.46727 | 119.2385 | 147.1537 | 2.94E-06 | 1.79E-05 | 286.3168 | 146.2865 | 0.510925 | -0.96882 | 2.311026 |
| VC1750 | 687.56862 | 745.0438 | 766.9033 | 544.09318 | 477.9313 | 448.14991 | 2.99E-05 | 0.00016 | 733.1719 | 490.0581 | 0.668408 | -0.5812 | 2.777727 |
| VC1753 | 918.85121 | 1124.617 | 943.5446 | 662.15113 | 593.2604 | 571.41503 | 2.56E-06 | 1.58E-05 | 995.6708 | 608.9422 | 0.61159 | -0.70936 | 2.891346 |
| VC1754 | 3956.9208 | 4049.56 | 4237.643 | 2877.5343 | 2832.403 | 2699.4105 | 8.54E-10 | 7.7E-09 | 4081.375 | 2803.116 | 0.686807 | -0.54202 | 3.529224 |
| VC1764 | 556.75267 | 522.9408 | 661.0934 | 383.945 | 377.2628 | 391.77284 | 7.35E-05 | 0.00037 | 580.2623 | 384.3269 | 0.662333 | -0.59437 | 2.674163 |
| VC1775 | 1313.3921 | 1272.685 | 1299.451 | 968.07522 | 1021.346 | 1022.4316 | 5.93E-05 | 0.000304 | 1295.176 | 1003.951 | 0.775146 | -0.36746 | 3.057021 |
| VC1784 | 612.21863 | 329.0414 | 821.9944 | 1077.9204 | 1988.937 | 2039.1299 | 0.000178 | 0.000816 | 587.7515 | 1701.996 | 2.895775 | 1.533949 | 3.000076 |
| VC1831 | 647.80057 | 674.5349 | 684.7038 | 1028.6441 | 1498.3 | 1510.7143 | 6.2E-07 | 4.13E-06 | 669.0131 | 1345.886 | 2.011749 | 1.00845 | 2.977221 |
| VC1834 | 10134.573 | 12770.33 | 10984.12 | 30570.85 | 25953.92 | 26805.862 | 1.5E-17 | 2.3E-16 | 11296.34 | 27776.88 | 2.458927 | 1.298029 | 4.248311 |
| VC1835 | 41347.258 | 48048.27 | 39635.87 | 87954.202 | 81333.36 | 80923.07 | 1.05E-13 | 1.32E-12 | 43010.47 | 83403.54 | 1.939145 | 0.955421 | 4.777379 |
| VC1836 | 19912.28 | 23615.77 | 20042.67 | 55771.604 | 50475.02 | 52001.634 | 4.45E-25 | 9.53E-24 | 21190.24 | 52749.42 | 2.489326 | 1.315755 | 4.524177 |
| VC1837 | 1659.7927 | 1562.947 | 1699.08 | 6120.535 | 6018.612 | 5539.2858 | 8.49E-73 | 5.8E-71 | 1640.606 | 5892.811 | 3.591849 | 1.844727 | 3.492663 |
| VC1838 | 2004.1003 | 2015.379 | 1693.833 | 6527.065 | 6062.593 | 6609.4945 | 5.04E-43 | 1.69E-41 | 1904.437 | 6399.718 | 3.360424 | 1.748643 | 3.542964 |
| VC1839 | 3330.0508 | 4090.69 | 2971.422 | 10355.222 | 9648.544 | 9896.5643 | 3.58E-20 | 6.06E-19 | 3464.054 | 9966.777 | 2.8772 | 1.524665 | 3.76907 |

| | | | | | | | | | | | | | |
|---|---|---|---|---|---|---|---|---|---|---|---|---|---|
| VC1854 | 1355.2532 | 1099.938 | 1218.126 | 625.19386 | 759.4124 | 780.67906 | 3.28E-06 | 1.97E-05 | 1224.439 | 721.7618 | 0.589463 | -0.76253 | 2.973166 |
| VC1866 | 28123.336 | 26853.3 | 38339.04 | 55022.193 | 68415.53 | 61204.474 | 1.3E-06 | 8.34E-06 | 31105.23 | 61547.4 | 1.978683 | 0.984541 | 4.641022 |
| VC1871 | 4733.4442 | 6412.782 | 4841.022 | 1809.8798 | 2314.4 | 2272.2825 | 4.04E-11 | 4.14E-10 | 5329.083 | 2132.187 | 0.400104 | -1.32155 | 3.527739 |
| VC1879 | 114.07151 | 116.3396 | 109.3078 | 179.65341 | 210.1334 | 175.82001 | 1.96E-05 | 0.000108 | 113.2396 | 188.5356 | 1.664926 | 0.735458 | 2.164696 |
| VC1880 | 570.35753 | 840.2307 | 547.4133 | 373.67909 | 354.7834 | 347.81784 | 0.000107 | 0.000519 | 652.6672 | 358.7601 | 0.549683 | -0.86333 | 2.684748 |
| VC1882 | 1762.3524 | 1615.828 | 1734.933 | 4778.7807 | 4500.764 | 4218.7247 | 2.72E-38 | 8.37E-37 | 1704.371 | 4499.423 | 2.639931 | 1.4005 | 3.442361 |
| VC1883 | 1315.4852 | 1270.335 | 1330.057 | 3664.9295 | 3238.987 | 3316.6916 | 3.14E-39 | 9.73E-38 | 1305.292 | 3406.869 | 2.610043 | 1.384074 | 3.324032 |
| VC1884 | 1597.0011 | 1480.686 | 1351.044 | 3904.1252 | 3371.908 | 3114.1163 | 4.91E-16 | 7.04E-15 | 1476.244 | 3463.383 | 2.346078 | 1.230251 | 3.354329 |
| VC1886 | 5792.5301 | 5906.293 | 6438.665 | 8712.677 | 8125.81 | 7567.9047 | 0.000182 | 0.00083 | 6045.829 | 8135.464 | 1.345632 | 0.428284 | 3.845919 |
| VC1887 | 3969.4791 | 4586.602 | 4168.561 | 10740.194 | 9017.166 | 8894.1991 | 1.06E-16 | 1.55E-15 | 4241.547 | 9550.52 | 2.251659 | 1.170989 | 3.803776 |
| VC1888 | 1444.2081 | 1608.777 | 1440.239 | 2666.0566 | 2422.887 | 2563.7233 | 3.01E-12 | 3.38E-11 | 1497.742 | 2550.889 | 1.703157 | 0.768211 | 3.291064 |
| VC1892 | 893.73455 | 730.942 | 1086.956 | 2613.7004 | 2389.657 | 2416.5696 | 2.41E-14 | 3.18E-13 | 903.8777 | 2473.309 | 2.736331 | 1.452243 | 3.174694 |
| VC1893 | 2509.5731 | 2271.561 | 2807.023 | 12159.969 | 11584.7 | 11690.119 | 8.5E-69 | 5.41E-67 | 2529.386 | 11811.6 | 4.669749 | 2.223345 | 3.737662 |
| VC1894 | 6646.4966 | 6594.93 | 7291.265 | 29687.982 | 27963.38 | 26802.996 | 1.15E-85 | 1.01E-83 | 6844.231 | 28151.45 | 4.113165 | 2.040249 | 4.142413 |
| VC1895 | 1367.8115 | 1380.799 | 1608.136 | 5964.4931 | 5365.732 | 5337.6661 | 4.07E-50 | 1.67E-48 | 1452.249 | 5555.964 | 3.825766 | 1.935749 | 3.4534 |
| VC1896 | 4495.8825 | 4150.622 | 4647.766 | 8784.5384 | 7643.969 | 7121.6659 | 1.43E-09 | 1.25E-08 | 4431.424 | 7850.058 | 1.771453 | 0.824933 | 3.770708 |
| VC1897 | 3517.3792 | 2788.626 | 3368.428 | 2039.8361 | 1720.162 | 1755.3335 | 3.19E-07 | 2.22E-06 | 3224.811 | 1838.444 | 0.570093 | -0.81073 | 3.386477 |
| VC1900 | 4077.2714 | 4889.79 | 3657.875 | 7803.1174 | 6313.776 | 6581.7838 | 6.48E-05 | 0.00033 | 4208.312 | 6899.559 | 1.639507 | 0.713262 | 3.731465 |
| VC1903 | 7166.6209 | 9761.954 | 6362.586 | 38075.23 | 33806.07 | 32693.922 | 1.28E-24 | 2.64E-23 | 7763.72 | 34858.41 | 4.48991 | 2.166687 | 4.216189 |
| VC1907 | 5888.8107 | 5299.917 | 5639.406 | 3567.4034 | 3704.212 | 3679.7981 | 2.08E-11 | 2.17E-10 | 5609.378 | 3650.471 | 0.65078 | -0.61976 | 3.655632 |
| VC1910 | 1949.6809 | 2745.145 | 1643.114 | 5372.1502 | 4447.987 | 4492.9656 | 1.12E-06 | 7.32E-06 | 2112.647 | 4771.034 | 2.258321 | 1.17525 | 3.50172 |
| VC1918 | 27718.329 | 33230.83 | 24977.26 | 51258.71 | 50779.96 | 52924.689 | 2.27E-08 | 1.79E-07 | 28642.14 | 51654.45 | 1.803442 | 0.850753 | 4.585057 |
| VC1944 | 150.69997 | 146.8935 | 144.2863 | 4627.8718 | 4578.953 | 4423.211 | 0 | 0 | 147.2932 | 4543.345 | 30.84558 | 4.946992 | 2.912779 |
| VC1945 | 192.56107 | 132.7917 | 221.2389 | 8051.5524 | 8123.855 | 7634.7927 | 6E-141 | 1.2E-138 | 182.1972 | 7936.733 | 43.56122 | 5.444972 | 3.080092 |
| VC1947 | 283.60897 | 283.2106 | 315.6808 | 2889.8534 | 3259.511 | 3323.3804 | 2.5E-146 | 5E-144 | 294.1668 | 3157.582 | 10.73398 | 3.424114 | 2.983974 |
| VC1948 | 3.1395827 | 0 | 4.372311 | 137.56318 | 144.65 | 146.19816 | 1.79E-34 | 5.03E-33 | 2.503964 | 142.8038 | 57.03107 | 5.833676 | 1.276684 |
| VC1949 | 53.372906 | 50.53136 | 53.34219 | 1195.9784 | 1361.469 | 1297.6281 | 8.6E-157 | 1.9E-154 | 52.41549 | 1285.025 | 24.51614 | 4.61566 | 2.414186 |
| VC1952 | 244.88745 | 267.9337 | 278.079 | 428.08841 | 518.0033 | 625.88101 | 6.63E-07 | 4.4E-06 | 263.6334 | 523.9909 | 1.987574 | 0.991009 | 2.570162 |
| VC1955 | 277.32981 | 383.0982 | 298.1916 | 978.34113 | 807.3032 | 850.43374 | 1.26E-14 | 1.7E-13 | 319.5399 | 878.6927 | 2.749869 | 1.459363 | 2.724181 |
| VC1956 | 4750.1887 | 5305.793 | 4869.005 | 19711.572 | 17923.89 | 18658.898 | 3.33E-76 | 2.36E-74 | 4974.996 | 18764.79 | 3.77182 | 1.915261 | 3.985068 |
| VC1959 | 2671.7849 | 3082.413 | 2456.364 | 5972.7059 | 5470.31 | 5640.5734 | 1.32E-14 | 1.77E-13 | 2736.854 | 5694.53 | 2.080685 | 1.057058 | 3.596355 |
| VC1960 | 5444.0365 | 6464.489 | 5277.379 | 10392.18 | 10185.12 | 10391.536 | 3.21E-11 | 3.32E-10 | 5728.635 | 10322.94 | 1.80199 | 0.849591 | 3.885927 |
| VC1961 | 1267.3449 | 1474.811 | 1144.671 | 2061.3945 | 2092.538 | 2220.6831 | 9.1E-07 | 5.95E-06 | 1295.609 | 2124.872 | 1.640057 | 0.713746 | 3.219903 |
| VC1962 | 2154.8003 | 2641.732 | 2049.739 | 94843.654 | 82743.69 | 85509.679 | 6.1E-248 | 2.5E-245 | 2282.091 | 87699.01 | 38.42924 | 5.264133 | 4.150664 |
| VC1964 | 3705.7542 | 3031.882 | 3984.924 | 6072.2852 | 5769.384 | 5844.1042 | 1.46E-06 | 9.3E-06 | 3574.187 | 5895.258 | 1.649398 | 0.72194 | 3.66184 |
| VC1974 | 1390.8352 | 1618.179 | 1411.382 | 921.87863 | 918.7228 | 913.49961 | 6.09E-09 | 5.03E-08 | 1473.465 | 918.0337 | 0.623044 | -0.68259 | 3.065599 |
| VC1975 | 3693.1958 | 3873.287 | 3643.009 | 3060.2675 | 2993.668 | 2933.5186 | 0.00012 | 0.000575 | 3736.497 | 2995.818 | 0.801772 | -0.31874 | 3.52449 |
| VC1976 | 2854.9272 | 3081.238 | 2655.741 | 1704.1409 | 1838.423 | 1724.7561 | 5.24E-10 | 4.87E-09 | 2863.969 | 1755.773 | 0.613056 | -0.70591 | 3.350718 |
| VC1978 | 308.72564 | 363.1207 | 381.2655 | 521.50818 | 572.7357 | 605.81459 | 6.92E-06 | 4.05E-05 | 351.0373 | 566.6862 | 1.614319 | 0.690926 | 2.649348 |
| VC1983 | 2780.6238 | 3150.571 | 2948.686 | 5647.2766 | 5648.19 | 5458.0646 | 9.94E-21 | 1.7E-19 | 2959.961 | 5584.511 | 1.886684 | 0.915853 | 3.609136 |
| VC1984 | 2003.0538 | 2795.677 | 2059.358 | 3980.0929 | 4061.928 | 4209.1692 | 3.84E-06 | 2.29E-05 | 2286.03 | 4083.73 | 1.786385 | 0.837043 | 3.485069 |
| VC1987 | 1983.1698 | 2036.531 | 1851.236 | 2690.6948 | 2605.654 | 2549.3901 | 1.06E-05 | 6.02E-05 | 1956.979 | 2615.246 | 1.336369 | 0.418318 | 3.354549 |
| VC1995 | 10720.629 | 10499.95 | 10480.43 | 4475.9363 | 4387.39 | 4741.407 | 4.41E-50 | 1.79E-48 | 10567 | 4534.911 | 0.429158 | -1.22042 | 3.84026 |
| VC2002 | 383.02909 | 412.4769 | 445.1012 | 685.76272 | 590.3283 | 601.03688 | 2.4E-05 | 0.00013 | 413.5357 | 625.7093 | 1.513072 | 0.59748 | 2.706443 |
| VC2005 | 368.37771 | 439.5053 | 424.1141 | 139.61636 | 124.1253 | 128.04283 | 7.61E-25 | 1.6E-23 | 410.6657 | 130.5948 | 0.318008 | -1.65287 | 2.364707 |
| VC2027 | 659.31237 | 642.8059 | 524.6773 | 401.39704 | 413.4253 | 401.32828 | 9.88E-05 | 0.000483 | 608.9318 | 405.3835 | 0.665729 | -0.58699 | 2.696217 |
| VC2032 | 168.49094 | 195.0746 | 148.6586 | 711.4275 | 705.6573 | 685.12471 | 4.7E-40 | 1.49E-38 | 170.7414 | 700.7365 | 4.104082 | 2.03706 | 2.538947 |
| VC2033 | 17980.39 | 19134.93 | 24641.47 | 49784.526 | 58266.57 | 48621.877 | 3.61E-13 | 4.36E-12 | 20585.6 | 52224.32 | 2.536935 | 1.343087 | 4.515718 |
| VC2037 | 20327.752 | 23471.23 | 18264.89 | 9478.5138 | 11770.4 | 13726.383 | 4.51E-05 | 0.000236 | 20687.96 | 11658.43 | 0.563537 | -0.82742 | 4.191179 |

| | | | | | | | | | | | | | |
|---|---|---|---|---|---|---|---|---|---|---|---|---|---|
| VC2046 | 248.02704 | 366.6461 | 308.6851 | 153.98864 | 136.8311 | 107.02087 | 4.96E-07 | 3.39E-06 | 307.7861 | 132.6135 | 0.430863 | -1.2147 | 2.305418 |
| VC2049 | 2481.3169 | 2909.666 | 2539.438 | 3748.0834 | 3863.523 | 3854.6626 | 2.92E-06 | 1.79E-05 | 2643.474 | 3822.09 | 1.445859 | 0.531927 | 3.502238 |
| VC2058 | 2695.855 | 3351.522 | 3218.895 | 2173.2929 | 2112.085 | 2163.3505 | 8.8E-05 | 0.000436 | 3088.757 | 2149.576 | 0.695936 | -0.52297 | 3.411068 |
| VC2067 | 4579.6047 | 4760.524 | 4937.213 | 3546.8716 | 3549.788 | 3714.1977 | 3.19E-06 | 1.93E-05 | 4759.114 | 3603.619 | 0.757204 | -0.40125 | 3.617133 |
| VC2069 | 2990.9758 | 3279.838 | 3073.734 | 2319.0688 | 2360.336 | 2467.2134 | 4.57E-05 | 0.000239 | 3114.849 | 2382.206 | 0.76479 | -0.38686 | 3.435208 |
| VC2076 | 550.47351 | 531.1668 | 421.4907 | 330.56227 | 325.4624 | 307.68501 | 0.000121 | 0.000577 | 501.0437 | 321.2366 | 0.641135 | -0.6413 | 2.60335 |
| VC2077 | 7124.7597 | 8381.155 | 6371.331 | 4435.8993 | 4156.732 | 3845.1071 | 3E-07 | 2.1E-06 | 7292.415 | 4145.913 | 0.568524 | -0.81471 | 3.740246 |
| VC2078 | 531.63601 | 639.2805 | 484.452 | 306.95068 | 219.9071 | 248.44132 | 1.37E-07 | 1E-06 | 551.7895 | 258.433 | 0.468354 | -1.09433 | 2.577061 |
| VC2095 | 11637.387 | 12125.18 | 11925.04 | 7444.8372 | 9011.302 | 8828.2666 | 6.55E-05 | 0.000333 | 11895.87 | 8428.135 | 0.708493 | -0.49718 | 4.000564 |
| VC2097 | 2529.4572 | 3212.854 | 2442.373 | 4814.7113 | 4220.261 | 3928.2394 | 0.00018 | 0.000822 | 2728.228 | 4321.07 | 1.583838 | 0.663425 | 3.535736 |
| VC2125 | 3202.3744 | 2988.401 | 3295.848 | 2065.5009 | 2338.834 | 2436.636 | 0.000107 | 0.000518 | 3162.208 | 2280.324 | 0.721118 | -0.47169 | 3.428993 |
| VC2127 | 2614.2259 | 2473.686 | 2624.261 | 1536.8066 | 1772.94 | 1673.1567 | 8.19E-09 | 6.66E-08 | 2570.724 | 1660.968 | 0.646109 | -0.63015 | 3.315208 |
| VC2128 | 2274.1044 | 2252.759 | 2816.643 | 827.43227 | 812.19 | 754.87938 | 7.29E-27 | 1.63E-25 | 2447.835 | 798.1672 | 0.326071 | -1.61674 | 3.145438 |
| VC2134 | 1224.4373 | 1101.114 | 1184.022 | 794.58136 | 849.3299 | 904.89972 | 8.71E-05 | 0.000432 | 1169.858 | 849.6037 | 0.726245 | -0.46147 | 2.998675 |
| VC2136 | 2032.3566 | 2089.413 | 2085.592 | 1578.8968 | 1598.969 | 1608.1797 | 2.7E-06 | 1.66E-05 | 2069.121 | 1595.348 | 0.771027 | -0.37515 | 3.259321 |
| VC2137 | 6284.3981 | 6987.429 | 6042.533 | 4352.7454 | 4792.996 | 4858.9388 | 9.36E-05 | 0.00046 | 6438.12 | 4668.227 | 0.725092 | -0.46376 | 3.738956 |
| VC2138 | 2104.567 | 1754.496 | 2016.51 | 640.59272 | 716.4083 | 684.16916 | 3.81E-30 | 9.75E-29 | 1958.524 | 680.3901 | 0.347399 | -1.52533 | 3.062343 |
| VC2139 | 1138.622 | 991.8248 | 1147.294 | 648.80545 | 644.0833 | 637.34753 | 1.37E-10 | 1.34E-09 | 1092.58 | 643.4121 | 0.588892 | -0.76392 | 2.923471 |
| VC2140 | 4988.797 | 5301.092 | 5572.947 | 3697.7804 | 3884.047 | 3785.8634 | 4.46E-07 | 3.07E-06 | 5287.612 | 3789.23 | 0.716624 | -0.48071 | 3.650905 |
| VC2141 | 4277.1582 | 4491.415 | 4642.519 | 3379.5373 | 3151.024 | 3130.3606 | 5.82E-07 | 3.91E-06 | 4470.364 | 3220.307 | 0.720368 | -0.47319 | 3.57912 |
| VC2142 | 4986.7039 | 5473.839 | 5100.738 | 2120.9368 | 2129.678 | 2176.7281 | 6.66E-45 | 2.34E-43 | 5187.093 | 2142.448 | 0.413034 | -1.27567 | 3.522917 |
| VC2143 | 12300.885 | 10781.98 | 14669.98 | 2024.4373 | 2354.471 | 2068.7517 | 2.17E-49 | 8.53E-48 | 12584.28 | 2149.22 | 0.170786 | -2.54974 | 3.716055 |
| VC2145 | 7139.4111 | 6109.594 | 7731.994 | 20098.597 | 19626.46 | 18769.741 | 2.9E-27 | 6.65E-26 | 6993.666 | 19498.27 | 2.787989 | 1.479225 | 4.06735 |
| VC2146 | 1183.6227 | 1090.537 | 1055.476 | 3191.6711 | 2933.071 | 2821.72 | 9.11E-35 | 2.59E-33 | 1109.879 | 2982.154 | 2.686919 | 1.425953 | 3.259903 |
| VC2156 | 10017.362 | 10455.29 | 9933.89 | 18658.29 | 21357.37 | 22952.155 | 8.73E-17 | 1.28E-15 | 10135.51 | 20989.27 | 2.070864 | 1.050233 | 4.163922 |
| VC2157 | 9967.1286 | 8282.442 | 9966.245 | 12995.614 | 12817.16 | 13126.301 | 0.000198 | 0.000899 | 9405.272 | 12979.69 | 1.380044 | 0.464715 | 4.043318 |
| VC2162 | 1460.9525 | 1575.873 | 1414.005 | 6508.5863 | 5627.666 | 5539.2858 | 3.28E-55 | 1.57E-53 | 1483.61 | 5891.846 | 3.971289 | 1.989607 | 3.470786 |
| VC2163 | 335.93535 | 457.1325 | 328.7978 | 1519.3545 | 1138.63 | 1027.2093 | 9.07E-13 | 1.05E-11 | 373.9552 | 1228.398 | 3.28488 | 1.715841 | 2.831079 |
| VC2164 | 3583.3104 | 3259.86 | 3754.94 | 6778.5797 | 6367.531 | 6054.3238 | 1.18E-13 | 1.47E-12 | 3532.704 | 6400.145 | 1.811685 | 0.857332 | 3.677149 |
| VC2165 | 1679.6768 | 1518.291 | 1844.241 | 3224.522 | 3119.748 | 3251.7146 | 2.92E-14 | 3.82E-13 | 1680.736 | 3198.662 | 1.903131 | 0.928375 | 3.365234 |
| VC2166 | 1332.2296 | 1329.092 | 1597.642 | 2383.7441 | 2372.064 | 2285.6601 | 2.12E-08 | 1.68E-07 | 1419.655 | 2347.156 | 1.653329 | 0.725374 | 3.261362 |
| VC2167 | 271.05064 | 299.6627 | 283.3257 | 653.9384 | 525.8222 | 537.971 | 1.56E-10 | 1.52E-09 | 284.6797 | 572.5772 | 2.011303 | 1.008131 | 2.606095 |
| VC2168 | 4487.5103 | 4363.324 | 3956.067 | 12281.107 | 11115.57 | 10845.419 | 4.21E-34 | 1.18E-32 | 4268.967 | 11414.03 | 2.673722 | 1.41885 | 3.843881 |
| VC2175 | 8751.0636 | 10692.67 | 8184.965 | 6334.0659 | 6155.443 | 6134.5894 | 0.000137 | 0.000648 | 9209.567 | 6208.033 | 0.674085 | -0.569 | 3.878597 |
| VC2179 | 8227.7998 | 10302.52 | 8443.806 | 6089.7372 | 6092.891 | 6008.4577 | 3.73E-05 | 0.000197 | 8991.376 | 6063.695 | 0.67439 | -0.56834 | 3.868282 |
| VC2187 | 1258.9727 | 864.9088 | 1854.734 | 335.69523 | 383.127 | 340.17349 | 5.6E-10 | 5.17E-09 | 1326.205 | 352.9986 | 0.266172 | -1.90957 | 2.835192 |
| VC2188 | 13913.584 | 14191.09 | 16374.3 | 2778.9816 | 2931.117 | 3181.0044 | 2.56E-71 | 1.68E-69 | 14826.32 | 2963.701 | 0.199895 | -2.32269 | 3.821434 |
| VC2190 | 1646.1879 | 1666.36 | 2164.294 | 372.6525 | 367.4891 | 299.08512 | 9.22E-36 | 2.64E-34 | 1825.614 | 346.4089 | 0.189749 | -2.39783 | 2.900499 |
| VC2191 | 3401.2146 | 3785.151 | 3971.807 | 714.50727 | 623.5587 | 707.10221 | 2.73E-77 | 2.06E-75 | 3719.391 | 681.7227 | 0.183289 | -2.44781 | 3.20204 |
| VC2192 | 1717.3518 | 1521.817 | 2118.822 | 377.78545 | 481.8408 | 421.39469 | 1.3E-25 | 2.83E-24 | 1785.997 | 427.007 | 0.239086 | -2.0644 | 2.941158 |
| VC2193 | 2692.7155 | 2420.805 | 3016.02 | 533.82727 | 573.7131 | 563.77068 | 7.29E-60 | 3.84E-58 | 2709.847 | 557.1037 | 0.205585 | -2.28219 | 3.08944 |
| VC2194 | 2473.9912 | 2693.439 | 2942.565 | 498.92318 | 523.8675 | 534.14883 | 4.44E-79 | 3.49E-77 | 2703.332 | 518.9798 | 0.191978 | -2.38099 | 3.073525 |
| VC2195 | 3275.6313 | 3548.947 | 3806.534 | 577.97068 | 667.5401 | 608.68122 | 1.97E-84 | 1.65E-82 | 3543.704 | 618.064 | 0.174412 | -2.51943 | 3.170245 |
| VC2196 | 3241.0959 | 4484.364 | 4419.532 | 775.07613 | 791.6654 | 773.03471 | 3.69E-40 | 1.18E-38 | 4048.331 | 779.9254 | 0.192654 | -2.37592 | 3.249665 |
| VC2197 | 15356.746 | 15907.98 | 19162.09 | 4447.1918 | 4520.312 | 4701.2741 | 8.71E-43 | 2.89E-41 | 16808.94 | 4556.259 | 0.271062 | -1.88331 | 3.942074 |
| VC2198 | 6655.9154 | 7177.803 | 7959.354 | 2137.3623 | 2197.116 | 2159.5284 | 1.81E-50 | 7.68E-49 | 7264.358 | 2164.669 | 0.297985 | -1.74669 | 3.598294 |
| VC2199 | 3721.4521 | 4023.706 | 4518.346 | 1246.2814 | 1125.924 | 1175.3185 | 3.25E-42 | 1.05E-40 | 4087.835 | 1182.508 | 0.289275 | -1.78949 | 3.342149 |
| VC2200 | 6563.821 | 7122.571 | 8370.351 | 1883.7943 | 1547.168 | 1531.7363 | 8.31E-37 | 2.45E-35 | 7352.248 | 1654.233 | 0.224997 | -2.15202 | 3.542508 |
| VC2201 | 7236.7382 | 7070.865 | 7956.731 | 4735.6638 | 4534.972 | 4604.7642 | 4.55E-12 | 5.01E-11 | 7421.445 | 4625.133 | 0.623212 | -0.68221 | 3.767806 |

| | | | | | | | | | | | | | |
|---|---|---|---|---|---|---|---|---|---|---|---|---|---|
| VC2202 | 7906.5158 | 7893.468 | 8192.836 | 4391.7559 | 4473.398 | 4594.2533 | 5.23E-28 | 1.25E-26 | 7997.607 | 4486.469 | 0.560976 | -0.83399 | 3.777432 |
| VC2206 | 1817.8184 | 2098.814 | 2252.614 | 416.79591 | 476.954 | 421.39469 | 7.63E-50 | 3.03E-48 | 2056.416 | 438.3815 | 0.213177 | -2.22987 | 2.977482 |
| VC2207 | 3516.3327 | 3694.665 | 4012.032 | 763.78363 | 783.8465 | 824.63406 | 1.47E-88 | 1.39E-86 | 3741.01 | 790.7547 | 0.211375 | -2.24213 | 3.235515 |
| VC2208 | 2408.06 | 2651.134 | 2570.919 | 1366.3925 | 1347.786 | 1323.4278 | 3.05E-22 | 5.62E-21 | 2543.371 | 1345.869 | 0.529167 | -0.9182 | 3.267206 |
| VC2209 | 3942.2694 | 4571.325 | 3642.135 | 1368.4457 | 1232.457 | 1159.0743 | 1.08E-28 | 2.66E-27 | 4051.91 | 1253.326 | 0.309317 | -1.69284 | 3.352862 |
| VC2210 | 2773.2981 | 3242.233 | 2172.164 | 874.65545 | 782.8691 | 763.47927 | 1.09E-18 | 1.74E-17 | 2729.232 | 807.0013 | 0.295688 | -1.75785 | 3.171457 |
| VC2211 | 1813.6323 | 2362.047 | 1456.854 | 678.57659 | 569.8036 | 577.14829 | 1.69E-12 | 1.93E-11 | 1877.511 | 608.5095 | 0.324104 | -1.62547 | 3.028925 |
| VC2212 | 1442.115 | 1889.638 | 1397.39 | 14544.74 | 12692.06 | 12169.802 | 4.97E-63 | 2.87E-61 | 1576.381 | 13135.53 | 8.332714 | 3.058787 | 3.658054 |
| VC2213 | 36695.443 | 34227.36 | 41688.23 | 252008.56 | 315774.8 | 271403.98 | 1.42E-76 | 1.03E-74 | 37537.01 | 279729.1 | 7.452088 | 2.897645 | 5.010599 |
| VC2232 | 1838.749 | 1776.824 | 1825.002 | 1409.5093 | 1420.111 | 1413.2489 | 1.11E-05 | 6.25E-05 | 1813.525 | 1414.29 | 0.779857 | -0.35872 | 3.204531 |
| VC2239 | 1773.8642 | 1714.541 | 1681.591 | 2513.0945 | 2965.324 | 3633.932 | 6.17E-06 | 3.63E-05 | 1723.332 | 3037.45 | 1.762545 | 0.81766 | 3.359439 |
| VC2240 | 603.84641 | 575.8225 | 765.1544 | 1751.3641 | 2495.212 | 2949.7629 | 1.49E-13 | 1.86E-12 | 648.2744 | 2398.78 | 3.700253 | 1.887624 | 3.095875 |
| VC2250 | 13368.343 | 16796.39 | 13054.85 | 25620.629 | 24957.98 | 25554.1 | 3.93E-08 | 3.04E-07 | 14406.53 | 25377.57 | 1.761533 | 0.816832 | 4.281505 |
| VC2251 | 5141.59 | 4311.618 | 5683.129 | 9230.0788 | 10900.55 | 9892.7421 | 1.29E-09 | 1.13E-08 | 5045.446 | 10007.79 | 1.983529 | 0.98807 | 3.851619 |
| VC2252 | 17425.731 | 20867.1 | 17174.44 | 38460.202 | 33964.4 | 35737.328 | 3.15E-12 | 3.52E-11 | 18489.09 | 36053.98 | 1.950014 | 0.963484 | 4.411934 |
| VC2253 | 5168.7997 | 7388.155 | 4675.749 | 12335.516 | 10469.53 | 10853.063 | 1.83E-05 | 0.000101 | 5744.234 | 11219.37 | 1.953153 | 0.965805 | 3.9046 |
| VC2264 | 604.89294 | 340.7929 | 679.4571 | 1751.3641 | 3224.326 | 3943.5281 | 4.16E-10 | 3.91E-09 | 541.7143 | 2973.073 | 5.488267 | 2.456351 | 3.103488 |
| VC2274 | 5030.6581 | 4799.304 | 4619.783 | 3631.052 | 3666.095 | 3472.4452 | 2.23E-06 | 1.39E-05 | 4816.582 | 3589.864 | 0.745314 | -0.42408 | 3.618908 |
| VC2275 | 317.09786 | 203.3006 | 320.0531 | 426.03522 | 673.4043 | 730.99079 | 0.000199 | 0.000905 | 280.1505 | 610.1434 | 2.177913 | 1.122946 | 2.616412 |
| VC2277 | 4712.5137 | 5359.85 | 4192.171 | 2692.7479 | 2652.568 | 2858.9862 | 1.91E-08 | 1.52E-07 | 4754.845 | 2734.767 | 0.575154 | -0.79798 | 3.557028 |
| VC2278 | 1094.6678 | 1418.404 | 937.4234 | 718.61363 | 608.8982 | 600.08133 | 7.93E-05 | 0.000395 | 1150.165 | 642.5311 | 0.558643 | -0.84 | 2.934327 |
| VC2283 | 692.80126 | 709.7893 | 657.5955 | 1295.5577 | 1198.249 | 1123.7192 | 5.85E-12 | 6.37E-11 | 686.7287 | 1205.842 | 1.755922 | 0.812229 | 2.959038 |
| VC2285 | 1101.9935 | 1084.662 | 1061.597 | 3884.62 | 3304.47 | 3164.7601 | 2.02E-36 | 5.88E-35 | 1082.751 | 3451.283 | 3.187514 | 1.672432 | 3.286255 |
| VC2286 | 772.33735 | 844.9313 | 834.2369 | 3845.6095 | 3293.719 | 3087.3611 | 8.93E-48 | 3.33E-46 | 817.1685 | 3408.897 | 4.171596 | 2.060599 | 3.222463 |
| VC2305 | 32518.751 | 21075.1 | 30399.8 | 3863.0616 | 8924.317 | 12844.416 | 0.000105 | 0.000507 | 27997.89 | 8543.931 | 0.305163 | -1.71235 | 4.189391 |
| VC2312 | 2865.3925 | 2805.078 | 2389.905 | 8025.8877 | 6888.466 | 6936.2904 | 6.64E-24 | 1.3E-22 | 2686.792 | 7283.548 | 2.710872 | 1.438757 | 3.645789 |
| VC2319 | 1422.231 | 1602.902 | 1641.365 | 2126.0698 | 2263.577 | 2228.3275 | 7.38E-06 | 4.3E-05 | 1555.499 | 2205.991 | 1.418188 | 0.504049 | 3.267737 |
| VC2323 | 843.50123 | 552.3195 | 696.0719 | 352.12068 | 364.557 | 383.17295 | 5.65E-06 | 3.33E-05 | 697.2975 | 366.6169 | 0.525768 | -0.9275 | 2.703815 |
| VC2336 | 155.93261 | 192.7243 | 197.6284 | 306.95068 | 305.9151 | 257.04121 | 0.000198 | 0.000901 | 182.0951 | 289.969 | 1.592404 | 0.671207 | 2.361325 |
| VC2339 | 187.32844 | 198.6 | 183.637 | 441.43409 | 504.3202 | 566.63731 | 1.14E-17 | 1.75E-16 | 189.8552 | 504.1305 | 2.655343 | 1.408898 | 2.490483 |
| VC2341 | 1128.1567 | 260.8828 | 2013.886 | 151.93545 | 219.9071 | 280.9298 | 0.000209 | 0.000948 | 1134.309 | 217.5908 | 0.191827 | -2.38212 | 2.696186 |
| VC2356 | 19170.292 | 23225.62 | 21496.9 | 9586.3058 | 10290.67 | 9825.854 | 1.83E-19 | 3.03E-18 | 21297.61 | 9900.944 | 0.464885 | -1.10505 | 4.162004 |
| VC2360 | 7893.9575 | 6339.923 | 8215.572 | 3791.2002 | 3807.813 | 3863.2625 | 4.63E-11 | 4.7E-10 | 7483.151 | 3820.758 | 0.510582 | -0.96979 | 3.728117 |
| VC2361 | 570.35753 | 722.716 | 781.7691 | 7414.0395 | 6941.244 | 5624.3292 | 1.13E-60 | 6.15E-59 | 691.6142 | 6659.871 | 9.629459 | 3.267455 | 3.331665 |
| VC2362 | 2682.2502 | 2816.829 | 2842.876 | 4812.6581 | 4711.875 | 4832.1836 | 1.09E-22 | 2.08E-21 | 2780.652 | 4785.572 | 1.721025 | 0.783268 | 3.56204 |
| VC2363 | 1725.724 | 2099.989 | 1985.029 | 3553.0311 | 3629.932 | 3363.5132 | 1.35E-11 | 1.42E-10 | 1936.914 | 3515.492 | 1.814996 | 0.859967 | 3.416548 |
| VC2364 | 3718.3125 | 4204.679 | 4263.877 | 7800.0377 | 7521.799 | 6895.2021 | 4.68E-13 | 5.56E-12 | 4062.29 | 7405.679 | 1.823031 | 0.866339 | 3.739168 |
| VC2366 | 885.36233 | 728.5917 | 908.5661 | 1644.5986 | 1662.497 | 1706.6007 | 5.12E-13 | 6.05E-12 | 840.8401 | 1671.232 | 1.987574 | 0.991009 | 3.073875 |
| VC2368 | 20949.389 | 21438.22 | 19739.23 | 14838.345 | 14477.7 | 14334.108 | 8.37E-10 | 7.56E-09 | 20708.95 | 14550.05 | 0.702597 | -0.50923 | 4.239511 |
| VC2371 | 116.16456 | 144.5432 | 113.6801 | 449.64682 | 298.0962 | 226.46381 | 7.62E-06 | 4.42E-05 | 124.7959 | 324.7356 | 2.602133 | 1.379695 | 2.303865 |
| VC2373 | 530.58948 | 1153.995 | 526.4262 | 6091.7904 | 5340.321 | 4194.8361 | 2.2E-14 | 2.9E-13 | 737.0036 | 5208.982 | 7.067784 | 2.821258 | 3.292111 |
| VC2374 | 377.79646 | 524.116 | 289.447 | 1793.4543 | 1876.54 | 1686.5343 | 4.6E-18 | 7.18E-17 | 397.1198 | 1785.51 | 4.496149 | 2.16869 | 2.925342 |
| VC2377 | 641.52141 | 562.8958 | 871.8387 | 1365.3659 | 1524.689 | 1497.3367 | 3.17E-07 | 2.21E-06 | 692.0853 | 1462.464 | 2.113126 | 1.079379 | 3.002622 |
| VC2397 | 27229.601 | 34264.96 | 28330.82 | 48889.339 | 47107.02 | 47641.489 | 8.73E-07 | 5.73E-06 | 29941.8 | 47879.28 | 1.599079 | 0.677241 | 4.578213 |
| VC2398 | 8823.274 | 11663.34 | 9682.045 | 18254.839 | 16460.78 | 17041.163 | 6.05E-07 | 4.03E-06 | 10056.22 | 17252.26 | 1.715581 | 0.778697 | 4.11964 |
| VC2400 | 7144.6438 | 9102.696 | 6450.033 | 24573.506 | 22610.35 | 22429.473 | 1.05E-19 | 1.76E-18 | 7565.791 | 23204.44 | 3.067022 | 1.616838 | 4.122213 |
| VC2401 | 2347.3614 | 3306.866 | 2340.061 | 9323.4986 | 8432.702 | 8251.1183 | 3.05E-18 | 4.88E-17 | 2664.763 | 8669.106 | 3.253238 | 1.701876 | 3.681816 |
| VC2402 | 1246.4143 | 1712.19 | 1293.329 | 5430.6659 | 4740.219 | 4535.0096 | 1.38E-21 | 2.48E-20 | 1417.311 | 4901.965 | 3.458636 | 1.790203 | 3.420918 |
| VC2403 | 3291.3292 | 4682.964 | 3453.251 | 11693.897 | 9899.727 | 10320.826 | 3.73E-14 | 4.83E-13 | 3809.182 | 10638.15 | 2.792765 | 1.481694 | 3.803849 |

| | | | | | | | | | | | | | |
|---|---|---|---|---|---|---|---|---|---|---|---|---|---|
| VC2404 | 3314.3528 | 4653.586 | 3600.161 | 16231.429 | 13428.01 | 13760.782 | 2.79E-23 | 5.39E-22 | 3856.033 | 14473.41 | 3.753445 | 1.908215 | 3.873356 |
| VC2405 | 5646.0163 | 6067.289 | 5982.195 | 23514.065 | 20383.92 | 19828.484 | 7.11E-53 | 3.16E-51 | 5898.5 | 21242.16 | 3.601281 | 1.84851 | 4.04897 |
| VC2406 | 6338.8175 | 6992.13 | 6958.97 | 22095.316 | 20677.13 | 19578.131 | 9.13E-51 | 3.91E-49 | 6763.306 | 20783.52 | 3.072983 | 1.61964 | 4.073939 |
| VC2407 | 2680.1571 | 3112.967 | 2654.867 | 12453.574 | 10503.74 | 9934.786 | 6.87E-37 | 2.04E-35 | 2815.997 | 10964.03 | 3.893482 | 1.961061 | 3.744801 |
| VC2408 | 355.81938 | 327.8663 | 303.4384 | 1150.8084 | 1042.848 | 959.3657 | 2.87E-30 | 7.44E-29 | 329.0413 | 1051.007 | 3.19415 | 1.675432 | 2.769428 |
| VC2409 | 3802.0347 | 4479.664 | 3562.559 | 15801.287 | 13553.12 | 12981.059 | 1.74E-30 | 4.6E-29 | 3948.086 | 14111.82 | 3.574345 | 1.837679 | 3.872985 |
| VC2418 | 4474.9519 | 4283.414 | 4418.657 | 5777.6536 | 5797.727 | 5988.3913 | 5.08E-08 | 3.88E-07 | 4392.341 | 5854.591 | 1.332909 | 0.414578 | 3.705096 |
| VC2423 | 94.187482 | 50.53136 | 89.19514 | 206.34477 | 189.6087 | 202.57523 | 3.91E-07 | 2.71E-06 | 77.97133 | 199.5096 | 2.558756 | 1.355442 | 2.095949 |
| VC2424 | 167.44441 | 173.9219 | 224.7368 | 432.19477 | 431.9952 | 428.0835 | 7.22E-12 | 7.81E-11 | 188.701 | 430.7578 | 2.282753 | 1.190775 | 2.455004 |
| VC2429 | 947.10746 | 997.7006 | 985.5188 | 551.27931 | 531.6864 | 605.81459 | 3.49E-12 | 3.89E-11 | 976.7756 | 562.9268 | 0.576311 | -0.79508 | 2.870123 |
| VC2430 | 20059.841 | 19724.86 | 17967.57 | 30610.888 | 36564.19 | 38646.958 | 1.25E-09 | 1.1E-08 | 19250.76 | 35274.01 | 1.832344 | 0.87369 | 4.415951 |
| VC2431 | 13470.903 | 14098.25 | 12973.52 | 20252.585 | 25347.95 | 24452.359 | 1.22E-08 | 9.85E-08 | 13514.22 | 23350.97 | 1.727881 | 0.789004 | 4.249548 |
| VC2432 | 885.36233 | 1236.256 | 1095.701 | 2965.8211 | 3789.243 | 3696.0423 | 6.76E-18 | 1.05E-16 | 1072.44 | 3483.702 | 3.24839 | 1.699725 | 3.286207 |
| VC2433 | 1704.7934 | 2399.652 | 2278.848 | 6444.9377 | 7696.747 | 7381.5737 | 2.16E-20 | 3.69E-19 | 2127.765 | 7174.419 | 3.371811 | 1.753524 | 3.591855 |
| VC2434 | 757.68597 | 1195.125 | 1222.498 | 2645.5248 | 2756.168 | 2291.3934 | 7.9E-08 | 5.89E-07 | 1058.436 | 2564.362 | 2.422783 | 1.276665 | 3.216822 |
| VC2435 | 964.89843 | 1530.043 | 1679.842 | 3550.9779 | 3650.457 | 2810.2535 | 1.72E-06 | 1.08E-05 | 1391.594 | 3337.229 | 2.398134 | 1.261912 | 3.333449 |
| VC2443 | 3482.8438 | 4491.415 | 3229.389 | 11410.558 | 11329.61 | 12467.932 | 7.02E-21 | 1.21E-19 | 3734.549 | 11736.03 | 3.142557 | 1.651939 | 3.82088 |
| VC2450 | 2376.6641 | 1914.316 | 2358.424 | 5817.6906 | 5565.114 | 5491.5086 | 1.9E-22 | 3.57E-21 | 2216.468 | 5624.771 | 2.537718 | 1.343532 | 3.547883 |
| VC2451 | 7013.8278 | 8086.193 | 7445.171 | 24334.311 | 23366.83 | 21959.346 | 1.22E-47 | 4.51E-46 | 7515.064 | 23220.16 | 3.089816 | 1.627521 | 4.120899 |
| VC2452 | 2715.7391 | 3216.38 | 2465.983 | 7507.4593 | 7087.849 | 6633.3831 | 3.47E-18 | 5.49E-17 | 2799.367 | 7076.23 | 2.527796 | 1.33788 | 3.648431 |
| VC2458 | 3869.0125 | 3505.466 | 3955.192 | 2805.6729 | 2835.335 | 2616.2782 | 1.99E-05 | 0.000109 | 3776.557 | 2752.429 | 0.72882 | -0.45637 | 3.508406 |
| VC2464 | 1499.674 | 1801.502 | 1594.144 | 3004.8316 | 2917.434 | 2978.4292 | 3.88E-13 | 4.65E-12 | 1631.773 | 2966.898 | 1.818205 | 0.862515 | 3.342481 |
| VC2465 | 7779.886 | 8578.58 | 8218.195 | 14436.948 | 12911.96 | 12883.593 | 4.61E-11 | 4.7E-10 | 8192.22 | 13410.84 | 1.637021 | 0.711073 | 4.020429 |
| VC2466 | 12915.197 | 13666.97 | 12219.73 | 24137.205 | 23235.87 | 24467.647 | 2.64E-21 | 4.68E-20 | 12933.97 | 23946.91 | 1.851474 | 0.888674 | 4.245491 |
| VC2467 | 13953.352 | 14920.85 | 13144.91 | 22520.325 | 20102.44 | 21041.068 | 5.34E-08 | 4.06E-07 | 14006.37 | 21221.28 | 1.515116 | 0.599428 | 4.236549 |
| VC2473 | 1019.3179 | 1152.82 | 1018.748 | 5055.9602 | 5316.864 | 5404.5542 | 3.4E-99 | 3.83E-97 | 1063.629 | 5259.126 | 4.944513 | 2.305828 | 3.373852 |
| VC2476 | 3280.864 | 3091.814 | 3127.077 | 2364.2388 | 2328.083 | 2215.9054 | 1.96E-07 | 1.4E-06 | 3166.585 | 2302.742 | 0.727201 | -0.45957 | 3.431418 |
| VC2479 | 933.5026 | 981.2485 | 828.9901 | 536.90704 | 569.8036 | 476.81622 | 4.8E-08 | 3.68E-07 | 914.5804 | 527.8423 | 0.577141 | -0.793 | 2.841863 |
| VC2481 | 1377.2303 | 1296.188 | 1762.916 | 3569.4566 | 3303.493 | 2932.5631 | 4.48E-10 | 4.19E-09 | 1478.778 | 3268.504 | 2.210274 | 1.144225 | 3.342126 |
| VC2490 | 1535.256 | 1542.969 | 1971.912 | 2487.4298 | 2625.202 | 2449.058 | 0.000187 | 0.000852 | 1683.379 | 2520.563 | 1.497324 | 0.582386 | 3.31384 |
| VC2512 | 8798.1573 | 8289.493 | 9576.235 | 5900.8445 | 6197.469 | 6401.1861 | 1.47E-06 | 9.37E-06 | 8887.962 | 6166.5 | 0.693804 | -0.5274 | 3.86942 |
| VC2513 | 1622.1177 | 2438.432 | 1363.286 | 6220.1143 | 5370.619 | 5985.5246 | 4.63E-11 | 4.7E-10 | 1807.945 | 5858.753 | 3.240558 | 1.696242 | 3.512495 |
| VC2514 | 17345.148 | 22883.65 | 17358.95 | 81191.021 | 64784.62 | 67206.243 | 2E-23 | 3.9E-22 | 19195.92 | 71060.63 | 3.701862 | 1.888251 | 4.567419 |
| VC2515 | 2606.9002 | 3447.884 | 2563.923 | 8112.1213 | 7081.007 | 6696.449 | 3.57E-13 | 4.33E-12 | 2872.902 | 7296.526 | 2.539775 | 1.344701 | 3.660718 |
| VC2516 | 1354.2067 | 1483.037 | 1593.27 | 10398.339 | 9721.846 | 8797.6892 | 8.15E-91 | 7.91E-89 | 1476.838 | 9639.292 | 6.526981 | 2.706416 | 3.576689 |
| VC2517 | 2280.3836 | 2568.873 | 2614.642 | 26011.76 | 24239.62 | 21611.528 | 2.2E-130 | 3.6E-128 | 2487.966 | 23954.3 | 9.628066 | 3.267246 | 3.887614 |
| VC2518 | 1680.7233 | 1705.14 | 1780.405 | 10586.205 | 9177.454 | 8766.1563 | 2.07E-85 | 1.78E-83 | 1722.089 | 9509.939 | 5.522326 | 2.465276 | 3.607117 |
| VC2519 | 1453.6268 | 1696.914 | 1489.209 | 7014.6956 | 6309.866 | 5994.1245 | 7.96E-54 | 3.67E-52 | 1546.583 | 6439.562 | 4.163735 | 2.057878 | 3.499115 |
| VC2520 | 2064.7989 | 2333.844 | 2105.705 | 9761.8529 | 8995.664 | 8346.6727 | 3.25E-62 | 1.85E-60 | 2168.116 | 9034.73 | 4.167088 | 2.05904 | 3.645999 |
| VC2535 | 8701.8768 | 9067.441 | 8390.464 | 6976.7118 | 7073.188 | 7023.2449 | 9.73E-05 | 0.000476 | 8719.927 | 7024.382 | 0.805555 | -0.31194 | 3.89356 |
| VC2540 | 418.61103 | 504.1384 | 395.2569 | 812.0334 | 923.6096 | 954.58798 | 2.1E-10 | 2.04E-09 | 439.3355 | 896.7437 | 2.041137 | 1.029373 | 2.797732 |
| VC2541 | 715.82486 | 667.484 | 857.8473 | 1741.0982 | 2380.86 | 2560.8566 | 2.76E-13 | 3.38E-12 | 747.0521 | 2227.605 | 2.98186 | 1.576213 | 3.110595 |
| VC2542 | 3865.8729 | 3469.037 | 3897.478 | 11309.952 | 15683.77 | 17206.472 | 4.58E-22 | 8.36E-21 | 3744.129 | 14733.4 | 3.935067 | 1.976388 | 3.870827 |
| VC2545 | 27258.904 | 33729.09 | 24941.41 | 13476.059 | 13433.88 | 14135.355 | 3.24E-11 | 3.33E-10 | 28643.14 | 13681.76 | 0.477663 | -1.06594 | 4.296581 |
| VC2546 | 659.31237 | 565.2461 | 746.7907 | 2896.0129 | 3002.464 | 3066.3392 | 2.55E-48 | 9.6E-47 | 657.1164 | 2988.272 | 4.547554 | 2.185091 | 3.146531 |
| VC2547 | 4309.6006 | 4415.031 | 4820.91 | 30763.85 | 28296.66 | 26754.263 | 1.1E-117 | 1.6E-115 | 4515.18 | 28604.92 | 6.335279 | 2.663408 | 4.055558 |
| VC2548 | 5301.7087 | 5003.78 | 4909.23 | 32215.449 | 28096.3 | 26929.128 | 1.59E-93 | 1.58E-91 | 5071.573 | 29080.29 | 5.733979 | 2.519537 | 4.084371 |
| VC2552 | 873.85053 | 732.1171 | 909.4406 | 510.21568 | 569.8036 | 449.10546 | 2.31E-05 | 0.000126 | 838.4694 | 509.7082 | 0.607903 | -0.71809 | 2.815404 |
| VC2553 | 406.0527 | 344.3183 | 435.4821 | 225.85 | 189.6087 | 182.50881 | 3.27E-08 | 2.54E-07 | 395.2844 | 199.3225 | 0.504251 | -0.98779 | 2.448233 |

| | | | | | | | | | | | | | |
|---|---|---|---|---|---|---|---|---|---|---|---|---|---|
| VC2560 | 58.605544 | 77.55976 | 60.33789 | 107.79204 | 113.3743 | 127.08729 | 0.000112 | 0.000537 | 65.50106 | 116.0845 | 1.772254 | 0.825586 | 1.940511 |
| VC2567 | 1364.672 | 1850.858 | 1214.628 | 3339.5002 | 3403.184 | 3095.0055 | 1.72E-08 | 1.37E-07 | 1476.719 | 3279.23 | 2.220618 | 1.150961 | 3.342535 |
| VC2599 | 19380.644 | 16126.55 | 21876.42 | 11845.832 | 13043.91 | 11793.318 | 7.7E-05 | 0.000385 | 19127.87 | 12227.69 | 0.63926 | -0.64552 | 4.184505 |
| VC2604 | 28125.429 | 30194.25 | 28948.19 | 22805.717 | 21589.01 | 22390.296 | 2.57E-06 | 1.58E-05 | 29089.29 | 22261.67 | 0.765288 | -0.38593 | 4.405645 |
| VC2607 | 1880.6101 | 1432.505 | 1625.625 | 944.46363 | 972.4778 | 942.16591 | 1.56E-07 | 1.13E-06 | 1646.247 | 953.0358 | 0.578914 | -0.78858 | 3.097802 |
| VC2608 | 3481.7973 | 3813.355 | 3376.298 | 5485.0752 | 5378.438 | 4905.7604 | 5.69E-07 | 3.84E-06 | 3557.15 | 5256.424 | 1.477707 | 0.56336 | 3.635896 |
| VC2609 | 1199.3206 | 956.5704 | 999.5102 | 5891.6052 | 5718.561 | 5617.6404 | 8.19E-72 | 5.49E-70 | 1051.8 | 5742.602 | 5.459783 | 2.448844 | 3.390521 |
| VC2610 | 1605.3733 | 1879.061 | 1596.768 | 5688.3402 | 5382.347 | 5357.7325 | 1.18E-44 | 4.1E-43 | 1693.734 | 5476.14 | 3.233176 | 1.692952 | 3.48366 |
| VC2612 | 135.00206 | 133.9669 | 113.6801 | 314.13682 | 305.9151 | 320.10708 | 3.7E-17 | 5.52E-16 | 127.5497 | 313.3863 | 2.456975 | 1.296883 | 2.30088 |
| VC2613 | 555.70614 | 507.6639 | 606.0023 | 1358.1798 | 1282.302 | 1321.5167 | 3.73E-24 | 7.5E-23 | 556.4574 | 1320.666 | 2.373347 | 1.246923 | 2.933113 |
| VC2633 | 52.326379 | 49.35621 | 53.34219 | 17.452045 | 25.41148 | 23.888588 | 2.89E-05 | 0.000155 | 51.67493 | 22.2507 | 0.43059 | -1.21561 | 1.530312 |
| VC2634 | 86.861789 | 70.50887 | 73.45482 | 22.585 | 28.34358 | 27.710762 | 4.95E-09 | 4.1E-08 | 76.94183 | 26.21311 | 0.340687 | -1.55348 | 1.652341 |
| VC2635 | 7688.8381 | 9968.78 | 7581.587 | 48195.363 | 40459.97 | 39285.261 | 8.12E-38 | 2.44E-36 | 8413.068 | 42646.86 | 5.069121 | 2.341736 | 4.277421 |
| VC2636 | 2370.385 | 2282.137 | 2270.104 | 1375.6318 | 1418.156 | 1327.25 | 3.45E-17 | 5.19E-16 | 2307.542 | 1373.679 | 0.5953 | -0.74831 | 3.250517 |
| VC2638 | 990.01509 | 1021.204 | 997.7613 | 703.21477 | 746.7066 | 765.39036 | 1.04E-05 | 5.92E-05 | 1002.993 | 738.4372 | 0.736233 | -0.44176 | 2.934806 |
| VC2651 | 4843.3296 | 4671.213 | 5764.454 | 3249.1602 | 3655.344 | 3670.2427 | 0.000222 | 0.000999 | 5092.999 | 3524.916 | 0.69211 | -0.53093 | 3.627061 |
| VC2662 | 16605.253 | 20459.32 | 15383.54 | 100233.26 | 86003.2 | 85729.454 | 3.89E-44 | 1.34E-42 | 17482.71 | 90655.3 | 5.185428 | 2.374463 | 4.600001 |
| VC2666 | 861.2922 | 968.3219 | 902.4449 | 601.58227 | 655.8117 | 610.59231 | 2.23E-06 | 1.39E-05 | 910.6863 | 622.6621 | 0.683728 | -0.5485 | 2.876811 |
| VC2667 | 114.07151 | 104.5882 | 90.94406 | 306.95068 | 454.4746 | 455.79426 | 1.55E-16 | 2.25E-15 | 103.2012 | 405.7398 | 3.93154 | 1.975095 | 2.310966 |
| VC2681 | 31767.345 | 29143.67 | 34678.54 | 12805.695 | 13713.4 | 14489.862 | 1.52E-22 | 2.88E-21 | 31863.19 | 13669.65 | 0.429011 | -1.22091 | 4.319523 |
| VC2686 | 6374.3995 | 6927.497 | 5317.604 | 2867.2684 | 2762.033 | 2774.8984 | 4.34E-15 | 5.95E-14 | 6206.5 | 2801.4 | 0.451365 | -1.14763 | 3.620111 |
| VC2691 | 106.74581 | 82.26035 | 122.4247 | 505.08272 | 479.8861 | 484.46057 | 2.1E-32 | 5.74E-31 | 103.8103 | 489.8098 | 4.718316 | 2.238272 | 2.353134 |
| VC2692 | 1325.9504 | 1412.528 | 1253.979 | 883.89477 | 1048.712 | 921.14396 | 0.000219 | 0.000989 | 1330.819 | 951.2503 | 0.714786 | -0.48442 | 3.051207 |
| VC2694 | 1556.1865 | 2479.562 | 1337.927 | 610.82159 | 517.0259 | 463.43861 | 1.07E-09 | 9.44E-09 | 1791.225 | 530.4287 | 0.296126 | -1.75572 | 2.988889 |
| VC2697 | 847.68734 | 420.7029 | 846.4793 | 1631.2529 | 1929.318 | 1966.5086 | 7.51E-06 | 4.36E-05 | 704.9565 | 1842.36 | 2.613437 | 1.385949 | 3.056768 |
| VC2703 | 2041.7753 | 2394.951 | 2012.137 | 15279.779 | 14946.84 | 14606.438 | 1.9E-119 | 2.6E-117 | 2149.621 | 14944.35 | 6.952086 | 2.797446 | 3.75342 |
| VC2714 | 3729.8243 | 2792.151 | 3754.066 | 6361.7838 | 5787.954 | 5712.2392 | 3.63E-06 | 2.17E-05 | 3425.347 | 5953.992 | 1.738216 | 0.797607 | 3.654756 |
| VC2715 | 1047.5741 | 1028.254 | 988.1422 | 2981.22 | 2560.695 | 2749.0987 | 3.88E-36 | 1.12E-34 | 1021.324 | 2763.671 | 2.70597 | 1.436146 | 3.225325 |
| VC2716 | 9332.9329 | 9039.238 | 10198.85 | 37832.955 | 38456.37 | 37491.706 | 1.97E-97 | 2.07E-95 | 9523.674 | 37927.01 | 3.982393 | 1.993635 | 4.278877 |
| VC2720 | 6754.289 | 6187.154 | 6517.366 | 10368.568 | 8784.554 | 9296.4829 | 3.27E-06 | 1.97E-05 | 6486.27 | 9483.202 | 1.462042 | 0.547985 | 3.894475 |
| VC2723 | 1285.1359 | 1421.929 | 1469.971 | 2140.442 | 2361.313 | 2303.8154 | 9.22E-10 | 8.25E-09 | 1392.345 | 2268.524 | 1.629282 | 0.704237 | 3.249745 |
| VC2724 | 984.78245 | 1192.775 | 1170.905 | 1630.2264 | 1904.884 | 1802.1551 | 3.21E-06 | 1.94E-05 | 1116.154 | 1779.088 | 1.593945 | 0.672602 | 3.148961 |
| VC2725 | 3099.8147 | 3493.715 | 3513.589 | 5537.4313 | 5964.857 | 5923.4143 | 4.83E-13 | 5.72E-12 | 3369.039 | 5808.567 | 1.724102 | 0.785845 | 3.645788 |
| VC2726 | 788.03527 | 908.3893 | 982.8954 | 1517.3014 | 1627.312 | 1552.7582 | 2.54E-09 | 2.16E-08 | 893.1067 | 1565.791 | 1.753195 | 0.809987 | 3.072819 |
| VC2727 | 681.28945 | 715.6651 | 838.6092 | 1500.8759 | 1526.644 | 1389.3603 | 1.66E-12 | 1.9E-11 | 745.1879 | 1472.293 | 1.975734 | 0.982389 | 3.02013 |
| VC2728 | 432.21589 | 408.9515 | 429.3609 | 888.00113 | 885.4924 | 807.43428 | 4.99E-19 | 8.11E-18 | 423.5094 | 860.3093 | 2.031382 | 1.022461 | 2.780759 |
| VC2729 | 1371.9977 | 1644.032 | 1536.43 | 2888.8268 | 2986.826 | 2988.9401 | 3.07E-16 | 4.44E-15 | 1517.487 | 2954.864 | 1.94721 | 0.961408 | 3.325831 |
| VC2730 | 2184.1031 | 1987.175 | 2101.332 | 3512.9941 | 4092.226 | 4421.2999 | 1.97E-11 | 2.07E-10 | 2090.87 | 4008.84 | 1.917307 | 0.939081 | 3.461673 |
| VC2731 | 2826.671 | 2961.373 | 2894.47 | 5033.3752 | 5552.409 | 5958.7694 | 3.21E-16 | 4.62E-15 | 2894.171 | 5514.851 | 1.905503 | 0.930172 | 3.601529 |
| VC2732 | 3656.5674 | 3973.175 | 4054.006 | 6212.9281 | 6940.267 | 7353.863 | 3.23E-11 | 3.33E-10 | 3894.583 | 6835.686 | 1.755178 | 0.811617 | 3.712622 |
| VC2733 | 7055.6889 | 7025.034 | 7406.694 | 11309.952 | 13233.52 | 14822.391 | 2.87E-09 | 2.43E-08 | 7162.472 | 13121.95 | 1.832042 | 0.873453 | 3.986531 |
| VC2734 | 1797.9344 | 2190.476 | 1896.708 | 3486.3027 | 3936.825 | 4138.459 | 2.26E-11 | 2.35E-10 | 1961.706 | 3853.862 | 1.964546 | 0.974196 | 3.439265 |
| VC2735 | 2685.3898 | 1879.061 | 4194.795 | 1115.9043 | 1310.646 | 1121.8081 | 4.62E-05 | 0.00024 | 2919.749 | 1182.786 | 0.405099 | -1.30365 | 3.269126 |
| VC2736 | 15003.019 | 13444.87 | 22520.9 | 9190.0418 | 9653.431 | 8455.6046 | 0.000187 | 0.000852 | 16989.59 | 9099.692 | 0.535604 | -0.90076 | 4.094605 |
| VC2742 | 659.31237 | 598.1503 | 605.1278 | 465.04568 | 449.5877 | 429.99459 | 6.35E-05 | 0.000325 | 620.8635 | 448.2093 | 0.721913 | -0.4701 | 2.722239 |
| VC2755 | 1116.6449 | 1078.786 | 1366.784 | 641.61931 | 757.4576 | 765.39036 | 1.66E-05 | 9.21E-05 | 1187.405 | 721.4891 | 0.607618 | -0.71876 | 2.966414 |
| VC2756 | 928.26996 | 1002.401 | 968.904 | 554.35909 | 706.6347 | 624.92546 | 1.75E-05 | 9.7E-05 | 966.525 | 628.6397 | 0.650412 | -0.62057 | 2.891807 |
| VC2757 | 848.73387 | 735.6426 | 890.2024 | 418.84909 | 388.9911 | 391.77284 | 3.82E-14 | 4.92E-13 | 824.8596 | 399.871 | 0.484775 | -1.04461 | 2.75915 |
| VC2761 | 17598.408 | 8840.638 | 25933.05 | 4089.9382 | 5722.47 | 5986.4802 | 2.04E-05 | 0.000111 | 17457.36 | 5266.296 | 0.301666 | -1.72898 | 3.981742 |

| | | | | | | | | | | | | | |
|---|---|---|---|---|---|---|---|---|---|---|---|---|---|
| VC2771 | 21603.469 | 29023.8 | 22166.74 | 36526.104 | 38247.21 | 44720.392 | 0.000117 | 0.000562 | 24264.67 | 39831.24 | 1.641532 | 0.715043 | 4.492599 |
| VCA0017 | 84.768734 | 77.55976 | 116.3035 | 242.27545 | 221.8618 | 222.64164 | 1.65E-09 | 1.43E-08 | 92.87732 | 228.9263 | 2.464824 | 1.301485 | 2.163803 |
| VCA0026 | 3504.8209 | 4304.567 | 3193.536 | 24352.789 | 21026.05 | 22101.722 | 2.28E-54 | 1.07E-52 | 3667.641 | 22493.52 | 6.132966 | 2.616585 | 3.958222 |
| VCA0027 | 433.26242 | 403.0757 | 422.3652 | 1383.8445 | 1514.915 | 1386.4937 | 2.2E-55 | 1.08E-53 | 419.5678 | 1428.418 | 3.404498 | 1.767442 | 2.888829 |
| VCA0029 | 1330.1366 | 1242.131 | 1287.208 | 1015.2984 | 964.6589 | 1002.3652 | 8.99E-05 | 0.000444 | 1286.492 | 994.1075 | 0.772727 | -0.37197 | 3.05342 |
| VCA0033 | 280.46939 | 303.1882 | 304.3128 | 530.7475 | 479.8861 | 466.30524 | 2.9E-08 | 2.26E-07 | 295.9901 | 492.3129 | 1.663275 | 0.734027 | 2.581759 |
| VCA0035 | 636.28877 | 855.5077 | 659.3444 | 22118.928 | 20199.2 | 20023.415 | 1.9E-183 | 7.1E-181 | 717.047 | 20780.51 | 28.98069 | 4.85702 | 3.586602 |
| VCA0036 | 3583.3104 | 5254.086 | 3814.404 | 630.32681 | 665.5853 | 703.28003 | 9.09E-40 | 2.86E-38 | 4217.267 | 666.3974 | 0.158016 | -2.66185 | 3.224382 |
| VCA0058 | 1353.1602 | 1387.85 | 1477.841 | 5644.1968 | 4908.325 | 4888.5607 | 1.11E-54 | 5.24E-53 | 1406.284 | 5147.028 | 3.660021 | 1.871852 | 3.429815 |
| VCA0059 | 35471.006 | 40379.26 | 37464.58 | 114664.04 | 111153.7 | 109923.82 | 1.49E-57 | 7.63E-56 | 37771.61 | 111913.9 | 2.962909 | 1.567014 | 4.813025 |
| VCA0064 | 3065.2793 | 6270.589 | 1773.409 | 912.63931 | 677.3137 | 577.14829 | 4.7E-07 | 3.24E-06 | 3703.093 | 722.3671 | 0.195071 | -2.35793 | 3.213661 |
| VCA0065 | 2247.9412 | 4383.302 | 1430.62 | 669.33727 | 487.705 | 469.17187 | 6.61E-08 | 4.98E-07 | 2687.288 | 542.0714 | 0.201717 | -2.3096 | 3.081685 |
| VCA0066 | 1290.3685 | 2452.534 | 871.8387 | 376.75886 | 291.2547 | 233.15262 | 1.67E-08 | 1.33E-07 | 1538.247 | 300.3887 | 0.19528 | -2.35638 | 2.832355 |
| VCA0067 | 662.45196 | 1115.215 | 462.5905 | 246.38182 | 209.156 | 164.35349 | 5.06E-07 | 3.45E-06 | 746.7526 | 206.6304 | 0.276705 | -1.85358 | 2.594186 |
| VCA0070 | 61.745127 | 74.03432 | 60.33789 | 153.98864 | 142.6952 | 156.70914 | 3.13E-10 | 2.97E-09 | 65.37244 | 151.131 | 2.311846 | 1.209045 | 1.997374 |
| VCA0071 | 70.117348 | 63.45799 | 74.32928 | 117.03136 | 124.1253 | 128.04283 | 1.64E-05 | 9.14E-05 | 69.30154 | 123.0665 | 1.775812 | 0.828479 | 1.965441 |
| VCA0074 | 270.00412 | 314.9396 | 301.6894 | 475.31159 | 412.4479 | 430.95013 | 6.52E-05 | 0.000331 | 295.5444 | 439.5699 | 1.487323 | 0.572718 | 2.556825 |
| VCA0078 | 102.5597 | 96.36213 | 112.8056 | 262.80727 | 269.7526 | 301.95175 | 3.58E-17 | 5.37E-16 | 103.9091 | 278.1706 | 2.677056 | 1.420647 | 2.230482 |
| VCA0079 | 15819.311 | 19575.61 | 16136.45 | 54113.66 | 49771.32 | 50227.19 | 6.35E-30 | 1.62E-28 | 17177.12 | 51370.72 | 2.990647 | 1.580458 | 4.472833 |
| VCA0095 | 64.88471 | 16.45207 | 61.21235 | 516.37522 | 1517.847 | 2434.7249 | 2.06E-14 | 2.73E-13 | 47.51638 | 1489.649 | 31.35023 | 4.970404 | 2.424964 |
| VCA0096 | 47.093741 | 48.18106 | 70.83143 | 662.15113 | 1762.189 | 2514.035 | 2.01E-23 | 3.9E-22 | 55.36875 | 1646.125 | 29.73022 | 4.893858 | 2.479864 |
| VCA0098 | 4980.4247 | 5390.403 | 4486.865 | 2710.2 | 2501.076 | 2408.9252 | 7.81E-14 | 9.9E-13 | 4952.564 | 2540.067 | 0.512879 | -0.96331 | 3.549838 |
| VCA0099 | 3644.009 | 3496.065 | 3651.754 | 2141.4686 | 2098.402 | 1990.3972 | 1.29E-19 | 2.16E-18 | 3597.276 | 2076.756 | 0.577313 | -0.79257 | 3.43668 |
| VCA0106 | 185.23538 | 204.4757 | 243.1005 | 395.2375 | 340.1229 | 306.72947 | 0.000204 | 0.000923 | 210.9372 | 347.3633 | 1.646762 | 0.719632 | 2.432469 |
| VCA0107 | 279.42286 | 303.1882 | 317.4298 | 467.09886 | 763.3218 | 852.34482 | 6.18E-06 | 3.63E-05 | 300.0136 | 694.2552 | 2.314079 | 1.210438 | 2.65933 |
| VCA0108 | 1035.0158 | 1043.531 | 1026.619 | 1862.2359 | 2383.792 | 2650.6777 | 8.25E-11 | 8.31E-10 | 1035.055 | 2298.902 | 2.221043 | 1.151237 | 3.188242 |
| VCA0109 | 156.97914 | 193.8994 | 195.0051 | 380.86522 | 440.7915 | 487.3272 | 6.96E-12 | 7.54E-11 | 181.9612 | 436.328 | 2.397918 | 1.261782 | 2.449896 |
| VCA0110 | 307.67911 | 323.1657 | 267.5854 | 537.93363 | 570.781 | 669.83601 | 2.71E-09 | 2.3E-08 | 299.4767 | 592.8502 | 1.97962 | 0.985224 | 2.624654 |
| VCA0112 | 276.28328 | 299.6627 | 341.9147 | 458.88613 | 471.0898 | 476.81622 | 1.76E-05 | 9.72E-05 | 305.9536 | 468.9307 | 1.532686 | 0.616062 | 2.578382 |
| VCA0114 | 311.86522 | 325.516 | 376.8932 | 508.1625 | 577.6225 | 542.74872 | 3.46E-06 | 2.08E-05 | 338.0915 | 542.8446 | 1.605615 | 0.683126 | 2.631855 |
| VCA0115 | 169.53747 | 206.826 | 189.7583 | 296.68477 | 262.9111 | 302.9073 | 0.000151 | 0.000704 | 188.7073 | 287.5011 | 1.523529 | 0.607417 | 2.367214 |
| VCA0117 | 371.51729 | 441.8556 | 389.1356 | 1102.5586 | 1073.146 | 1023.3871 | 1.85E-28 | 4.46E-27 | 400.8362 | 1066.364 | 2.660349 | 1.411615 | 2.815436 |
| VCA0118 | 73.25693 | 71.68402 | 74.32928 | 155.01523 | 156.3783 | 116.57631 | 7.07E-06 | 4.13E-05 | 73.09008 | 142.6566 | 1.951792 | 0.964799 | 2.009075 |
| VCA0120 | 949.20051 | 1040.006 | 985.5188 | 1707.2207 | 1790.532 | 1765.8444 | 1.95E-17 | 2.96E-16 | 991.5751 | 1754.532 | 1.76944 | 0.823293 | 3.120243 |
| VCA0121 | 284.6555 | 290.2615 | 309.5596 | 441.43409 | 486.7276 | 517.90459 | 2.35E-07 | 1.67E-06 | 294.8255 | 482.0221 | 1.63494 | 0.709238 | 2.576316 |
| VCA0125 | 567.21795 | 712.1396 | 581.5173 | 9169.5099 | 8079.874 | 7299.397 | 1.4E-106 | 1.8E-104 | 620.2916 | 8182.927 | 13.19206 | 3.721598 | 3.352752 |
| VCA0137 | 5020.1928 | 1360.821 | 7013.186 | 282.3125 | 844.4431 | 1221.1846 | 0.000179 | 0.000819 | 4464.733 | 782.6467 | 0.175295 | -2.51214 | 3.271681 |
| VCA0139 | 556.75267 | 538.2177 | 490.5733 | 99681.977 | 89447.44 | 94959.049 | 0 | 0 | 528.5146 | 94696.15 | 179.1742 | 7.485219 | 3.849695 |
| VCA0140 | 283.60897 | 257.3574 | 355.0316 | 7310.3538 | 9164.749 | 9826.8096 | 1.2E-122 | 1.8E-120 | 298.666 | 8767.304 | 29.35488 | 4.875528 | 3.209026 |
| VCA0159 | 47.093741 | 38.77988 | 43.72311 | 233.03613 | 207.2013 | 208.30849 | 2.02E-30 | 5.27E-29 | 43.19891 | 216.182 | 5.004339 | 2.323179 | 1.985146 |
| VCA0163 | 208.25899 | 186.8485 | 235.2303 | 111.89841 | 115.329 | 135.68718 | 2.02E-05 | 0.000111 | 210.1126 | 120.9715 | 0.575746 | -0.7965 | 2.202568 |
| VCA0167 | 3634.5903 | 3075.362 | 4146.699 | 8775.299 | 8426.838 | 8435.5382 | 1.4E-15 | 1.97E-14 | 3618.884 | 8545.892 | 2.361472 | 1.239686 | 3.745166 |
| VCA0176 | 4433.0908 | 3257.51 | 4727.342 | 1151.835 | 1538.372 | 1658.8236 | 9.79E-11 | 9.81E-10 | 4139.314 | 1449.677 | 0.350221 | -1.51366 | 3.3891 |
| VCA0177 | 937.68871 | 1077.611 | 904.1938 | 729.90613 | 653.857 | 644.03633 | 0.00015 | 0.000703 | 973.1644 | 675.9331 | 0.694572 | -0.5258 | 2.909045 |
| VCA0178 | 225.00343 | 188.0237 | 146.9096 | 389.07795 | 350.8739 | 364.06208 | 1.99E-06 | 1.25E-05 | 186.6456 | 368.0047 | 1.971676 | 0.979423 | 2.418436 |
| VCA0180 | 187.32844 | 206.826 | 204.6241 | 312.08363 | 318.6209 | 279.97425 | 3.29E-05 | 0.000175 | 199.5929 | 303.5596 | 1.520894 | 0.60492 | 2.391194 |
| VCA0189 | 1632.583 | 1525.342 | 1524.187 | 1189.8189 | 1218.774 | 1241.251 | 0.000123 | 0.000584 | 1560.704 | 1216.615 | 0.779529 | -0.35932 | 3.139237 |
| VCA0195 | 69.07082 | 41.13018 | 55.96558 | 274.09977 | 288.3226 | 232.19708 | 8.2E-21 | 1.41E-19 | 55.38886 | 264.8731 | 4.782065 | 2.257634 | 2.08323 |
| VCA0208 | 765.01166 | 656.9077 | 620.8681 | 383.945 | 388.0138 | 372.66197 | 4.76E-09 | 3.96E-08 | 680.9291 | 381.5402 | 0.560323 | -0.83567 | 2.707321 |

| | | | | | | | | | | | | | |
|---|---|---|---|---|---|---|---|---|---|---|---|---|---|
| VCA0214 | 506.51935 | 551.1444 | 548.2878 | 824.35249 | 826.8505 | 769.21254 | 1.17E-07 | 8.62E-07 | 535.3172 | 806.8052 | 1.507154 | 0.591826 | 2.81769 |
| VCA0216 | 283.60897 | 327.8663 | 169.6457 | 57.489091 | 62.55134 | 73.576851 | 1.22E-10 | 1.21E-09 | 260.3736 | 64.53909 | 0.247871 | -2.01234 | 2.11271 |
| VCA0225 | 3125.9779 | 2966.073 | 2702.088 | 5519.9793 | 4842.842 | 5097.8247 | 1.77E-11 | 1.86E-10 | 2931.38 | 5153.549 | 1.758062 | 0.813986 | 3.589589 |
| VCA0227 | 14176.263 | 19240.7 | 11703.8 | 5442.985 | 3767.741 | 3613.8656 | 4.76E-11 | 4.82E-10 | 15040.25 | 4274.864 | 0.284228 | -1.81488 | 3.904089 |
| VCA0228 | 1974.7975 | 2785.101 | 1410.507 | 674.47022 | 470.1124 | 467.26078 | 6.63E-10 | 6.08E-09 | 2056.802 | 537.2811 | 0.261222 | -1.93665 | 3.021697 |
| VCA0229 | 1918.285 | 2942.57 | 1396.516 | 719.64022 | 512.1391 | 507.39361 | 1.93E-08 | 1.53E-07 | 2085.79 | 579.7243 | 0.27794 | -1.84716 | 3.041246 |
| VCA0230 | 4040.643 | 5194.154 | 2894.47 | 1340.7277 | 1062.395 | 1027.2093 | 9.89E-12 | 1.06E-10 | 4043.089 | 1143.444 | 0.282815 | -1.82207 | 3.332464 |
| VCA0231 | 398.72701 | 695.6876 | 373.3953 | 176.57364 | 144.65 | 132.82055 | 4.77E-08 | 3.66E-07 | 489.27 | 151.3481 | 0.309334 | -1.69276 | 2.434763 |
| VCA0233 | 51.279851 | 69.33373 | 57.7145 | 25.664773 | 26.38885 | 21.977501 | 8.92E-06 | 5.12E-05 | 59.44269 | 24.67704 | 0.41514 | -1.26833 | 1.583196 |
| VCA0235 | 3391.7959 | 6891.067 | 5232.781 | 19698.226 | 18594.36 | 14774.614 | 3.32E-09 | 2.77E-08 | 5171.881 | 17689.07 | 3.420238 | 1.774097 | 3.980677 |
| VCA0236 | 276.28328 | 454.7822 | 474.8329 | 1126.1702 | 1096.603 | 914.45515 | 9.06E-08 | 6.7E-07 | 401.9661 | 1045.743 | 2.601569 | 1.379382 | 2.811807 |
| VCA0249 | 159.07219 | 121.0402 | 137.2906 | 332.61545 | 363.5797 | 362.151 | 1.17E-15 | 1.66E-14 | 139.1343 | 352.782 | 2.53555 | 1.342299 | 2.34547 |
| VCA0254 | 120.35067 | 108.1136 | 138.165 | 1306.8502 | 1271.551 | 1240.2955 | 2.4E-114 | 3.1E-112 | 122.2098 | 1272.899 | 10.41569 | 3.380687 | 2.59595 |
| VCA0255 | 136.04859 | 119.8651 | 127.6715 | 983.47408 | 1060.441 | 1004.2762 | 2.3E-106 | 2.7E-104 | 127.8617 | 1016.064 | 7.946583 | 2.990335 | 2.556831 |
| VCA0256 | 208.25899 | 243.2556 | 223.8623 | 2092.1923 | 2101.334 | 2127.9954 | 3.8E-163 | 8.9E-161 | 225.1256 | 2107.174 | 9.359991 | 3.226507 | 2.838063 |
| VCA0257 | 375.7034 | 420.7029 | 438.98 | 3883.5934 | 4008.173 | 4002.7718 | 2E-180 | 6.7E-178 | 411.7954 | 3964.846 | 9.628193 | 3.267265 | 3.106454 |
| VCA0258 | 253.25967 | 230.329 | 238.7282 | 6313.534 | 7052.663 | 7159.8876 | 0 | 0 | 240.7723 | 6842.028 | 28.41701 | 4.828683 | 3.108396 |
| VCA0271 | 1082.1095 | 1146.944 | 1093.952 | 9709.4968 | 9562.536 | 9564.0351 | 0 | 0 | 1107.669 | 9612.023 | 8.677706 | 3.117314 | 3.513612 |
| VCA0332 | 4354.6013 | 4175.3 | 3652.628 | 2943.2361 | 2647.681 | 2675.5219 | 1.22E-05 | 6.86E-05 | 4060.843 | 2755.48 | 0.678549 | -0.55948 | 3.524407 |
| VCA0345 | 386.16868 | 262.058 | 522.0539 | 185.81295 | 154.4236 | 196.84197 | 0.000164 | 0.000763 | 390.0935 | 179.0262 | 0.458931 | -1.12365 | 2.422043 |
| VCA0361a | 23.023607 | 39.95503 | 24.48494 | 100.60591 | 63.5287 | 60.199242 | 0.00015 | 0.000701 | 29.15452 | 74.77795 | 2.564883 | 1.358893 | 1.66924 |
| VCA0455 | 232.32912 | 249.1314 | 232.6069 | 482.49772 | 440.7915 | 456.7498 | 1.02E-13 | 1.29E-12 | 238.0225 | 460.013 | 1.932645 | 0.950577 | 2.519694 |
| VCA0473 | 81.629151 | 106.9385 | 79.57605 | 206.34477 | 172.0162 | 176.77555 | 3.35E-07 | 2.32E-06 | 89.38122 | 185.0455 | 2.070295 | 1.049836 | 2.109262 |
| VCA0497 | 2548.2947 | 2285.663 | 2297.212 | 1386.9243 | 1319.442 | 1417.071 | 1.12E-13 | 1.41E-12 | 2377.056 | 1374.479 | 0.578227 | -0.79029 | 3.257089 |
| VCA0498 | 5777.8788 | 4440.884 | 5516.982 | 3657.7434 | 3310.334 | 3561.3107 | 0.000166 | 0.000769 | 5245.248 | 3509.796 | 0.669138 | -0.57962 | 3.632524 |
| VCA0501 | 248.02704 | 264.4083 | 309.5596 | 186.83954 | 166.152 | 162.4424 | 8.07E-05 | 0.000401 | 273.9983 | 171.8113 | 0.627052 | -0.67334 | 2.3364 |
| VCA0510 | 1343.7414 | 1061.159 | 1382.525 | 1833.4914 | 2341.766 | 2568.501 | 4.6E-05 | 0.00024 | 1262.475 | 2247.919 | 1.780566 | 0.832336 | 3.226502 |
| VCA0511 | 328.60966 | 370.1716 | 350.6593 | 568.73136 | 665.5853 | 636.39199 | 9.52E-10 | 8.48E-09 | 349.8135 | 623.5696 | 1.782577 | 0.833964 | 2.669361 |
| VCA0514 | 3729.8243 | 4443.234 | 3599.286 | 21885.891 | 19939.22 | 19187.314 | 4.23E-62 | 2.36E-60 | 3924.115 | 20337.47 | 5.182691 | 2.373701 | 3.951019 |
| VCA0516 | 30201.739 | 7741.874 | 39277.34 | 657.01818 | 3144.182 | 5961.636 | 0.000148 | 0.000693 | 25740.32 | 3254.279 | 0.126427 | -2.98362 | 3.961534 |
| VCA0520 | 2693.762 | 2936.695 | 2747.56 | 1597.3754 | 1731.89 | 1915.8648 | 5.19E-08 | 3.96E-07 | 2792.672 | 1748.377 | 0.626059 | -0.67563 | 3.344327 |
| VCA0534 | 160.11872 | 237.3799 | 138.165 | 520.48159 | 475.9766 | 363.10654 | 1.26E-06 | 8.13E-06 | 178.5545 | 453.1882 | 2.538094 | 1.343746 | 2.454025 |
| VCA0538 | 954.43315 | 732.1171 | 800.1328 | 1603.535 | 1543.259 | 1727.6227 | 3.75E-10 | 3.53E-09 | 828.8944 | 1624.805 | 1.960208 | 0.971007 | 3.06465 |
| VCA0539 | 2245.8482 | 1938.994 | 2874.357 | 5828.9831 | 5699.991 | 5863.2151 | 4.64E-12 | 5.1E-11 | 2353.066 | 5797.396 | 2.463762 | 1.300863 | 3.567434 |
| VCA0540 | 544.19434 | 626.3538 | 1186.645 | 2020.3309 | 2374.019 | 2478.6799 | 3.92E-06 | 2.32E-05 | 785.7311 | 2291.01 | 2.915768 | 1.543876 | 3.12765 |
| VCA0542 | 519.07768 | 506.4887 | 530.7985 | 716.56045 | 731.0688 | 751.05721 | 1.32E-06 | 8.46E-06 | 518.7883 | 732.8955 | 1.412706 | 0.498461 | 2.790016 |
| VCA0545 | 3851.2215 | 3650.009 | 4864.633 | 2710.2 | 2650.613 | 2636.3446 | 7.89E-05 | 0.000394 | 4121.955 | 2665.719 | 0.646712 | -0.6288 | 3.520459 |
| VCA0546 | 376.74993 | 403.0757 | 298.1916 | 614.92795 | 604.9887 | 624.92546 | 2.53E-06 | 1.57E-05 | 359.3391 | 614.9474 | 1.711329 | 0.775117 | 2.672171 |
| VCA0547 | 2095.1482 | 2340.895 | 2181.783 | 4298.3361 | 3640.683 | 4326.7011 | 9.56E-12 | 1.02E-10 | 2205.942 | 4088.574 | 1.853437 | 0.890203 | 3.477583 |
| VCA0551 | 516.98462 | 491.2118 | 564.0281 | 1110.7714 | 990.0704 | 1139.9634 | 2.53E-15 | 3.52E-14 | 524.0748 | 1080.268 | 2.061287 | 1.043545 | 2.876462 |
| VCA0553 | 99.42012 | 131.6166 | 122.4247 | 258.70091 | 256.0695 | 237.93034 | 5.78E-10 | 5.33E-09 | 117.8205 | 250.9003 | 2.129514 | 1.090524 | 2.235361 |
| VCA0557 | 344.30757 | 287.9112 | 318.3042 | 590.28977 | 522.8901 | 558.99296 | 6.42E-09 | 5.29E-08 | 316.841 | 557.3909 | 1.759213 | 0.81493 | 2.623501 |
| VCA0558 | 327.56313 | 420.7029 | 359.4039 | 572.83772 | 558.0752 | 547.52644 | 5.38E-05 | 0.000278 | 369.2233 | 559.4798 | 1.515288 | 0.599592 | 2.657537 |
| VCA0563 | 11931.461 | 11923.05 | 14580.78 | 24498.565 | 24236.69 | 23516.882 | 8.99E-12 | 9.67E-11 | 12811.76 | 24084.05 | 1.879838 | 0.910609 | 4.244669 |
| VCA0564 | 8252.9165 | 8961.678 | 10477.81 | 19506.254 | 19036.13 | 19284.779 | 5.32E-15 | 7.24E-14 | 9230.8 | 19275.72 | 2.088196 | 1.062257 | 4.125125 |
| VCA0565 | 2382.9433 | 2798.027 | 2364.546 | 15949.116 | 12327.5 | 12603.619 | 3.85E-46 | 1.39E-44 | 2515.172 | 13626.75 | 5.417818 | 2.437712 | 3.76748 |
| VCA0566 | 5716.1336 | 6566.726 | 5293.119 | 202710.64 | 179931.9 | 188134.1 | 8.8E-292 | 4.1E-289 | 5858.66 | 190258.9 | 32.47481 | 5.021249 | 4.523572 |
| VCA0567 | 1187.8088 | 1316.166 | 1246.109 | 3884.62 | 3135.386 | 3257.4479 | 1.13E-24 | 2.35E-23 | 1250.028 | 3425.818 | 2.740594 | 1.454488 | 3.315842 |
| VCA0568 | 3387.6098 | 4027.232 | 3415.649 | 5934.722 | 5092.07 | 5297.5333 | 2.2E-05 | 0.00012 | 3610.164 | 5441.442 | 1.507256 | 0.591925 | 3.64662 |

| | | | | | | | | | | | | | |
|---|---|---|---|---|---|---|---|---|---|---|---|---|---|
| VCA0572 | 6974.0598 | 5127.17 | 7863.163 | 12772.844 | 13141.65 | 12821.483 | 1.5E-06 | 9.51E-06 | 6654.798 | 12911.99 | 1.940253 | 0.956245 | 3.967064 |
| VCA0574 | 740.94152 | 538.2177 | 814.1242 | 1151.835 | 1988.937 | 2288.5267 | 1.59E-05 | 8.88E-05 | 697.7612 | 1809.766 | 2.593676 | 1.374998 | 3.050665 |
| VCA0576 | 20524.499 | 28236.45 | 13792.02 | 2332.4145 | 1915.635 | 1869.0431 | 1.44E-29 | 3.62E-28 | 20850.99 | 2039.031 | 0.097791 | -3.35416 | 3.814275 |
| VCA0582 | 440.58811 | 539.3929 | 519.4305 | 319.26977 | 343.055 | 300.04067 | 2.7E-05 | 0.000145 | 499.8038 | 320.7885 | 0.641829 | -0.63974 | 2.602509 |
| VCA0588 | 1055.9463 | 867.2591 | 1535.555 | 1974.1343 | 2517.691 | 2545.5679 | 0.000121 | 0.000576 | 1152.92 | 2345.798 | 2.034657 | 1.024786 | 3.216045 |
| VCA0589 | 749.31375 | 545.2686 | 961.9083 | 1283.2386 | 1571.602 | 1636.8461 | 0.000115 | 0.000554 | 752.1636 | 1497.229 | 1.990563 | 0.993177 | 3.0258 |
| VCA0591 | 2973.1848 | 1890.813 | 3263.493 | 4786.9934 | 6325.504 | 6850.2915 | 3.06E-05 | 0.000164 | 2709.163 | 5987.596 | 2.210127 | 1.14413 | 3.605044 |
| VCA0594 | 1017.2248 | 1023.554 | 1113.19 | 14745.952 | 12285.47 | 12596.93 | 1.5E-166 | 3.9E-164 | 1051.323 | 13209.45 | 12.5646 | 3.651293 | 3.57131 |
| VCA0604 | 24.070134 | 18.80237 | 22.73602 | 123.19091 | 106.5327 | 93.643265 | 6.83E-17 | 1.01E-15 | 21.86951 | 107.789 | 4.928734 | 2.301217 | 1.686207 |
| VCA0605 | 29.302772 | 27.0284 | 37.60187 | 71.861363 | 57.66452 | 63.065873 | 0.000161 | 0.000751 | 31.31102 | 64.19725 | 2.050309 | 1.035841 | 1.651607 |
| VCA0625 | 398.72701 | 719.1905 | 234.3558 | 160.14818 | 124.1253 | 116.57631 | 3.81E-05 | 0.000201 | 450.7578 | 133.6166 | 0.296427 | -1.75425 | 2.389902 |
| VCA0652 | 8176.52 | 8141.425 | 6999.195 | 4345.5593 | 3928.029 | 4036.2158 | 6.67E-14 | 8.51E-13 | 7772.38 | 4103.268 | 0.527929 | -0.92158 | 3.751842 |
| VCA0656 | 1340.6018 | 981.2485 | 1317.814 | 372.6525 | 549.2789 | 702.32449 | 4.97E-05 | 0.000258 | 1213.222 | 541.4186 | 0.446265 | -1.16403 | 2.908737 |
| VCA0657 | 3593.7757 | 3250.459 | 3852.006 | 826.40568 | 1470.934 | 1613.913 | 4E-07 | 2.76E-06 | 3565.413 | 1303.751 | 0.365666 | -1.4514 | 3.333652 |
| VCA0659 | 516.98462 | 431.2793 | 417.9929 | 179.65341 | 272.6847 | 225.50827 | 3.24E-06 | 1.95E-05 | 455.4189 | 225.9488 | 0.496134 | -1.0112 | 2.506211 |
| VCA0663 | 1119.7845 | 1055.283 | 1226.87 | 630.32681 | 614.7624 | 645.94742 | 8.29E-13 | 9.69E-12 | 1133.979 | 630.3455 | 0.55587 | -0.84718 | 2.927092 |
| VCA0665 | 1080.0165 | 1075.26 | 1199.762 | 263.83386 | 255.0922 | 250.3524 | 8.17E-66 | 5.02E-64 | 1118.346 | 256.4261 | 0.22929 | -2.12475 | 2.728769 |
| VCA0673 | 1504.9067 | 1137.543 | 1296.827 | 721.6934 | 741.8198 | 784.50123 | 2.61E-07 | 1.84E-06 | 1313.092 | 749.3381 | 0.570667 | -0.80928 | 2.996487 |
| VCA0677 | 435.35547 | 491.2118 | 494.9456 | 914.69249 | 930.4512 | 769.21254 | 2.71E-09 | 2.3E-08 | 473.8376 | 871.4521 | 1.839137 | 0.879029 | 2.807937 |
| VCA0678 | 4926.0053 | 5442.11 | 5676.134 | 10921.901 | 11724.47 | 11680.564 | 6.39E-24 | 1.26E-22 | 5348.083 | 11442.31 | 2.139516 | 1.097285 | 3.893356 |
| VCA0679 | 687.56862 | 834.355 | 886.7046 | 1838.6243 | 1686.931 | 1659.7791 | 3.39E-13 | 4.14E-12 | 802.8761 | 1728.445 | 2.152817 | 1.106225 | 3.071152 |
| VCA0680 | 1279.9032 | 1349.07 | 1358.914 | 2814.9123 | 2703.391 | 2576.1453 | 3.3E-26 | 7.24E-25 | 1329.296 | 2698.149 | 2.029759 | 1.021308 | 3.277344 |
| VCA0697 | 1314.4386 | 1279.736 | 1222.498 | 267.94023 | 254.1148 | 288.57414 | 7.84E-77 | 5.78E-75 | 1272.224 | 270.2097 | 0.212392 | -2.2352 | 2.768132 |
| VCA0699 | 161.16525 | 101.0627 | 181.0137 | 29.771136 | 31.27567 | 26.755219 | 1.17E-13 | 1.47E-12 | 147.7472 | 29.26734 | 0.198091 | -2.33577 | 1.817951 |
| VCA0704 | 335.93535 | 357.245 | 327.0488 | 652.91181 | 631.3776 | 611.54785 | 1.57E-14 | 2.1E-13 | 340.0764 | 631.9457 | 1.858247 | 0.893942 | 2.666128 |
| VCA0705 | 255.35273 | 293.787 | 263.2131 | 729.90613 | 646.038 | 608.68122 | 4.61E-18 | 7.18E-17 | 270.7843 | 661.5418 | 2.443058 | 1.288688 | 2.62659 |
| VCA0706 | 205.11941 | 265.5834 | 218.6155 | 633.40659 | 584.4641 | 503.57144 | 3.83E-13 | 4.6E-12 | 229.7728 | 573.814 | 2.497311 | 1.320375 | 2.560035 |
| VCA0715 | 46.047213 | 41.13018 | 50.7188 | 99.579318 | 94.80437 | 87.910004 | 3.5E-06 | 2.09E-05 | 45.9654 | 94.0979 | 2.047146 | 1.033614 | 1.818005 |
| VCA0716 | 61.745127 | 61.10769 | 51.59327 | 111.89841 | 98.71383 | 93.643265 | 0.000166 | 0.000771 | 58.14869 | 101.4185 | 1.744123 | 0.802502 | 1.885329 |
| VCA0717 | 2077.3572 | 2580.625 | 2068.977 | 5440.9318 | 4463.624 | 4400.2779 | 2.53E-10 | 2.44E-09 | 2242.32 | 4768.278 | 2.126493 | 1.088476 | 3.51453 |
| VCA0721 | 333.8423 | 289.0864 | 290.3214 | 2858.0291 | 2860.746 | 2860.8973 | 1.6E-169 | 4.4E-167 | 304.4167 | 2859.891 | 9.394658 | 3.231841 | 2.969909 |
| VCA0722 | 164.30483 | 180.9728 | 195.0051 | 1655.8911 | 1704.524 | 1626.3351 | 3.1E-136 | 5.4E-134 | 180.0942 | 1662.25 | 9.229891 | 3.206314 | 2.738098 |
| VCA0723 | 876.99011 | 1356.121 | 887.5791 | 5108.3163 | 4856.525 | 4451.8773 | 2.11E-22 | 3.96E-21 | 1040.23 | 4805.573 | 4.619722 | 2.207806 | 3.349437 |
| VCA0732 | 400.82006 | 426.5787 | 378.6421 | 1000.9261 | 943.1569 | 935.47711 | 3.18E-27 | 7.24E-26 | 402.0136 | 959.8534 | 2.387614 | 1.25557 | 2.793223 |
| VCA0734 | 835.12901 | 970.6722 | 666.3401 | 435.27454 | 450.5651 | 408.97263 | 1.06E-06 | 6.92E-06 | 824.0471 | 431.6041 | 0.523761 | -0.93302 | 2.775519 |
| VCA0735 | 1553.0469 | 1515.941 | 1316.94 | 6112.3222 | 5605.186 | 5498.1974 | 1.42E-55 | 7.16E-54 | 1461.976 | 5738.569 | 3.925214 | 1.972771 | 3.461872 |
| VCA0736 | 2860.1599 | 2379.674 | 3133.198 | 1786.2682 | 1754.37 | 1744.8225 | 9.13E-06 | 5.23E-05 | 2791.011 | 1761.82 | 0.631248 | -0.66372 | 3.345862 |
| VCA0737 | 4160.9936 | 3271.612 | 3602.784 | 2155.8409 | 2162.908 | 2187.2391 | 2.84E-08 | 2.22E-07 | 3678.463 | 2168.663 | 0.589557 | -0.7623 | 3.450929 |
| VCA0738 | 516.98462 | 497.0876 | 545.6644 | 258.70091 | 280.5037 | 269.46327 | 2.83E-14 | 3.71E-13 | 519.9122 | 269.5559 | 0.518464 | -0.94768 | 2.573289 |
| VCA0739 | 534.77559 | 524.116 | 536.0453 | 200.18523 | 218.9297 | 210.21958 | 1.48E-27 | 3.49E-26 | 531.6456 | 209.7782 | 0.394583 | -1.3416 | 2.523691 |
| VCA0740 | 4217.5061 | 4116.543 | 4445.765 | 2204.0907 | 2272.373 | 2026.7078 | 2.82E-21 | 4.95E-20 | 4259.938 | 2167.724 | 0.508863 | -0.97465 | 3.482704 |
| VCA0772 | 1713.1656 | 1756.846 | 1271.468 | 717.58704 | 687.0874 | 744.3684 | 2.25E-10 | 2.18E-09 | 1580.493 | 716.3476 | 0.453243 | -1.14164 | 3.026958 |
| VCA0785 | 63.838182 | 81.0852 | 98.81422 | 146.8025 | 171.0388 | 154.79805 | 2.9E-05 | 0.000156 | 81.24587 | 157.5465 | 1.939132 | 0.955411 | 2.053605 |
| VCA0789 | 86.861789 | 56.4071 | 75.20374 | 120.11114 | 217.9523 | 159.57577 | 0.000134 | 0.000633 | 72.82421 | 165.8797 | 2.27781 | 1.187648 | 2.041035 |
| VCA0799 | 1081.063 | 870.7846 | 888.4535 | 546.14636 | 489.6597 | 514.08242 | 7.27E-09 | 5.95E-08 | 946.767 | 516.6295 | 0.545678 | -0.87388 | 2.844711 |
| VCA0802 | 9163.3955 | 6871.09 | 10681.55 | 4094.0445 | 4216.351 | 4757.6512 | 9.55E-07 | 6.23E-06 | 8905.347 | 4356.016 | 0.489146 | -1.03166 | 3.79437 |
| VCA0811 | 5403.2219 | 4452.635 | 5892.126 | 30328.575 | 40939.85 | 40694.688 | 4.15E-45 | 1.48E-43 | 5249.328 | 37321.04 | 7.10968 | 2.829785 | 4.146029 |
| VCA0817 | 928.26996 | 859.0331 | 770.4011 | 528.69431 | 512.1391 | 471.08296 | 4.22E-08 | 3.26E-07 | 852.5681 | 503.9721 | 0.591122 | -0.75847 | 2.816568 |
| VCA0822 | 619.54433 | 722.716 | 778.2713 | 1009.1389 | 1037.961 | 1020.5205 | 0.000108 | 0.000519 | 706.8439 | 1022.54 | 1.446628 | 0.532694 | 2.929502 |

| | | | | | | | | | | | | | |
|---|---|---|---|---|---|---|---|---|---|---|---|---|---|
| VCA0845 | 369.42423 | 487.6864 | 414.495 | 3145.4745 | 3002.464 | 2739.5433 | 2.76E-69 | 1.79E-67 | 423.8686 | 2962.494 | 6.989181 | 2.805123 | 3.049444 |
| VCA0847 | 101.51318 | 103.413 | 109.3078 | 334.66863 | 332.304 | 312.46273 | 2.58E-27 | 6.03E-26 | 104.7447 | 326.4785 | 3.116899 | 1.640111 | 2.266993 |
| VCA0849 | 2601.6676 | 2251.583 | 3281.856 | 4313.735 | 4741.196 | 4379.256 | 0.000105 | 0.000509 | 2711.702 | 4478.062 | 1.651384 | 0.723676 | 3.542166 |
| VCA0868 | 91.047899 | 47.00592 | 87.44621 | 20.531818 | 24.43412 | 37.266197 | 0.000174 | 0.000802 | 75.16668 | 27.41071 | 0.364666 | -1.45535 | 1.656973 |
| VCA0900 | 390.35479 | 324.3408 | 397.8803 | 615.95454 | 648.9701 | 618.23666 | 7.03E-08 | 5.27E-07 | 370.8586 | 627.7204 | 1.692614 | 0.759253 | 2.683487 |
| VCA0901 | 276.28328 | 297.3124 | 308.6851 | 471.20522 | 462.2935 | 437.63893 | 5.16E-07 | 3.52E-06 | 294.0936 | 457.0459 | 1.554083 | 0.636064 | 2.564223 |
| VCA0902 | 102.5597 | 84.61065 | 122.4247 | 209.42454 | 215.0202 | 209.26403 | 1.41E-07 | 1.03E-06 | 103.1983 | 211.2363 | 2.046896 | 1.033438 | 2.169221 |
| VCA0907 | 13913.584 | 19223.07 | 11047.95 | 2740.9977 | 2345.675 | 2244.5717 | 4.57E-25 | 9.74E-24 | 14728.2 | 2443.748 | 0.165923 | -2.59141 | 3.778103 |
| VCA0908 | 6150.4426 | 9344.776 | 4728.217 | 1171.3402 | 883.5377 | 743.41286 | 6.99E-19 | 1.13E-17 | 6741.145 | 932.7636 | 0.138369 | -2.85341 | 3.399253 |
| VCA0909 | 4469.7193 | 8076.791 | 3272.237 | 1220.6166 | 876.6961 | 821.76743 | 1.05E-10 | 1.05E-09 | 5272.916 | 973.0267 | 0.184533 | -2.43805 | 3.355088 |
| VCA0910 | 1221.2977 | 2040.057 | 1047.606 | 371.62591 | 310.802 | 248.44132 | 5.06E-12 | 5.53E-11 | 1436.32 | 310.2897 | 0.216031 | -2.21069 | 2.824509 |
| VCA0911 | 2289.8023 | 4597.179 | 1781.279 | 674.47022 | 444.7009 | 462.48307 | 1.03E-09 | 9.14E-09 | 2889.42 | 527.2181 | 0.182465 | -2.45431 | 3.0914 |
| VCA0912 | 1234.9025 | 2343.245 | 865.7175 | 297.71136 | 245.3185 | 196.84197 | 1.34E-10 | 1.32E-09 | 1481.288 | 246.624 | 0.166493 | -2.58647 | 2.781337 |
| VCA0913 | 1671.3045 | 3405.579 | 1246.109 | 514.32204 | 379.2175 | 357.37328 | 1.01E-08 | 8.21E-08 | 2107.664 | 416.9709 | 0.197836 | -2.33763 | 2.971954 |
| VCA0914 | 1476.6504 | 3149.396 | 1032.74 | 457.85954 | 327.4172 | 276.15208 | 6.99E-08 | 5.25E-07 | 1886.262 | 353.8096 | 0.187572 | -2.41449 | 2.912186 |
| VCA0915 | 1115.5984 | 1982.474 | 699.5697 | 286.41886 | 175.9256 | 163.39794 | 2.02E-09 | 1.74E-08 | 1265.881 | 208.5808 | 0.164771 | -2.60146 | 2.710834 |
| VCA0917 | 2658.18 | 2223.38 | 2967.924 | 24300.433 | 28282.98 | 31217.607 | 2.09E-78 | 1.6E-76 | 2616.495 | 27933.67 | 10.67599 | 3.416298 | 3.931924 |
| VCA0935 | 3.1395827 | 4.700592 | 3.497849 | 18.478636 | 14.66047 | 16.24424 | 0.000135 | 0.00064 | 3.779341 | 16.46112 | 4.355552 | 2.122856 | 0.896938 |
| VCA0936 | 3369.8188 | 2759.247 | 3683.234 | 1535.78 | 1376.129 | 1279.4728 | 8.85E-13 | 1.03E-11 | 3270.767 | 1397.127 | 0.427156 | -1.22717 | 3.329943 |
| VCA0947 | 2289.8023 | 1955.446 | 2506.208 | 14349.688 | 13859.03 | 13113.879 | 7.94E-75 | 5.53E-73 | 2250.486 | 13774.2 | 6.120545 | 2.61366 | 3.745671 |
| VCA0948 | 1320.7178 | 908.3893 | 1527.685 | 6934.6215 | 6179.877 | 6116.4341 | 3.74E-24 | 7.5E-23 | 1252.264 | 6410.311 | 5.118977 | 2.355855 | 3.452288 |
| VCA0954 | 6295.9099 | 5534.947 | 5531.847 | 2569.557 | 2510.85 | 2559.9011 | 9.34E-29 | 2.31E-27 | 5787.568 | 2546.769 | 0.440041 | -1.18429 | 3.584243 |
| VCA0962 | 500.24018 | 460.658 | 505.4391 | 18336.967 | 17027.65 | 16465.926 | 0 | 0 | 488.7791 | 17276.85 | 35.34694 | 5.143514 | 3.463289 |
| VCA0972 | 375.7034 | 364.2958 | 350.6593 | 2311.8827 | 1598.969 | 1556.5804 | 1.17E-27 | 2.77E-26 | 363.5529 | 1822.477 | 5.012964 | 2.325664 | 2.910615 |
| VCA0976 | 471.98394 | 836.7053 | 313.9319 | 119.08454 | 96.7591 | 88.865548 | 7.63E-10 | 6.95E-09 | 540.8737 | 101.5697 | 0.187788 | -2.41282 | 2.36993 |
| VCA0977 | 1513.2789 | 2128.193 | 1181.398 | 677.55 | 534.6185 | 478.7273 | 7.93E-08 | 5.91E-07 | 1607.623 | 563.6319 | 0.350599 | -1.5121 | 2.97859 |
| VCA0981 | 325.47008 | 99.88757 | 302.5639 | 53.382727 | 78.18917 | 73.576851 | 8.33E-05 | 0.000413 | 242.6405 | 68.38292 | 0.281828 | -1.82711 | 2.109955 |
| VCA0982 | 2063.7524 | 605.2012 | 1917.695 | 197.10545 | 272.6847 | 289.52969 | 2.47E-08 | 1.94E-07 | 1528.883 | 253.1066 | 0.16555 | -2.59466 | 2.793839 |
| VCA0983 | 728.38319 | 273.8095 | 1054.601 | 156.04182 | 183.7446 | 219.77501 | 7.25E-05 | 0.000365 | 685.598 | 186.5205 | 0.272055 | -1.87803 | 2.553398 |
| VCA0989 | 1134.4359 | 594.6248 | 2094.337 | 312.08363 | 300.051 | 262.77447 | 1.63E-06 | 1.03E-05 | 1274.466 | 291.6364 | 0.22883 | -2.12765 | 2.785085 |
| VCA1006 | 131.86247 | 185.6734 | 149.533 | 333.64204 | 267.7979 | 316.28491 | 1.5E-06 | 9.51E-06 | 155.6896 | 305.9083 | 1.96486 | 0.974426 | 2.338925 |
| VCA1008 | 51.279851 | 57.58225 | 58.58896 | 128.32386 | 108.4875 | 105.10979 | 1.42E-06 | 9.06E-06 | 55.81702 | 113.9737 | 2.041917 | 1.029924 | 1.901786 |
| VCA1021 | 9788.1724 | 8462.24 | 8693.902 | 5790.9993 | 5918.92 | 6387.8084 | 8.28E-07 | 5.45E-06 | 8981.438 | 6032.576 | 0.671671 | -0.57417 | 3.866924 |
| VCA1025 | 1550.9539 | 2054.159 | 1467.347 | 3084.9057 | 2963.37 | 2823.6311 | 7.64E-06 | 4.42E-05 | 1690.82 | 2957.302 | 1.749034 | 0.806559 | 3.349497 |
| VCA1032 | 394.5409 | 400.7254 | 417.9929 | 4351.7188 | 3800.971 | 3872.8179 | 1.4E-164 | 3.4E-162 | 404.4197 | 4008.503 | 9.911738 | 3.309138 | 3.104907 |
| VCA1041 | 23592.918 | 14587.11 | 25850.85 | 4830.1102 | 8079.874 | 8554.0256 | 1.35E-06 | 8.65E-06 | 21343.63 | 7154.67 | 0.335213 | -1.57685 | 4.091929 |
| VCA1074 | 1226.5303 | 1211.577 | 1130.68 | 876.70863 | 854.2167 | 921.14396 | 3.62E-05 | 0.000192 | 1189.596 | 884.0231 | 0.743129 | -0.42832 | 3.010932 |
| VCA1101 | 198.84024 | 172.7467 | 212.4943 | 100.60591 | 120.2159 | 102.24316 | 3.74E-06 | 2.23E-05 | 194.6938 | 107.6883 | 0.553116 | -0.85435 | 2.16076 |
| VCA1105 | 422.79714 | 312.5893 | 463.4649 | 160.14818 | 235.5449 | 228.3749 | 0.000175 | 0.000806 | 399.6171 | 208.0227 | 0.520555 | -0.94188 | 2.459877 |
| VCA1114 | 5412.6406 | 5106.018 | 6115.988 | 8104.9352 | 10657.18 | 12126.803 | 1.06E-05 | 6.05E-05 | 5544.882 | 10296.31 | 1.856903 | 0.892898 | 3.878287 |
| VCr024 | 206.16593 | 195.0746 | 167.0223 | 69.808181 | 63.5287 | 117.53185 | 5.58E-05 | 0.000288 | 189.4209 | 83.62291 | 0.441466 | -1.17963 | 2.099877 |

#### Supplementary Table S5. List of genes that are  exhibited in both RNA-Seq and VxrB ChIP-Seq

### This file is generated by MACS version 2.0.10.20120913 (tag:beta)
### ARGUMENTS LIST:
### name = peaks
### format = AUTO
### ChIP-seq file = ['CHIPWigRHis100PG3H.bam']
### control file = ['InpWigRHis100PG3H.bam']
### effective genome size = 2.70e+09
### band width = 300
### model fold = [5, 50]
### qvalue cutoff = 1.00e-10
### Larger dataset will be scaled towards smaller dataset.
### Range for calculating regional lambda is: 1000 bps and 10000 bps
### Broad region calling is off

### tag size is determined as 150 bps
### total tags in treatment: 6618387
### tags after filtering in treatment: 2812499
### maximum duplicate tags at the same position in treatment = 1
### Redundant rate in treatment: 0.58
### total tags in control: 9724075
### tags after filtering in control: 4734726
### maximum duplicate tags at the same position in control = 1
### Redundant rate in control: 0.51
# d = 200

| chr | start | end | length | abs_summit (summit position) | pileup (stacked bases) | -LOG10(pvalue) of summit | fold enrichment | -LOG10(qvalue) | name | ID | Rank | Downstream Gene_FWD | Downstream Gene_REV | Start FWD | Start Rev | Dist FWD | Dist Rev | FWD Gene Log2FC | FWD Gene Padj | REV Gene Log2FC | REV Gene Padj | FWD DEG? | REV DEG? | Meet criteria? | Regulation |
|---|---|---|---|---|---|---|---|---|---|---|---|---|---|---|---|---|---|---|---|---|---|---|---|---|---|
| NC_002505 | 20807 | 21475 | 669 | 21301 | 266 | 14.49967 | 1.67129 | 12.77599 | peaks_peak_6 | Peak_chr1_1 | 345 | VC0025 | NA | 21656 | | 355 | | -0.5679 | 0.3083 | | | N | | | |
| NC_002505 | 44654 | 45468 | 815 | 44885 | 339 | 34.27369 | 2.10523 | 32.06459 | peaks_peak_10 | Peak_chr1_2 | 156 | VC0047 | def (NC_002505 | 44900 | 44808 | 15 | 77 | -0.0134 | 0.9650 | -0.4789 | 0.0061 | N | N | | |
| NC_002505 | 46815 | 47221 | 407 | 46981 | 323 | 25.01373 | 1.88941 | 23.01955 | peaks_peak_11 | Peak_chr1_3 | 219 | VC0049 | NA | 47193 | | 212 | | -1.1550 | 0.0000 | | | Y | | 1 | Down |
| NC_002505 | 90658 | 91009 | 352 | 90836 | 342 | 36.61581 | 2.15971 | 34.35271 | peaks_peak_19 | Peak_chr1_4 | 144 | VC0093 | VC0092 | 90962 | 90642 | 126 | 194 | -0.7426 | 0.0015 | -0.0059 | 0.9794 | N | N | | |
| NC_002505 | 94532 | 94854 | 323 | 94699 | 290 | 20.73194 | 1.83064 | 18.84114 | peaks_peak_20 | Peak_chr1_5 | 256 | VC0097 | ubiA | 95062 | 94318 | 363 | 381 | -0.2611 | 0.1390 | -0.2833 | 0.1168 | N | N | | |
| NC_002505 | 105834 | 106133 | 300 | 105986 | 263 | 14.81652 | 1.68766 | 13.08221 | peaks_peak_24 | Peak_chr1_6 | 335 | NA | NA | | | | | | | | | | | | |
| NC_002505 | 107057 | 107654 | 598 | 107259 | 285 | 18.64637 | 1.77478 | 16.80893 | peaks_peak_25 | Peak_chr1_7 | 279 | VC0112 | NA | 107409 | | 150 | | -0.3609 | 0.2562 | | | N | | | |
| NC_002505 | 109874 | 110220 | 347 | 110052 | 280 | 17.74968 | 1.7554 | 15.93566 | peaks_peak_26 | Peak_chr1_8 | 289 | NA | NA | | | | | | | | | | | | |
| NC_002505 | 134439 | 135147 | 709 | 134779 | 248 | 12.11572 | 1.61507 | 10.47253 | peaks_peak_29 | Peak_chr1_9 | 402 | VC0143 | VC0142a | 134836 | 134475 | 57 | 304 | -1.6547 | 0.2598 | 0.9757 | 0.1868 | N | N | | |
| NC_002505 | 144823 | 145272 | 450 | 145036 | 342 | 35.85617 | 2.13954 | 33.6104 | peaks_peak_30 | Peak_chr1_10 | 148 | VC0156 | VC0154 | 145370 | 144972 | 334 | 64 | -0.6682 | 0.0581 | -0.1310 | 0.8706 | N | N | | |
| NC_002505 | 148899 | 149306 | 408 | 149063 | 332 | 34.08725 | 2.11845 | 31.88239 | peaks_peak_32 | Peak_chr1_11 | 159 | VC0158 | VC0157 | 149278 | 148885 | 215 | 178 | -0.0502 | 0.8784 | 6.0040 | 0.0000 | N | | | |
| NC_002505 | 166827 | 167265 | 439 | 167038 | 280 | 19.36474 | 1.80786 | 17.50901 | peaks_peak_40 | Peak_chr1_12 | 274 | tRNA-Arg-1 | VC0168 | 167187 | 166897 | 149 | 141 | 1.7129 | 0.7322 | -0.5865 | 0.0035 | N | N | | |
| NC_002505 | 167995 | 168563 | 569 | 168415 | 301 | 24.73361 | 1.92882 | 22.74656 | peaks_peak_41 | Peak_chr1_13 | 222 | VC0170 | VC0169 | 168532 | 168399 | 117 | 16 | -1.0089 | 0.0788 | 0.0823 | 0.9988 | N | N | | |
| NC_002505 | 176732 | 177439 | 708 | 177245 | 240 | 12.52953 | 1.64481 | 10.87126 | peaks_peak_43 | Peak_chr1_14 | 390 | NA | VC0175 | | 176941 | | 304 | | | -0.3125 | 0.2428 | | N | | |
| NC_002505 | 178100 | 178964 | 865 | 178744 | 248 | 14.59802 | 1.70799 | 12.87071 | peaks_peak_44 | Peak_chr1_15 | 340 | NA | NA | | | | | | | | | | | | |
| NC_002505 | 200455 | 200811 | 357 | 200646 | 334 | 30.24522 | 2.00923 | 28.13066 | peaks_peak_50 | Peak_chr1_16 | 184 | NA | VC0194 | | 200599 | | 47 | | | -0.4250 | 0.5606 | | N | | |
| NC_002505 | 221440 | 221779 | 340 | 221598 | 340 | 31.56861 | 2.03075 | 29.42199 | peaks_peak_51 | Peak_chr1_17 | 176 | NA | VC0213 | | 221341 | | 257 | | | 1.6599 | 0.0000 | | Y | 2 | Up |
| NC_002505 | 222028 | 222331 | 304 | 222199 | 325 | 31.36217 | 2.06161 | 29.22062 | peaks_peak_52 | Peak_chr1_18 | 178 | NA | slmA | | 221928 | | 271 | | | -0.3315 | 0.0046 | | N | | |
| NC_002505 | 231691 | 232116 | 426 | 231929 | 259 | 12.94695 | 1.62784 | 11.27414 | peaks_peak_53 | Peak_chr1_19 | 375 | VC0227 | VC0225 | 231950 | 231870 | 21 | 59 | -0.4493 | 0.0436 | -0.5154 | 0.0000 | N | Y | 3 | Down |
| NC_002505 | 234082 | 235308 | 1227 | 234218 | 252 | 13.30765 | 1.65286 | 11.62344 | peaks_peak_54 | Peak_chr1_20 | 368 | NA | NA | | | | | | | | | | | | |
| NC_002505 | 254215 | 254887 | 673 | 254465 | 242 | 14.75966 | 1.7262 | 13.02708 | peaks_peak_61 | Peak_chr1_21 | 337 | VC0248 | NA | 254505 | | 40 | | -0.7239 | 0.0000 | | | Y | | 4 | Down |
| NC_002505 | 257131 | 257694 | 564 | 257299 | 314 | 32.87756 | 2.1341 | 30.70063 | peaks_peak_62 | Peak_chr1_22 | 167 | NA | NA | | | | | | | | | | | | |
| NC_002505 | 278124 | 278605 | 482 | 278322 | 335 | 37.61663 | 2.20623 | 35.32866 | peaks_peak_69 | Peak_chr1_23 | 140 | VC0273 | VC0271 | 278432 | 278060 | 110 | 262 | 0.3988 | 0.1057 | -1.2497 | 0.0052 | N | N | | |
| NC_002505 | 282520 | 282820 | 301 | 282661 | 303 | 25.71098 | 1.9533 | 23.69982 | peaks_peak_70 | Peak_chr1_24 | 214 | zntR | purH | 282765 | 282479 | 104 | 182 | -0.9269 | 0.0000 | -0.2799 | 0.2083 | Y | N | 5 | Down |
| NC_002505 | 299628 | 300102 | 475 | 299865 | 363 | 47.27649 | 2.37526 | 44.72988 | peaks_peak_72 | Peak_chr1_25 | 88 | NA | VC0289 | | 299750 | | 115 | | | -0.3883 | 0.0307 | | N | | |
| NC_002505 | 329636 | 330077 | 442 | 329861 | 329 | 30.51612 | 2.02839 | 28.39545 | peaks_peak_79 | Peak_chr1_26 | 181 | NA | NA | | | | | | | | | | | | |
| NC_002505 | 330913 | 331812 | 900 | 331127 | 378 | 51.33161 | 2.42908 | 48.66179 | peaks_peak_80 | Peak_chr1_27 | 72 | murB | pssA | 331605 | 331051 | 478 | 76 | 1.7627 | 0.0000 | 1.1488 | 0.0000 | Y | Y | 6 | Up |
| NC_002505 | 360326 | 360666 | 341 | 360516 | 312 | 28.78175 | 2.02131 | 26.70035 | peaks_peak_85 | Peak_chr1_28 | 193 | NA | NA | | | | | | | | | | | | |
| NC_002505 | 375201 | 375689 | 489 | 375416 | 377 | 45.47874 | 2.28833 | 42.98847 | peaks_peak_87 | Peak_chr1_29 | 98 | VC0354 | NA | 375590 | | 174 | | 1.8005 | 0.0000 | | | Y | | 7 | Up |
| NC_002505 | 386245 | 386718 | 474 | 386488 | 368 | 48.25588 | 2.38423 | 45.68023 | peaks_peak_91 | Peak_chr1_30 | 82 | VC0371 | VC0370 | 386594 | 386468 | 106 | 20 | 0.8594 | 0.0000 | 3.5291 | 0.0000 | Y | Y | 8 | Up |
| NC_002505 | 391449 | 391892 | 444 | 391683 | 327 | 32.37654 | 2.08459 | 30.21124 | peaks_peak_92 | Peak_chr1_31 | 171 | VC0376 | NA | 391741 | | 58 | | 0.1601 | 0.6648 | | | N | | | |
| NC_002505 | 421820 | 422526 | 707 | 422057 | 298 | 24.80823 | 1.93804 | 22.81907 | peaks_peak_98 | Peak_chr1_32 | 221 | NA | NA | | | | | | | | | | | | |
| NC_002505 | 444943 | 445442 | 500 | 445160 | 315 | 28.11467 | 1.99477 | 26.04833 | peaks_peak_103 | Peak_chr1_33 | 198 | VC0419 | NA | 445206 | | 46 | | 0.3017 | 0.0002 | | | Y | | 9 | Up |
| NC_002505 | 461809 | 462249 | 441 | 462099 | 292 | 21.9732 | 1.86497 | 20.05072 | peaks_peak_105 | Peak_chr1_34 | 242 | VC0432 | VC0431 | 462243 | 461915 | 144 | 184 | -0.9260 | 0.1172 | 0.1111 | 0.5923 | N | N | | |
| NC_002505 | 495119 | 495405 | 287 | 495269 | 252 | 13.23831 | 1.6503 | 11.55661 | peaks_peak_106 | Peak_chr1_35 | 369 | NA | NA | | | | | | | | | | | | |
| NC_002505 | 510481 | 510957 | 477 | 510718 | 322 | 35.10947 | 2.17465 | 32.88123 | peaks_peak_108 | Peak_chr1_36 | 151 | VC0480 | NA | 510777 | | 59 | | 1.7278 | 0.0000 | | | Y | | 10 | Up |
| NC_002505 | 513176 | 513778 | 603 | 513417 | 330 | 35.12903 | 2.15242 | 32.90027 | peaks_peak_109 | Peak_chr1_37 | 150 | VC0483 | NA | 513480 | | 63 | | 4.3679 | 0.0000 | | | | | | |
| NC_002505 | 532424 | 532873 | 450 | 532656 | 314 | 32.52176 | 2.12385 | 30.3532 | peaks_peak_115 | Peak_chr1_38 | 170 | NA | VC0500 | | 532411 | | 245 | | | 1.4763 | 0.0000 | | Y | 11 | Up |
| NC_002505 | 535096 | 535478 | 383 | 535240 | 253 | 15.91523 | 1.74599 | 14.14931 | peaks_peak_116 | Peak_chr1_39 | 319 | VC0503 | NA | 535418 | | 178 | | -0.8703 | 0.0000 | | | Y | | 12 | Down |
| NC_002505 | 539946 | 540528 | 583 | 540179 | 261 | 20.92893 | 1.90374 | 19.03273 | peaks_peak_117 | Peak_chr1_40 | 253 | NA | VC0510 | | 540004 | | 175 | | | -0.9160 | 0.0011 | | N | | |
| NC_002505 | 544022 | 546240 | 2219 | 544242 | 277 | 31.57719 | 2.21148 | 29.43033 | peaks_peak_120 | Peak_chr1_41 | 175 | VC0513 | NA | 544362 | | 120 | | -0.7888 | 0.0002 | | | Y | | 13 | Down |
| NC_002505 | 548496 | 549009 | 514 | 548676 | 213 | 11.97412 | 1.67545 | 10.33628 | peaks_peak_122 | Peak_chr1_42 | 410 | NA | NA | | | | | | | | | | | | |
| NC_002505 | 563077 | 564843 | 1767 | 564169 | 397 | 63.44891 | 2.65981 | 60.24549 | peaks_peak_125 | Peak_chr1_43 | 32 | NA | NA | | | | | | | | | | | | |
| NC_002505 | 580292 | 580694 | 403 | 580481 | 263 | 16.64312 | 1.75137 | 14.85751 | peaks_peak_127 | Peak_chr1_44 | 304 | VC0548 | NA | 580589 | | 108 | | -0.2980 | 0.0976 | | | N | | | |
| NC_002505 | 594734 | 595026 | 293 | 594899 | 254 | 14.55521 | 1.69487 | 12.8293 | peaks_peak_130 | Peak_chr1_45 | 343 | NA | VC0559 | | 594461 | | 438 | | | 0.1186 | 0.5193 | | N | | |
| NC_002505 | 597378 | 597707 | 330 | 597544 | 288 | 22.66558 | 1.89541 | 20.72677 | peaks_peak_131 | Peak_chr1_46 | 237 | rplS | NA | 597929 | | 385 | | -0.1981 | 0.7669 | | | N | | | |
| NC_002505 | 601440 | 601755 | 316 | 601591 | 284 | 23.08668 | 1.91804 | 21.13806 | peaks_peak_132 | Peak_chr1_47 | 230 | VC0568 | VC0567 | 601728 | 601574 | 137 | 17 | -0.3939 | 0.0149 | 0.8739 | 0.0000 | N | Y | | Up |
| NC_002505 | 604838 | 605114 | 277 | 604981 | 250 | 12.93047 | 1.64234 | 11.2583 | peaks_peak_134 | Peak_chr1_48 | 378 | NA | NA | | | | | | | | | | | | |
| NC_002505 | 611443 | 611969 | 527 | 611803 | 358 | 42.75315 | 2.27474 | 40.33649 | peaks_peak_136 | Peak_chr1_49 | 113 | NA | VC0581 | | 611416 | | 387 | | | 1.8976 | 0.0000 | | Y | 14 | Up |
| NC_002505 | 622413 | 622744 | 332 | 622582 | 265 | 17.57383 | 1.77936 | 15.76401 | peaks_peak_139 | Peak_chr1_50 | 292 | NA | VC0594 | | 622245 | | 337 | | | -0.0350 | 0.9108 | | N | | |
| NC_002505 | 623676 | 624061 | 386 | 623874 | 305 | 28.52184 | 2.0316 | 26.44569 | peaks_peak_140 | Peak_chr1_51 | 194 | NA | VC0596 | | 623745 | | 129 | | | 0.2568 | 0.1560 | | N | | |
| NC_002505 | 628061 | 628402 | 342 | 628239 | 303 | 22.27448 | 1.851 | 20.34448 | peaks_peak_141 | Peak_chr1_52 | 239 | NA | NA | | | | | | | | | | | | |
| NC_002505 | 630404 | 630675 | 272 | 630558 | 259 | 12.32331 | 1.60516 | 10.67227 | peaks_peak_142 | Peak_chr1_53 | 394 | NA | NA | | | | | | | | | | | | |
| NC_002505 | 636584 | 636984 | 401 | 636772 | 334 | 40.10639 | 2.2773 | 37.7582 | peaks_peak_143 | Peak_chr1_54 | 122 | VC0606 | NA | 637173 | | 401 | | -0.3406 | 0.4812 | | | N | | | |
| NC_002505 | 663446 | 664103 | 658 | 663590 | 257 | 16.04509 | 1.74246 | 14.27558 | peaks_peak_147 | Peak_chr1_55 | 316 | NA | NA | | | | | | | | | | | | |
| NC_002505 | 670926 | 673059 | 2134 | 671328 | 391 | 61.32818 | 2.63037 | 58.25627 | peaks_peak_148 | Peak_chr1_56 | 38 | NA | NA | | | | | | | | | | | | |
| NC_002505 | 684383 | 684609 | 227 | 684469 | 264 | 16.15863 | 1.7326 | 14.38593 | peaks_peak_149 | Peak_chr1_57 | 312 | NA | NA | | | | | | | | | | | | |
| NC_002505 | 694082 | 694533 | 452 | 694375 | 281 | 20.72178 | 1.8495 | 18.83117 | peaks_peak_150 | Peak_chr1_58 | 258 | NA | NA | | | | | | | | | | | | |
| NC_002505 | 695034 | 695758 | 725 | 695527 | 364 | 41.6158 | 2.22887 | 39.22874 | peaks_peak_151 | Peak_chr1_59 | 118 | NA | NA | | | | | | | | | | | | |
| NC_002505 | 707646 | 707941 | 296 | 707807 | 250 | 15.22877 | 1.7272 | 13.48215 | peaks_peak_152 | Peak_chr1_60 | 329 | VC0662 | VC0661 | 707977 | 707668 | 170 | 139 | -0.3764 | 0.0231 | -0.1562 | 0.6228 | N | N | | |
| NC_002505 | 713890 | 714295 | 406 | 714034 | 241 | 13.14983 | 1.6669 | 11.47079 | peaks_peak_153 | Peak_chr1_61 | 371 | VC0666 | NA | 714307 | | 273 | | 1.2472 | 0.0000 | | | Y | | 15 | Up |
| NC_002505 | 716634 | 717027 | 394 | 716786 | 267 | 19.57212 | 1.84298 | 17.71095 | peaks_peak_154 | Peak_chr1_62 | 272 | VC0671 | VC0668 | 717080 | 716512 | 294 | 274 | -0.1681 | 0.3344 | -0.4268 | 0.0309 | N | N | | |
| NC_002505 | 723568 | 724154 | 587 | 723755 | 366 | 53.47551 | 2.5244 | 50.73292 | peaks_peak_155 | Peak_chr1_63 | 66 | nhaR | VC0676 | 724055 | 723660 | 300 | 95 | 0.3371 | 0.1996 | 1.2349 | 0.0000 | N | Y | 16 | Up |
| NC_002505 | 725780 | 726246 | 467 | 726005 | 352 | 47.12893 | 2.40764 | 44.58707 | peaks_peak_156 | Peak_chr1_64 | 90 | VC0680 | rpsT | 726042 | 725799 | 37 | 206 | 3.4632 | 0.0000 | -0.4972 | 0.3030 | Y | N | | Up |
| NC_002505 | 739071 | 740155 | 1085 | 739806 | 392 | 60.14623 | 2.59773 | 57.13753 | peaks_peak_159 | Peak_chr1_65 | 45 | VC0692 | anmK | 739874 | 739768 | 68 | 38 | 0.2386 | 0.3228 | 0.0107 | 0.9455 | N | N | | |
| NC_002505 | 753655 | 753990 | 336 | 753807 | 274 | 21.07038 | 1.8767 | 19.17101 | peaks_peak_160 | Peak_chr1_66 | 251 | NA | VC0703 | | 753622 | | 185 | | | 1.7744 | 0.0000 | | Y | 17 | Up |
| NC_002505 | 755918 | 756924 | 1007 | 756599 | 258 | 16.85218 | 1.76895 | 15.06098 | peaks_peak_161 | Peak_chr1_67 | 298 | NA | VC0706 | | 756569 | | 30 | | | -1.8754 | 0.0006 | | Y | 18 | Down |
| NC_002505 | 778869 | 779260 | 392 | 779065 | 326 | 37.61255 | 2.23302 | 35.32477 | peaks_peak_165 | Peak_chr1_68 | 141 | VC0724 | VC0723 | 779108 | 778909 | 43 | 156 | -0.2723 | 0.0142 | 0.7671 | 0.0350 | N | N | | |
| NC_002505 | 788318 | 789169 | 852 | 788569 | 265 | 19.87583 | 1.85791 | 18.00656 | peaks_peak_166 | Peak_chr1_69 | 268 | VC0734 | NA | 788949 | | 380 | | 0.7095 | 0.0111 | | | N | | | |
| NC_002505 | 799421 | 800013 | 593 | 799747 | 375 | 57.04038 | 2.58271 | 54.15988 | peaks_peak_167 | Peak_chr1_70 | 52 | VC0746 | VC0745 | 799804 | 799625 | 57 | 122 | -1.4962 | 0.0000 | -0.2152 | 0.8743 | Y | N | 19 | Down |
| NC_002505 | 809556 | 810043 | 488 | 809824 | 372 | 56.08322 | 2.56945 | 53.23766 | peaks_peak_170 | Peak_chr1_71 | 54 | VC0758 | NA | 810001 | | 177 | | 0.4977 | 0.0027 | | | N | | | |
| NC_002505 | 819478 | 820020 | 543 | 819598 | 255 | 13.98841 | 1.67246 | 12.28215 | peaks_peak_172 | Peak_chr1_72 | 352 | VC0767 | NA | 820012 | | 414 | | -0.5309 | 0.4459 | | | N | | | |
| NC_002505 | 841988 | 842432 | 445 | 842242 | 308 | 32.75872 | 2.14809 | 30.58434 | peaks_peak_174 | Peak_chr1_73 | 168 | NA | VC0784 | | 841989 | | 253 | | | -0.8588 | 0.2052 | | N | | |
| NC_002505 | 893481 | 894476 | 996 | 894065 | 276 | 30.27129 | 2.17212 | 28.1561 | peaks_peak_186 | Peak_chr1_74 | 183 | VC0832 | NA | 894412 | | 347 | | 0.0478 | 0.8979 | | | N | | | |
| NC_002505 | 898815 | 902363 | 3549 | 900213 | 226 | 22.77344 | 2.07955 | 20.83233 | peaks_peak_190 | Peak_chr1_75 | 234 | NA | NA | | | | | | | | | | | | |
| NC_002505 | 911578 | 911925 | 348 | 911756 | 225 | 13.77408 | 1.72433 | 12.07463 | peaks_peak_194 | Peak_chr1_76 | 358 | NA | NA | | | | | | | | | | | | |
| NC_002505 | 922998 | 923287 | 290 | 923135 | 298 | 28.20002 | 2.04049 | 26.13175 | peaks_peak_197 | Peak_chr1_77 | 197 | NA | NA | | | | | | | | | | | | |
| NC_002505 | 925000 | 925301 | 302 | 925128 | 255 | 18.43201 | 1.83137 | 16.60048 | peaks_peak_198 | Peak_chr1_78 | 281 | VC0860 | NA | 925340 | | 212 | | 1.2109 | 0.0000 | | | Y | | 20 | Up |
| NC_002505 | 928351 | 928654 | 304 | 928500 | 279 | 22.14977 | 1.90004 | 20.22357 | peaks_peak_199 | Peak_chr1_79 | 241 | NA | NA | | | | | | | | | | | | |
| NC_002505 | 940191 | 941659 | 1469 | 940776 | 392 | 68.56988 | 2.80668 | 65.03608 | peaks_peak_202 | Peak_chr1_80 | 16 | VC0880 | VC0876 | 941187 | 940490 | 411 | 286 | 0.8455 | 0.0142 | -0.0771 | 0.6140 | N | N | | |
| NC_002505 | 941956 | 942705 | 750 | 942350 | 374 | 57.53819 | 2.59918 | 54.63897 | peaks_peak_203 | Peak_chr1_81 | 49 | NA | NA | | | | | | | | | | | | |
| NC_002505 | 962070 | 962304 | 235 | 962196 | 262 | 13.47029 | 1.64174 | 11.78094 | peaks_peak_208 | Peak_chr1_82 | 365 | NA | NA | | | | | | | | | | | | |
| NC_002505 | 980401 | 980774 | 374 | 980582 | 341 | 43.00624 | 2.33393 | 40.58389 | peaks_peak_211 | Peak_chr1_83 | 111 | wecC | NA | 980595 | | 13 | | 1.1917 | 0.0000 | | | Y | | 21 | Up |
| NC_002505 | 982618 | 983065 | 448 | 982814 | 266 | 21.6414 | 1.91528 | 19.72722 | peaks_peak_212 | Peak_chr1_84 | 243 | NA | NA | | | | | | | | | | | | |
| NC_002505 | 1000738 | 1000982 | 245 | 1000863 | 244 | 13.79156 | 1.68564 | 12.0915 | peaks_peak_213 | Peak_chr1_85 | 356 | NA | NA | | | | | | | | | | | | |
| NC_002505 | 1001676 | 1002056 | 381 | 1001862 | 322 | 35.09714 | 2.1743 | 32.86915 | peaks_peak_214 | Peak_chr1_86 | 152 | VC0936 | NA | 1001910 | | 48 | | 0.5203 | 0.0389 | | | N | | | |
| NC_002505 | 1007451 | 1007692 | 242 | 1007605 | 243 | 11.83999 | 1.61284 | 10.20728 | peaks_peak_216 | Peak_chr1_87 | 416 | NA | NA | | | | | | | | | | | | |
| NC_002505 | 1012163 | 1012597 | 435 | 1012381 | 321 | 38.7357 | 2.28065 | 36.41997 | peaks_peak_217 | Peak_chr1_88 | 131 | NA | VC0947 | | 1012213 | | 168 | | | 1.3366 | 0.0000 | | Y | 22 | Up |
| NC_002505 | 1013181 | 1013593 | 413 | 1013406 | 375 | 61.14527 | 2.688 | 58.08503 | peaks_peak_218 | Peak_chr1_89 | 40 | NA | VC0948 | | 1013096 | | 310 | | | 2.4147 | 0.0000 | | Y | 23 | Up |
| NC_002505 | 1031364 | 1032184 | 821 | 1031990 | 350 | 52.35248 | 2.55485 | 49.64951 | peaks_peak_219 | Peak_chr1_90 | 69 | NA | VC0966 | | 1031872 | | 118 | | | 0.3184 | 0.5330 | | N | | |
| NC_002505 | 1033848 | 1034264 | 417 | 1034054 | 327 | 38.50333 | 2.25485 | 36.19363 | peaks_peak_220 | Peak_chr1_91 | 133 | VC0970 | VC0969 | 1034243 | 1034013 | 189 | 41 | 1.1308 | 0.0000 | 3.0456 | 0.0000 | Y | Y | 24 | Up |
| NC_002505 | 1039043 | 1039454 | 412 | 1039229 | 276 | 26.54198 | 2.05026 | 24.51179 | peaks_peak_223 | Peak_chr1_92 | 210 | NA | NA | | | | | | | | | | | | |
| NC_002505 | 1040761 | 1041819 | 1059 | 1041312 | 381 | 63.22756 | 2.71705 | 60.03635 | peaks_peak_224 | Peak_chr1_93 | 34 | NA | NA | | | | | | | | | | | | |
| NC_002505 | 1043254 | 1043698 | 445 | 1043465 | 257 | 20.55158 | 1.90094 | 18.6651 | peaks_peak_225 | Peak_chr1_94 | 261 | NA | VC0977 | | 1043084 | | 381 | | | -1.7569 | 0.0004 | | Y | 25 | Down |
| NC_002505 | 1047978 | 1048312 | 335 | 1048127 | 261 | 21.13986 | 1.911 | 19.2382 | peaks_peak_226 | Peak_chr1_95 | 250 | VC0985 | VC0984 | 1048252 | 1048024 | 125 | 103 | -1.3958 | 0.0000 | 0.4134 | 0.0001 | Y | Y | | Down |
| NC_002505 | 1061042 | 1061553 | 512 | 1061246 | 339 | 47.28175 | 2.45778 | 44.73499 | peaks_peak_227 | Peak_chr1_96 | 87 | VC0995 | nagA | 1061324 | 1060855 | 78 | 391 | -0.3880 | 0.3583 | -0.0965 | 0.3836 | N | N | | |
| NC_002505 | 1079426 | 1079721 | 296 | 1079572 | 252 | 17.13282 | 1.79194 | 15.33441 | peaks_peak_228 | Peak_chr1_97 | 294 | NA | NA | | | | | | | | | | | | |
| NC_002505 | 1127484 | 1127990 | 507 | 1127752 | 377 | 56.38986 | 2.55886 | 53.5333 | peaks_peak_233 | Peak_chr1_98 | 53 | NA | VC1059 | | 1127555 | | 197 | | | -0.4302 | 0.0001 | | Y | 26 | Down |
| NC_002505 | 1130019 | 1130926 | 908 | 1130436 | 381 | 65.26976 | 2.76947 | 61.969 | peaks_peak_234 | Peak_chr1_99 | 26 | NA | VC1061 | | 1129967 | | 469 | | | 0.4230 | 0.1577 | | N | | |
| NC_002505 | 1131282 | 1131669 | 388 | 1131456 | 330 | 43.9232 | 2.39684 | 41.47543 | peaks_peak_235 | Peak_chr1_100 | 109 | VC1064 | VC1063 | 1131567 | 1131416 | 111 | 40 | 1.8794 | 0.0000 | -1.2040 | 0.0000 | Y | Y | | Up |
| NC_002505 | 1136930 | 1137428 | 499 | 1137205 | 352 | 51.79832 | 2.53218 | 49.11334 | peaks_peak_239 | Peak_chr1_101 | 71 | VC1070 | NA | 1137311 | | 106 | | 0.4062 | 0.0061 | | | N | | | |
| NC_002505 | 1158175 | 1158702 | 528 | 1158508 | 232 | 14.50461 | 1.73788 | 12.78068 | peaks_peak_241 | Peak_chr1_102 | 344 | VC1091 | VC1089 | 1158668 | 1158044 | 160 | 464 | 0.2359 | 0.6783 | 0.3733 | 0.2169 | N | N | | |
| NC_002505 | 1160441 | 1161270 | 830 | 1160644 | 306 | 36.19617 | 2.25624 | 33.94256 | peaks_peak_242 | Peak_chr1_103 | 146 | NA | NA | | | | | | | | | | | | |
| NC_002505 | 1163290 | 1163733 | 444 | 1163514 | 359 | 52.09127 | 2.51384 | 49.39691 | peaks_peak_243 | Peak_chr1_104 | 70 | NA | NA | | | | | | | | | | | | |
| NC_002505 | 1168930 | 1169238 | 309 | 1169052 | 229 | 11.87188 | 1.63897 | 10.23802 | peaks_peak_244 | Peak_chr1_105 | 414 | VC1099 | VC1098 | 1169290 | 1168943 | 238 | 109 | -0.0773 | 0.7366 | 0.4753 | 0.4310 | N | N | | |
| NC_002505 | 1170128 | 1171137 | 1010 | 1170581 | 380 | 63.5905 | 2.73047 | 60.37962 | peaks_peak_245 | Peak_chr1_106 | 31 | VC1101 | VC1100 | 1170719 | 1170281 | 138 | 300 | -0.7194 | 0.0704 | -0.8203 | 0.0840 | N | N | | |
| NC_002505 | 1176898 | 1177146 | 249 | 1177003 | 247 | 15.45983 | 1.74187 | 13.7065 | peaks_peak_247 | Peak_chr1_107 | 325 | lolA | NA | 1177056 | | 53 | | 2.0842 | 0.0000 | | | Y | | 27 | Up |
| NC_002505 | 1190461 | 1190704 | 244 | 1190564 | 234 | 12.92274 | 1.67148 | 11.25088 | peaks_peak_249 | Peak_chr1_108 | 379 | NA | NA | | | | | | | | | | | | |
| NC_002505 | 1200348 | 1201617 | 1270 | 1200981 | 380 | 67.2246 | 2.8245 | 63.77274 | peaks_peak_251 | Peak_chr1_109 | 21 | hisG | NA | 1201260 | | 279 | | 0.2533 | 0.0643 | | | N | | | |
| NC_002505 | 1206486 | 1206803 | 318 | 1206631 | 268 | 22.8204 | 1.94973 | 20.87807 | peaks_peak_252 | Peak_chr1_110 | 233 | VC1138 | NA | 1207016 | | 385 | | 0.6678 | 0.0000 | | | Y | | 28 | Up |
| NC_002505 | 1221427 | 1221812 | 386 | 1221587 | 329 | 47.6357 | 2.50601 | 45.0792 | peaks_peak_253 | Peak_chr1_111 | 85 | NA | NA | | | | | | | | | | | | |
| NC_002505 | 1226195 | 1226531 | 337 | 1226360 | 304 | 36.5589 | 2.2739 | 34.29718 | peaks_peak_256 | Peak_chr1_112 | 145 | NA | VC1154 | | 1226227 | | 133 | | | 1.3459 | 0.0000 | | Y | 29 | Up |
| NC_002505 | 1228464 | 1229387 | 924 | 1229018 | 366 | 60.42415 | 2.70641 | 57.40019 | peaks_peak_257 | Peak_chr1_113 | 43 | VC1158 | VC1156 | 1229089 | 1228866 | 71 | 152 | 3.1421 | 0.0000 | 0.5858 | 0.2204 | Y | N | 30 | Up |
| NC_002505 | 1232282 | 1232549 | 268 | 1232405 | 250 | 18.10946 | 1.83172 | 16.28624 | peaks_peak_258 | Peak_chr1_114 | 286 | NA | NA | | | | | | | | | | | | |
| NC_002505 | 1234761 | 1235199 | 439 | 1234975 | 380 | 60.15955 | 2.64294 | 57.1497 | peaks_peak_259 | Peak_chr1_115 | 44 | NA | VC1162 | | 1234894 | | 81 | | | 1.9036 | 0.0000 | | Y | 31 | Up |
| NC_002505 | 1237653 | 1238266 | 614 | 1237934 | 384 | 67.91038 | 2.82455 | 64.4163 | peaks_peak_260 | Peak_chr1_116 | 19 | aspS | VC1165 | 1238124 | 1237832 | 190 | 102 | -0.3016 | 0.0870 | -0.6362 | 0.0001 | N | Y | 32 | Down |
| NC_002505 | 1241832 | 1242131 | 300 | 1241951 | 224 | 11.87577 | 1.6487 | 10.24178 | peaks_peak_261 | Peak_chr1_117 | 413 | NA | VC1168 | | 1241932 | | 19 | | | -0.2138 | 0.5238 | | N | | |
| NC_002505 | 1247167 | 1247475 | 309 | 1247338 | 284 | 26.64048 | 2.03055 | 24.60815 | peaks_peak_262 | Peak_chr1_118 | 207 | NA | VC1173 | | 1247330 | | 8 | | | 0.2688 | 0.4885 | | N | | |
| NC_002505 | 1252942 | 1253314 | 373 | 1253149 | 301 | 29.43087 | 2.06935 | 27.33464 | peaks_peak_264 | Peak_chr1_119 | 188 | NA | NA | | | | | | | | | | | | |
| NC_002505 | 1265964 | 1266311 | 348 | 1266155 | 248 | 19.47701 | 1.88614 | 17.61835 | peaks_peak_267 | Peak_chr1_120 | 273 | NA | VC1191 | | 1266106 | | 49 | | | 1.2186 | 0.0000 | | Y | 33 | Up |
| NC_002505 | 1319097 | 1319419 | 323 | 1319271 | 323 | 39.46406 | 2.29482 | 37.1312 | peaks_peak_270 | Peak_chr1_121 | 126 | NA | VC1247 | | 1319198 | | 73 | | | -0.4792 | 0.0079 | | N | | |
| NC_002505 | 1344451 | 1344770 | 320 | 1344599 | 307 | 37.91693 | 2.30423 | 35.6218 | peaks_peak_272 | Peak_chr1_122 | 138 | VC1269 | NA | 1344755 | | 156 | | 2.4890 | 0.0000 | | | Y | | 34 | Up |
| NC_002505 | 1346546 | 1346920 | 375 | 1346762 | 298 | 34.84215 | 2.24228 | 32.61919 | peaks_peak_274 | Peak_chr1_123 | 154 | VC1272 | VC1271 | 1346837 | 1346733 | 75 | 29 | -2.0542 | 0.0000 | 2.4333 | 0.0000 | Y | Y | | Down |
| NC_002505 | 1353242 | 1353692 | 451 | 1353468 | 306 | 39.11659 | 2.34383 | 36.79183 | peaks_peak_275 | Peak_chr1_124 | 129 | VC1279 | VC1278 | 1353601 | 1353244 | 133 | 224 | 0.0954 | 0.7136 | -0.3137 | 0.1515 | N | N | | |
| NC_002505 | 1359521 | 1359832 | 312 | 1359665 | 237 | 15.92963 | 1.78165 | 14.1633 | peaks_peak_276 | Peak_chr1_125 | 317 | NA | NA | | | | | | | | | | | | |
| NC_002505 | 1371320 | 1371683 | 364 | 1371486 | 298 | 34.25888 | 2.22444 | 32.05008 | peaks_peak_279 | Peak_chr1_126 | 157 | NA | VC1291 | | 1371254 | | 232 | | | 0.0820 | 0.7784 | | N | | |
| NC_002505 | 1378973 | 1379438 | 466 | 1379264 | 278 | 28.20484 | 2.09827 | 26.13639 | peaks_peak_280 | Peak_chr1_127 | 196 | VC1299 | VC1298 | 1379571 | 1379173 | 307 | 91 | -0.6923 | 0.1048 | -0.8865 | 0.0936 | N | N | | |
| NC_002505 | 1381460 | 1381675 | 216 | 1381584 | 224 | 12.45057 | 1.67243 | 10.79516 | peaks_peak_281 | Peak_chr1_128 | 392 | NA | VC1300 | | 1381389 | | 195 | | | -0.7926 | 0.0015 | | N | | |
| NC_002505 | 1408733 | 1409052 | 320 | 1408889 | 255 | 22.90015 | 1.98873 | 20.95604 | peaks_peak_286 | Peak_chr1_129 | 232 | VC1325 | VC1323 | 1409080 | 1408581 | 191 | 308 | -0.0430 | 0.9698 | 0.5593 | 0.0164 | N | N | | |
| NC_002505 | 1410101 | 1410445 | 345 | 1410283 | 298 | 39.15545 | 2.37517 | 36.82971 | peaks_peak_287 | Peak_chr1_130 | 128 | NA | NA | | | | | | | | | | | | |

| | | | | | | | | | | | | | | | | | | | | | | | | | |
|---|---|---|---|---|---|---|---|---|---|---|---|---|---|---|---|---|---|---|---|---|---|---|---|---|---|
| NC_002505 | 1418508 | 1418763 | 256 | 1418643 | 232 | 16.46606 | 1.81441 | 14.68497 | peaks_peak_288 | Peak_chr1_131 | 307 | VC1335 | VC1334 | 1419022 | 1418392 | 379 | 251 | 0.1677 | 0.4385 | -0.0221 | 0.9712 | N | N | | |
| NC_002505 | 1430890 | 1431498 | 609 | 1431270 | 275 | 27.84762 | 2.09594 | 25.78715 | peaks_peak_290 | Peak_chr1_132 | 201 | VC1344 | VC1343 | 1431464 | 1431040 | 194 | 230 | 1.8009 | 0.0000 | 0.9879 | 0.0000 | Y | Y | 35 | Up |
| NC_002505 | 1436288 | 1436877 | 590 | 1436548 | 348 | 55.25502 | 2.64243 | 52.44188 | peaks_peak_291 | Peak_chr1_133 | 60 | VC1349 | VC1348 | 1436615 | 1436493 | 67 | 55 | 0.6645 | 0.0006 | 0.0119 | 0.9849 | Y | N | 36 | Up |
| NC_002505 | 1438533 | 1439028 | 496 | 1438795 | 370 | 55.54283 | 2.56301 | 52.71869 | peaks_peak_292 | Peak_chr1_134 | 59 | NA | NA | | | | | | | | | | | | |
| NC_002505 | 1452230 | 1452741 | 512 | 1452526 | 290 | 33.28684 | 2.22127 | 31.09996 | peaks_peak_294 | Peak_chr1_135 | 164 | VC1363 | VC1362 | 1452643 | 1452188 | 117 | 338 | 0.1104 | 0.4496 | -0.0907 | 0.8724 | N | N | | |
| NC_002505 | 1456251 | 1458124 | 1874 | 1456815 | 388 | 74.12511 | 2.96744 | 70.05345 | peaks_peak_295 | Peak_chr1_136 | 7 | NA | NA | | | | | | | | | | | | |
| NC_002505 | 1470438 | 1470712 | 275 | 1470572 | 225 | 12.28747 | 1.6637 | 10.63777 | peaks_peak_297 | Peak_chr1_137 | 397 | VC1377 | NA | 1471032 | | 460 | | -0.7349 | 0.0000 | | | Y | | 37 | Down |
| NC_002505 | 1489948 | 1490786 | 839 | 1490522 | 285 | 33.10244 | 2.23271 | 30.91968 | peaks_peak_299 | Peak_chr1_138 | 165 | NA | NA | | | | | | | | | | | | |
| NC_002505 | 1493377 | 1493853 | 477 | 1493524 | 279 | 25.5172 | 2.00853 | 23.51076 | peaks_peak_300 | Peak_chr1_139 | 216 | NA | NA | | | | | | | | | | | | |
| NC_002505 | 1495201 | 1495767 | 567 | 1495437 | 376 | 72.76059 | 2.99055 | 68.84422 | peaks_peak_301 | Peak_chr1_140 | 9 | NA | NA | | | | | | | | | | | | |
| NC_002505 | 1496486 | 1497581 | 1096 | 1496692 | 323 | 44.10071 | 2.4276 | 41.64748 | peaks_peak_302 | Peak_chr1_141 | 108 | NA | NA | | | | | | | | | | | | |
| NC_002505 | 1499226 | 1499830 | 605 | 1499591 | 384 | 71.32064 | 2.91317 | 67.55944 | peaks_peak_303 | Peak_chr1_142 | 11 | VC1407 | NA | 1499666 | | 75 | | -0.4132 | 0.1615 | | | N | | | |
| NC_002505 | 1503483 | 1503777 | 295 | 1503624 | 231 | 15.09689 | 1.76339 | 13.35428 | peaks_peak_305 | Peak_chr1_143 | 331 | NA | NA | | | | | | | | | | | | |
| NC_002505 | 1506967 | 1507449 | 483 | 1507168 | 334 | 48.86776 | 2.52124 | 46.27251 | peaks_peak_307 | Peak_chr1_144 | 79 | NA | NA | | | | | | | | | | | | |
| NC_002505 | 1514270 | 1514662 | 393 | 1514480 | 347 | 49.77359 | 2.49631 | 47.15183 | peaks_peak_308 | Peak_chr1_145 | 76 | VC1418 | NA | 1514611 | | 131 | | 3.7077 | 0.0000 | | | Y | | 38 | Up |
| NC_002505 | 1520497 | 1521037 | 541 | 1520739 | 298 | 41.98685 | 2.46352 | 39.59053 | peaks_peak_310 | Peak_chr1_146 | 116 | VC1422a | VC1422 | 1520836 | 1520333 | 97 | 406 | 1.9960 | 0.0000 | 0.1743 | 0.7804 | Y | N | 39 | Up |
| NC_002505 | 1529918 | 1531283 | 1366 | 1530487 | 382 | 72.80769 | 2.96194 | 68.88506 | peaks_peak_312 | Peak_chr1_147 | 8 | NA | NA | | | | | | | | | | | | |
| NC_002505 | 1539519 | 1540471 | 953 | 1540146 | 350 | 59.66706 | 2.75625 | 56.68484 | peaks_peak_316 | Peak_chr1_148 | 47 | VC1444 | NA | 1540238 | | 92 | | 1.4196 | 0.0000 | | | Y | | 40 | Up |
| NC_002505 | 1541125 | 1541427 | 303 | 1541276 | 229 | 15.8136 | 1.79637 | 14.05062 | peaks_peak_317 | Peak_chr1_149 | 322 | VC1445 | NA | 1541452 | | 176 | | 1.1560 | 0.0000 | | | Y | | 41 | Up |
| NC_002505 | 1545158 | 1545733 | 576 | 1545545 | 309 | 38.08369 | 2.30221 | 35.78411 | peaks_peak_319 | Peak_chr1_150 | 136 | NA | VC1446 | | 1545396 | | 149 | | | -1.0665 | 0.2083 | | N | | |
| NC_002505 | 1559023 | 1559395 | 373 | 1559212 | 330 | 37.36329 | 2.21399 | 35.08251 | peaks_peak_321 | Peak_chr1_151 | 142 | NA | NA | | | | | | | | | | | | |
| NC_002505 | 1560639 | 1560964 | 326 | 1560798 | 266 | 21.6414 | 1.91528 | 19.72722 | peaks_peak_322 | Peak_chr1_152 | 243 | NA | NA | | | | | | | | | | | | |
| NC_002505 | 1588411 | 1588657 | 247 | 1588548 | 212 | 11.62994 | 1.66267 | 10.00567 | peaks_peak_335 | Peak_chr1_153 | 423 | NA | NA | | | | | | | | | | | | |
| NC_002505 | 1589110 | 1589595 | 486 | 1589352 | 323 | 44.25067 | 2.43193 | 41.79332 | peaks_peak_336 | Peak_chr1_154 | 107 | NA | VC1484 | | 1589199 | | 153 | | | 3.6302 | 0.0000 | | Y | 42 | Up |
| NC_002505 | 1602504 | 1602840 | 337 | 1602689 | 261 | 24.52824 | 2.02749 | 22.54644 | peaks_peak_339 | Peak_chr1_155 | 225 | NA | VC1492 | | 1602422 | | 267 | | | 0.2161 | 0.8303 | | N | | |
| NC_002505 | 1605819 | 1606174 | 356 | 1605989 | 249 | 21.35905 | 1.95131 | 19.45225 | peaks_peak_340 | Peak_chr1_156 | 248 | NA | NA | | | | | | | | | | | | |
| NC_002505 | 1608918 | 1609261 | 344 | 1609102 | 219 | 12.93398 | 1.70309 | 11.26161 | peaks_peak_341 | Peak_chr1_157 | 377 | NA | NA | | | | | | | | | | | | |
| NC_002505 | 1630659 | 1630920 | 262 | 1630790 | 253 | 20.56675 | 1.91197 | 18.67993 | peaks_peak_343 | Peak_chr1_158 | 260 | NA | VC1516 | | 1630456 | | 334 | | | 3.4211 | 0.0000 | | Y | 43 | Up |
| NC_002505 | 1649234 | 1649609 | 376 | 1649419 | 307 | 40.71469 | 2.38829 | 38.35043 | peaks_peak_344 | Peak_chr1_159 | 120 | VC1537 | VC1535 | 1649484 | 1649395 | 65 | 24 | -1.2446 | 0.0000 | 0.0057 | 0.9986 | Y | N | 44 | Down |
| NC_002505 | 1660617 | 1662098 | 1482 | 1661145 | 391 | 69.56259 | 2.83611 | 65.96381 | peaks_peak_347 | Peak_chr1_160 | 14 | VC1549 | VC1548 | 1661257 | 1660938 | 112 | 207 | 0.3768 | 0.5255 | -1.1256 | 0.0003 | N | Y | | Down |
| NC_002505 | 1675395 | 1675715 | 321 | 1675546 | 263 | 23.72265 | 1.99402 | 21.76017 | peaks_peak_349 | Peak_chr1_161 | 227 | VC1563 | NA | 1675622 | | 76 | | 0.5683 | 0.0230 | | | N | | | |
| NC_002505 | 1680451 | 1680780 | 330 | 1680629 | 310 | 38.31422 | 2.30556 | 36.00933 | peaks_peak_350 | Peak_chr1_162 | 135 | NA | NA | | | | | | | | | | | | |
| NC_002505 | 1703465 | 1703730 | 266 | 1703612 | 232 | 16.08478 | 1.79959 | 14.31404 | peaks_peak_352 | Peak_chr1_163 | 315 | VC1592 | NA | 1703680 | | 68 | | 0.6994 | 0.0874 | | | N | | | |
| NC_002505 | 1710937 | 1711727 | 791 | 1711531 | 351 | 57.26861 | 2.68519 | 54.37827 | peaks_peak_353 | Peak_chr1_164 | 50 | NA | VC1597 | | 1711152 | | 379 | | | -0.3091 | 0.0919 | | N | | |
| NC_002505 | 1731810 | 1732947 | 1138 | 1732605 | 393 | 69.04184 | 2.81429 | 65.47992 | peaks_peak_357 | Peak_chr1_165 | 15 | NA | NA | | | | | | | | | | | | |
| NC_002505 | 1741242 | 1741781 | 540 | 1741499 | 327 | 45.9857 | 2.46657 | 43.48005 | peaks_peak_358 | Peak_chr1_166 | 96 | VC1621 | VC1620 | 1741764 | 1741367 | 265 | 132 | -0.2742 | 0.3771 | 0.1993 | 0.2309 | N | N | | |
| NC_002505 | 1751033 | 1751571 | 539 | 1751417 | 279 | 28.8311 | 2.11543 | 26.74864 | peaks_peak_361 | Peak_chr1_167 | 192 | NA | nhaA | | 1751164 | | 253 | | | 0.6059 | 0.0062 | | N | | |
| NC_002505 | 1768082 | 1768578 | 497 | 1768273 | 252 | 20.87522 | 1.92562 | 18.98046 | peaks_peak_363 | Peak_chr1_168 | 254 | VC1643 | VC1642 | 1768345 | 1768154 | 72 | 119 | -0.0315 | 0.9291 | 1.8129 | 0.0000 | N | Y | 45 | Up |
| NC_002505 | 1770147 | 1770445 | 299 | 1770304 | 246 | 16.83594 | 1.79475 | 15.04515 | peaks_peak_364 | Peak_chr1_169 | 299 | VC1644 | NA | 1770467 | | 163 | | 0.1265 | 0.8636 | | | N | | | |
| NC_002505 | 1775883 | 1776219 | 337 | 1776017 | 243 | 19.18969 | 1.88889 | 17.33858 | peaks_peak_365 | Peak_chr1_170 | 275 | NA | NA | | | | | | | | | | | | |
| NC_002505 | 1777424 | 1778206 | 783 | 1777847 | 385 | 72.19291 | 2.93127 | 68.33851 | peaks_peak_366 | Peak_chr1_171 | 10 | NA | NA | | | | | | | | | | | | |
| NC_002505 | 1783846 | 1784106 | 261 | 1783981 | 240 | 12.6295 | 1.64868 | 10.9676 | peaks_peak_368 | Peak_chr1_172 | 386 | NA | NA | | | | | | | | | | | | |
| NC_002505 | 1786141 | 1786893 | 753 | 1786535 | 388 | 75.28632 | 2.99787 | 71.048 | peaks_peak_370 | Peak_chr1_173 | 4 | VC1655 | VC1653 | 1786971 | 1786270 | 436 | 265 | 1.6557 | 0.0000 | 0.9548 | 0.0000 | Y | Y | 46 | Up |
| NC_002505 | 1793963 | 1794261 | 299 | 1794082 | 229 | 13.62516 | 1.7096 | 11.93066 | peaks_peak_371 | Peak_chr1_174 | 362 | VC1661 | NA | 1794166 | | 84 | | 0.2336 | 0.6794 | | | N | | | |
| NC_002505 | 1794468 | 1795203 | 736 | 1794757 | 374 | 66.29622 | 2.82708 | 62.91714 | peaks_peak_372 | Peak_chr1_175 | 23 | VC1663 | VC1662 | 1794806 | 1794659 | 49 | 98 | 2.4647 | 0.0000 | -0.3044 | 0.2693 | Y | N | | Up |
| NC_002505 | 1801774 | 1802087 | 314 | 1801931 | 230 | 14.49445 | 1.74195 | 12.77098 | peaks_peak_374 | Peak_chr1_176 | 346 | cls | VC1669 | 1802327 | 1801771 | 396 | 160 | 0.3048 | 0.0218 | -0.2306 | 0.5191 | N | N | | |
| NC_002505 | 1818115 | 1818376 | 262 | 1818232 | 237 | 14.9176 | 1.7429 | 13.18049 | peaks_peak_375 | Peak_chr1_177 | 333 | VC1684 | NA | 1818269 | | 37 | | -0.2241 | 0.0846 | | | N | | | |
| NC_002505 | 1848438 | 1848701 | 264 | 1848582 | 229 | 12.84147 | 1.67818 | 11.17226 | peaks_peak_377 | Peak_chr1_178 | 382 | VC1713 | VC1712 | 1848654 | 1848576 | 72 | 6 | 0.1025 | 0.8002 | 1.5269 | 0.0000 | N | Y | 47 | Up |
| NC_002505 | 1856281 | 1856625 | 345 | 1856448 | 291 | 32.11894 | 2.18143 | 29.9595 | peaks_peak_378 | Peak_chr1_179 | 173 | VC1718 | NA | 1856575 | | 127 | | -0.4343 | 0.0664 | | | N | | | |
| NC_002505 | 1860133 | 1861124 | 992 | 1860800 | 392 | 75.10676 | 2.9737 | 70.89553 | peaks_peak_379 | Peak_chr1_180 | 5 | VC1722 | NA | 1860890 | | 90 | | 1.5473 | 0.0000 | | | Y | | 48 | Up |
| NC_002505 | 1866617 | 1867533 | 917 | 1866772 | 249 | 19.62275 | 1.88878 | 17.76006 | peaks_peak_380 | Peak_chr1_181 | 271 | NA | glgC (NC_002506 637338.. | | 1866550 | | 222 | | | -2.3358 | 0.0000 | | Y | 49 | Down |
| NC_002505 | 1871146 | 1871403 | 258 | 1871288 | 232 | 15.1064 | 1.76145 | 13.36355 | peaks_peak_381 | Peak_chr1_182 | 330 | NA | NA | | | | | | | | | | | | |
| NC_002505 | 1884634 | 1886001 | 1368 | 1885471 | 395 | 77.61683 | 3.02413 | 73.12509 | peaks_peak_385 | Peak_chr1_183 | 3 | NA | NA | | | | | | | | | | | | |
| NC_002505 | 1902135 | 1903009 | 875 | 1902587 | 268 | 28.26749 | 2.13243 | 26.1974 | peaks_peak_387 | Peak_chr1_184 | 195 | NA | NA | | | | | | | | | | | | |
| NC_002505 | 1904036 | 1905934 | 1899 | 1904429 | 371 | 67.54165 | 2.87423 | 64.07333 | peaks_peak_388 | Peak_chr1_185 | 20 | NA | NA | | | | | | | | | | | | |
| NC_002505 | 1907479 | 1907775 | 297 | 1907618 | 232 | 14.66618 | 1.74422 | 12.93692 | peaks_peak_389 | Peak_chr1_186 | 339 | NA | NA | | | | | | | | | | | | |
| NC_002505 | 1919382 | 1919980 | 599 | 1919665 | 221 | 13.60292 | 1.72636 | 11.90901 | peaks_peak_392 | Peak_chr1_187 | 363 | NA | NA | | | | | | | | | | | | |
| NC_002505 | 1924876 | 1925272 | 397 | 1925049 | 245 | 22.21598 | 1.99424 | 20.28794 | peaks_peak_393 | Peak_chr1_188 | 240 | NA | VC1774 | | 1925049 | | 0 | | | 0.6883 | 0.0023 | | N | | |
| NC_002505 | 1926525 | 1927012 | 488 | 1926646 | 223 | 14.25346 | 1.74837 | 12.53788 | peaks_peak_394 | Peak_chr1_189 | 350 | NA | NA | | | | | | | | | | | | |
| NC_002505 | 1934823 | 1935255 | 433 | 1934950 | 256 | 20.72513 | 1.90961 | 18.83448 | peaks_peak_397 | Peak_chr1_190 | 257 | NA | NA | | | | | | | | | | | | |
| NC_002505 | 1935561 | 1935943 | 383 | 1935674 | 223 | 12.27462 | 1.66721 | 10.62531 | peaks_peak_398 | Peak_chr1_191 | 398 | VC1786 | NA | 1936121 | | 447 | | -0.6797 | 0.5885 | | | N | | | |
| NC_002505 | 1940285 | 1940580 | 296 | 1940423 | 220 | 11.81862 | 1.65424 | 10.18675 | peaks_peak_401 | Peak_chr1_192 | 417 | NA | VC1792 | | 1940273 | | 150 | | | -0.1961 | 0.5202 | | N | | |
| NC_002505 | 1940817 | 1942085 | 1269 | 1941333 | 374 | 64.14154 | 2.77026 | 60.90097 | peaks_peak_402 | Peak_chr1_193 | 30 | NA | VC1794 | | 1941332 | | 1 | | | -0.4981 | 0.2141 | | N | | |
| NC_002505 | 1946678 | 1948278 | 1601 | 1947249 | 325 | 48.4058 | 2.5445 | 45.82543 | peaks_peak_403 | Peak_chr1_194 | 81 | NA | VC1800 | | 1947122 | | 127 | | | 0.2482 | 0.5164 | | N | | |
| NC_002505 | 1950284 | 1950600 | 317 | 1950429 | 223 | 13.85928 | 1.73231 | 12.15692 | peaks_peak_405 | Peak_chr1_195 | 354 | NA | NA | | | | | | | | | | | | |
| NC_002505 | 1952639 | 1953348 | 710 | 1952854 | 362 | 68.41977 | 2.94244 | 64.8958 | peaks_peak_406 | Peak_chr1_196 | 17 | NA | NA | | | | | | | | | | | | |
| NC_002505 | 1958958 | 1959516 | 559 | 1959334 | 234 | 14.02983 | 1.71496 | 12.32189 | peaks_peak_407 | Peak_chr1_197 | 351 | NA | VC1817 | | 1958964 | | 370 | | | -0.8076 | 0.0572 | | N | | |
| NC_002505 | 1966906 | 1967560 | 655 | 1967065 | 239 | 15.49718 | 1.76061 | 13.74299 | peaks_peak_408 | Peak_chr1_198 | 323 | VC1826 | VC1825 | 1967400 | 1967028 | 335 | 37 | -0.2674 | 0.8582 | -0.3700 | 0.5777 | N | N | | |
| NC_002505 | 1970976 | 1971512 | 537 | 1971352 | 253 | 20.17913 | 1.89822 | 18.30221 | peaks_peak_409 | Peak_chr1_199 | 265 | NA | VC1828 | | 1971155 | | 197 | | | 0.4839 | 0.1804 | | N | | |
| NC_002505 | 1977073 | 1978060 | 988 | 1977293 | 329 | 44.92846 | 2.42888 | 42.45257 | peaks_peak_410 | Peak_chr1_200 | 102 | NA | VC1835 | | 1977088 | | 205 | | | 0.9554 | 0.0000 | | Y | 50 | Up |
| NC_002505 | 1980655 | 1981227 | 573 | 1980811 | 293 | 32.05616 | 2.17303 | 29.89841 | peaks_peak_411 | Peak_chr1_201 | 174 | NA | VC1839 | | 1980701 | | 110 | | | 1.5247 | 0.0000 | | Y | 51 | Up |
| NC_002505 | 1984013 | 1984938 | 926 | 1984283 | 365 | 61.15433 | 2.73013 | 58.09334 | peaks_peak_412 | Peak_chr1_202 | 39 | NA | NA | | | | | | | | | | | | |
| NC_002505 | 2003225 | 2003449 | 225 | 2003345 | 228 | 13.69969 | 1.71473 | 12.00307 | peaks_peak_416 | Peak_chr1_203 | 360 | NA | NA | | | | | | | | | | | | |
| NC_002505 | 2004154 | 2005117 | 964 | 2004306 | 263 | 23.59181 | 1.98955 | 21.63222 | peaks_peak_417 | Peak_chr1_204 | 228 | NA | NA | | | | | | | | | | | | |
| NC_002505 | 2009528 | 2009970 | 443 | 2009796 | 314 | 34.02568 | 2.16725 | 31.82214 | peaks_peak_418 | Peak_chr1_205 | 160 | NA | NA | | | | | | | | | | | | |
| NC_002505 | 2014072 | 2014464 | 393 | 2014220 | 243 | 16.81041 | 1.80078 | 15.02028 | peaks_peak_419 | Peak_chr1_206 | 300 | NA | NA | | | | | | | | | | | | |
| NC_002505 | 2025471 | 2026004 | 534 | 2025618 | 232 | 13.07351 | 1.68141 | 11.39682 | peaks_peak_420 | Peak_chr1_207 | 373 | VC1880 | NA | 2025912 | | 294 | | -0.8633 | 0.0005 | | | Y | | 52 | Down |
| NC_002505 | 2029629 | 2029960 | 332 | 2029796 | 296 | 32.25659 | 2.16967 | 30.0943 | peaks_peak_421 | Peak_chr1_208 | 172 | VC1885 | VC1884 | 2029865 | 2029701 | 69 | 95 | 0.1097 | 0.6648 | 1.2303 | 0.0000 | N | Y | 53 | Up |
| NC_002505 | 2044258 | 2044513 | 256 | 2044386 | 224 | 12.04996 | 1.65591 | 10.4093 | peaks_peak_423 | Peak_chr1_209 | 405 | VC1898 | VC1897 | 2044494 | 2044261 | 108 | 125 | -0.2244 | 0.6910 | -0.8107 | 0.0000 | N | Y | 54 | Down |
| NC_002505 | 2048877 | 2049556 | 680 | 2049272 | 382 | 66.09913 | 2.78656 | 62.73773 | peaks_peak_424 | Peak_chr1_210 | 25 | nhaB | VC1900 | 2049552 | 2048907 | 280 | 365 | -0.3596 | 0.0383 | 0.7133 | 0.0003 | N | Y | | Up |
| NC_002505 | 2054694 | 2055066 | 373 | 2054821 | 251 | 17.78704 | 1.81775 | 15.97208 | peaks_peak_425 | Peak_chr1_211 | 287 | NA | VC1903 | | 2054702 | | 119 | | | 2.1667 | 0.0000 | | Y | 55 | Up |
| NC_002505 | 2056865 | 2057197 | 333 | 2057030 | 262 | 19.9925 | 1.86906 | 18.12009 | peaks_peak_426 | Peak_chr1_212 | 266 | NA | NA | | | | | | | | | | | | |
| NC_002505 | 2081505 | 2082065 | 561 | 2081776 | 323 | 41.88313 | 2.36384 | 39.4894 | peaks_peak_430 | Peak_chr1_213 | 117 | NA | NA | | | | | | | | | | | | |
| NC_002505 | 2083971 | 2084520 | 550 | 2084160 | 320 | 41.00801 | 2.34928 | 38.63639 | peaks_peak_431 | Peak_chr1_214 | 119 | VC1933 | VC1931 | 2084346 | 2083843 | 186 | 317 | 0.0916 | 0.7944 | -0.0296 | 0.8813 | N | N | | |
| NC_002505 | 2088752 | 2089286 | 535 | 2088930 | 245 | 16.74436 | 1.79368 | 14.95598 | peaks_peak_433 | Peak_chr1_215 | 302 | VC1938 | NA | 2089288 | | 358 | | 0.1500 | 0.7063 | | | N | | | |
| NC_002505 | 2093453 | 2093772 | 320 | 2093614 | 319 | 39.02094 | 2.29546 | 36.69834 | peaks_peak_434 | Peak_chr1_216 | 130 | NA | tRNA-Arg-2 | | 2093396 | | 218 | | | NA | | | | | |
| NC_002505 | 2097286 | 2097620 | 335 | 2097446 | 280 | 26.2572 | 2.02954 | 24.23336 | peaks_peak_435 | Peak_chr1_217 | 212 | NA | NA | | | | | | | | | | | | |
| NC_002505 | 2098708 | 2098946 | 239 | 2098837 | 227 | 12.93994 | 1.68624 | 11.26735 | peaks_peak_437 | Peak_chr1_218 | 376 | NA | NA | | | | | | | | | | | | |
| NC_002505 | 2110710 | 2111106 | 397 | 2110857 | 266 | 21.45356 | 1.90893 | 19.54412 | peaks_peak_438 | Peak_chr1_219 | 246 | minC | VC1956 | 2111116 | 2110613 | 259 | 244 | 1.0571 | 0.0000 | 1.9153 | 0.0000 | Y | Y | 56 | Up |
| NC_002505 | 2112731 | 2113460 | 730 | 2113103 | 399 | 64.40525 | 2.67552 | 61.15176 | peaks_peak_439 | Peak_chr1_220 | 28 | VC1962 | NA | 2113168 | | 65 | | 5.2641 | 0.0000 | | | Y | | 57 | Up |
| NC_002505 | 2150420 | 2150680 | 261 | 2150572 | 237 | 12.03379 | 1.63081 | 10.39366 | peaks_peak_442 | Peak_chr1_221 | 406 | NA | VC1996 | | 2150433 | | 139 | | | -0.9958 | 0.0025 | | N | | |
| NC_002505 | 2151626 | 2152376 | 751 | 2152020 | 280 | 25.82623 | 2.0157 | 23.81227 | peaks_peak_443 | Peak_chr1_222 | 213 | VC2000 | VC1998 | 2152265 | 2151965 | 245 | 55 | 0.1812 | 0.4990 | 0.2723 | 0.2113 | N | N | | |
| NC_002505 | 2163389 | 2163841 | 453 | 2163682 | 232 | 13.5519 | 1.70036 | 11.8597 | peaks_peak_448 | Peak_chr1_223 | 364 | VC2011 | VC2009 | 2163877 | 2163645 | 195 | 37 | 0.8284 | 0.0021 | 0.1484 | 0.7883 | N | N | | |
| NC_002505 | 2167590 | 2169159 | 1570 | 2168572 | 386 | 70.45755 | 2.88142 | 66.79388 | peaks_peak_449 | Peak_chr1_224 | 13 | NA | VC2013 | | 2168149 | | 423 | | | -0.1747 | 0.6669 | | N | | |
| NC_002505 | 2178714 | 2179002 | 289 | 2178857 | 225 | 11.6279 | 1.63651 | 10.00375 | peaks_peak_450 | Peak_chr1_225 | 424 | maf | rpmF | 2179203 | 2178496 | 346 | 361 | | | -0.2049 | 0.7019 | | N | | |
| NC_002505 | 2179714 | 2180002 | 289 | 2179857 | 230 | 12.48497 | 1.66185 | 10.82839 | peaks_peak_451 | Peak_chr1_226 | 391 | NA | NA | | | | | | | | | | | | |
| NC_002505 | 2190152 | 2191083 | 932 | 2190742 | 379 | 62.88959 | 2.7166 | 59.7188 | peaks_peak_453 | Peak_chr1_227 | 36 | VC2035 | VC2033 | 2190929 | 2190302 | 187 | 440 | 0.5594 | 0.0803 | 1.3431 | 0.0000 | N | Y | 58 | Up |
| NC_002505 | 2192242 | 2193699 | 1458 | 2192947 | 389 | 68.01542 | 2.80548 | 64.51865 | peaks_peak_454 | Peak_chr1_228 | 18 | VC2037 | VC2036 | 2193194 | 2192769 | 247 | 178 | -0.8274 | 0.0002 | -0.1946 | 0.6617 | Y | N | 59 | Down |
| NC_002505 | 2228853 | 2229429 | 577 | 2229140 | 341 | 45.22181 | 2.39411 | 42.73884 | peaks_peak_459 | Peak_chr1_229 | 101 | NA | NA | | | | | | | | | | | | |
| NC_002505 | 2255133 | 2255592 | 460 | 2255340 | 306 | 32.89452 | 2.15807 | 30.71711 | peaks_peak_464 | Peak_chr1_230 | 166 | NA | VC2093 | | 2255278 | | 62 | | | -0.3458 | 0.3812 | | N | | |
| NC_002505 | 2258349 | 2258668 | 320 | 2258525 | 252 | 15.47674 | 1.73218 | 13.72293 | peaks_peak_465 | Peak_chr1_231 | 324 | VC2097 | VC2096 | 2258535 | 2258442 | 10 | 83 | 0.6634 | 0.0008 | -0.4500 | 0.0705 | Y | N | | Up |
| NC_002505 | 2262885 | 2263482 | 598 | 2263274 | 353 | 44.33174 | 2.33079 | 41.87215 | peaks_peak_466 | Peak_chr1_232 | 106 | fur | VC2105 | 2263275 | 2262909 | 1 | 365 | -0.2507 | 0.0890 | -0.4504 | 0.2441 | N | N | | |
| NC_002505 | 2307372 | 2307875 | 504 | 2307741 | 230 | 11.76643 | 1.63282 | 10.13671 | peaks_peak_467 | Peak_chr1_233 | 418 | bcp | dapA | 2308172 | 2307337 | 431 | 404 | -0.0699 | 0.7183 | 0.4647 | 0.0009 | N | Y | 60 | Up |
| NC_002505 | 2311812 | 2312244 | 433 | 2311927 | 250 | 16.1283 | 1.76001 | 14.35619 | peaks_peak_468 | Peak_chr1_234 | 313 | VC2164 | VC2162 | 2312280 | 2311867 | 353 | 60 | 0.8573 | 0.0000 | 1.9896 | 0.0000 | Y | Y | 61 | Up |
| NC_002505 | 2316422 | 2316843 | 422 | 2316566 | 288 | 26.5921 | 2.01805 | 24.56085 | peaks_peak_469 | Peak_chr1_235 | 208 | VC2171 | VC2168 | 2316680 | 2316459 | 114 | 107 | 0.2981 | 0.7042 | 1.4188 | 0.0000 | N | Y | 62 | Up |
| NC_002505 | 2341810 | 2342025 | 216 | 2341913 | 237 | 12.03379 | 1.63081 | 10.39366 | peaks_peak_470 | Peak_chr1_236 | 406 | NA | flgI | | 2341497 | | 416 | | | -2.2822 | 0.0000 | | Y | 63 | Down |
| NC_002505 | 2350313 | 2350657 | 345 | 2350498 | 256 | 16.99026 | 1.77808 | 15.19549 | peaks_peak_471 | Peak_chr1_237 | 296 | NA | NA | | | | | | | | | | | | |
| NC_002505 | 2359620 | 2359826 | 207 | 2359706 | 249 | 12.81151 | 1.63963 | 11.14335 | peaks_peak_472 | Peak_chr1_238 | 383 | NA | NA | | | | | | | | | | | | |
| NC_002505 | 2364278 | 2364957 | 680 | 2364620 | 379 | 60.64729 | 2.65923 | 57.61192 | peaks_peak_473 | Peak_chr1_239 | 42 | NA | NA | | | | | | | | | | | | |
| NC_002505 | 2365783 | 2366768 | 986 | 2366407 | 356 | 50.36884 | 2.47932 | 47.72803 | peaks_peak_475 | Peak_chr1_240 | 75 | NA | VC2213 | | 2366258 | | 149 | | | 2.8976 | 0.0000 | | Y | 64 | Up |
| NC_002505 | 2375759 | 2376218 | 460 | 2375948 | 279 | 22.34035 | 1.90619 | 20.40896 | peaks_peak_476 | Peak_chr1_241 | 238 | VC2222 | tRNA-Pro-4 | 2376252 | 2375700 | 304 | 248 | -0.3475 | 0.0324 | -0.1866 | 0.5366 | N | N | | |
| NC_002505 | 2389003 | 2389297 | 295 | 2389145 | 258 | 13.78647 | 1.65985 | 12.08658 | peaks_peak_477 | Peak_chr1_242 | 357 | NA | VC2233 | | 2388767 | | 378 | | | -0.3757 | 0.0152 | | N | | |
| NC_002505 | 2397666 | 2398537 | 872 | 2398059 | 390 | 64.22018 | 2.70584 | 60.97736 | peaks_peak_478 | Peak_chr1_243 | 29 | NA | VC2244 | | 2397677 | | 382 | | | -0.2257 | 0.4541 | | N | | |
| NC_002505 | 2399980 | 2400326 | 347 | 2400153 | 328 | 37.68598 | 2.22895 | 35.39604 | peaks_peak_479 | Peak_chr1_244 | 139 | NA | NA | | | | | | | | | | | | |
| NC_002505 | 2409899 | 2410204 | 306 | 2410063 | 288 | 21.05943 | 1.84509 | 19.16049 | peaks_peak_480 | Peak_chr1_245 | 252 | NA | VC2253 | | 2409929 | | 134 | | | 0.9658 | 0.0001 | | Y | 65 | Up |
| NC_002505 | 2414410 | 2414661 | 252 | 2414536 | 263 | 16.50026 | 1.74641 | 14.71828 | peaks_peak_481 | Peak_chr1_246 | 306 | NA | pyrH | | 2414229 | | 307 | | | 0.0503 | 0.9275 | | N | | |
| NC_002505 | 2429431 | 2430257 | 827 | 2430080 | 314 | 34.20999 | 2.17257 | 32.00211 | peaks_peak_482 | Peak_chr1_247 | 158 | NA | VC2275 | | 2429904 | | 176 | | | 1.1229 | 0.0009 | | Y | 66 | Up |
| NC_002505 | 2432558 | 2433519 | 962 | 2433268 | 392 | 66.52154 | 2.75525 | 63.12645 | peaks_peak_483 | Peak_chr1_248 | 22 | NA | VC2278 | | 2433176 | | 92 | | | -0.8400 | 0.0004 | | Y | 67 | Down |
| NC_002505 | 2439507 | 2439925 | 419 | 2439751 | 349 | 48.61334 | 2.45767 | 46.02553 | peaks_peak_484 | Peak_chr1_249 | 80 | NA | VC2283 | | 2439578 | | 173 | | | 0.8122 | 0.0000 | | Y | 68 | Up |
| NC_002505 | 2441165 | 2442264 | 1100 | 2441834 | 301 | 30.7536 | 2.10901 | 28.62682 | peaks_peak_485 | Peak_chr1_250 | 180 | NA | NA | | | | | | | | | | | | |
| NC_002505 | 2444672 | 2444958 | 287 | 2444812 | 244 | 13.3911 | 1.67048 | 11.704 | peaks_peak_486 | Peak_chr1_251 | 367 | NA | VC2289 | | 2444777 | | 35 | | | -0.2884 | 0.2684 | | N | | |
| NC_002505 | 2447236 | 2447614 | 379 | 2447431 | 307 | 27.771 | 2.0044 | 25.71236 | peaks_peak_488 | Peak_chr1_252 | 202 | NA | VC2292 | | 2447419 | | 12 | | | 0.0044 | 0.9897 | | N | | |
| NC_002505 | 2450804 | 2451408 | 605 | 2450947 | 264 | 16.70915 | 1.75165 | 14.9219 | peaks_peak_489 | Peak_chr1_253 | 303 | NA | VC2295 | | 2450820 | | 127 | | | -0.0656 | 0.8512 | | N | | |
| NC_002505 | 2463857 | 2464193 | 337 | 2464037 | 282 | 22.93151 | 1.91792 | 20.98681 | peaks_peak_491 | Peak_chr1_254 | 231 | NA | mltA | | 2463993 | | 44 | | | 1.4388 | 0.0000 | | Y | 69 | Up |
| NC_002505 | 2466074 | 2466329 | 256 | 2466183 | 241 | 12.59403 | 1.64549 | 10.93357 | peaks_peak_492 | Peak_chr1_255 | 388 | VC2318 | NA | 2466301 | | 118 | | 0.1551 | 0.8403 | | | N | | | |
| NC_002505 | 2469735 | 2469958 | 224 | 2469850 | 266 | 16.22054 | 1.73089 | 14.44592 | peaks_peak_493 | Peak_chr1_256 | 310 | NA | NA | | | | | | | | | | | | |
| NC_002505 | 2493348 | 2494176 | 829 | 2494013 | 267 | 17.26072 | 1.76455 | 15.45893 | peaks_peak_496 | Peak_chr1_257 | 293 | NA | NA | | | | | | | | | | | | |
| NC_002505 | 2506747 | 2507075 | 329 | 2506934 | 255 | 16.09884 | 1.74846 | 14.3277 | peaks_peak_499 | Peak_chr1_258 | 314 | NA | NA | | | | | | | | | | | | |
| NC_002505 | 2513015 | 2514934 | 1920 | 2513763 | 391 | 63.29185 | 2.67884 | 60.09526 | peaks_peak_501 | Peak_chr1_259 | 33 | VC2357 | NA | 2513867 | | 104 | | 0.4818 | 0.1161 | | | N | | | |
| NC_002505 | 2516932 | 2517455 | 524 | 2517313 | 246 | 12.32225 | 1.62629 | 10.6713 | peaks_peak_502 | Peak_chr1_260 | 395 | NA | NA | | | | | | | | | | | | |
| NC_002505 | 2547537 | 2547916 | 380 | 2547705 | 351 | 42.22856 | 2.28192 | 39.82602 | peaks_peak_505 | Peak_chr1_261 | 115 | NA | NA | | | | | | | | | | | | |
| NC_002505 | 2549158 | 2550239 | 1082 | 2549465 | 226 | 13.64807 | 1.71702 | 11.95295 | peaks_peak_506 | Peak_chr1_262 | 361 | VC2385 | VC2384 | 2549642 | 2549180 | 177 | 285 | -0.1587 | 0.4909 | -0.0681 | 0.7964 | N | N | | |
| NC_002505 | 2551522 | 2552640 | 1119 | 2552287 | 252 | 19.78759 | 1.88689 | 17.92083 | peaks_peak_508 | Peak_chr1_263 | 269 | NA | NA | | | | | | | | | | | | |
| NC_002505 | 2558398 | 2558842 | 445 | 2558585 | 322 | 33.66742 | 2.13403 | 31.47216 | peaks_peak_509 | Peak_chr1_264 | 161 | NA | VC2390 | | 2558435 | | 150 | | | 0.3021 | 0.4253 | | N | | |
| NC_002505 | 2560731 | 2561108 | 378 | 2560933 | 362 | 44.89297 | 2.31753 | 42.41769 | peaks_peak_510 | Peak_chr1_265 | 103 | NA | NA | | | | | | | | | | | | |
| NC_002505 | 2564882 | 2565175 | 294 | 2565034 | 263 | 16.88326 | 1.75969 | 15.09127 | peaks_peak_511 | Peak_chr1_266 | 297 | NA | lpxC | | 2564949 | | 85 | | | 0.2450 | 0.2467 | | N | | |
| NC_002505 | 2572267 | 2572771 | 505 | 2572637 | 273 | 14.90072 | 1.67318 | 13.16413 | peaks_peak_513 | Peak_chr1_267 | 334 | NA | VC2402 | | 2572211 | | 426 | | | 1.7902 | 0.0000 | | Y | 70 | Up |
| NC_002505 | 2577059 | 2577416 | 358 | 2577232 | 316 | 31.51037 | 2.0893 | 29.36487 | peaks_peak_514 | Peak_chr1_268 | 177 | NA | NA | | | | | | | | | | | | |
| NC_002505 | 2580487 | 2580826 | 340 | 2580693 | 290 | 21.60489 | 1.85786 | 19.69174 | peaks_peak_515 | Peak_chr1_269 | 245 | NA | mraW | | 2580534 | | 159 | | | 1.8377 | 0.0000 | | Y | 71 | Up |
| NC_002505 | 2588413 | 2588649 | 237 | 2588529 | 248 | 13.43775 | 1.66485 | 11.74939 | peaks_peak_516 | Peak_chr1_270 | 366 | VC2416 | pdhR | 2588859 | 2588375 | 330 | 154 | -0.2601 | 0.7162 | 0.4668 | 0.1832 | N | N | | |
| NC_002505 | 2590329 | 2590821 | 493 | 2590597 | 347 | 42.80388 | 2.30935 | 40.38606 | peaks_peak_517 | Peak_chr1_271 | 112 | NA | NA | | | | | | | | | | | | |
| NC_002505 | 2607350 | 2607804 | 455 | 2607543 | 314 | 31.13592 | 2.08395 | 28.9999 | peaks_peak_519 | Peak_chr1_272 | 179 | NA | VC2431 | | 2607478 | | 65 | | | 0.7890 | 0.0000 | | Y | 72 | Up |
| NC_002505 | 2633970 | 2634237 | 268 | 2634115 | 268 | 15.44893 | 1.70063 | 13.69605 | peaks_peak_525 | Peak_chr1_273 | 326 | VC2453 | NA | 2634597 | | 482 | | 0.0734 | 0.7570 | | | N | | | |
| NC_002505 | 2649097 | 2649669 | 573 | 2649303 | 274 | 17.6103 | 1.76239 | 15.79966 | peaks_peak_526 | Peak_chr1_274 | 291 | NA | VC2466 | | 2649160 | | 143 | | | 0.8887 | 0.0000 | | Y | 73 | Up |
| NC_002505 | 2653899 | 2654509 | 611 | 2654073 | 311 | 29.79268 | 2.05313 | 27.68839 | peaks_peak_527 | Peak_chr1_275 | 186 | VC2473 | NA | 2654268 | | 195 | | 2.3058 | 0.0000 | | | Y | | 74 | Up |
| NC_002505 | 2664714 | 2665004 | 291 | 2664854 | 259 | 16.78023 | 1.76432 | 14.99078 | peaks_peak_528 | Peak_chr1_276 | 301 | NA | NA | | | | | | | | | | | | |
| NC_002505 | 2686261 | 2686551 | 291 | 2686397 | 275 | 19.89759 | 1.83589 | 18.02765 | peaks_peak_534 | Peak_chr1_277 | 267 | VC2501 | VC2500 | 2686560 | 2686366 | 163 | 31 | 0.2403 | 0.0346 | 0.1255 | 0.6859 | N | N | | |
| NC_002505 | 2697164 | 2697449 | 286 | 2697326 | 258 | 14.35616 | 1.68032 | 12.6374 | peaks_peak_535 | Peak_chr1_278 | 349 | NA | NA | | | | | | | | | | | | |
| NC_002505 | 2701459 | 2702002 | 544 | 2701644 | 344 | 40.16289 | 2.24832 | 37.81338 | peaks_peak_538 | Peak_chr1_279 | 121 | NA | NA | | | | | | | | | | | | |
| NC_002505 | 2704958 | 2705447 | 490 | 2705260 | 371 | 46.49415 | 2.3308 | 43.97294 | peaks_peak_540 | Peak_chr1_280 | 93 | NA | VC2517 | | 2704870 | | 390 | | | 3.2672 | 0.0000 | | Y | 75 | Up |
| NC_002505 | 2706835 | 2707379 | 545 | 2707059 | 285 | 20.74975 | 1.84172 | 18.8584 | peaks_peak_541 | Peak_chr1_281 | 255 | VC2522 | VC2520 | 2707506 | 2706949 | 447 | 110 | 0.0955 | 0.7912 | 2.0590 | 0.0000 | N | Y | 76 | Up |
| NC_002505 | 2721640 | 2721864 | 225 | 2721777 | 246 | 11.92308 | 1.61104 | 10.28713 | peaks_peak_542 | Peak_chr1_282 | 412 | NA | thiP | | 2721334 | | 443 | | | 0.3441 | 0.5428 | | N | | |
| NC_002505 | 2749395 | 2749721 | 327 | 2749586 | 282 | 18.91072 | 1.78914 | 17.06665 | peaks_peak_543 | Peak_chr1_283 | 277 | VC2568 | VC2567 | 2749721 | 2749510 | 135 | 76 | -0.5161 | 0.0078 | 1.1510 | 0.0000 | N | Y | | Up |
| NC_002505 | 2755117 | 2755448 | 332 | 2755296 | 281 | 19.11284 | 1.79767 | 17.26369 | peaks_peak_544 | Peak_chr1_284 | 276 | NA | NA | | | | | | | | | | | | |
| NC_002505 | 2805248 | 2805740 | 493 | 2805484 | 362 | 47.01187 | 2.37168 | 44.47338 | peaks_peak_547 | Peak_chr1_285 | 92 | VC2635 | NA | 2805704 | | 220 | | 2.3417 | 0.0000 | | | Y | | 77 | Up |
| NC_002505 | 2818278 | 2818581 | 304 | 2818443 | 304 | 26.87741 | 1.98549 | 24.83968 | peaks_peak_548 | Peak_chr1_286 | 205 | VC2646 | NA | 2818512 | | 69 | | 0.7227 | 0.0151 | | | N | | | |
| NC_002505 | 2826479 | 2826900 | 422 | 2826738 | 272 | 15.39293 | 1.69162 | 13.6417 | peaks_peak_550 | Peak_chr1_287 | 327 | VC2656 | VC2655 | 2826752 | 2826443 | 14 | 295 | 0.0008 | 0.9988 | 0.0437 | 0.8876 | N | N | | |
| NC_002505 | 2830424 | 2830761 | 338 | 2830609 | 347 | 38.72068 | 2.20156 | 36.40562 | peaks_peak_551 | Peak_chr1_288 | 132 | VC2661 | NA | 2830790 | | 181 | | -0.2254 | 0.3103 | | | N | | | |
| NC_002505 | 2832131 | 2832745 | 615 | 2832496 | 393 | 55.9684 | 2.49314 | 53.12749 | peaks_peak_552 | Peak_chr1_289 | 57 | NA | VC2662 | | 2832424 | | 72 | | | 2.3745 | 0.0000 | | Y | 78 | Up |
| NC_002505 | 2836481 | 2836846 | 366 | 2836654 | 294 | 22.76004 | 1.88483 | 20.8193 | peaks_peak_554 | Peak_chr1_290 | 235 | NA | VC2668 | | 2836640 | | 14 | | | -0.2271 | 0.2272 | | N | | |

| | | | | | | | | | | | | | | | | | | | | | | | | | |
|---|---|---|---|---|---|---|---|---|---|---|---|---|---|---|---|---|---|---|---|---|---|---|---|---|---|
| NC_002505 | 2845054 | 2845381 | 328 | 2845232 | 294 | 22.69947 | 1.88297 | 20.75995 | peaks_peak_555 | Peak_chr1_291 | 236 | NA | VC2677 | | 2845045 | | 187 | | | 0.3891 | 0.1567 | | N | | |
| NC_002505 | 2855130 | 2855952 | 823 | 2855511 | 388 | 54.50028 | 2.47415 | 51.71504 | peaks_peak_556 | Peak_chr1_292 | 62 | glpX | VC2686 | 2855699 | 2855378 | 188 | 133 | -0.2366 | 0.5902 | -1.1476 | 0.0000 | N | Y | 79 | Down |
| NC_002505 | 2864607 | 2865242 | 636 | 2864824 | 315 | 29.25192 | 2.02731 | 27.15967 | peaks_peak_558 | Peak_chr1_293 | 190 | aspA | VC2697 | 2864958 | 2864713 | 134 | 111 | -0.9619 | 0.0343 | 1.3859 | 0.0000 | N | Y | | Up |
| NC_002505 | 2892820 | 2893190 | 371 | 2893002 | 339 | 36.97895 | 2.17763 | 34.70722 | peaks_peak_561 | Peak_chr1_294 | 143 | NA | VC2717 | | 2892951 | | 51 | | | -0.0782 | 0.7435 | | N | | |
| NC_002505 | 2909856 | 2910160 | 305 | 2909995 | 288 | 19.75394 | 1.80404 | 17.8879 | peaks_peak_562 | Peak_chr1_295 | 270 | VC2738 | NA | 2910015 | | 20 | | 0.0923 | 0.8803 | | | N | | | |
| NC_002505 | 2918258 | 2919188 | 931 | 2918675 | 393 | 56.04652 | 2.49501 | 53.20255 | peaks_peak_564 | Peak_chr1_296 | 55 | *glnA* | VC2744 | 2919083 | 2918595 | 408 | 80 | 0.3242 | 0.5365 | -0.0881 | 0.8636 | N | N | | |
| NC_002505 | 2931481 | 2932034 | 554 | 2931611 | 264 | 11.63325 | 1.57259 | 10.00887 | peaks_peak_567 | Peak_chr1_297 | 422 | NA | *tRNA-Lys-1* | | 2931580 | | 31 | | | -0.0522 | 0.9507 | | N | | |
| NC_002505 | 2933226 | 2934008 | 783 | 2933453 | 310 | 12.87404 | 1.55504 | 11.2038 | peaks_peak_568 | Peak_chr1_298 | 381 | NA | 16Sg | | 2933279 | | 174 | | | -0.8103 | 0.1363 | | N | | |
| NC_002505 | 2940101 | 2940918 | 818 | 2940551 | 394 | 52.70198 | 2.41206 | 49.98661 | peaks_peak_571 | Peak_chr1_299 | 68 | NA | NA | | | | | | | | | | | | |
| NC_002505 | 2941937 | 2942457 | 521 | 2942239 | 370 | 47.48534 | 2.35859 | 44.93339 | peaks_peak_573 | Peak_chr1_300 | 86 | fadB | VC2757 | 2942322 | 2942111 | 83 | 128 | -1.0935 | 0.1235 | -1.0446 | 0.0000 | N | Y | 80 | Down |
| NC_002505 | 2949398 | 2949829 | 432 | 2949561 | 268 | 14.7944 | 1.67807 | 13.06071 | peaks_peak_574 | Peak_chr1_301 | 336 | NA | glmU | | 2949438 | | 123 | | | 0.5785 | 0.0292 | | N | | |
| NC_002505 | 2954151 | 2954488 | 338 | 2954354 | 307 | 24.69427 | 1.91413 | 22.70812 | peaks_peak_575 | Peak_chr1_302 | 223 | NA | VC2766 | | 2954046 | | 308 | | | 0.3347 | 0.0953 | | N | | |
| NC_002505 | 2960414 | 2961069 | 656 | 2960692 | 384 | 54.45431 | 2.48628 | 51.67185 | peaks_peak_578 | Peak_chr1_303 | 64 | NA | NA | | | | | | | | | | | | |
| NC_002506 | 73 | 906 | 834 | 598 | 281 | 24.5601 | 1.97245 | 22.57747 | peaks_peak_579 | Peak_chr2_1 | 224 | NA | NA | | | | | | | | | | | | |
| NC_002506 | 2889 | 3565 | 677 | 3223 | 395 | 62.51577 | 2.64456 | 59.3657 | peaks_peak_581 | Peak_chr2_2 | 37 | NA | NA | | | | | | | | | | | | |
| NC_002506 | 32517 | 32901 | 385 | 32696 | 306 | 34.35072 | 2.20127 | 32.13993 | peaks_peak_585 | Peak_chr2_3 | 155 | VCA0027 | VCA0026 | 33035 | 32603 | 339 | 93 | 1.7674 | 0.0000 | 2.6166 | 0.0000 | Y | Y | 81 | Up |
| NC_002506 | 40382 | 40714 | 333 | 40572 | 324 | 39.21965 | 2.2846 | 36.8925 | peaks_peak_586 | Peak_chr2_4 | 127 | NA | NA | | | | | | | | | | | | |
| NC_002506 | 43050 | 43425 | 376 | 43238 | 338 | 45.32961 | 2.40754 | 42.84315 | peaks_peak_587 | Peak_chr2_5 | 100 | VCA0035 | VCA0034 | 43274 | 43086 | 36 | 152 | 4.8570 | 0.0000 | 0.6027 | 0.0670 | Y | N | 82 | Up |
| NC_002506 | 45037 | 45400 | 364 | 45217 | 232 | 12.74975 | 1.66854 | 11.0836 | peaks_peak_589 | Peak_chr2_6 | 384 | NA | VCA0036 | | 45297 | | -80 | | | -2.6619 | 0.0000 | | Y | 83 | Down |
| NC_002506 | 48167 | 48546 | 380 | 48302 | 233 | 13.12791 | 1.68157 | 11.44959 | peaks_peak_590 | Peak_chr2_7 | 372 | VCA0041 | VCA0040 | 48768 | 48229 | 466 | 73 | 0.0742 | 0.8632 | -0.0848 | 0.8217 | N | N | | |
| NC_002506 | 60697 | 61026 | 330 | 60884 | 322 | 32.52867 | 2.10202 | 30.35982 | peaks_peak_592 | Peak_chr2_8 | 169 | NA | NA | | | | | | | | | | | | |
| NC_002506 | 75330 | 75598 | 269 | 75479 | 230 | 12.07005 | 1.64512 | 10.42854 | peaks_peak_594 | Peak_chr2_9 | 404 | NA | NA | | | | | | | | | | | | |
| NC_002506 | 79135 | 79507 | 373 | 79340 | 335 | 44.70268 | 2.40082 | 42.23217 | peaks_peak_595 | Peak_chr2_10 | 105 | VCA0070 | VCA0068 | 79473 | 79102 | 133 | 238 | 1.2090 | 0.0000 | -0.0791 | 0.7693 | Y | N | 84 | Up |
| NC_002506 | 80197 | 80568 | 372 | 80388 | 309 | 33.42539 | 2.1647 | 31.23545 | peaks_peak_596 | Peak_chr2_11 | 162 | VCA0071 | NA | 80372 | | -16 | | 0.8285 | 0.0001 | | | Y | | | Up |
| NC_002506 | 83962 | 84318 | 357 | 84121 | 235 | 14.48789 | 1.73066 | 12.76454 | peaks_peak_597 | Peak_chr2_12 | 347 | NA | VCA0074 | | 84092 | | 29 | | | 0.5727 | 0.0003 | | Y | 85 | Up |
| NC_002506 | 86222 | 86769 | 548 | 86414 | 348 | 53.08484 | 2.58272 | 50.35648 | peaks_peak_599 | Peak_chr2_13 | 67 | NA | VCA0076 | | 86223 | | 191 | | | -0.6977 | 0.4684 | | N | | |
| NC_002506 | 88942 | 89224 | 283 | 89058 | 229 | 13.79219 | 1.71627 | 12.09208 | peaks_peak_600 | Peak_chr2_14 | 355 | VCA0079 | NA | 89148 | | 90 | | 1.5805 | 0.0000 | | | Y | | 86 | Up |
| NC_002506 | 92145 | 92558 | 414 | 92389 | 334 | 47.8334 | 2.49205 | 45.27089 | peaks_peak_602 | Peak_chr2_15 | 84 | NA | VCA0080 | | 92381 | | 8 | | | -0.3381 | 0.2229 | | N | | |
| NC_002506 | 101796 | 102378 | 583 | 102202 | 257 | 20.20156 | 1.8887 | 18.3242 | peaks_peak_603 | Peak_chr2_16 | 264 | VCA0095 | NA | 102421 | | 219 | | 4.9704 | 0.0000 | | | Y | | 87 | Up |
| NC_002506 | 113127 | 113455 | 329 | 113293 | 296 | 29.51802 | 2.08603 | 27.41978 | peaks_peak_606 | Peak_chr2_17 | 187 | NA | NA | | | | | | | | | | | | |
| NC_002506 | 114472 | 115460 | 989 | 114985 | 274 | 25.58486 | 2.02477 | 23.57684 | peaks_peak_607 | Peak_chr2_18 | 215 | VCA0105 | NA | 115142 | | 157 | | 0.5873 | 0.0039 | | | N | | | |
| NC_002506 | 153218 | 154043 | 826 | 153646 | 399 | 71.20307 | 2.84268 | 67.47878 | peaks_peak_612 | Peak_chr2_19 | 12 | VCA0140 | VCA0139 | 153872 | 153464 | 226 | 182 | 4.8755 | 0.0000 | 7.4852 | 0.0000 | Y | | 88 | Up |
| NC_002506 | 185471 | 185759 | 289 | 185628 | 304 | 27.9092 | 2.01605 | 25.84709 | peaks_peak_613 | Peak_chr2_20 | 200 | NA | VCA0167 | | 185474 | | 154 | | | 1.2397 | 0.0000 | | Y | 89 | Up |
| NC_002506 | 212515 | 213323 | 809 | 212704 | 325 | 49.0576 | 2.56352 | 46.45624 | peaks_peak_615 | Peak_chr2_21 | 78 | NA | VCA0195 | | 212549 | | 155 | | | 2.2576 | 0.0000 | | Y | 90 | Up |
| NC_002506 | 219869 | 220356 | 488 | 219989 | 236 | 14.70996 | 1.73709 | 12.97906 | peaks_peak_617 | Peak_chr2_22 | 338 | NA | NA | | | | | | | | | | | | |
| NC_002506 | 236497 | 236797 | 301 | 236676 | 222 | 12.92199 | 1.69606 | 11.25017 | peaks_peak_619 | Peak_chr2_23 | 380 | NA | VCA0218 | | 236620 | | 56 | | | -0.0988 | 0.8413 | | N | | |
| NC_002506 | 237693 | 238080 | 388 | 237908 | 279 | 26.87406 | 2.05225 | 24.83637 | peaks_peak_620 | Peak_chr2_24 | 206 | NA | NA | | | | | | | | | | | | |
| NC_002506 | 241776 | 242067 | 292 | 241936 | 221 | 11.85549 | 1.65378 | 10.2222 | peaks_peak_621 | Peak_chr2_25 | 415 | VCA0221 | NA | 242108 | | 172 | | 0.5684 | 0.1481 | | | N | | | |
| NC_002506 | 244139 | 245125 | 987 | 244353 | 319 | 38.34626 | 2.27606 | 36.04035 | peaks_peak_622 | Peak_chr2_26 | 134 | NA | NA | | | | | | | | | | | | |
| NC_002506 | 248673 | 248959 | 287 | 248796 | 241 | 14.57455 | 1.72127 | 12.84786 | peaks_peak_624 | Peak_chr2_27 | 341 | VCA0227 | NA | 248766 | | -30 | | -1.8149 | 0.0000 | | | Y | | 91 | Down |
| NC_002506 | 253335 | 253642 | 308 | 253451 | 248 | 18.12274 | 1.83704 | 16.29917 | peaks_peak_625 | Peak_chr2_28 | 285 | VCA0232 | NA | 253614 | | 163 | | -0.8394 | 0.0014 | | | N | | | |
| NC_002506 | 255593 | 256030 | 438 | 255839 | 366 | 60.82727 | 2.71713 | 57.78638 | peaks_peak_626 | Peak_chr2_29 | 41 | VCA0235 | NA | 256240 | | 401 | | 1.7741 | 0.0000 | | | Y | | 92 | Up |
| NC_002506 | 275460 | 275783 | 324 | 275655 | 271 | 23.35266 | 1.95949 | 21.39851 | peaks_peak_627 | Peak_chr2_30 | 229 | NA | VCA0254 | | 275421 | | 234 | | | 3.3807 | 0.0000 | | Y | 93 | Up |
| NC_002506 | 278891 | 279307 | 417 | 279130 | 366 | 58.57586 | 2.65752 | 55.63623 | peaks_peak_628 | Peak_chr2_31 | 48 | NA | VCA0258 | | 279053 | | 77 | | | 4.8287 | 0.0000 | | | | |
| NC_002506 | 289860 | 290363 | 504 | 290109 | 396 | 54.60437 | 2.45095 | 51.81646 | peaks_peak_630 | Peak_chr2_32 | 61 | VCA0271 | NA | 290166 | | 57 | | 3.1173 | 0.0000 | | | | | | |
| NC_002506 | 292292 | 292652 | 361 | 292428 | 228 | 11.74622 | 1.63572 | 10.11734 | peaks_peak_631 | Peak_chr2_33 | 419 | NA | NA | | | | | | | | | | | | |
| NC_002506 | 310117 | 310437 | 321 | 310260 | 298 | 36.10144 | 2.28089 | 33.84974 | peaks_peak_634 | Peak_chr2_34 | 147 | NA | NA | | | | | | | | | | | | |
| NC_002506 | 322819 | 323710 | 892 | 323139 | 234 | 14.57007 | 1.73603 | 12.84357 | peaks_peak_641 | Peak_chr2_35 | 342 | VCA0313 | VCA0311 | 323508 | 323021 | 369 | 118 | -0.1594 | 0.5343 | -0.3589 | 0.1177 | N | N | | |
| NC_002506 | 334843 | 335321 | 479 | 335044 | 342 | 54.47573 | 2.64627 | 51.69189 | peaks_peak_649 | Peak_chr2_36 | 63 | NA | NA | | | | | | | | | | | | |
| NC_002506 | 344826 | 345123 | 298 | 345005 | 216 | 12.1809 | 1.67787 | 10.53529 | peaks_peak_657 | Peak_chr2_37 | 400 | NA | NA | | | | | | | | | | | | |
| NC_002506 | 351921 | 353042 | 1122 | 352252 | 236 | 20.27205 | 1.94925 | 18.39255 | peaks_peak_660 | Peak_chr2_38 | 262 | VCA0364 | NA | 352409 | | 157 | | 0.1572 | 0.6849 | | | N | | | |
| NC_002506 | 365576 | 366112 | 537 | 365926 | 236 | 11.98309 | 1.6306 | 10.34495 | peaks_peak_663 | Peak_chr2_39 | 409 | VCA0387 | VCA0386 | 366171 | 365929 | 245 | -3 | -0.0541 | 0.8746 | 0.1027 | 0.7789 | N | N | | |
| NC_002506 | 373927 | 375021 | 1095 | 374096 | 248 | 21.41506 | 1.9562 | 19.50654 | peaks_peak_666 | Peak_chr2_40 | 247 | VCA0400 | NA | 374494 | | 398 | | 0.2744 | 0.2652 | | | N | | | |
| NC_002506 | 379893 | 380359 | 467 | 380136 | 217 | 11.70457 | 1.65549 | 10.07729 | peaks_peak_668 | Peak_chr2_41 | 420 | VCA0410 | NA | 380380 | | 244 | | -0.1401 | 0.6152 | | | N | | | |
| NC_002506 | 399891 | 400666 | 776 | 400418 | 318 | 51.13597 | 2.65638 | 48.47186 | peaks_peak_674 | Peak_chr2_42 | 73 | VCA0447 | NA | 400356 | | -62 | | 0.1611 | 0.7116 | | | N | | | |
| NC_002506 | 403548 | 405172 | 1625 | 404753 | 263 | 30.23884 | 2.21725 | 28.12459 | peaks_peak_676 | Peak_chr2_43 | 185 | NA | VCA0453 | | 404784 | | -31 | | | -0.2888 | 0.8313 | | N | | |
| NC_002506 | 409978 | 410361 | 384 | 410142 | 230 | 15.91808 | 1.79801 | 14.1521 | peaks_peak_678 | Peak_chr2_44 | 318 | VCA0460 | NA | 410246 | | 104 | | -0.3281 | 0.7230 | | | N | | | |
| NC_002506 | 410832 | 411408 | 577 | 411146 | 277 | 24.38416 | 1.97733 | 22.40575 | peaks_peak_679 | Peak_chr2_45 | 226 | NA | NA | | | | | | | | | | | | |
| NC_002506 | 412175 | 412655 | 481 | 412378 | 234 | 12.22156 | 1.64371 | 10.57436 | peaks_peak_680 | Peak_chr2_46 | 399 | VCA0463 | NA | 412362 | | -16 | | 0.3504 | 0.1582 | | | N | | | |
| NC_002506 | 420832 | 421183 | 352 | 421029 | 220 | 13.04854 | 1.70566 | 11.37273 | peaks_peak_681 | Peak_chr2_47 | 374 | VCA0480 | NA | 421433 | | 404 | | 0.3349 | 0.6308 | | | N | | | |
| NC_002506 | 427864 | 428746 | 883 | 428229 | 321 | 38.06873 | 2.26162 | 35.76943 | peaks_peak_683 | Peak_chr2_48 | 137 | VCA0493 | NA | 428335 | | 106 | | 0.1886 | 0.3907 | | | N | | | |
| NC_002506 | 431223 | 431972 | 750 | 431668 | 347 | 53.8356 | 2.60742 | 51.08102 | peaks_peak_685 | Peak_chr2_49 | 65 | NA | NA | | | | | | | | | | | | |
| NC_002506 | 438968 | 439300 | 333 | 439138 | 314 | 39.58649 | 2.32906 | 37.25007 | peaks_peak_688 | Peak_chr2_50 | 125 | NA | NA | | | | | | | | | | | | |
| NC_002506 | 443572 | 444200 | 629 | 443906 | 350 | 57.17034 | 2.68682 | 54.28672 | peaks_peak_689 | Peak_chr2_51 | 51 | VCA0514 | NA | 444244 | | 338 | | 2.3737 | 0.0000 | | | Y | | 94 | Up |
| NC_002506 | 472019 | 472326 | 308 | 472165 | 273 | 26.5208 | 2.05846 | 24.49101 | peaks_peak_692 | Peak_chr2_52 | 211 | NA | NA | | | | | | | | | | | | |
| NC_002506 | 473816 | 474116 | 301 | 473990 | 232 | 15.83577 | 1.7899 | 14.07216 | peaks_peak_694 | Peak_chr2_53 | 321 | NA | VCA0536 | | 474038 | | -48 | | | -0.0706 | 0.7208 | | N | | |
| NC_002506 | 485302 | 485776 | 475 | 485497 | 312 | 42.58051 | 2.42505 | 40.16882 | peaks_peak_695 | Peak_chr2_54 | 114 | VCA0547 | NA | 485540 | | 43 | | 0.8902 | 0.0000 | | | Y | | 95 | Up |
| NC_002506 | 491965 | 493769 | 1805 | 492436 | 392 | 78.44988 | 3.06093 | 73.86951 | peaks_peak_696 | Peak_chr2_55 | 2 | VCA0556 | VCA0554 | 492527 | 492116 | 91 | 320 | 2.2209 | 0.0012 | -0.3586 | 0.7337 | N | N | | |
| NC_002506 | 503142 | 503409 | 268 | 503262 | 216 | 12.74059 | 1.70162 | 11.07488 | peaks_peak_697 | Peak_chr2_56 | 385 | VCA0565 | NA | 503254 | | -8 | | 2.4377 | 0.0000 | | | Y | | 96 | Up |
| NC_002506 | 509601 | 510294 | 694 | 509747 | 277 | 27.92431 | 2.09225 | 25.86183 | peaks_peak_700 | Peak_chr2_57 | 199 | NA | NA | | | | | | | | | | | | |
| NC_002506 | 510930 | 511268 | 339 | 511082 | 237 | 17.03689 | 1.82382 | 15.24093 | peaks_peak_701 | Peak_chr2_58 | 295 | VCA0574 | VCA0573 | 511108 | 510901 | 26 | 181 | 1.3750 | 0.0001 | 0.5191 | 0.1361 | Y | N | 97 | Up |
| NC_002506 | 512263 | 512810 | 548 | 512392 | 214 | 12.08645 | 1.67812 | 10.44449 | peaks_peak_702 | Peak_chr2_59 | 403 | NA | NA | | | | | | | | | | | | |
| NC_002506 | 517158 | 517769 | 612 | 517396 | 348 | 50.96916 | 2.52501 | 48.30945 | peaks_peak_703 | Peak_chr2_60 | 74 | NA | NA | | | | | | | | | | | | |
| NC_002506 | 520842 | 521969 | 1128 | 521410 | 345 | 55.9913 | 2.6758 | 53.14935 | peaks_peak_704 | Peak_chr2_61 | 56 | VCA0583 | VCA0582 | 521497 | 521381 | 87 | 29 | 0.1050 | 0.8805 | -0.6397 | 0.0001 | N | Y | | Down |
| NC_002506 | 528860 | 530078 | 1219 | 529512 | 391 | 74.32182 | 2.95816 | 70.20187 | peaks_peak_705 | Peak_chr2_62 | 6 | NA | NA | | | | | | | | | | | | |
| NC_002506 | 532096 | 532665 | 570 | 532338 | 321 | 45.41952 | 2.4736 | 42.93078 | peaks_peak_706 | Peak_chr2_63 | 99 | VCA0594 | VCA0593 | 532353 | 532143 | 15 | 195 | 3.6513 | 0.0000 | -0.4694 | 0.0091 | Y | N | | Up |
| NC_002506 | 539914 | 540318 | 405 | 540090 | 329 | 47.24244 | 2.49476 | 44.69712 | peaks_peak_708 | Peak_chr2_64 | 89 | NA | VCA0602 | | 540083 | | 7 | | | 0.1005 | 0.9173 | | N | | |
| NC_002506 | 564922 | 565224 | 303 | 565081 | 237 | 20.60973 | 1.95901 | 18.72177 | peaks_peak_712 | Peak_chr2_65 | 259 | NA | NA | | | | | | | | | | | | |
| NC_002506 | 591822 | 592134 | 313 | 591970 | 237 | 11.70062 | 1.61764 | 10.0735 | peaks_peak_715 | Peak_chr2_66 | 421 | NA | NA | | | | | | | | | | | | |
| NC_002506 | 592351 | 592789 | 439 | 592641 | 269 | 27.16223 | 2.09209 | 25.11757 | peaks_peak_716 | Peak_chr2_67 | 203 | glpD | VCA0656 | 592946 | 592599 | 305 | 42 | -1.4514 | 0.0000 | -1.1640 | 0.0003 | Y | Y | 98 | Down |
| NC_002506 | 598182 | 598989 | 808 | 598364 | 238 | 18.63022 | 1.8815 | 16.79317 | peaks_peak_717 | Peak_chr2_68 | 280 | VCA0663 | VCA0662 | 598557 | 598410 | 193 | -46 | -0.8472 | 0.0000 | -0.1770 | 0.4300 | Y | N | 99 | Down |
| NC_002506 | 600066 | 601115 | 1050 | 600671 | 395 | 85.00883 | 3.21892 | 78.50765 | peaks_peak_719 | Peak_chr2_69 | 1 | dcuC | NA | 600714 | | 43 | | -2.1248 | 0.0000 | | | Y | | 100 | Down |
| NC_002506 | 601927 | 602550 | 624 | 602146 | 308 | 39.80518 | 2.35717 | 37.46393 | peaks_peak_720 | Peak_chr2_70 | 124 | NA | NA | | | | | | | | | | | | |
| NC_002506 | 606691 | 607009 | 319 | 606836 | 293 | 35.49644 | 2.27994 | 33.25877 | peaks_peak_722 | Peak_chr2_71 | 149 | VCA0671 | NA | 606844 | | 8 | | -0.0352 | 0.9075 | | | N | | | |
| NC_002506 | 611001 | 611334 | 334 | 611160 | 335 | 48.08622 | 2.4953 | 45.51633 | peaks_peak_724 | Peak_chr2_72 | 83 | NA | NA | | | | | | | | | | | | |
| NC_002506 | 621877 | 622829 | 953 | 622375 | 379 | 64.61762 | 2.76116 | 61.35179 | peaks_peak_727 | Peak_chr2_73 | 27 | VCA0686 | NA | 622639 | | 264 | | -0.3429 | 0.4694 | | | N | | | |
| NC_002506 | 634670 | 635098 | 429 | 634853 | 302 | 40.09607 | 2.38868 | 37.74814 | peaks_peak_728 | Peak_chr2_74 | 123 | VCA0697 | VCA0695 | 635108 | 634750 | 255 | 103 | -2.2352 | 0.0000 | 0.3851 | 0.1803 | Y | N | | Down |
| NC_002506 | 649341 | 650010 | 670 | 649596 | 302 | 34.90301 | 2.23074 | 32.67854 | peaks_peak_730 | Peak_chr2_75 | 153 | VCA0707 | VCA0706 | 649740 | 649493 | 144 | 103 | -0.1484 | 0.8130 | 1.3204 | 0.0000 | N | Y | | Up |
| NC_002506 | 658660 | 658936 | 277 | 658787 | 220 | 12.60279 | 1.6871 | 10.94198 | peaks_peak_731 | Peak_chr2_76 | 387 | VCA0715 | NA | 658854 | | 67 | | 1.0336 | 0.0000 | | | Y | | 101 | Up |
| NC_002506 | 665620 | 666163 | 544 | 665856 | 368 | 63.08979 | 2.76871 | 59.9067 | peaks_peak_734 | Peak_chr2_77 | 35 | VCA0721 | VCA0720 | 665990 | 665712 | 134 | 144 | 3.2318 | 0.0000 | -0.6798 | 0.0222 | Y | N | | Up |
| NC_002506 | 673489 | 673883 | 395 | 673678 | 318 | 46.07541 | 2.50497 | 43.56647 | peaks_peak_735 | Peak_chr2_78 | 95 | NA | NA | | | | | | | | | | | | |
| NC_002506 | 683255 | 683514 | 260 | 683380 | 244 | 15.30901 | 1.74261 | 13.56002 | peaks_peak_738 | Peak_chr2_79 | 328 | NA | VCA0736 | | 683045 | | 335 | | | -0.6637 | 0.0001 | | Y | 102 | Down |
| NC_002506 | 703420 | 703676 | 257 | 703558 | 218 | 12.57471 | 1.69022 | 10.91489 | peaks_peak_740 | Peak_chr2_80 | 389 | NA | NA | | | | | | | | | | | | |
| NC_002506 | 717822 | 718211 | 390 | 718055 | 272 | 25.22093 | 2.01852 | 23.22158 | peaks_peak_741 | Peak_chr2_81 | 217 | VCA0776 | VCA0774 | 718129 | 717809 | 74 | 246 | -0.0020 | 0.9882 | 0.0452 | 0.8840 | N | N | | |
| NC_002506 | 733266 | 733624 | 359 | 733437 | 339 | 47.05308 | 2.45146 | 44.51347 | peaks_peak_742 | Peak_chr2_82 | 91 | VCA0789 | VCA0788 | 733638 | 733402 | 201 | 35 | 1.1876 | 0.0006 | -0.2818 | 0.2555 | Y | N | | Up |
| NC_002506 | 735567 | 736889 | 1323 | 736198 | 243 | 13.2017 | 1.66515 | 11.52116 | peaks_peak_743 | Peak_chr2_83 | 370 | NA | NA | | | | | | | | | | | | |
| NC_002506 | 757837 | 758354 | 518 | 758078 | 330 | 44.88443 | 2.42394 | 42.40934 | peaks_peak_745 | Peak_chr2_84 | 104 | NA | NA | | | | | | | | | | | | |
| NC_002506 | 776812 | 777195 | 384 | 777026 | 277 | 27.0724 | 2.06456 | 25.0301 | peaks_peak_746 | Peak_chr2_85 | 204 | NA | NA | | | | | | | | | | | | |
| NC_002506 | 787029 | 788437 | 1409 | 787470 | 372 | 55.55176 | 2.55585 | 52.72683 | peaks_peak_750 | Peak_chr2_86 | 58 | VCA0845 | NA | 787558 | | 88 | | 2.8051 | 0.0000 | | | Y | | 103 | Up |
| NC_002506 | 808798 | 809083 | 286 | 808944 | 289 | 30.3577 | 2.13279 | 28.24091 | peaks_peak_754 | Peak_chr2_87 | 182 | malS | VCA0859 | 809269 | 808977 | 325 | -33 | 0.0044 | 0.9988 | -0.6365 | 0.1006 | N | N | | |
| NC_002506 | 811361 | 811681 | 321 | 811541 | 249 | 18.19344 | 1.83717 | 16.36813 | peaks_peak_755 | Peak_chr2_88 | 283 | VCA0862 | NA | 811876 | | 335 | | -1.3452 | 0.0323 | | | N | | | |
| NC_002506 | 819916 | 820312 | 397 | 820067 | 220 | 12.30839 | 1.67479 | 10.65793 | peaks_peak_756 | Peak_chr2_89 | 396 | VCA0868 | NA | 820464 | | 397 | | -1.4554 | 0.0008 | | | Y | | 104 | Down |
| NC_002506 | 823846 | 824237 | 392 | 824086 | 256 | 16.50905 | 1.761 | 14.72679 | peaks_peak_757 | Peak_chr2_90 | 305 | NA | NA | | | | | | | | | | | | |
| NC_002506 | 826794 | 827131 | 338 | 826958 | 321 | 45.59758 | 2.47881 | 43.10437 | peaks_peak_758 | Peak_chr2_91 | 97 | NA | NA | | | | | | | | | | | | |
| NC_002506 | 829092 | 829602 | 511 | 829432 | 211 | 12.03206 | 1.68232 | 10.39208 | peaks_peak_759 | Peak_chr2_92 | 408 | VCA0876 | VCA0875 | 829467 | 829259 | 35 | 173 | -0.5000 | 0.0783 | 0.0489 | 0.8986 | N | N | | |
| NC_002506 | 830091 | 830368 | 278 | 830248 | 237 | 17.70211 | 1.84907 | 15.88927 | peaks_peak_760 | Peak_chr2_93 | 290 | VCA0877 | NA | 830711 | | 463 | | 0.2914 | 0.3940 | | | N | | | |
| NC_002506 | 870731 | 871263 | 533 | 871071 | 284 | 28.90216 | 2.10226 | 26.81796 | peaks_peak_765 | Peak_chr2_94 | 191 | VCA0919 | VCA0917 | 871322 | 870823 | 251 | 248 | 0.3142 | 0.0596 | 3.4163 | 0.0000 | N | Y | 105 | Up |
| NC_002506 | 881845 | 882203 | 359 | 882059 | 270 | 24.86878 | 2.01257 | 22.87787 | peaks_peak_767 | Peak_chr2_95 | 220 | NA | NA | | | | | | | | | | | | |
| NC_002506 | 894453 | 894807 | 355 | 894671 | 244 | 14.91848 | 1.72801 | 13.18128 | peaks_peak_768 | Peak_chr2_96 | 332 | NA | malG | | 894347 | | 324 | | | 0.4568 | 0.0498 | | N | | |
| NC_002506 | 899626 | 899887 | 262 | 899732 | 244 | 17.77796 | 1.83421 | 15.96319 | peaks_peak_769 | Peak_chr2_97 | 288 | VCA0948 | NA | 899732 | | 0 | | 2.3559 | 0.0000 | | | Y | | 106 | Up |
| NC_002506 | 901578 | 902158 | 581 | 901906 | 271 | 26.55833 | 2.06575 | 24.52762 | peaks_peak_770 | Peak_chr2_98 | 209 | NA | NA | | | | | | | | | | | | |
| NC_002506 | 902582 | 903156 | 575 | 902722 | 243 | 18.37082 | 1.85863 | 16.54076 | peaks_peak_771 | Peak_chr2_99 | 282 | VCA0954 | NA | 903131 | | 409 | | -1.1843 | 0.0000 | | | Y | | 107 | Down |
| NC_002506 | 906318 | 906741 | 424 | 906437 | 232 | 13.73932 | 1.70777 | 12.0412 | peaks_peak_772 | Peak_chr2_100 | 359 | VCA0958 | NA | 906685 | | 248 | | -0.2950 | 0.6198 | | | N | | | |
| NC_002506 | 908228 | 908535 | 308 | 908391 | 249 | 18.18457 | 1.83685 | 16.3594 | peaks_peak_773 | Peak_chr2_101 | 284 | NA | NA | | | | | | | | | | | | |
| NC_002506 | 910786 | 911257 | 472 | 911031 | 391 | 60.1454 | 2.60134 | 57.13753 | peaks_peak_775 | Peak_chr2_102 | 45 | VCA0962 | NA | 911092 | | 61 | | 5.1435 | 0.0000 | | | | | | |
| NC_002506 | 913971 | 914268 | 298 | 914133 | 276 | 25.06067 | 2.00202 | 23.0656 | peaks_peak_776 | Peak_chr2_103 | 218 | VCA0965 | NA | 914406 | | 273 | | -0.6339 | 0.0461 | | | N | | | |
| NC_002506 | 918610 | 918989 | 380 | 918776 | 330 | 43.63111 | 2.38862 | 41.19163 | peaks_peak_777 | Peak_chr2_104 | 110 | NA | NA | | | | | | | | | | | | |
| NC_002506 | 919332 | 919937 | 606 | 919500 | 243 | 16.41068 | 1.7859 | 14.63103 | peaks_peak_778 | Peak_chr2_105 | 308 | VCA0973 | NA | 919692 | | 192 | | -1.4919 | 0.5708 | | | N | | | |
| NC_002506 | 942238 | 942540 | 303 | 942395 | 232 | 12.17506 | 1.64558 | 10.52957 | peaks_peak_780 | Peak_chr2_106 | 401 | VCA0989 | NA | 942593 | | 198 | | -2.1277 | 0.0000 | | | Y | | 108 | Down |
| NC_002506 | 949426 | 949704 | 279 | 949541 | 228 | 12.42931 | 1.66353 | 10.7747 | peaks_peak_781 | Peak_chr2_107 | 393 | VCA0996 | VCA0994 | 949658 | 949301 | 117 | 240 | -0.2052 | 0.4937 | 0.4586 | 0.0519 | N | N | | |
| NC_002506 | 954872 | 955200 | 329 | 955066 | 259 | 20.23023 | 1.88466 | 18.3518 | peaks_peak_782 | Peak_chr2_108 | 263 | VCA1001 | VCA1000 | 955133 | 954932 | 67 | 134 | 0.3725 | 0.1219 | -0.8328 | 0.3364 | N | N | | |
| NC_002506 | 957417 | 957793 | 377 | 957557 | 287 | 21.2352 | 1.85276 | 19.33132 | peaks_peak_783 | Peak_chr2_109 | 249 | NA | NA | | | | | | | | | | | | |
| NC_002506 | 958930 | 959228 | 299 | 959079 | 256 | 18.84393 | 1.84354 | 17.00179 | peaks_peak_784 | Peak_chr2_110 | 278 | VCA1008 | NA | 959181 | | 102 | | 1.0299 | 0.0000 | | | Y | | 109 | Up |
| NC_002506 | 962782 | 963037 | 256 | 962896 | 237 | 13.91362 | 1.7042 | 12.20993 | peaks_peak_785 | Peak_chr2_111 | 353 | NA | VCA1012 | | 962579 | | 317 | | | 0.3476 | 0.2205 | | N | | |
| NC_002506 | 963455 | 963760 | 306 | 963601 | 247 | 16.3119 | 1.77324 | 14.53477 | peaks_peak_786 | Peak_chr2_112 | 309 | VCA1015 | VCA1013 | 963699 | 963307 | 98 | 294 | 1.0846 | 0.0016 | 0.2280 | 0.5705 | N | N | | |
| NC_002506 | 965164 | 965427 | 264 | 965278 | 253 | 14.43087 | 1.69223 | 12.70965 | peaks_peak_787 | Peak_chr2_113 | 348 | NA | NA | | | | | | | | | | | | |
| NC_002506 | 976203 | 976854 | 652 | 976629 | 390 | 66.29358 | 2.75776 | 62.91476 | peaks_peak_789 | Peak_chr2_114 | 24 | NA | NA | | | | | | | | | | | | |
| NC_002506 | 1013415 | 1014018 | 604 | 1013567 | 248 | 15.90752 | 1.75622 | 14.14206 | peaks_peak_793 | Peak_chr2_115 | 320 | NA | potE | | 1013118 | | 449 | | | -0.6870 | 0.4080 | | N | | |
| NC_002506 | 1015396 | 1016124 | 729 | 1015893 | 304 | 33.36004 | 2.17802 | 31.17187 | peaks_peak_794 | Peak_chr2_116 | 163 | NA | NA | | | | | | | | | | | | |
| NC_002506 | 1026643 | 1026985 | 343 | 1026828 | 295 | 29.27958 | 2.08161 | 27.18674 | peaks_peak_796 | Peak_chr2_117 | 189 | NA | VCA1073 | | 1026682 | | 146 | | | -0.3555 | 0.4023 | | N | | |
| NC_002506 | 1029255 | 1029538 | 284 | 1029347 | 233 | 11.93217 | 1.634 | 10.29596 | peaks_peak_798 | Peak_chr2_118 | 411 | VCA1077 | NA | 1029543 | | 196 | | 0.0108 | 0.9522 | | | N | | | |
| NC_002506 | 1037451 | 1037860 | 410 | 1037652 | 343 | 46.28771 | 2.41614 | 43.77232 | peaks_peak_800 | Peak_chr2_119 | 94 | NA | NA | | | | | | | | | | | | |
| NC_002506 | 1039473 | 1039846 | 374 | 1039662 | 348 | 49.39413 | 2.48233 | 46.78255 | peaks_peak_801 | Peak_chr2_120 | 77 | NA | NA | | | | | | | | | | | | |
| NC_002506 | 1069967 | 1070202 | 236 | 1070059 | 246 | 16.17197 | 1.7703 | 14.39874 | peaks_peak_803 | Peak_chr2_121 | 311 | NA | NA | | | | | | | | | | | | |

**Supplementary Table S6. Summary of genes list and validation results**

O = Confirmed
X = Not changed/ Not bound
U = Unclear

| **UP** | 102 genes | RNA-Seq | ChIP-Seq | S1 Mapping | EMSA | Footprinting | Protected region by VxrB (N=nucleotide number/**Bold**=Start codon) |
|---|---|---|---|---|---|---|---|
| | VC0213 | O | O | O | | | |
| | VC0315 | O | O | | | | |
| | VC0318 | O | O | | | | |
| | VC0354 | O | O | | | | |
| | VC0370 | O | O | O | O | O | CGAAAGTTTGTGCGCCGCCTAACCAGAATTGACCTAGGGTTGACCCAAACTCAG--2N--**ATG** |
| | VC0371 | O | O | | | | |
| | VC0419 | O | O | | | | |
| | VC0480 | O | O | | | | |
| | VC0500 | O | O | | | | |
| | VC0567 | O | O | | | | |
| | VC0581 | O | O | | | | |
| | VC0666 | O | O | | | | |
| | VC0676 | O | O | | | | |
| | VC0680 | O | O | O | O | O | ATTCTATCAGCATTTTTTTACCAATGCTAATACTTATCTGATCTCACTATGACATT--37N--**TTG** |
| | VC0703 | O | O | | | | |
| | VC0860 | O | O | | | | |
| | VC0918 | O | O | | | | |
| | VC0947 | O | O | O | | | |
| | VC0948 | O | O | O | | X | |
| | VC0969 | O | O | O | O | O | AACTTAATGTGAGAAGTTGAAAAGATGAC–83N--**ATG** |
| | VC0970 | O | O | | | | |
| | VC1063 | O | O | | | | |
| | VC1064 | O | O | | | | |
| | VC1107 | O | O | | | | |
| | VC1138 | O | O | | | | |
| | VC1154 | O | O | | | | |
| | VC1158 | O | O | | | | |
| | VC1162 | O | O | | | | |
| | VC1191 | O | O | | | | |
| | VC1269 | O | O | O | | | |
| | VC1343 | O | O | | | | |
| | VC1344 | O | O | | | | |
| | VC1349 | O | O | | | | |
| | VC1418 | O | O | | | | |
| | VC1422a | O | O | | | | |
| | VC1444 | O | O | | | | |
| | VC1445 | O | O | | | | |
| | VC1484 | O | O | | | | |
| | VC1516 | O | O | | | | |
| | VC1642 | O | O | | | | |
| | VC1653 | O | O | | | | |
| | VC1655 | O | O | O | O | O | AAGCAATATCATGCAAATGCATGGTTCTGACCAAAACTTTACGTAACTATTACATTTTT--410N--**GTG** |
| | VC1663 | O | O | | | | |
| | VC1712 | O | O | | | | |
| | VC1722 | O | O | | | | |
| | VC1835 | O | O | | | | |
| | VC1839 | O | O | | | | |
| | VC1884 | O | O | | | | |
| | VC1900 | O | O | | | | |
| | VC1903 | O | O | | | | |
| | VC1956 | O | O | X | | | |
| | VC1959 | O | O | | | | |
| | VC1962 | O | O | O | O | | |
| | VC2033 | O | O | | | | |
| | VC2097 | O | O | | | | |
| | VC2157 | O | O | | | | |
| | VC2162 | O | O | | | | |
| | VC2164 | O | O | | | | |
| | VC2168 | O | O | | | | |
| | VC2212 | O | O | O | | | |
| | VC2213 | O | O | O | O | O | TGTTTTAGTCTTTTCTCATTTTTAGCAAAGACGTGACAAAATAATAACCGAAAGGCTCT--134N--**ATG** |
| | VC2253 | O | O | | | | |
| | VC2275 | O | O | | | | |
| | VC2283 | O | O | | | | |
| | VC2312 | O | O | O | | | |
| | VC2402 | O | O | | | | |
| | VC2409 | O | O | O | | | |
| | VC2431 | O | O | | | | |
| | VC2466 | O | O | | | | |
| | VC2473 | O | O | | | | |
| | VC2517 | O | O | O | | | |
| | VC2520 | O | O | | | | |
| | VC2567 | O | O | | | | |
| | VC2635 | O | O | O | O | O | AGGCAAACCCTTTTTTAGTTCTTTTAGTTTCTTTACAATTTTCTGATAATCGAG--463N--**GTG** |
| | VC2662 | O | O | | | | |
| | VC2697 | O | O | | | | |
| | VCA0026 | O | O | | | | |
| | VCA0027 | O | O | | | | |
| | VCA0035 | O | O | | | | |
| | VCA0070 | O | O | | | | |
| | VCA0071 | O | O | | | | |
| | VCA0074 | O | O | | | | |
| | VCA0079 | O | O | | | | |
| | VCA0095 | O | O | | | | |
| | VCA0140 | O | O | X | | | |
| | VCA0167 | O | O | | | | |
| | VCA0195 | O | O | | | | |
| | VCA0235 | O | O | X | | | |
| | VCA0254 | O | O | | | | |
| | VCA0514 | O | O | | | | |
| | VCA0547 | O | O | | | | |

| | | | | | | | |
|---|---|---|---|---|---|---|---|
| | VCA0565 | O | O | O | O | X | |
| | VCA0574 | O | O | | | | |
| | VCA0594 | O | O | O | O | O | **ATG**--32N--TAAAACCGCCCTATTACTCACTGTTGTTGGTGCCG |
| | VCA0706 | O | O | | | | |
| | VCA0715 | O | O | | | | |
| | VCA0721 | O | O | | | | |
| | VCA0789 | O | O | | | | |
| | VCA0845 | O | O | | | | |
| | VCA0917 | O | O | | | | |
| | VCA0948 | O | O | | | | |
| | VCA1008 | O | O | | | | |

| **DOWN** | 38 genes | RNA-Seq | ChIP-Seq | S1 Mapping | EMSA | Footprinting | Protected region by VxrB |
|---|---|---|---|---|---|---|---|
| | VC0049 | O | O | O | | X | |
| | VC0225 | O | O | | | | |
| | VC0248 | O | O | | | | |
| | VC0277 | O | O | O | | | |
| | VC0503 | O | O | | | | |
| | VC0513 | O | O | | | | |
| | VC0706 | O | O | | | | |
| | VC0746 | O | O | | | | |
| | VC0977 | O | O | | | | |
| | VC0984 | O | O | | | | |
| | VC0985 | O | O | | | | |
| | VC1059 | O | O | | | | |
| | VC1165 | O | O | | | | |
| | VC1271 | O | O | | | | |
| | VC1272 | O | O | U | O | O | GATCATTTG TCGCTGAGAACTTGACCTCATCCCTACAAAC—49N--**TTG** |
| | VC1377 | O | O | | | | |
| | VC1537 | O | O | | | | |
| | VC1548 | O | O | | | | |
| | VC1727 | O | O | | | | |
| | VC1880 | O | O | | | | |
| | VC1897 | O | O | | | | |
| | VC2037 | O | O | O | O | O | ACTCTCACTTTTTAACCAAGTTAGAGTATTGACGTCAGATTTTTACCAAAAAACTTATTTC--211N--**ATG** |
| | VC2193 | O | O | | | | |
| | VC2278 | O | O | | | | |
| | VC2686 | O | O | O | O | O | CTCGTTAAGTTAAGCTTGGGTCAAATTGCGCGGTTTTTTTAACCGAATAGC--106N--**ATG** |
| | VC2757 | O | O | | | | |
| | VCA0036 | O | O | | | | |
| | VCA0227 | O | O | X | | | |
| | VCA0582 | O | O | | | | |
| | VCA0656 | O | O | | | | |
| | VCA0657 | O | O | X | | | |
| | VCA0663 | O | O | | | | |
| | VCA0665 | O | O | X | | | |
| | VCA0699 | O | O | | | | |
| | VCA0736 | O | O | | | | |
| | VCA0868 | O | O | | | | |
| | VCA0954 | O | O | | | | |
| | VCA0989 | O | O | | | | |

### Figure S1

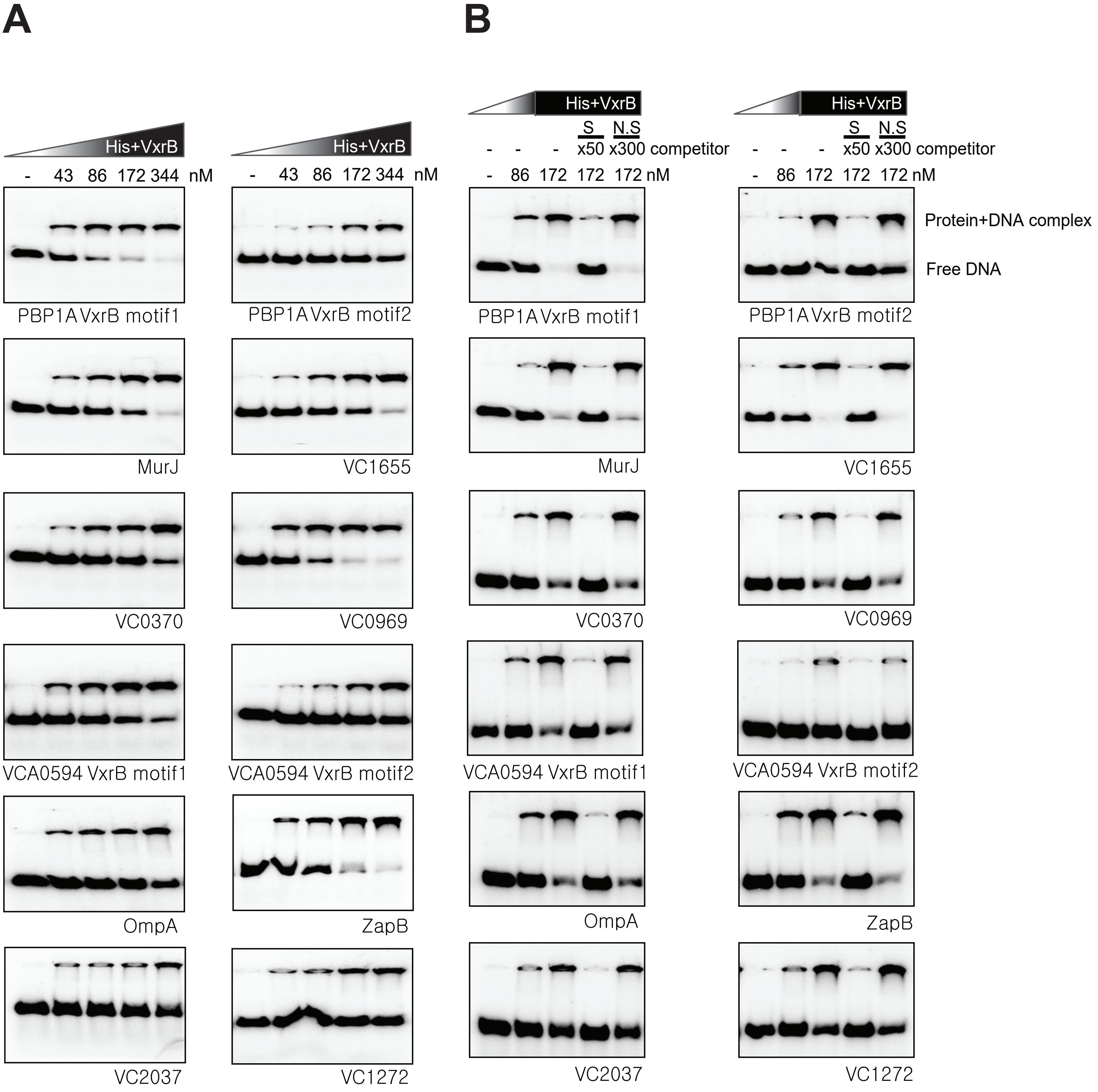

A
B
His+VxrB
- 43 86 172 344 nM
S N.S
- - - x50 x300 competitor
- 86 172 172 172 nM
Protein+DNA complex
Free DNA
PBP1A VxrB motif1
PBP1A VxrB motif2
MurJ
VC1655
VC0370
VC0969
VCA0594 VxrB motif1
VCA0594 VxrB motif2
OmpA
ZapB
VC2037
VC1272

Figure S2

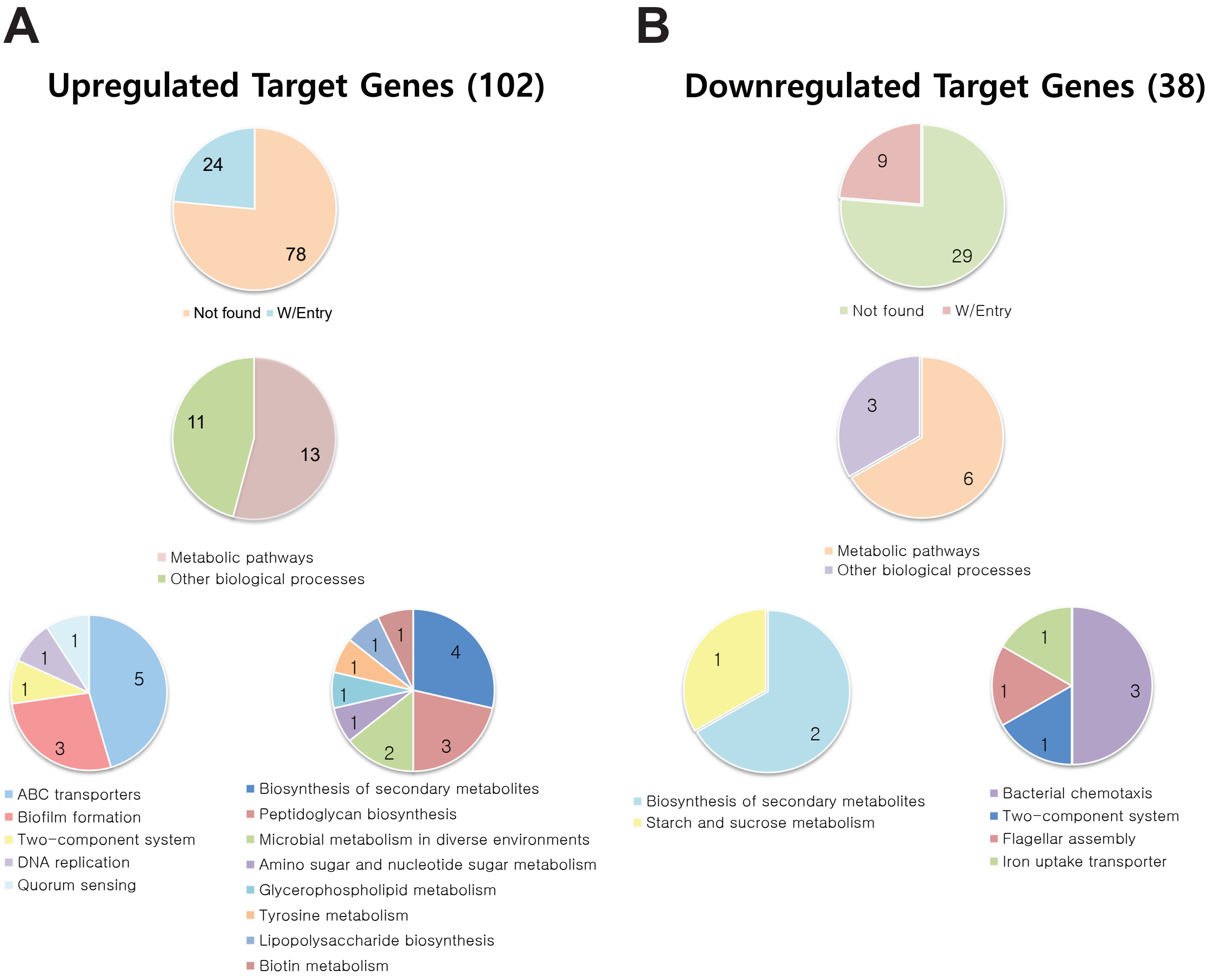
A
Upregulated Target Genes (102)
24
78
Not found
W/Entry
11
13
Metabolic pathways
Other biological processes
1
1
1
5
3
ABC transporters
Biofilm formation
Two−component system
DNA replication
Quorum sensing
1
1
1
1
1
4
3
2
Biosynthesis of secondary metabolites
Peptidoglycan biosynthesis
Microbial metabolism in diverse environments
Amino sugar and nucleotide sugar metabolism
Glycerophospholipid metabolism
Tyrosine metabolism
Lipopolysaccharide biosynthesis
Biotin metabolism
B
Downregulated Target Genes (38)
9
29
Not found
W/Entry
3
6
Metabolic pathways
Other biological processes
1
2
Biosynthesis of secondary metabolites
Starch and sucrose metabolism
1
1
3
1
Bacterial chemotaxis
Two−component system
Flagellar assembly
Iron uptake transporter

### Figure S3

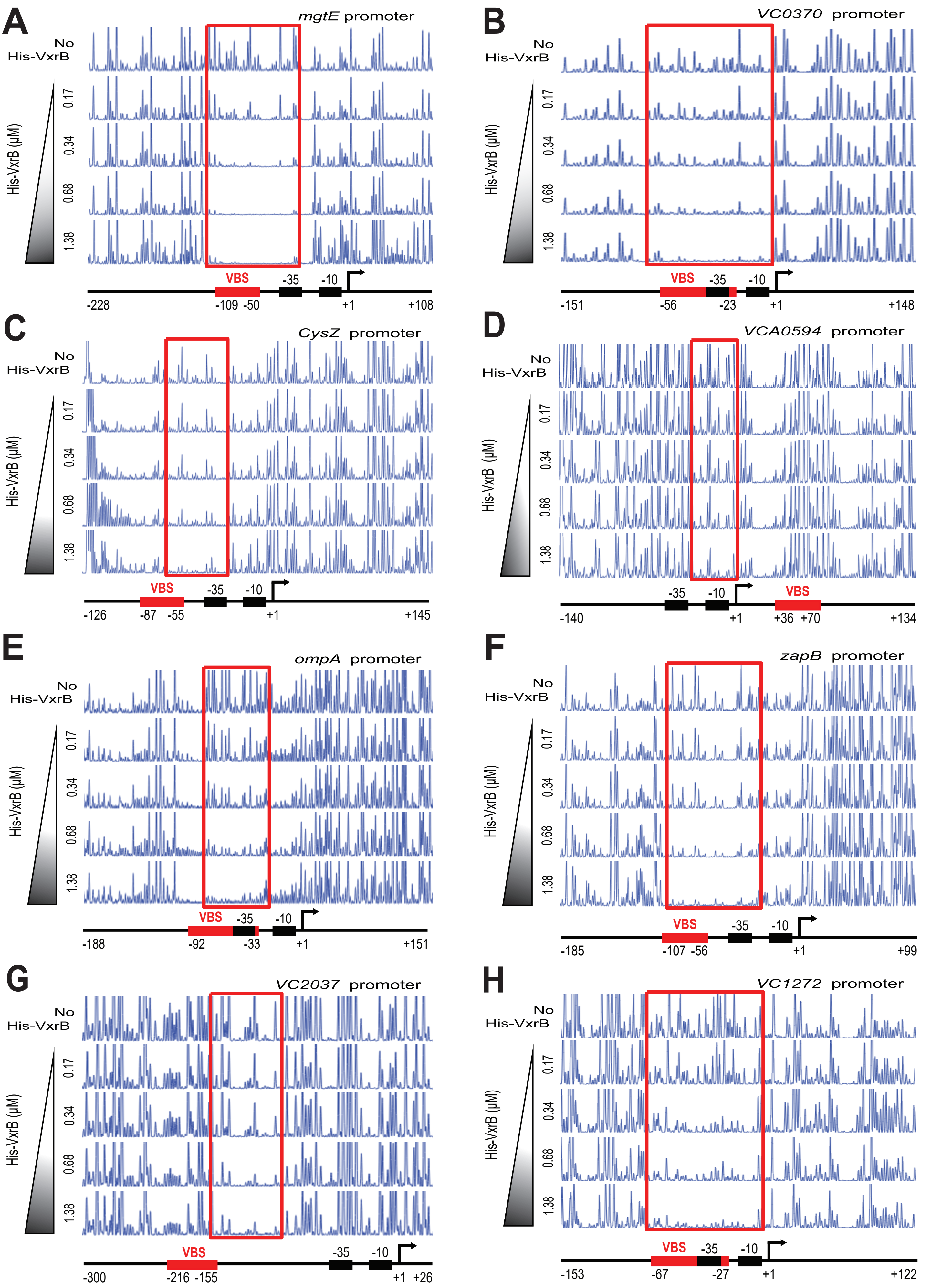
A
mgtE promoter
No His-VxrB
His-VxrB (μM)
0.17
0.34
0.68
1.38
VBS
-35
-10
-228
-109 -50
+1
+108
B
VC0370 promoter
No His-VxrB
His-VxrB (μM)
0.17
0.34
0.68
1.38
VBS
-35
-10
-151
-56
-23
+1
+148
C
CysZ promoter
No His-VxrB
His-VxrB (μM)
0.17
0.34
0.68
1.38
VBS
-35
-10
-126
-87 -55
+1
+145
D
VCA0594 promoter
No His-VxrB
His-VxrB (μM)
0.17
0.34
0.68
1.38
-35
-10
VBS
-140
+1
+36 +70
+134
E
ompA promoter
No His-VxrB
His-VxrB (μM)
0.17
0.34
0.68
1.38
VBS
-35
-10
-188
-92
-33
+1
+151
F
zapB promoter
No His-VxrB
His-VxrB (μM)
0.17
0.34
0.68
1.38
VBS
-35
-10
-185
-107 -56
+1
+99
G
VC2037 promoter
No His-VxrB
His-VxrB (μM)
0.17
0.34
0.68
1.38
VBS
-35
-10
-300
-216 -155
+1 +26
H
VC1272 promoter
No His-VxrB
His-VxrB (μM)
0.17
0.34
0.68
1.38
VBS
-35
-10
-153
-67
-27
+1
+122

Figure S4

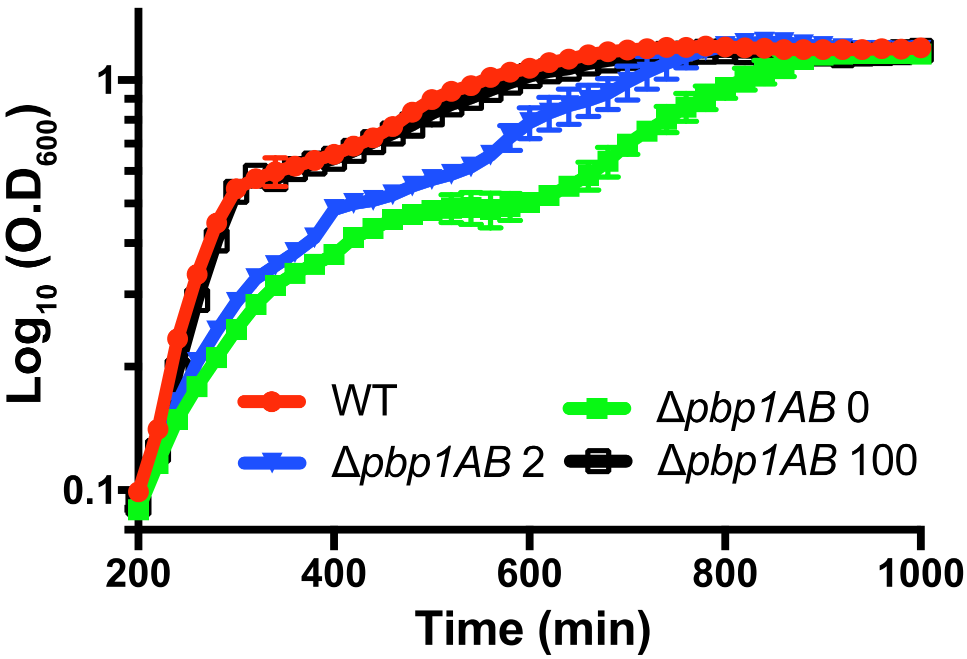

Log$_{10}$ (O.D$_{600}$)
1
0.1
200
400
600
800
1000
Time (min)
WT
Δ*pbp1AB* 2
Δ*pbp1AB* 0
Δ*pbp1AB* 100

### Figure S5

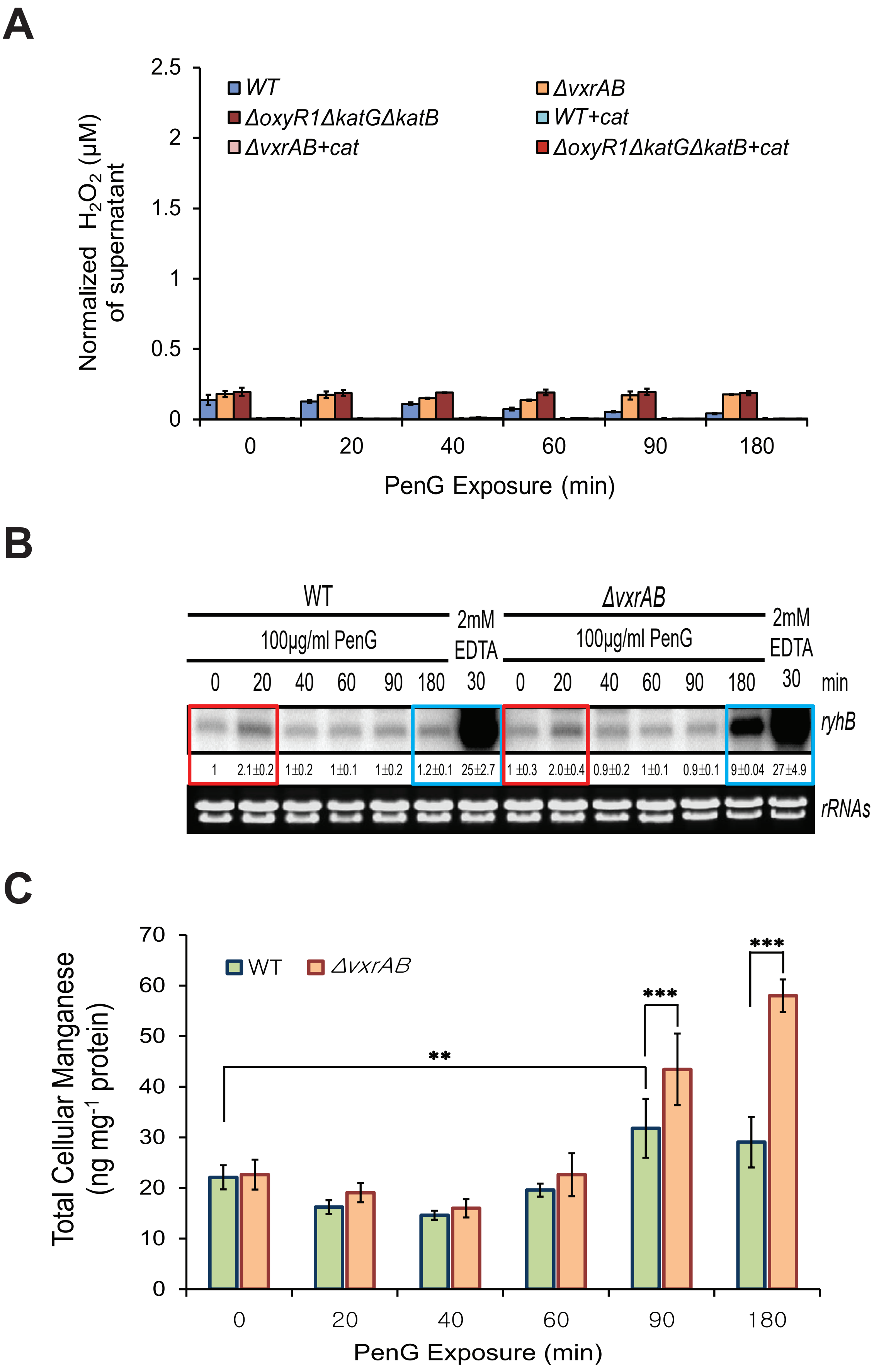

A
WT
ΔvxrAB
ΔoxyR1ΔkatGΔkatB
WT+cat
ΔvxrAB+cat
ΔoxyR1ΔkatGΔkatB+cat
Normalized $H_2O_2$ (μM) of supernatant
2.5
2
1.5
1
0.5
0
0
20
40
60
90
180
PenG Exposure (min)
B
WT
100μg/ml PenG
2mM EDTA
ΔvxrAB
100μg/ml PenG
2mM EDTA
0 20 40 60 90 180 30 0 20 40 60 90 180 30 min
ryhB
1
2.1±0.2
1±0.2
1±0.1
1±0.2
1.2±0.1
25±2.7
1 ±0.3
2.0±0.4
0.9±0.2
1±0.1
0.9±0.1
9±0.04
27±4.9
rRNAs
C
WT
ΔvxrAB
Total Cellular Manganese (ng mg$^{-1}$ protein)
70
60
50
40
30
20
10
0
**
***
***
0
20
40
60
90
180
PenG Exposure (min)

### Figure S6

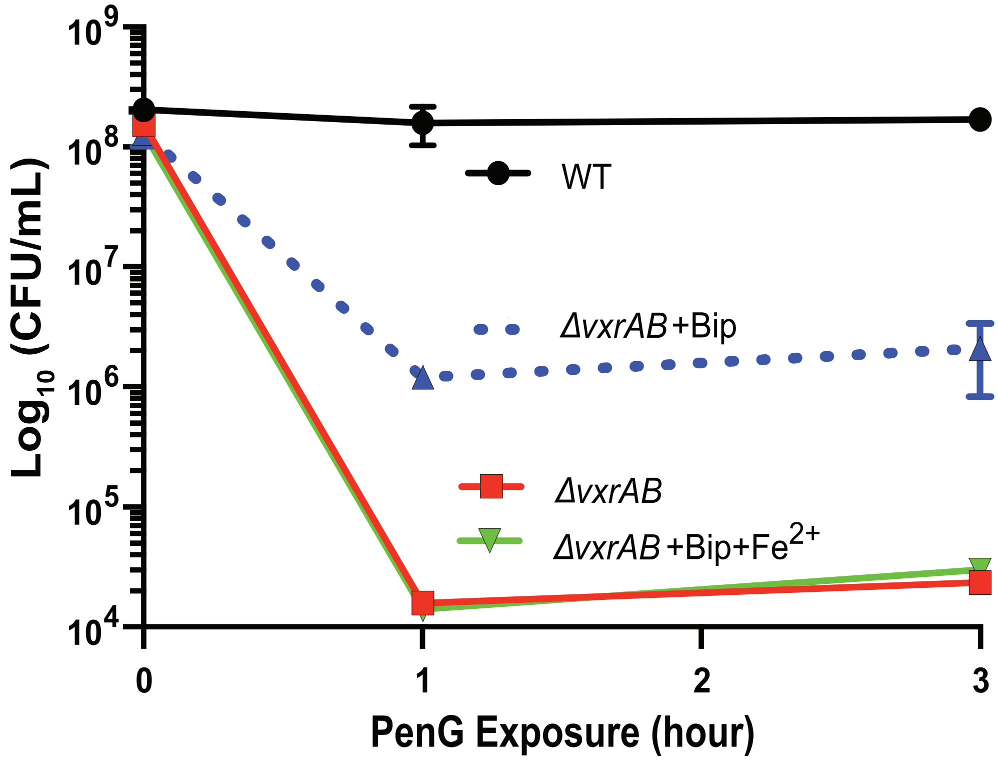

Log$_{10}$ (CFU/mL)
$10^9$
$10^8$
$10^7$
$10^6$
$10^5$
$10^4$
WT
ΔvxrAB+Bip
ΔvxrAB
ΔvxrAB+Bip+$Fe^{2+}$
0
1
2
3
PenG Exposure (hour)

Figure S7

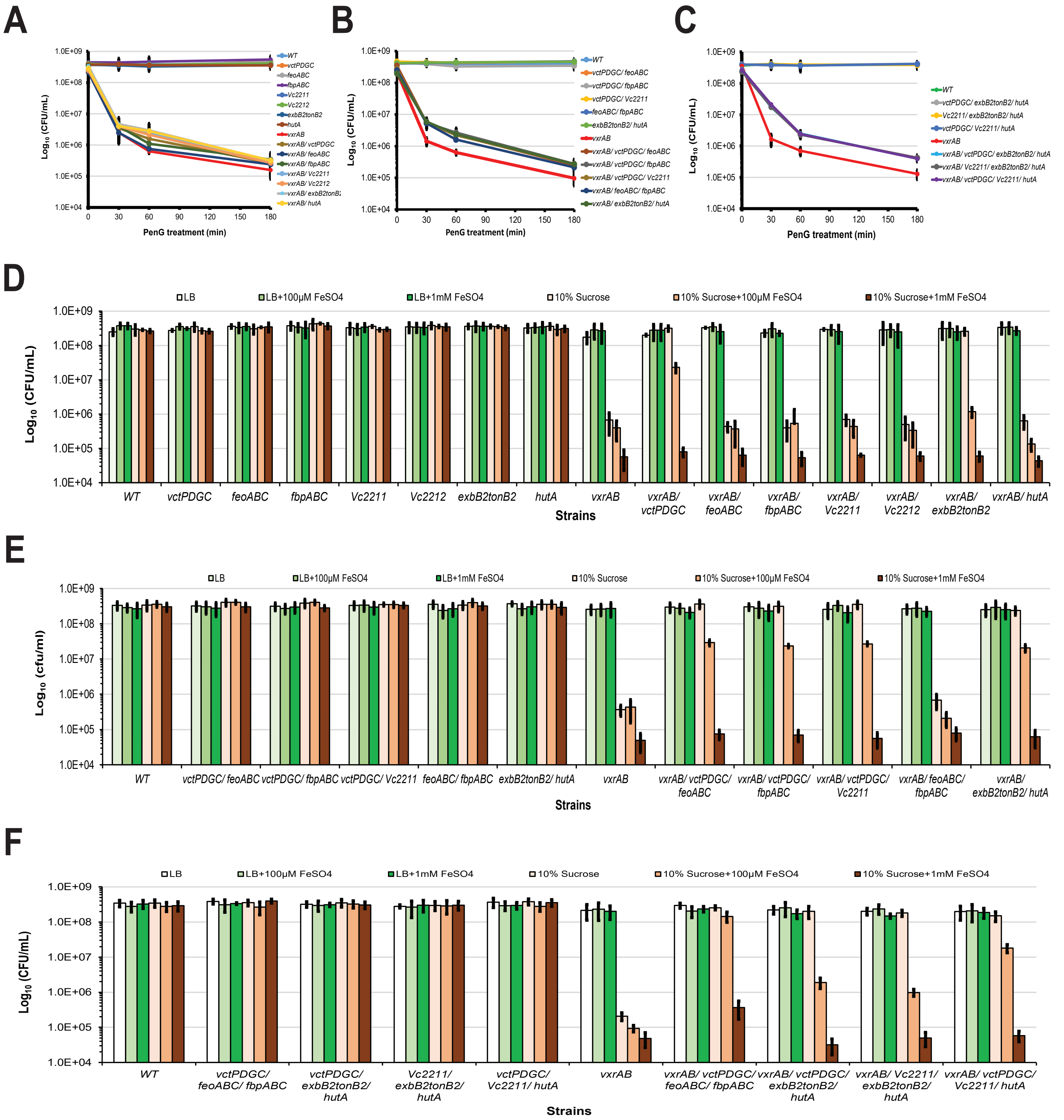
A
B
C
D
E
F
Log10 (CFU/mL)
Log10 (cfu/ml)
PenG treatment (min)
0
30
60
90
120
150
180
1.0E+04
1.0E+05
1.0E+06
1.0E+07
1.0E+08
1.0E+09
WT
vctPDGC
feoABC
fbpABC
Vc2211
Vc2212
exbB2tonB2
hutA
vxrAB
vxrAB/ vctPDGC
vxrAB/ feoABC
vxrAB/ fbpABC
vxrAB/ Vc2211
vxrAB/ Vc2212
vxrAB/ hutA
vctPDGC/ feoABC
vctPDGC/ fbpABC
vctPDGC/ Vc2211
feoABC/ fbpABC
exbB2tonB2/ hutA
vxrAB/ vctPDGC/ feoABC
vxrAB/ vctPDGC/ fbpABC
vxrAB/ vctPDGC/ Vc2211
vxrAB/ feoABC/ fbpABC
vxrAB/ exbB2tonB2/ hutA
vctPDGC/ exbB2tonB2/ hutA
Vc2211/ exbB2tonB2/ hutA
vctPDGC/ Vc2211/ hutA
vxrAB/ vctPDGC/ exbB2tonB2/ hutA
vxrAB/ Vc2211/ exbB2tonB2/ hutA
vxrAB/ vctPDGC/ Vc2211/ hutA
LB
LB+100μM FeSO4
LB+1mM FeSO4
10% Sucrose
10% Sucrose+100μM FeSO4
10% Sucrose+1mM FeSO4
Strains
vxrAB/ exbB2tonB2
vctPDGC/ feoABC/ fbpABC
vctPDGC/ exbB2tonB2/ hutA
Vc2211/ exbB2tonB2/ hutA
vctPDGC/ Vc2211/ hutA
vxrAB/ vctPDGC/ feoABC/ fbpABC
vxrAB/ vctPDGC/ exbB2tonB2/ hutA
vxrAB/ Vc2211/ exbB2tonB2/ hutA
vxrAB/ vctPDGC/ Vc2211/ hutA

Figure S8

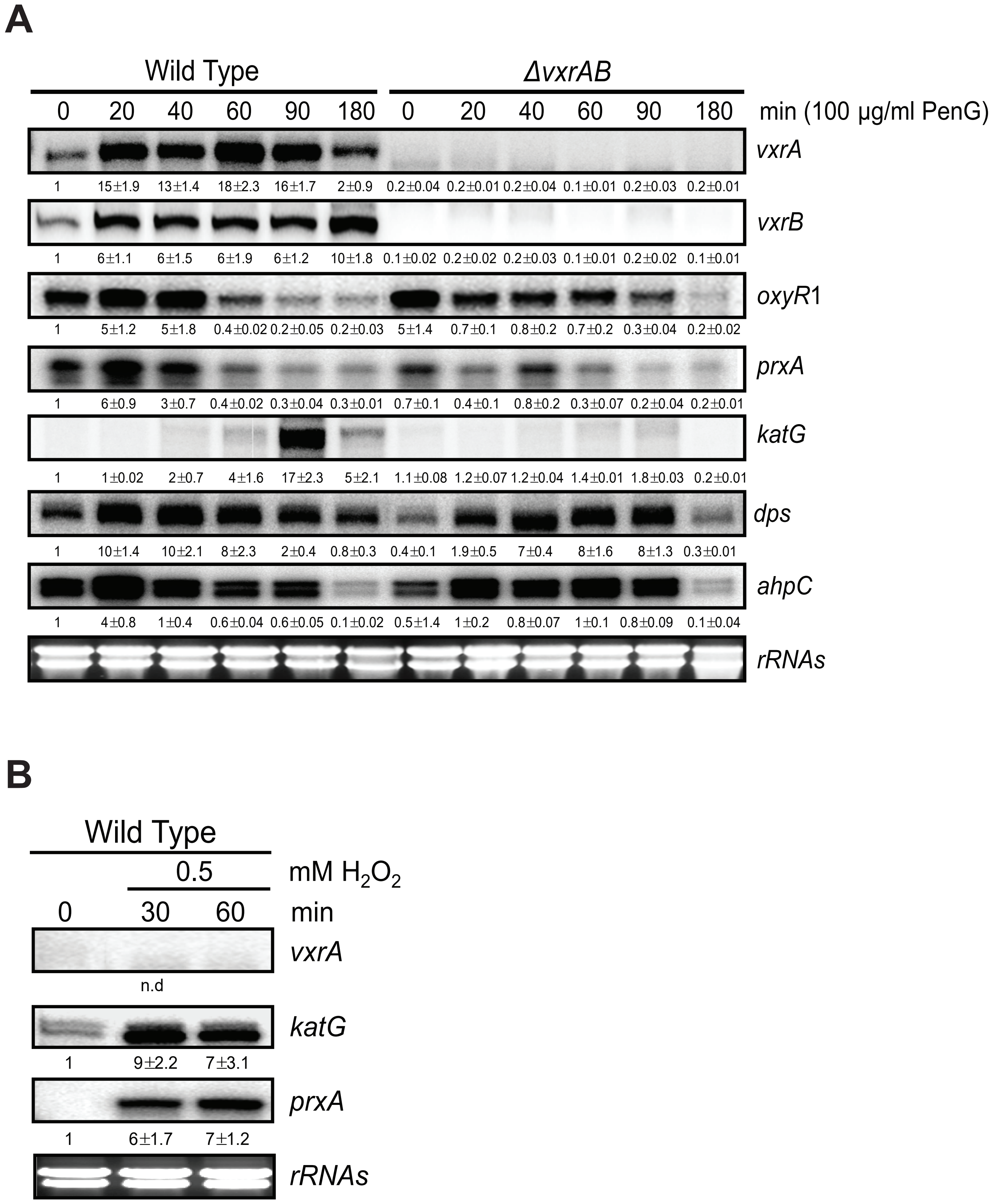
A
Wild Type
ΔvxrAB
0 20 40 60 90 180 0 20 40 60 90 180 min (100 µg/ml PenG)
vxrA
1 15±1.9 13±1.4 18±2.3 16±1.7 2±0.9 0.2±0.04 0.2±0.01 0.2±0.04 0.1±0.01 0.2±0.03 0.2±0.01
vxrB
1 6±1.1 6±1.5 6±1.9 6±1.2 10±1.8 0.1±0.02 0.2±0.02 0.2±0.03 0.1±0.01 0.2±0.02 0.1±0.01
oxyR1
1 5±1.2 5±1.8 0.4±0.02 0.2±0.05 0.2±0.03 5±1.4 0.7±0.1 0.8±0.2 0.7±0.2 0.3±0.04 0.2±0.02
prxA
1 6±0.9 3±0.7 0.4±0.02 0.3±0.04 0.3±0.01 0.7±0.1 0.4±0.1 0.8±0.2 0.3±0.07 0.2±0.04 0.2±0.01
katG
1 1±0.02 2±0.7 4±1.6 17±2.3 5±2.1 1.1±0.08 1.2±0.07 1.2±0.04 1.4±0.01 1.8±0.03 0.2±0.01
dps
1 10±1.4 10±2.1 8±2.3 2±0.4 0.8±0.3 0.4±0.1 1.9±0.5 7±0.4 8±1.6 8±1.3 0.3±0.01
ahpC
1 4±0.8 1±0.4 0.6±0.04 0.6±0.05 0.1±0.02 0.5±1.4 1±0.2 0.8±0.07 1±0.1 0.8±0.09 0.1±0.04
rRNAs
B
Wild Type
0.5 mM H2O2
0 30 60 min
vxrA
n.d
katG
1 9±2.2 7±3.1
prxA
1 6±1.7 7±1.2
rRNAs

### Figure S9

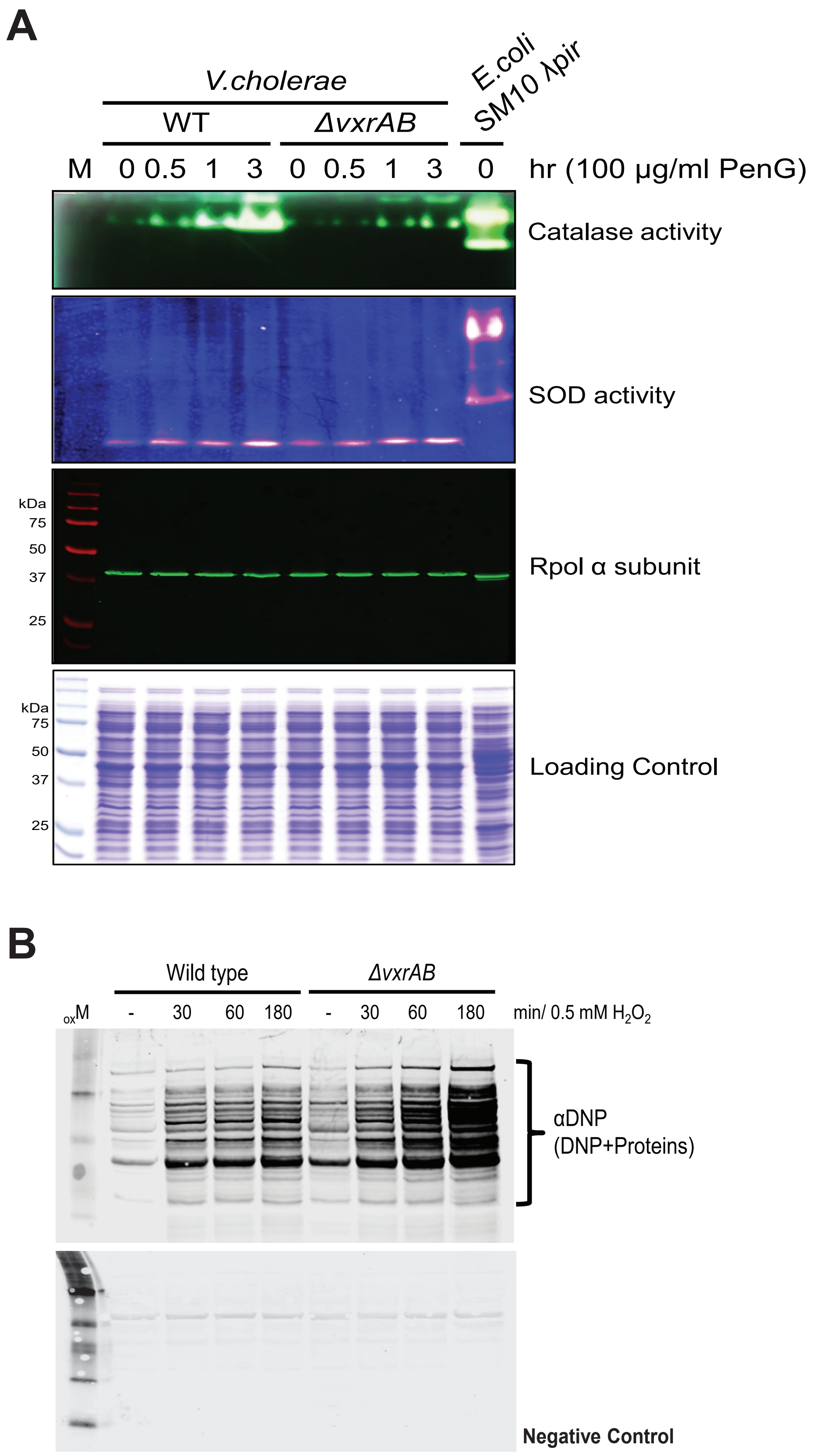
A
V.cholerae
WT
ΔvxrAB
E.coli SM10 λpir
M 0 0.5 1 3 0 0.5 1 3 0 hr (100 µg/ml PenG)
Catalase activity
SOD activity
kDa
75
50
37
25
Rpol α subunit
Loading Control
B
Wild type
ΔvxrAB
oxM - 30 60 180 - 30 60 180 min/ 0.5 mM H2O2
αDNP
(DNP+Proteins)
Negative Control

### Figure S10

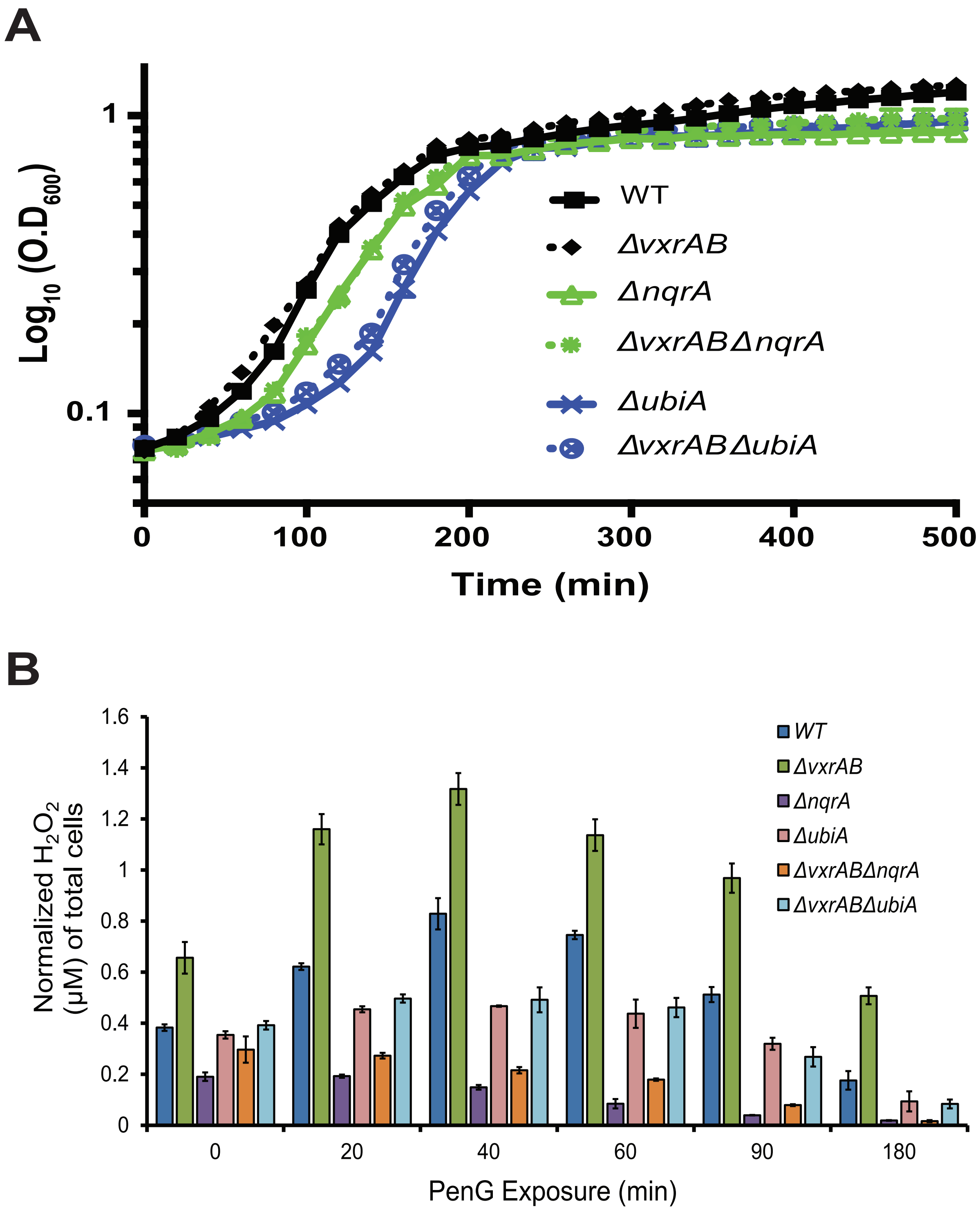
A
1
0.1
Log10 (O.D600)
0
100
200
300
400
500
Time (min)
WT
ΔvxrAB
ΔnqrA
ΔvxrABΔnqrA
ΔubiA
ΔvxrABΔubiA
B
1.6
1.4
1.2
1
0.8
0.6
0.4
0.2
0
Normalized H2O2 (μM) of total cells
0
20
40
60
90
180
PenG Exposure (min)
WT
ΔvxrAB
ΔnqrA
ΔubiA
ΔvxrABΔnqrA
ΔvxrABΔubiA
